# Conformational Changes of Surfactant Protein B Due to the Alveolar Air/Liquid Interface Using Molecular Dynamics

**DOI:** 10.1101/2025.04.06.647458

**Authors:** Tyler Locke, Amanda Ferrante, Deng Li, Mona Minkara

## Abstract

Surfactant Protein B (SP-B) is a critically important component of pulmonary surfactant (PS), responsible for much of the lipid restructuring activity necessary to maintain proper respiratory function. Despite its functional and biological importance, there is a significant lack of knowledge regarding the structural characteristics of SP-B, exacerbated by a lack of a complete experimentally determined structure. Comparative modeling with homologous Saposin-family proteins was used to generate predicted structures for SP-B in both an open (hydrophobic residues exposed) and closed (hydrophobic residues buried) conformation. These structures were then used for further study with Molecular Dynamics. Five replicate simulation systems were prepared for both conformations in different solvent conditions, including water and chloroform, a hydrophobic solvent. For each system, a minimum of 900 ns production time per simulation was collected in 5 replicate simulations. Overall RMSD, per-residue RMSF, specific geometric parameters, and solvent distribution information were collected over the course of the simulations and analyzed. Results of these analyses indicate the relative stability of the closed conformation protein in water, with the open conformation structure undergoing a large conformational change due to hydrophobic forces in water quantified by relevant intramolecular distances. Solvent distribution analysis elucidated the varying affinity of different regions of the protein to hydrophobic and hydrophilic environments, providing insight into the structural-functional characteristics of SP-B in the varied PS environment. Helix 3 was identified as a region of particular interest, demonstrating structure-dependent increased affinity to hydrophobic solvent molecules.

## Introduction

Pulmonary surfactant (PS) is a complex, dynamic system composed of lipids and proteins that exists at the air-liquid interface in the alveoli of the lungs, as shown in figure 1. PS is a vital functional component of the respiratory system, reducing the work of breathing and preventing alveolar collapse by reducing surface tension at the interface. Additionally, as the first point of contact for airborne pathogens, PS plays a key role in the body’s innate immune response. PS surface tension modulation occurs through the spreading of surfactant lipids in a monolayer at the air-liquid interface. Surface tension regulation over the wide range of areas and pressures that occur during breathing requires that PS undergo rapid adsorption, compression, and respreading on a constantly expanding and contracting surface (1). The understanding of this system is of clinical significance, as numerous pulmonary disease states are associated with insufficient or non-functioning surfactant. Neonatal respiratory distress syndrome (NRDS), acute respiratory distress syndrome, and pulmonary alveolar proteinosis are all serious medical conditions associated with measurable effects on surfactant composition and functional characteristics (2). In addition to these established relationships between PS function and disease, recent research has suggested that PS may be implicated in numerous other pulmonary diseases. Changes in PS quantity, composition, and function were observed in lung infections, chronic lung diseases such as asthma, and in smoking-associated lung damage (3). More recently, experiments and clinical observations have demonstrated changes in PS function in severe cases of COVID-19, potentially contributing to poor clinical outcomes (4,5).

**Figure 1:**
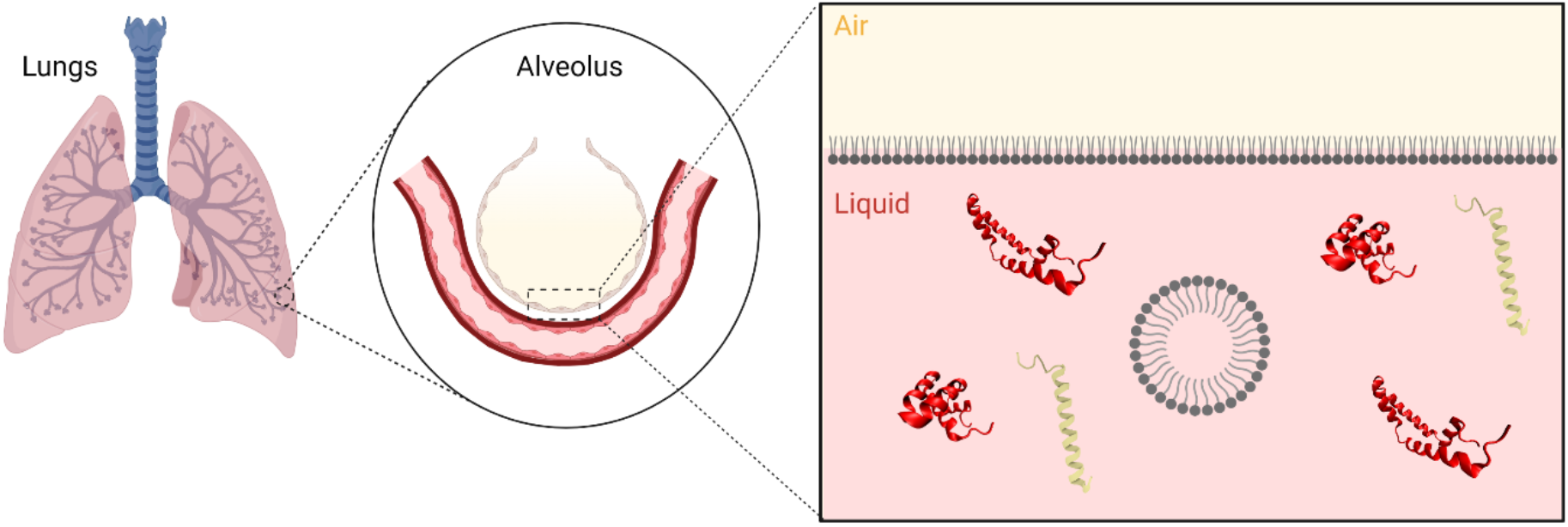
Cartoon detailing the location of hydrophobic surfactant protein components SP-B (red ribbon structure) and SP-C (yellow-green ribbon structure) in surfactant lipid (grey structures) complex at the alveolar air-liquid interface.

Healthy PS activity is dependent on the complex interactions between the lipid and protein components. Although the PS lipid monolayer is the primary force in reducing surface tension at the air-liquid interface, lipids alone are insufficient for maintaining surfactant function (6), and lipid-only surfactant replacement therapies have been shown to be less effective than their protein-containing counterparts (7). The four surfactant proteins (SP) are denoted SP-A, SP-B, SP-C, and SP-D and can be classified as either hydrophobic or hydrophilic surfactant proteins. The oligomeric SP-A and SP-D are much larger, forming complex oligomeric structures, and function primarily in the body’s innate immune response to airborne pathogens. The hydrophobic proteins SP-B and SP-C are smaller, lipid-associated proteins, and are involved primarily in modulating lipid restructuring at the interface (8). In vivo and in vitro models of environmental-associated lung disease have shown significant changes in the expression of surfactant proteins in the alveolar epithelial cells (9). SP-B is critically important to the function of PS, with innate SP-B deficiency in humans leading to death within the first year of life (10). This finding was confirmed in mice, where knockout of the gene encoding SP-B caused lethal respiratory failure (11). Recent research has shown that higher levels of synthetic SP-B analogues in replacement PS improve surfactant function in vitro and in vivo, highlighting the importance of functional SP-B mimics in developing surfactant replacement therapies (12).

SP-B is a 79-residue, primarily hydrophobic protein of the saposin-like protein (SAPLIP) superfamily. The SAPLIP proteins have a variety of functions, including membrane lysis, pore-formation, and other forms of lipid structural modulation in both healthy physiology and pathological response. Saposins A, B, C, and D catalyze physiologically important lipid catabolism and transfer activities, while NK-Lysin and Granulysin perform key antimicrobial membrane lysis functions in the innate immune system. Generally, the saposins are a family of lipid-associated proteins characterized by a motif of 6 conserved cysteine residues that form 3 intramolecular disulfide bonds. Conserved disulfide bonds create the signature “saposin fold” in which the N-terminal and C-Terminal regions of the protein are in close proximity in a U-shaped secondary structure, as shown in figure 2. The secondary structure of SAPLIP proteins is semi-conserved, generally forming 4 distinct helical regions, which are connected to one-another through disulfide bonds, forming the characteristic saposin fold (13). As a SAPLIP protein, SP-B is unique in that it is water-insoluble and contains a 7^th^ cysteine residue involved in the formation of a covalently linked homodimer. It is unknown whether this dimerization is critical to SP-B function, and modified SP-B has been shown to form dimers at high concentration even in the absence of the 7th cysteine. At pH=7.4, SP-B has 9 positively charged and 2 negatively charged amino acids resulting in a net +7 charge. This suggests that SP-B may interact preferentially with anionic surfactant lipids such as POPG (14). Electron microscopy and atomic force microscopy experiments have proposed the formation of higher order multimers, possibly forming larger ring-like structures which facilitate lipid transport (15).

**Figure 2:**
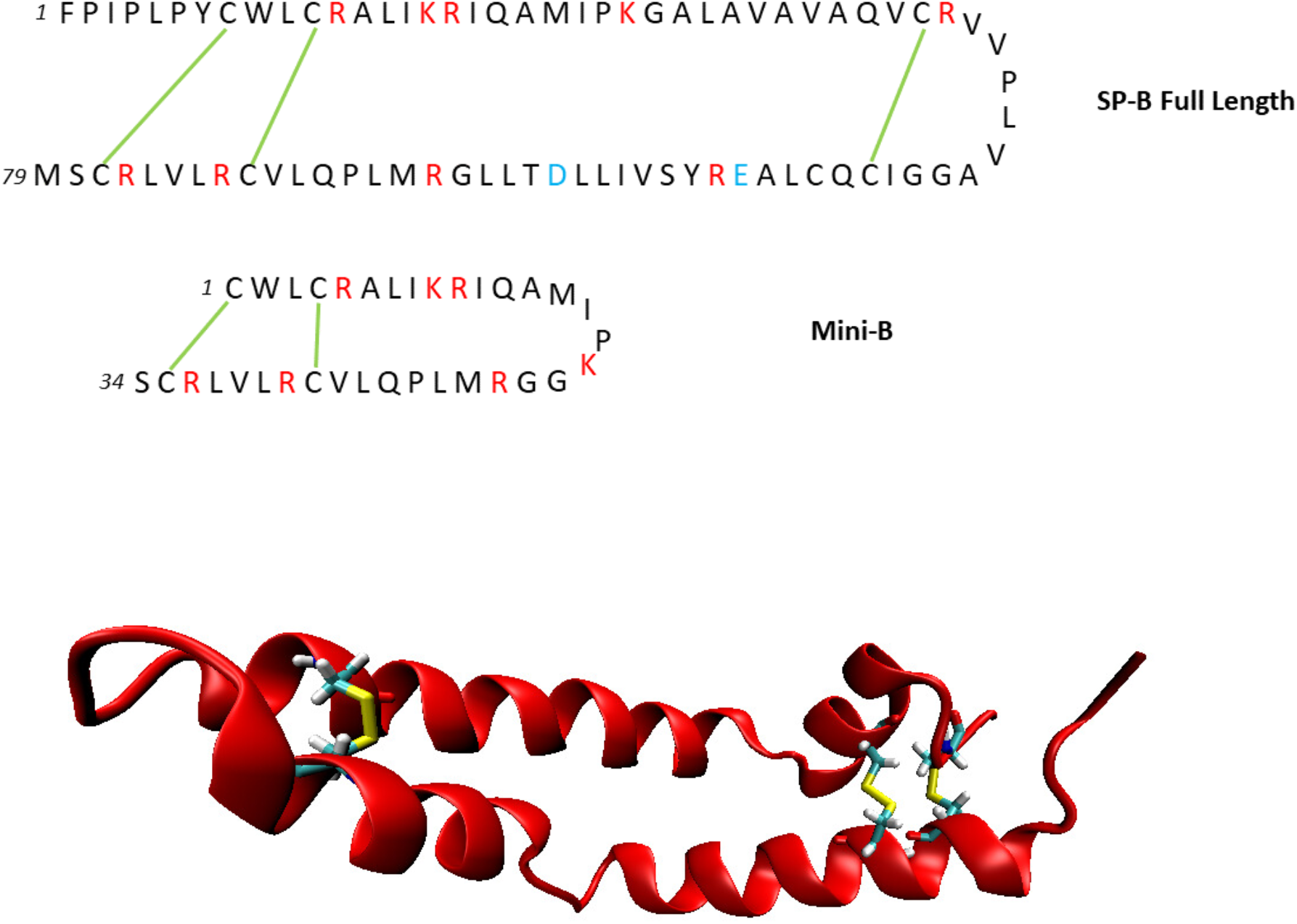
Top: Amino acid sequences of SP-B and the 25-residue fragment structure Mini-B. Disulfide bonds are shown as green connections. Positively charged residues are shown in red, and negatively charged residues are shown in blue. Bottom: Hypothesized structure of SP-B, with disulfide bonds highlighted.

Despite its functional importance, the structural and dynamic properties of SP-B are not well understood. Difficulty with obtaining isolated SP-B in forms suitable for structural analysis has prevented experimental determination of the 3D structure of SP-B. Structural determination efforts have been successful with synthetic fragments of SP-B, providing the structure of several fragments of SP-B, including the 34-residue functional fragment Mini-B, consisting of the N- and C-terminal helices of SP-B (16). The amino acid sequences of SP-B and Mini-B are compared in figure 2. Mini-B contains 7 of the positively charged amino acids and neither of the negatively charged amino acids of SP-B, maintaining its net +7 charge at pH 7.4. Experimental studies examining bulk PS structure and function have provided limited insight into the functional characteristics of SP-B. X-ray diffuse scattering experiments have suggested a superficial location within a lipid bilayer, suggesting SP-B may be able to interact with multiple lipid structures simultaneously. However, these findings were based off a static bilayer system and may not be fully representative of conditions in vivo (17). Experimentation with different surfactant systems has revealed the formation of complex lipid structures during respiration that allow stable compression and respreading of lipids at the interface. Due to its presence at the interface and its lipophilic character, SP-B is thought to play a key role in the formation and stabilization of these intermediate structures (8, 14). Recent studies have demonstrated significant structural effects in surfactant systems correlated with the presence of hydrophobic surfactant proteins but have yet to find definitive evidence of the specific molecular mechanisms responsible for that function (18,19).

Circular dichroism and Fourier transform infrared spectroscopy experiments suggest that SP-B has an overall helicity of 40-50% (15), largely independent of the solvent environment. The dynamic nature of the PS system makes high-resolution investigation of SP-B’s structure and interactions in PS prohibitively difficult. Specific details on the protein-protein and protein-lipid interaction mechanisms which may be responsible for SP-B’s function are currently unknown. Computational study provides an alternative pathway to investigate these important details and further our understanding of the molecular-scale function of SP-B.

The lack of an experimentally determined structure for SP-B presents a significant challenge for computational exploration of its structural and functional properties. Previous molecular dynamics studies have made use of available fragment structures for SP-B such as the 25-residue N-terminal helix region (18) or Mini-B (23). These structures have allowed researchers to examine how SP-B may interact with different surfactant components, but lacks insight into the full-protein scale interactions that may be crucial to the function of SP-B. Full-length SP-B models have been developed for this purpose based on the known homology of SP-B with the SAPLIP family of proteins as well as experimentally determined fragment structures (24). These studies have allowed researchers to perform molecular dynamics simulations involving full-length SP-B peptides in a variety of PS-like environments. These include simulations of SP-B in different locations within a lipid multilayer(25, 26), as well as SP-B in the presence of surfactant lipid vesicles (27).

Previous approaches to developing full-sequence models for SP-B have used fragment structures such as Mini-B as templates for secondary structures, building on the additional residues using the secondary structure of SAPLIP proteins such as NK-Lysin. Using this approach, researchers can explicitly define tertiary structures with different geometries (25, 26, 27). However, the diversity seen in tertiary structures of different saposin proteins involved in different types of lipid interaction and function suggests that the choice of a proper full-length starting structure may be key to understanding the functional characteristics of SP-B (13). Additionally, simulations by Khatami et al. identified a strong dependence on initial structural conditions when simulating SP-B in complex with lipid structures (25).

The initial orientation of helices in the SP-B molecule and the placement of the molecule relative to the lipid bilayer center both have major influence on subsequent structural changes in MD. These results emphasize the importance of determining native-like structural models of SP-B in the alveolar air-liquid environment. In this study, a hybridized comparative modeling approach was taken, using multiple SAPLIP template structures in different conformations. This modeling strategy makes use of the large amount of structural data available for the saposin proteins, while still allowing for structural flexibility to incorporate important differences between SP-B and the template proteins. Templates were segregated into “closed” and “open” conformations based on geometric factors, allowing for the creation of two distinct conformational models for the full-length SP-B. These models can then be used to examine the relative stability of different conformations of SP-B, as well as any large-scale conformational changes that may occur in response to environmental conditions.

Molecular dynamics studies have previously been conducted examining the interaction of different SP-B models with different systems of surfactant components(25, 26, 27). One major difficulty with this approach is the lack of concrete information on the location and orientation of SP-B within the surfactant monolayer. This, combined with the fact that membrane dynamics are sensitive to environmental changes in surface tension, makes it difficult to construct accurate systems and draw conclusions in vivo. Preparation of multiple initial system geometries is common and has demonstrated that protein behavior is sensitive to initial membrane orientation (25). The aim of this study is to further elucidate the process of SP-B association with lipids through analysis of SP-B structure and dynamics in various environments though molecular dynamics simulation of open and closed conformation SP-B in hydrophobic and hydrophilic solvents. Full-length SP-B structural models were created using a hybridized comparative modeling approach to fully predict the distinct tertiary conformations that may occur in the native protein environment. Multiple conformations of SP-B were simulated in distinct explicit hydrophilic and hydrophobic environments to examine potential differences in solvent interaction. This approach allows a large amount of data to be collected, with minimal complication from larger, more complex environments. Analysis of these simulation data has revealed previously unknown details on the relationship between SP-B’s conformational flexibility and its solvent interactions. Determining these interaction details is a key step toward performing more accurate and informed investigation of SP-B-Lipid interactions in PS function. This study combines comparative modeling and molecular dynamics simulation of SP-B in multiple solvents to investigate structural changes and solvent interactions which occur due to the PS environment.

## Methods

### Structure Generation and Preparation

We used the Monte-Carlo based RosettaCM (28) comparative modeling (CM) method to generate starting structures for simulation. Two separate CM runs were performed to reflect the multiple distinct conformations present in other saposin family proteins (29). The first run utilized a wide selection of saposin protein template structures, as well as the fragment structure Mini-B, which consists of the N- and C-terminal domains of SP-B (30). The full list of template structures used for comparative modeling is shown in Table 1, along with their respective RCSB PDB ID and classification as open or closed conformation. A second run was performed using only the open conformation saposin template structures. Template structures used in this open model were open conformation saposin A (PDB: 4DDJ 37), saposin B (PDB: 1N69 32), and saposin C (PDB: 1SN6 34). Both open and closed conformation structures were used in the closed conformation CM run, as this resulted in only closed conformation predicted structures. This closed conformation bias is likely because the closed conformation is more energetically favorable in water. The amino acid sequence for the full SP-B preprotein was downloaded from the NCBI Protein database (38) and residues 201-279 were selected, as these comprise the final SP-B peptide (39). To align the template structures with the mature SP-B sequence, the Clustal Omega multiple sequence alignment tool (40) was used. Alignments were verified by confirming the positions of the six conserved cysteine residues were in the correct position across each sequence. Three disulfide bonds were defined between residues Cys8-Cys77, Cys11-Cys71, and Cys35-Cys46 based on known structural motifs in the saposin protein family (39). Using these parameters, 15,000 structural models were generated. The representative structures for each CM run were chosen using the Calibur clustering method to determine the structures with the highest number of neighbors under a chosen RMSD threshold (41). Following clustering, all structural models were sorted by RMSD to the representative structure and the top 4 closest structures were selected for use in replicate simulations. Examples of the open and closed conformation structures used for simulation are shown in figure 2. To prepare the structural models for simulation, the H++ online protein protonation server (http://biophysics.cs.vt.edu/H++ 42,43,44) was used to generate Amber-compatible protonation states at pH 7.4.

**Table 1:** Template structures used for comparative modeling. Only structures classified as open were used in the open conformation run, while all structures were used in the closed conformation run.

| Template Protein | Classification | RCSB PDB ID | Reference |
| --- | --- | --- | --- |
| Granulysin | Closed | 1L9L | 31 |
| Saposin B | Open | 1N69 | 32 |
| NK-Lysin | Closed | 1NKL | 33 |
| Saposin C | Open | 1SN6 | 34 |
|  | Closed | 2GTG | 35 |
| Mini-B | N/A | 2DWF | 30 |
| Saposin D | Closed | 3BQQ | 36 |
| Saposin A | Open | 2DOB | 35 |
|  | Closed | 4DDJ | 27 |

### System Building

For each conformation of SP-B, 5 replicate simulation systems were prepared in both water and the hydrophobic solvent chloroform as a model hydrophobic polar liquid. Simulation systems with open conformation SP-B solvated by water were prepared in tLeap by first neutralizing the prepared structures with 7 Cl^—^ions placed by tLeap, then solvating in a 10 Å periodically replicated octahedral water box using the OPC water model (45). Na^+^ and Cl^-^ ions were added by tLeap to a NaCl concentration of 150 mM. Closed conformation systems were prepared similarly with a 14 Å periodically replicated octahedral water box to allow for potential expansion of the smaller peptide conformation. A minimum of 6,975 and a maximum of 8,361 water molecules were used to hydrate the open structures, while a minimum of 7,912 and a maximum of 10,696 water molecules were used to hydrate the closed structures. Variability in the geometry of the peptide starting structures resulted in different box sizes, reflected by the range of solvent molecule counts in replicate simulations. Simulation systems with open conformation SP-B solvated by chloroform were prepared by solvating the prepared structures in a 12 Å periodically replicated octahedral box using the AMBER solvents library model parameters. Similarly, a 17 Å periodically replicated octahedral box was used for the closed conformation simulation to allow for potential peptide expansion. Counterions were not added to these systems, as NaCl is not soluble in chloroform, and compatible ion parameters are therefore not available. A minimum of 1821 and a maximum of 1997 chloroform molecules were used to hydrate the open structures, while a minimum of 1880 and a maximum of 2144 chloroform molecules were used for the closed conformation.

### Minimization, Heating, and Production

Each simulation system was subject to the same molecular dynamics protocol using the GPU version of Amber20 and the FF19SB protein force field. First, the prepared systems were subject to a 2-stage energy minimization consisting of 1000 steps steepest descent minimization followed by 1000 steps conjugate gradient descent. The systems were then heated from 0 to 310 K over 10,000 steps with a step size of 0.002 ps using the Langevin thermostat (46) with a collision frequency of 2 ps^-1^. Production simulations were then run in the isobaric, isothermal (NPT) ensemble using the Berendsen barostat (47) for pressure regulation and Langevin thermostat for temperature regulation. During heating and production simulations, the SHAKE (48) algorithm was used to constrain all hydrogen-containing bonds. A non-bonded interaction cutoff distance of 8 Å was used for all simulation stages. Simulations were run in production for a minimum of 900 ns after the protein RMSD was determined to have reached a quasi-stable value. The 5 simulations for each condition with equilibration and production simulation times are shown in table 2.

**Table 2:** Equilibration and production time of each simulation.

| SP-B Conformation | Solvent | Replicate | Time to Equilibration (ns) | Production Time (ns) |
| --- | --- | --- | --- | --- |
| Open | Water | 1 | 23 | 1189 |
|  |  | 2 | 125 | 1087 |
|  |  | 3 | 80 | 1132 |
|  |  | 4 | 190 | 1022 |
|  |  | 5 | 107 | 1115 |
|  | Chloroform | 1 | 22 | 1490 |
|  |  | 2 | 34 | 1478 |
|  |  | 3 | 21 | 1491 |
|  |  | 4 | 500 | 1012 |
|  |  | 5 | 215 | 1297 |
| Closed | Water | 1 | 147 | 1065 |
|  |  | 2 | 88 | 1124 |
|  |  | 3 | 76 | 1136 |
|  |  | 4 | 105 | 1107 |
|  |  | 5 | 264 | 948 |
|  | Chloroform | 1 | 54 | 1458 |
|  |  | 2 | 31 | 1481 |
|  |  | 3 | 45 | 1469 |
|  |  | 4 | 20 | 1492 |
|  |  | 5 | 64 | 1448 |

### Analysis

Root mean square deviation (RMSD) was calculated for each simulation using the minimized structure as a reference geometry. Per-residue root mean square fluctuation (RMSF) was also calculated using the coordinates of the protein backbone atoms. To characterize the large-scale conformational differences in the protein, the representative residue separations TYR7-VAL34 and ILE45-VAL70 (figure 5) were calculated and analyzed. These residues were chosen for their location within opposing helices in the full-length SP-B structure. For comparison with other saposin proteins, the residue separation CYS8-CYS35 was also recorded, as these residues are conserved across all saposin proteins. To examine the interaction of solvent molecules with different regions of the protein, the radial distribution function of each solvent was calculated around several regions of interest in the protein. In this computation, the density of solvent was calculated in binned spherical shells about the regions center of mass as calculated by CPPtraj and normalized according to the volume of the simulation box. This allowed for direct comparison of density profiles against solvents with different solution densities.

### Free Energy Heatmaps

To further examine the different stable conformations of SP-B, free energy heatmaps were constructed using the methodology described in Minkara et al. (49) and Toba et al. (50). First, the produced frames were grouped into states consisting of combinations of the representative residue separations discussed previously. Distinct states were created by binning each parameter with a bin width of 0.1 Å. The probability of each state was determined by summing the number of frames in each state by the total number of frames across all replicates. The relative free energy is determined by Eq. 1, where R is the ideal gas constant, T is the simulation temperature, and P_i_ is the probability of an individual state occurring in a randomly selected frame.

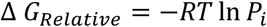

Data were collected using an in-house script. Heatmaps were generated using the final 500 ns of simulation, ensuring any equilibration was discarded.

## Results

### Quantification of Open/Closed Conformations in Different Environments

Based on structural/functional information available for other proteins, SP-B is hypothesized to exist in multiple conformations, characterized as “open” or “closed”, with the open conformation exposing hydrophobic residues for interaction with lipid complexes, and the closed conformation burying hydrophobic residues for better stability in water. SP-B likely retains the 4-helix secondary structure observed in other Saposin proteins, and it is hypothesized that this conformational flexibility is made possible by flexible “hinge” regions in the amino acids separating helices 1&2 and helices 3&4. Since the N-and C-terminal domains of the protein are connected by disulfide bonds, these hinges divide the molecule into two distinct flaps consisting of helices 1&4 and helices 2&3. The bulk motion of these two flaps relative to one another results in distinct open and closed conformations. Analysis of the MD simulation data was performed to characterize the impact solvent hydrophobicity has on the open/closed conformations of SP-B.

The RMSD (Figure 4, SI Figure 1) was obtained for the entire protein structure in each simulation, using the first frame after minimization as a reference point. Simulations were run for a minimum 900 ns after the RMSD was determined to have reached a stable value. The RMSD of the open conformation SP-B molecules solvated by water (“Open-water” simulations) rapidly increased to an average value of 8 angstroms in the first 200 ns of the simulation, after which they slowly increased to equilibrium values between 9 and 12 Å. For the open conformation SP-B solvated in chloroform (“Open-chloroform” simulations), the RMSD quickly reaches an average value of 8 Å in the first 50 ns of simulation, with final equilibrium values between 9 and 12 Å. For the closed conformation of SP-B solvated by water (“Closed-water” simulations), the RMSD levels off at an average value of 4 Å in the first 50 ns of simulation, with final equilibrium values between 3 and 6 Å. For the closed conformation SP-B solvated in chloroform (“Closed-Chloroform” simulations), the RMSD levels off at an average value of 4 Å in the first 50 ns of simulation and slowly increases to final equilibrium values between 5 and 8 Å. Consistently lower RMSD values seen in the closed conformation simulations suggest that there is less deviation from the initial CM structure in this conformation.

**Figure 3:**
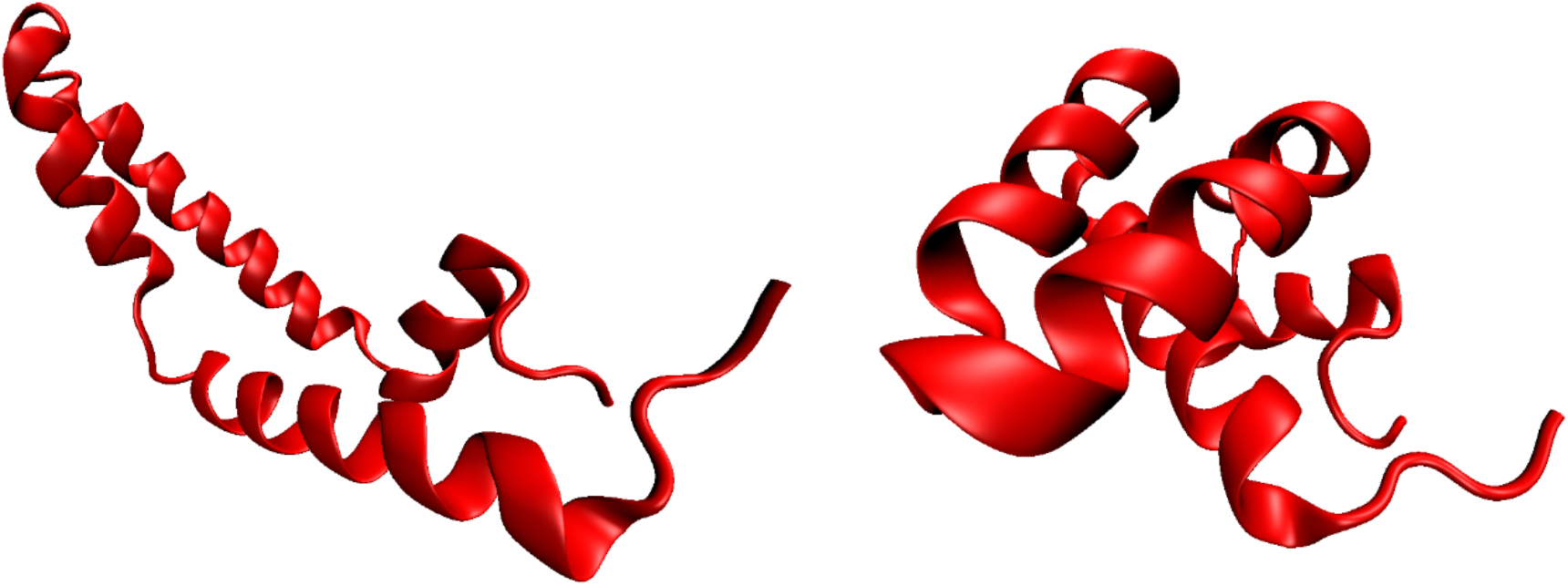
Open (left) and closed (right) conformations of Surfactant Protein B. Final structures from comparative modeling selected for simulation.

**Figure 4:**
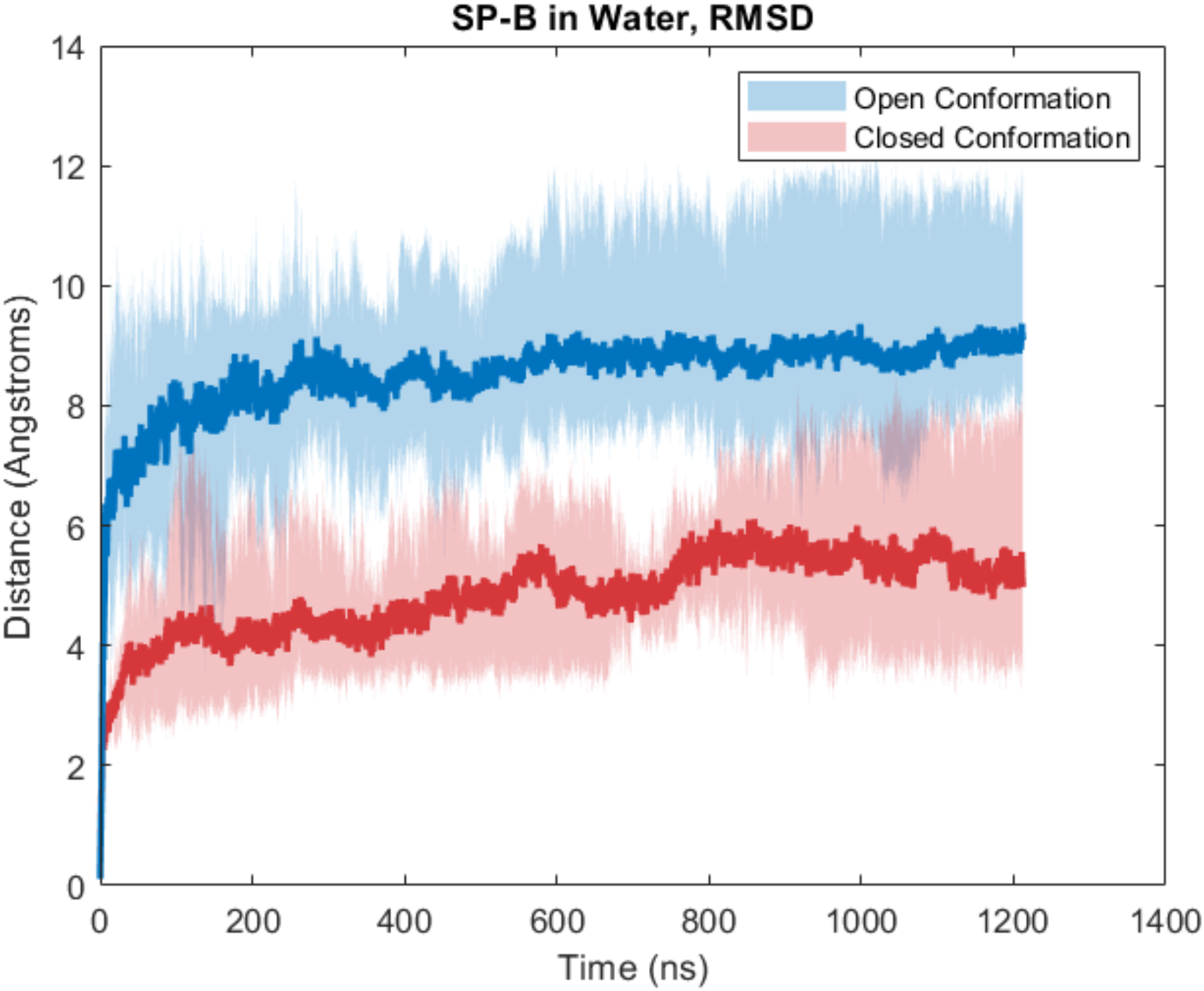
Peptide backbone RMSD of SP-B/water systems. Initial open conformation systems shown in blue, initial closed conformation in red. Dark lines represent the mean trajectory over all replicates, while shaded areas show the minimum and maximum values at a given time point. RMSD data for chloroform systems is included in supplemental information.

### TYR7/VAL34 and ILE45-VAL70 Separation in Water and Chloroform

To quantify the difference between the open and closed conformations, 2 pairs of residues were chosen, spanning the two flaps of the molecule. The separation between residues TYR7 in helix 1 and VAL34 in helix 2, as well as the separation between ILE45 in helix 3 and VAL70 in helix 4 were recorded over the entirety of each simulation (both residue pairs are indicated in figure 5). Both TYR7-VAL34 separation and ILE45-VAL70 separation are indicative of flap position, with significant differences observed in ensembles of structures produced by RosettaCM. Open conformations generally exhibit TYR7-VAL34 and ILE45-VAL70 separations of 20-40 Å, while closed conformations exhibit separations of 15-17 Å. These distances were monitored over the course of each simulation to determine the effects of solvent interactions on the conformation of the protein.

**Figure 5:**
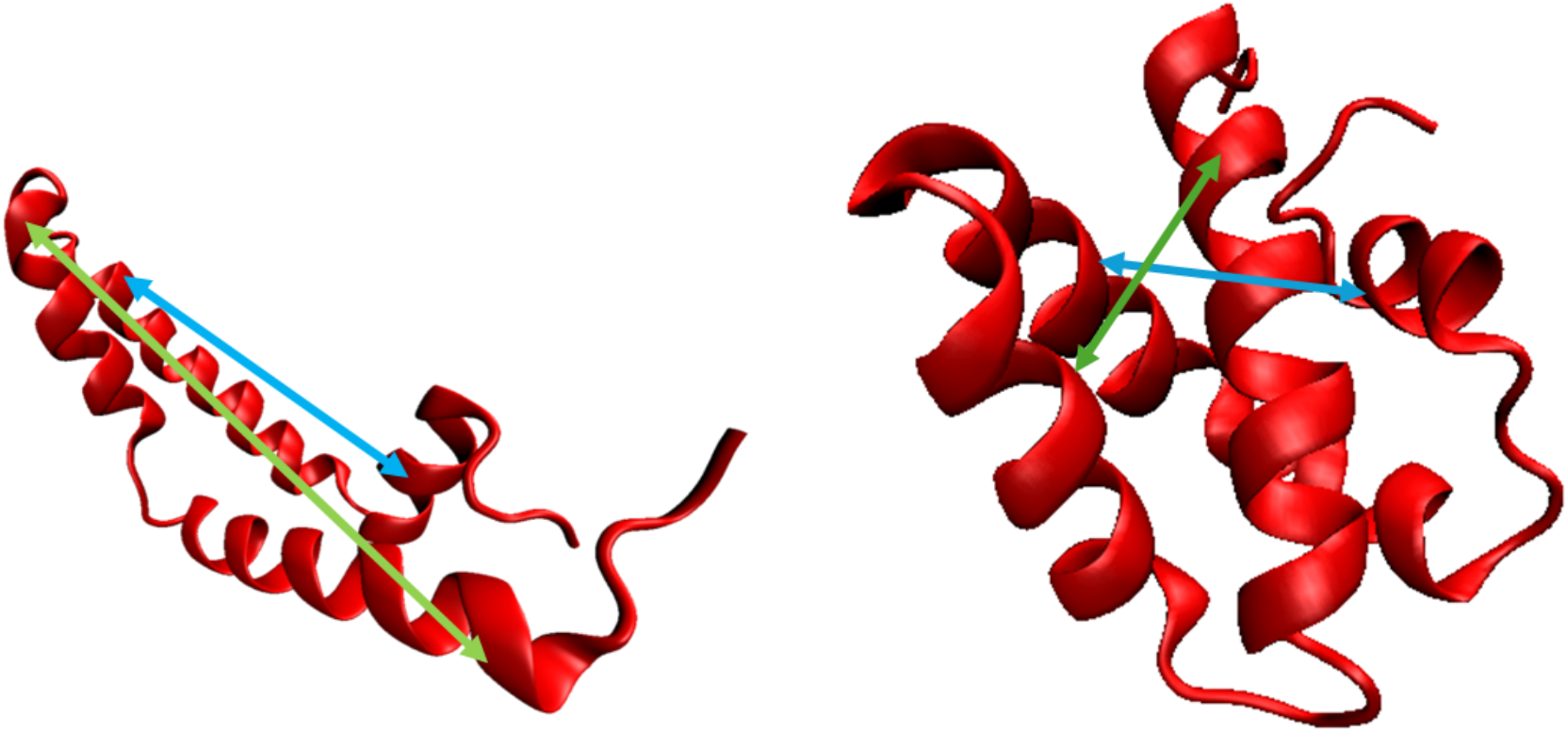
Open conformation (left) and Closed Conformation (right) SP-B structures with TYR7-VAL34 and ILE45-VAL70 distances indicated by green and blue lines, respectively.

The TYR7-VAL34 and ILE45-VAL70 separation data for both SP-B water systems are shown in figure 6. In the closed-water simulations, both separations remained roughly constant across all replicates, fluctuating minimally around equilibrium values between 15 and 20 Å for TYR7-VAL34 and between 8 and 15 Å for ILE45-VAL70. While the final equilibrium structures are all classified as closed conformation, there is some variation in the final residue separations between each replicate, suggesting there are several distinct stable closed conformations in water. In the open-water simulations, the average TYR7-VAL34 separation decreased from an initial value of 25 Å to 17 Å, matching that of the closed-water simulations. The variation among replicates was much higher in the open-water condition, with final equilibrium values between 12 and 30 Å. The average ILE45-VAL70 separation in the open-water system decreased from an initial value of 27 Å to 20 Å, which was significantly higher than the closed-water systems. However, there was significant variation among replicates in this condition, with final equilibrium values between 10 and 35 Å. The significant decrease of mean TYR7-VAL34 and ILE45-VAL70 in the open-water simulation shows that the hydrophilic solvent induces a large-scale conformational change toward the closed conformation. However, the significant variation seen in the equilibrium values of the open-water simulations compared to closed-water suggests that several distinct close conformation structures exist and are accessible from different starting conformations. Additionally, the convergence of the mean TYR7-VAL34 in the open-water and closed-water simulations while ILE45-VAL70 remained higher in the open-water simulations suggests the possibility of a partially open state in which helices 3 and 4 separate while helices 1 and 2 remain in close proximity. These observations are explored further in the conformational free energy heatmaps.

**Figure 6:**
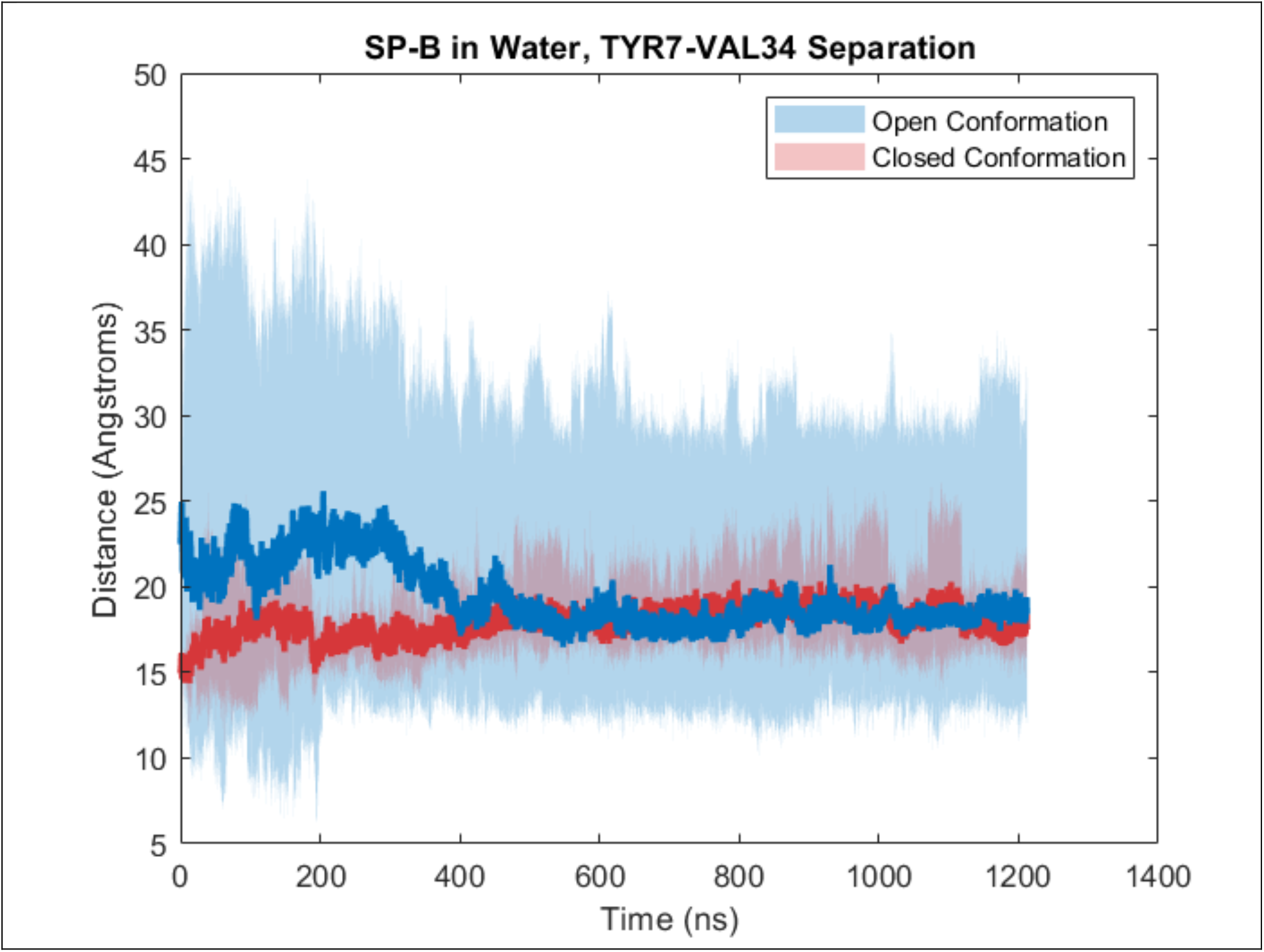
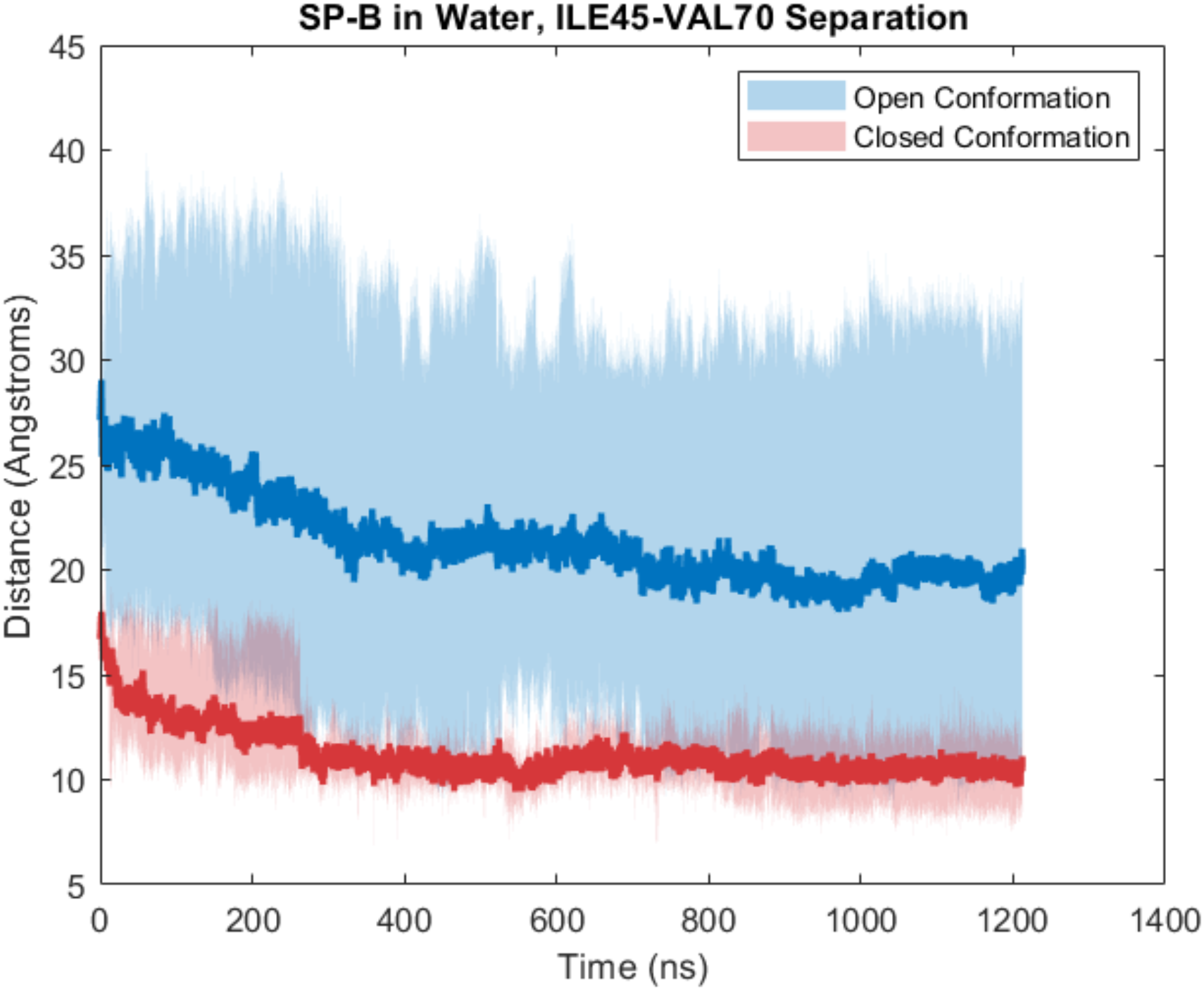
TYR7-VAL34 (top) and ILE45-VAL70 (bottom) separation in water simulations. Initial open conformation systems shown in blue, initial closed conformation in red. Dark lines represent the mean trajectory over all replicates, while shaded areas show the minimum and maximum values at a given time point. Residue separation data for chloroform simulations are included in supplemental information.

Residue separation data for both SP-B chloroform systems are included in the supplemental information. In the open-chloroform simulation, TYR7-VAL34 separations increased rapidly from initial values of approximately 25 Å to approximately 45 Å and fluctuated around that value for the remainder of the simulation, while ILE45-VAL70 similarly increased from approximately 27 Å to 35 Å. This increased residue separation corresponds with a further opening/elongation of the molecule, in contrast with the closure observed in the open-water simulation. In the closed-chloroform simulation, the mean TYR7-VAL34 separation remained roughly equal to its initial value of approximately 17 Å, with replicates varying between 12 and 20 Å. The mean ILE45-VAL70 separation also remained stable between 15 and 20 Å during the simulation with more significant variation between replicates. Final equilibrium ILE45-VAL70 in the closed-chloroform simulation were between 15 and 25 Å. The higher maximum ILE45-VAL70 seen in the closed-chloroform simulation suggests that some closed conformation structures may open partially in a hydrophobic environment, taking on a similar state to those observed in the open-water simulation. It is possible that this may represent a stable transition state between the open and closed conformation, however further opening was not observed from this state in our simulations.

### Conformational Free-Energy Heat Maps

To further investigate the stability of the conformations described previously, free-energy heatmaps were created using TYR7-VAL34 and ILE45-VAL70 separations to parameterize different conformations present in the simulations. Data were collected and analyzed using the previously described process. For each condition, a separate heatmap was generated combining all replicates to compare the different stable conformations in each environment, as shown in figure 7. The open-water simulation heatmap shows several distinct minima in the region of the map classified as closed conformation, as well several weaker minima in the open conformation region, corresponding to the replicate simulations which remained in an open conformation state. The closed-water simulation heatmap shows a more compact region of multiple minima distinct from those seen in the open-water simulation, suggesting that there may be different closed conformation states accessible to SP-B when starting in the open conformation than when starting in the closed conformation. It is likely that the initially closed conformation structures are in a relatively stable minimum and are not perturbed enough to access the other stable closed conformation structures. The open-chloroform simulation shows a more continuous distribution in the open conformation region, showing that this conformation may be more flexible in solution, rather than being confined to discrete states as in the closed conformation. Finally, the closed-chloroform consisted of several strong minima in the closed conformation region, as well as several distinct minima between the closed and open regions. This corresponds to the partial opening seen in some replicate simulations and may represent a stable transition state between the open and closed conformations.

**Figure 7:**
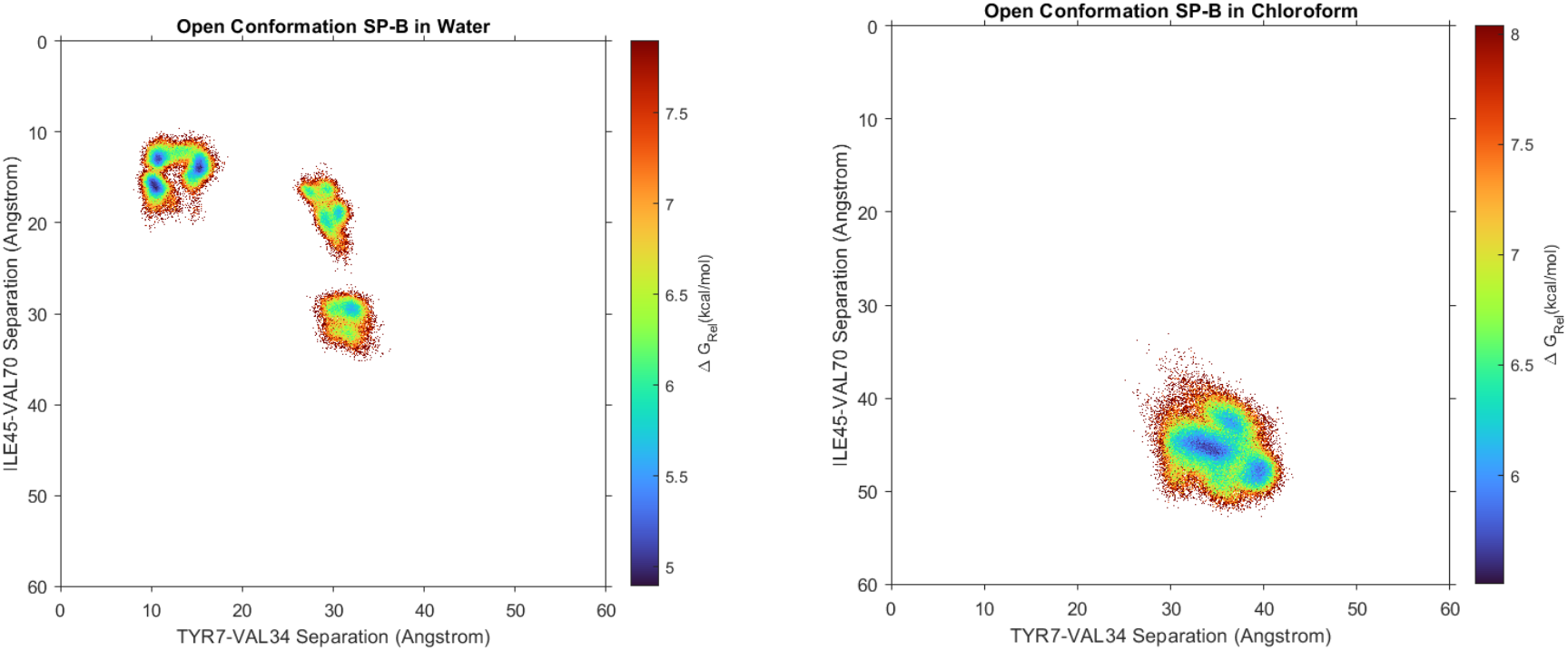
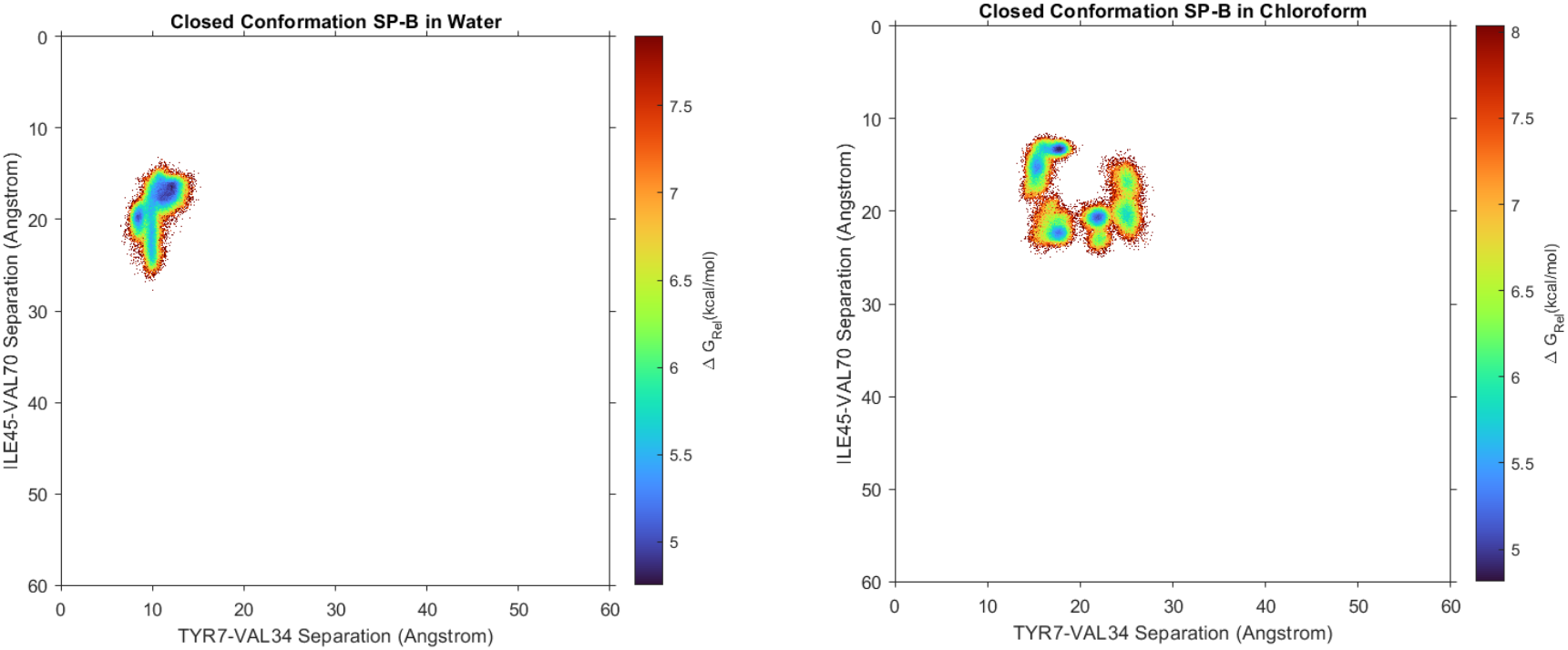
Free Energy heatmaps for each simulation. Open-water (top left), open-chloroform (top right), closed-water (bottom left), and closed-chloroform (bottom right). Relative free energy is shown by a color gradient, with blue corresponding to the lowest free-energy, most probable states, and red corresponding to the highest free-energy, least probable states.

### Helical Conformational Changes in Diverse Environments

The conformational flexibility of SP-B was further assessed using the-per residue root-mean-square fluctuation (RMSF) to identify regions of high variation within the protein. Regions of high variation correspond to more mobile residues in SP-B which would likely contribute to the conformational flexibility observed during the MD simulations.

To calculate the per-residue root-mean-square fluctuation, first the average protein structure was computed for each simulation. Each trajectory was then rotated/translated to minimize RMSD to the average structure. For each of these adjusted trajectories, the mass-weighted per-residue RMSF was calculated over the production simulation time (figure 8). High regions of RMSF were identified in each simulation as peaks with RMSF greater than the simulation mean plus one standard deviation. In the open-water simulation, the average RMSF was 11.6 Å, with 4 notable peaks of height greater than 14.3 Å located at residue indices 1-5, 24, 39-41, and 78-79. These regions correspond to the N- and C-terminal domains of the proteins, the hinge region between helices 1 and 2, and the bend between helices 2 and 3. In the open/chloroform simulation, the average RMSF was 10.5 Å with high RMSF peaks at residues 1-4, 37-42, and 45, corresponding to the N-terminal domain and the region between helices 2 and 3. In the closed-water simulation, the average RMSF was 11.3 Å with high RMSF peaks at residues 1-5, 24-26, 36, 64, and 79. Similarly to the open-water simulations, these peaks correspond to the N- and C-terminal domains, as well as the interhelical domains of SP-B. In the closed-chloroform simulation, the average RMSF was 11.3 Å with high RMSF peaks at residues 1-5, 23, 39-41, 64 and 79.

**Figure 8:**
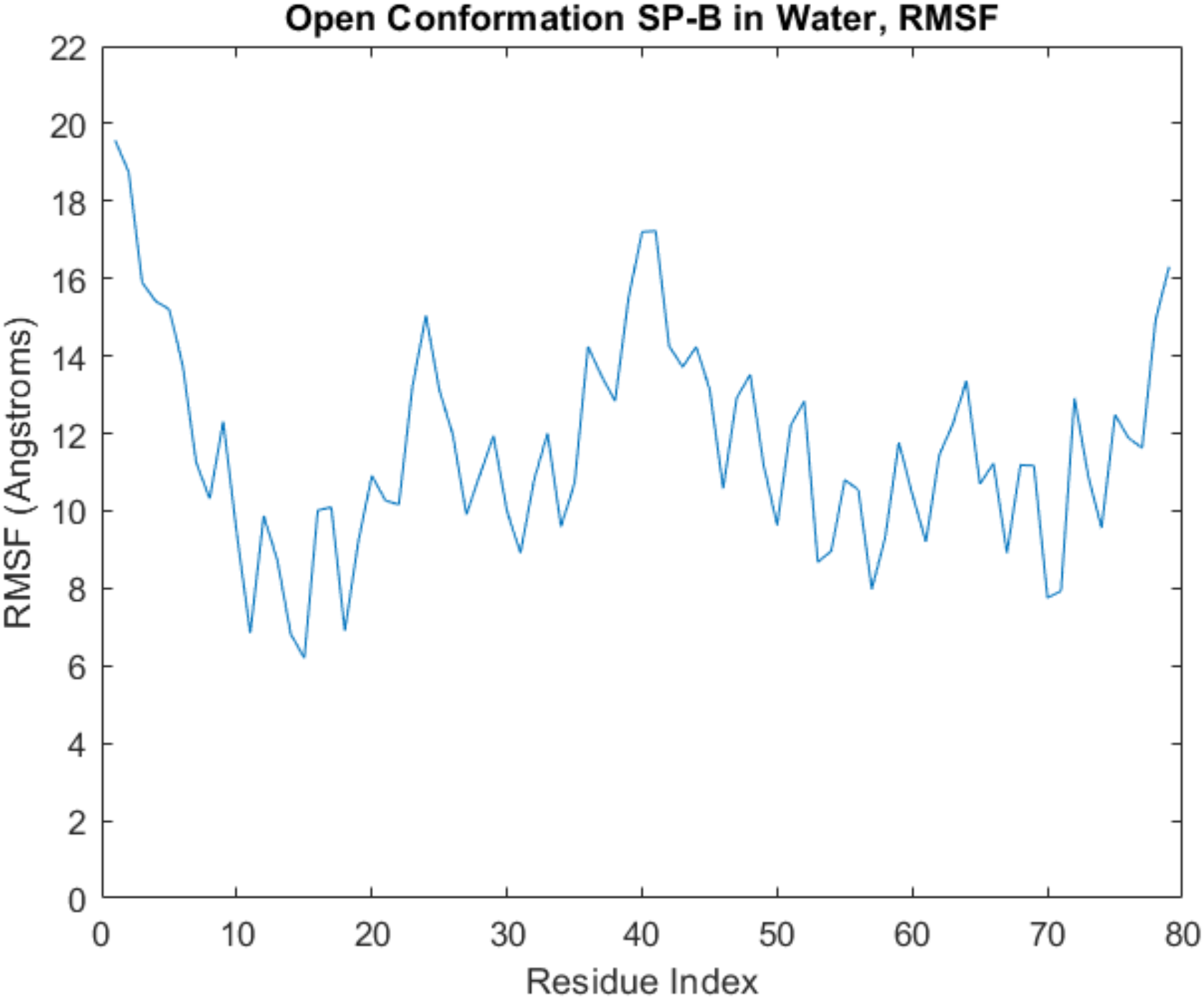
Per residue RMSF in open conformation/water simulation system. RMSF data for other simulation systems is included in supplemental information.

These were similar to peaks seen in other conditions, at the N-terminal, C-terminal and interhelical domains of the protein. It should be noted that the secondary structure of SP-B was not constant over the course of each simulation, and thus the position of each helix varied between and within the 4 simulations. Therefore, the high RMSF regions in this analysis identified as potential hinge regions should be considered an approximation of the hinge residue positions in SP-B.

As a member of the Saposin protein family, SP-B likely maintains the 4-helix secondary structure seen in other saposin proteins. Variation in the tertiary/quaternary structures of those proteins suggests that the specific orientation of the helices is an important factor in the biological function of Saposin proteins. These analyses seek to characterize the helical regions of SP-B when exposed to different solvents and establish how changes in the conformation of these helices impact its solvent interactions.

The secondary structure of the SP-B molecule was obtained at the end of each simulation using the Stride web interface (51). The secondary structures of SP-B at the end of each replicate are shown below in figure 9. Helices 1, 2, and 4 are generally well conserved in each system, with variations of 1-5 residues on each end, and minimal breaks in the middle of each helix. Helix 3 saw much more significant change over the course of the simulations. This was particularly pronounced in the open-water and closed-chloroform simulations, with loss of helicity on the terminal ends and multiple breaks seen in these simulations. This suggests that loss of helicity in helix 3 may be associated with larger scale conformational changes seen in SP-B, including closure in a hydrophilic environment and partial opening in a hydrophobic environment. These findings are further supported by the residue separation and heatmap analyses which identified a partially opened state with increased ILE45-VAL70 separation corresponding to a change in the helix 3-4 geometry.

**Figure 9:**
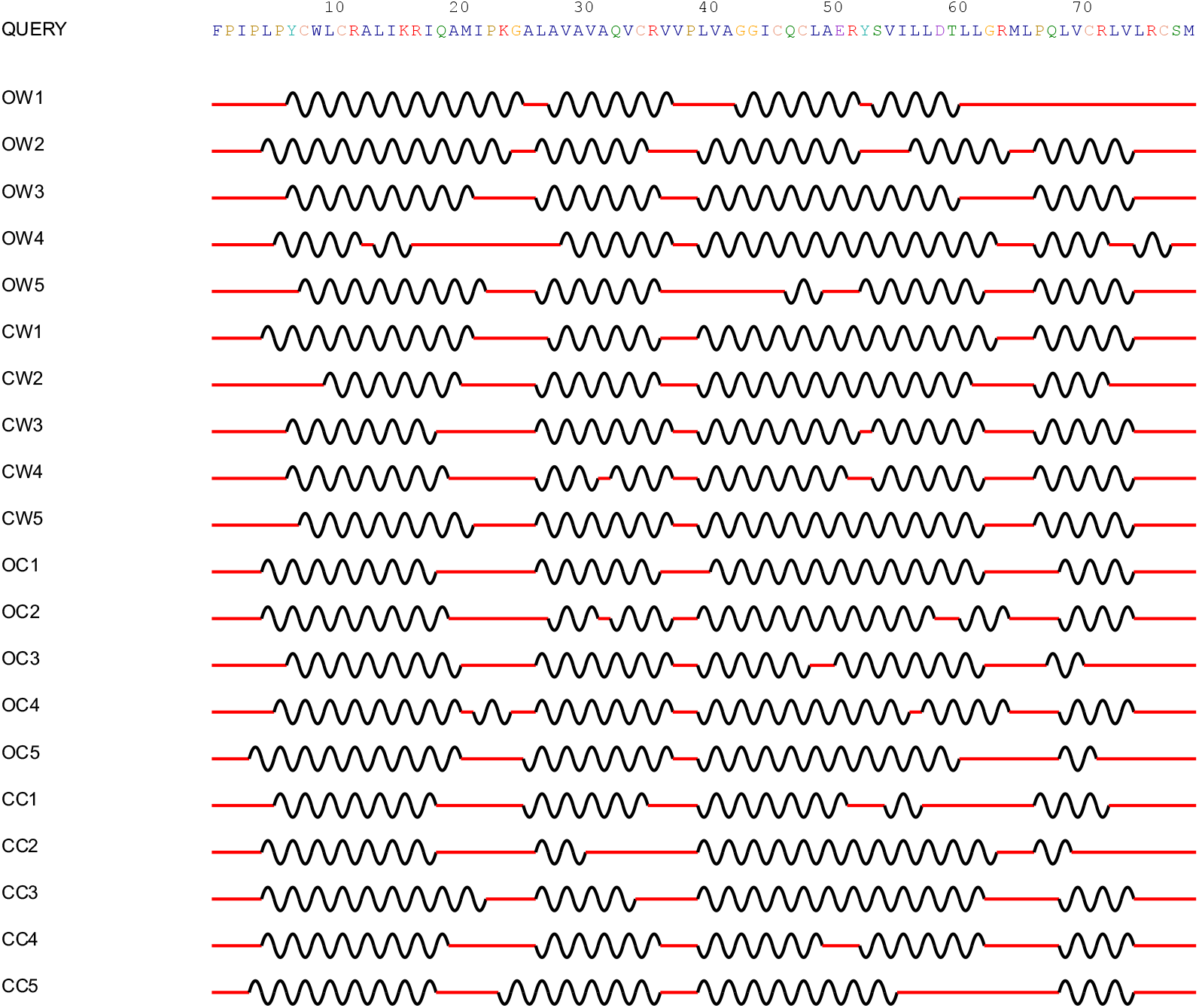
Secondary structure of protein molecule in simulation as reported by stride. Helical regions are denoted by a black curved line, while non-helical regions are denoted by a straight red line. Simulations are denoted by the initial conformation (O-open, C-closed), the solvent (C-chloroform, W-water), and the replicate number.

The precise positions of the helical regions of SP-B were not conserved across the 4 simulations performed. To compare solvent affinity of these regions across simulations, 4 helical regions for analysis were identified as being conserved among most replicate simulations. These regions are referred to as helix 1 (residues 9-19), helix 2 (residues 27-33), helix 3 (residues 46-61), and helix 4 (residues 69-72). For each region, the radial solvent density distribution was calculated about the center of mass of the region. This calculation was performed in CPPTraj, which calculates a histogram of solvent particle count as a function of the radial distance from the point of interest. This histogram is then normalized according to the bin volume, radius, and baseline solvent density so that the value of the SDF far from the solute molecule is equal to 1. The solvent distribution is compared across different regions and simulations to determine the effect of conformational change on potential solvent interactions. Since the open conformation quickly converged to a closed conformation in the water system, only the first 45 ns of this simulation were used.

The solvent distribution plot for helix 1 is shown in figure 10. Each solvent increases to a steady state at approximately 20 Å, representing the baseline density of the solvent in absence of a protein molecule. Closer to the molecule, there is some variation between the individual simulations. At approximately 5 Å, there is a shoulder in the distribution, with a higher amplitude in both chloroform systems likely due to the high hydrophobic character of this helix. Between 8 and 20 Å, the amplitude of both closed conformation systems is significantly lower than the corresponding open conformation simulations. This supports the hypothesis that the closed conformation reduced overall interaction with solvent molecules.

**Figure 10:**
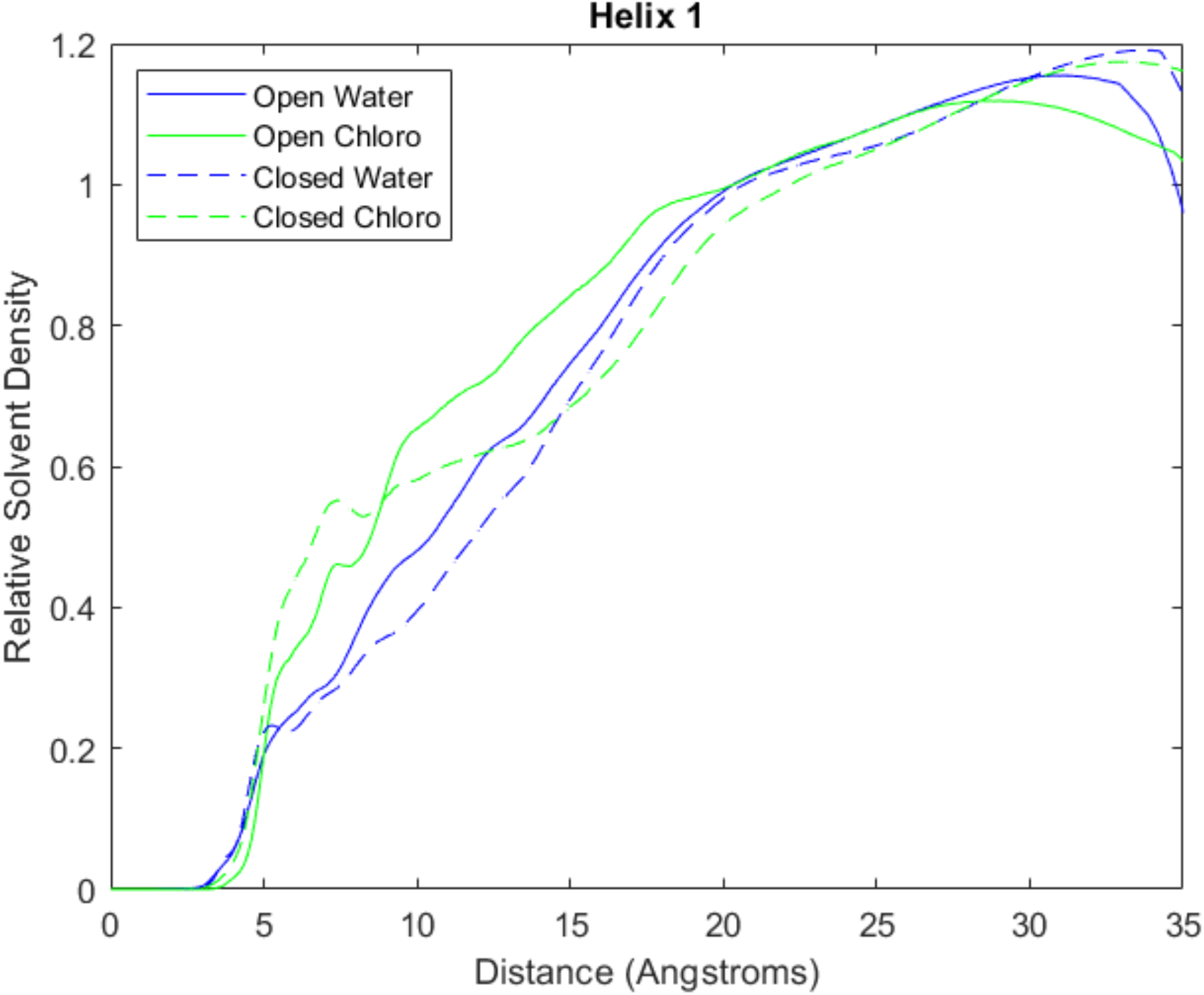
Radial solvent distribution function of four simulation systems centered at helix 1.

The helix 2 solvent distribution plot, shown in figure 11, reveals similar behavior. All 4 systems have an initial peak at approximately 8 Å, with the closed-chloroform systems having a higher density. This higher density is likely due to the extreme hydrophobicity of helix 2 compared to the other helices, with 6 hydrophobic amino acids in the 7-residue region. The initial peak in the closed-chloroform system is much lower than the open-chloroform system, suggesting that the closed conformation attenuates some of this interaction.

**Figure 11:**
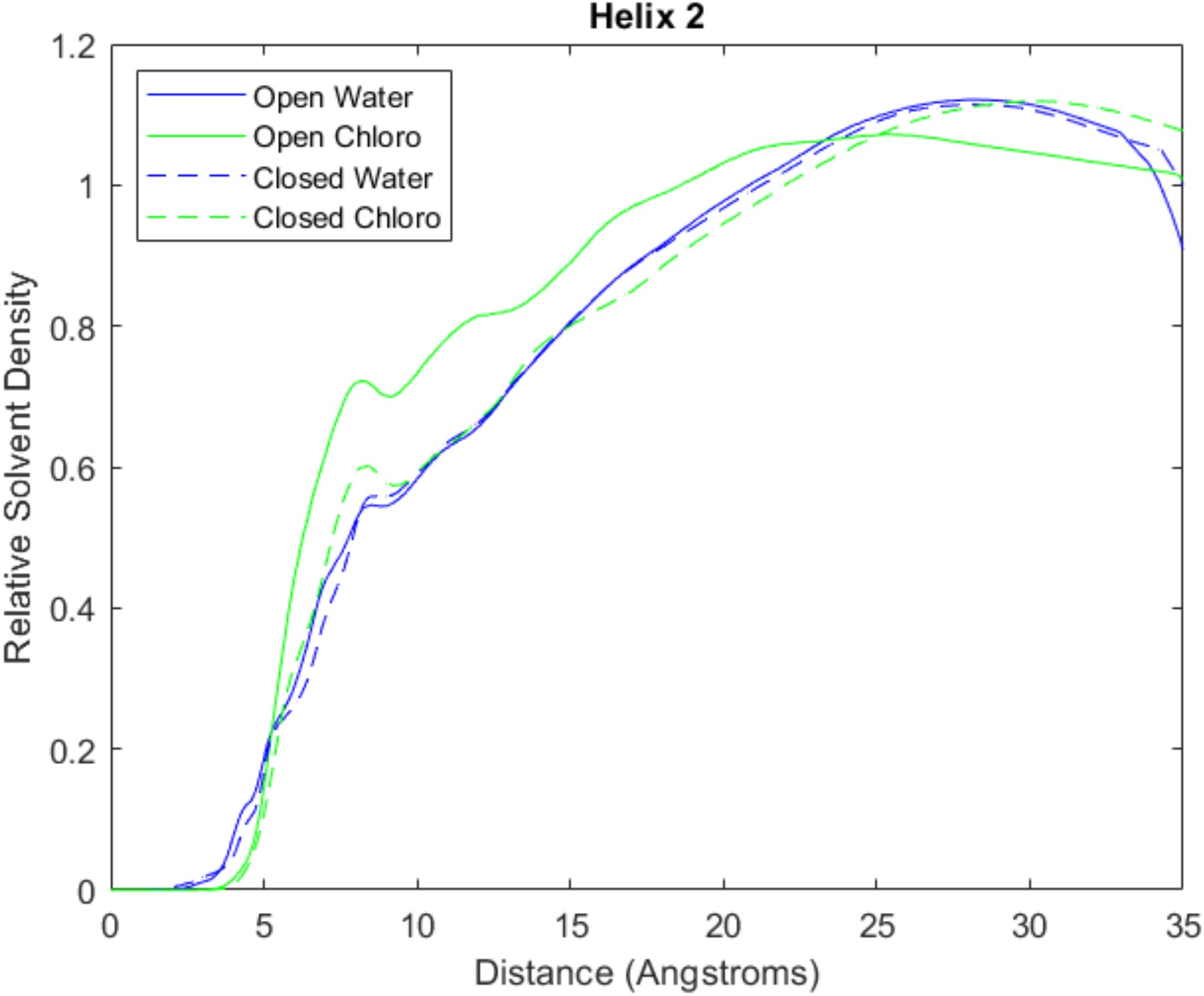
Radial solvent distribution function of four simulation systems centered at helix 2.

Similarly interesting phenomena are observed in the solvent distribution plot for helix 3, shown in figure 12. As observed in helix 1 and 2, there is a shoulder in all 4 systems at approximately 10 Å, with higher density in open/chloroform system. However, the closed-chloroform system exhibits significantly higher density in the 0-5 Å range which is not seen in any other condition. This information, in addition to the secondary structure variation seen in helix 3 in this region suggest that there is significant interaction between closed conformation SP-B and a hydrophobic solvent at helix 3. It is possible that the partial opening observed in combination with disruption of the secondary structure in helix 3 creates a pocket of favorable interaction for hydrophobic solvent molecules.

**Figure 12:**
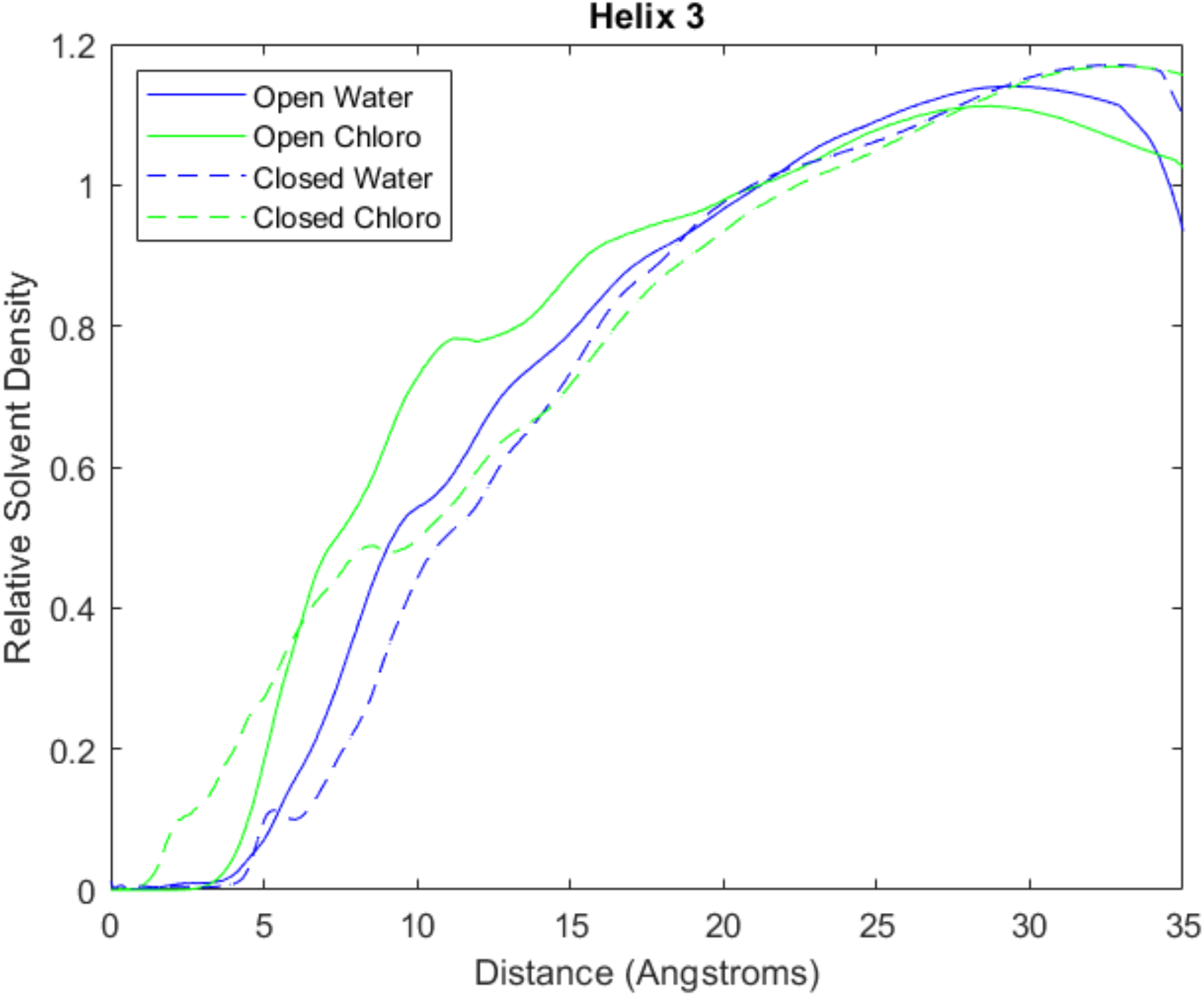
Radial solvent distribution function of four simulation systems centered at helix 3.

The solvent distribution plot for helix 4 is shown in figure 13 and exhibits similar features to that of helices 1 and 2, with similarly elevated open/chloroform densities in the 5-20 Å range likely due to the largely hydrophobic nature of SP-B.

**Figure 13:**
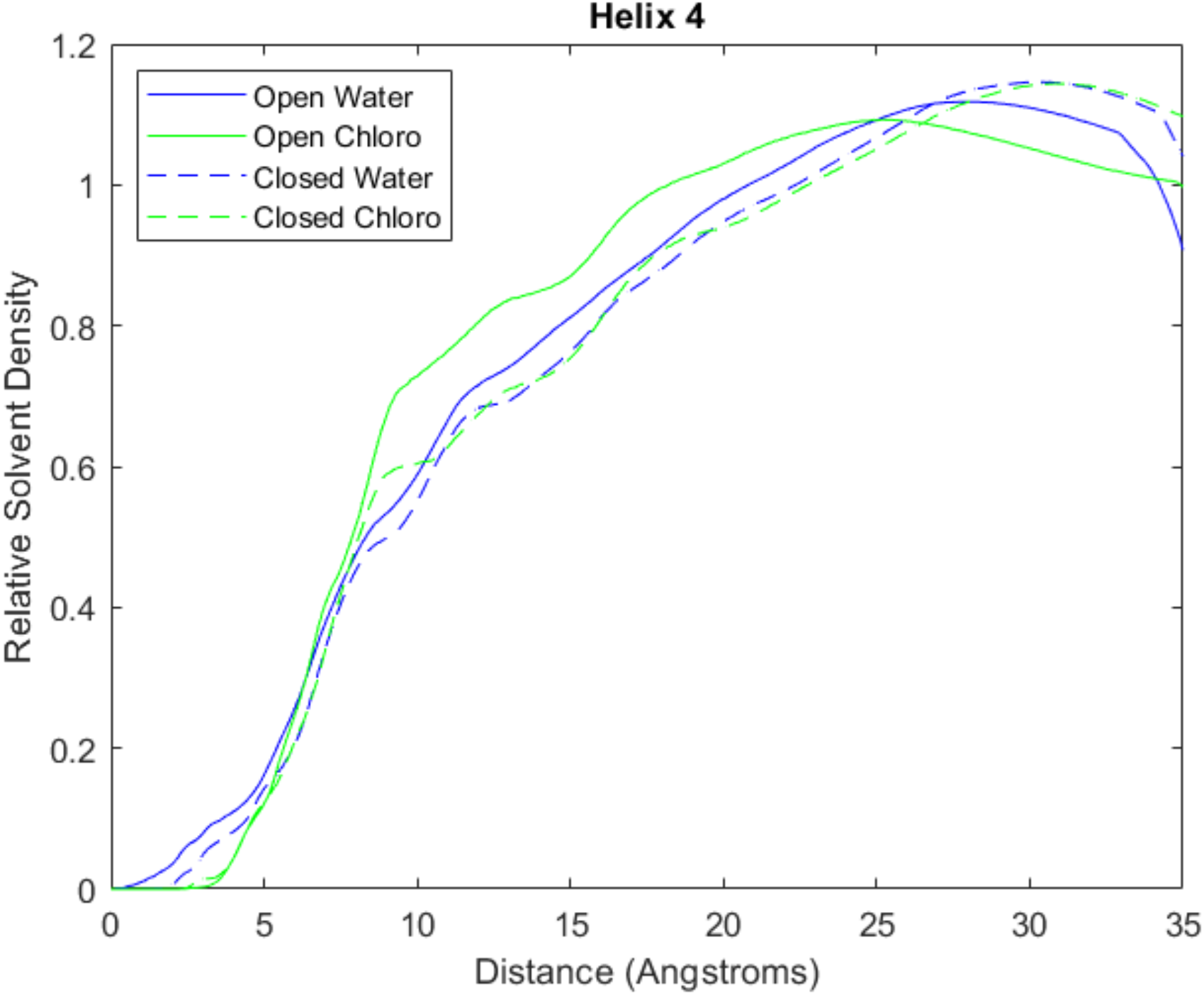
Radial solvent distribution function of four simulation systems centered at helix 4.

Our analysis of SP-B’s helicity and solvent interaction characteristics have revealed significant variations in the third helical region of SP-B when exposed to a hydrophobic solvent. This finding indicates that the specific conformation of this region of the protein may be an important factor in SP-B’s lipid manipulation function. Additionally, our solvent distribution data shows a region of favorable interaction with chloroform surrounding helix 3 only in the closed conformation. The dependence of this solvent interaction on conformation indicates that SP-B’s conformational flexibility allows for the 4 helices to be positioned in orientations that maximize interactions with different types of molecules.

This function may be an important factor in SP-B’s function in the varied and dynamic PS system, allowing it to interact with and stabilize different lipid structures that occur during different stages of respiration.

## Discussion

This study explores two hypothetical structural models of Surfactant Protein B (SP-B) - a closed conformation that minimizes hydrophobic exposure in aqueous solutions and an open conformation that presents a substantial hydrophobic surface for lipid interaction. The basis for these models draws on their homology with the saposin protein family, known for exhibiting a plethora of tertiary and quaternary structures that are tailored to their unique biological functions.

Our findings propose that SP-B is capable of adapting to diverse environments, leveraging its flexible structure for both hydrophobic interactions and insulation. This affirms SP-B’s ability to facilitate lipid restructuring due to its response to hydrophilic and hydrophobic environments at the pulmonary air-liquid interface, in which significant disruption and reformation of the surfactant monolayer regularly occurs. Through molecular dynamics simulations, we noted a predominant stability of the closed conformation, especially in a hydrophilic solvation environment. The closed conformation remained largely consistent across both hydrophilic and hydrophobic environments, albeit with minor secondary structural and conformational changes in the latter.

Interestingly, the open conformation demonstrated a significant conformational shift in a hydrophilic environment, transitioning towards a more energy-efficient closed conformation in some cases.

However, in a hydrophobic environment, the open conformation maintained its stability, taking on a further extended conformation. This suggests that the drastic conformational shift in hydrophilic conditions is likely an outcome of unfavorable protein-solvent interactions with the exposed hydrophobic residues. The free energy heatmaps further corroborated this, revealing more compact minima for the closed conformation in both solvent types. Additionally, a partially open conformation in which the C-terminal half of SP-B opens before the N-terminal half was observed in the open-water and closed-chloroform simulations. This may represent a stable transition structure between the open and closed conformations and could be involved in determining solvent accessibility.

RMSF and secondary structure analysis spotlighted significant variations in the protein’s third helical region, hinting at its crucial role in SP-B’s overall functionality. While the functions of individual helices within SP-B remain partially understood, it is plausible that the specific arrangement of polar and non-polar residues within each helix significantly impacts the protein’s functionality. Large-scale conformational shifts can manipulate these helices to expose or conceal these regions, thereby influencing potential lipid-protein interactions in pulmonary surfactant.

Our analysis of solvent distribution characteristics unveiled notable differences in solvent proximity between water and chloroform at various molecular points. The divergence in the spatial distribution of the solvent density near different molecule parts indicates how conformation and solvent type modulate local protein-solvent interactions. Particularly, the closed conformation of SP-B in chloroform simulation resulted in a radial solvent density peak in the <5-Å range surrounding helix 3. The specific orientation of these helices appears to significantly impact the interactions between SP-B and hydrophobic molecules.

## Conclusions

Our exploration of SP-B’s structure through homology modeling and molecular dynamics simulations has shed light on several critical aspects of this protein’s function within the pulmonary surfactant system.

The study emphasizes the existence of two distinct stable conformations subject to varying solvation environments and highlights the profound effect of solvent hydrophobicity on conformational stability.

Moreover, the protein’s conformation heavily influenced its interaction with hydrophobic solvent molecules, especially in the regions of helix 3. The critical positioning of these regions marks an essential discovery, paving the way for further research to fully comprehend their dynamics and interactions.

Large conformational changes may allow SP-B to modulate its interaction with surfactant lipids throughout the respiratory cycle, stabilizing the surfactant monolayer and maintaining alveolar surface tension.

Our work explores several less-explored SP-B mechanisms that could be instrumental in its pulmonary surfactant function. A comprehensive understanding of SP-B’s structural and functional characteristics at the atomic/molecular level could enhance our knowledge of pulmonary surfactants and their impact on respiratory physiology. Although we are still on the brink of achieving such comprehensive understanding, this study offers valuable insights that could guide future research in the right direction.

