## Supporting Information for "Conformational Changes of Surfactant Protein B Due to the Alveolar Air/Liquid Interface Using Molecular Dynamics"

### **Supplemental Information**

Table of Contents

S2 Peptide backbone RMSD of SP-B/chloroform systems

S3 TYR7-VAL34 and ILE45-VAL70 separation in chloroform simulations

S4 Per residue RMSF in all simulation systems

S5 SP-B Closed PDB

S43 SP-B Open PDB

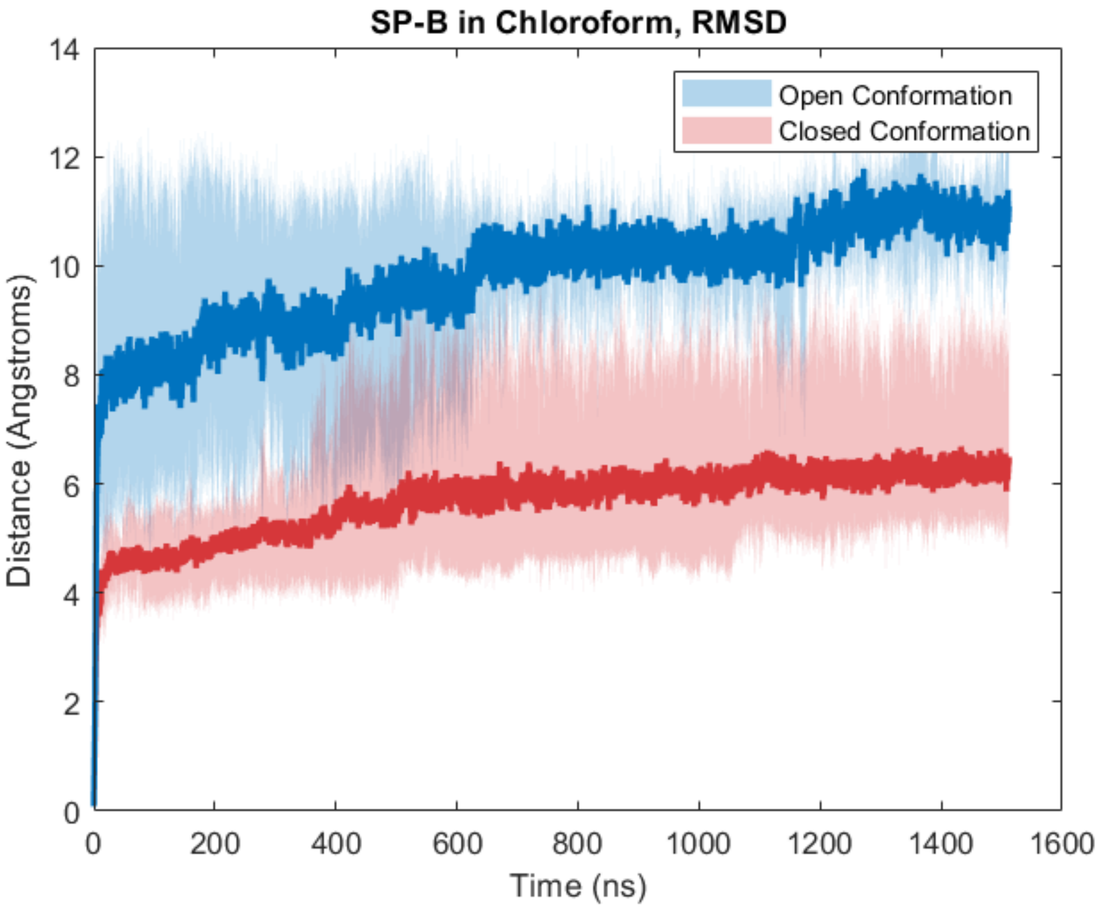

Figure S1: Peptide backbone RMSD of SP-B/chloroform systems. Initial open conformation systems shown in blue, initial closed conformation in red. Dark lines represent the mean trajectory over all replicates, while shaded areas show the minimum and maximum values at a given time point.

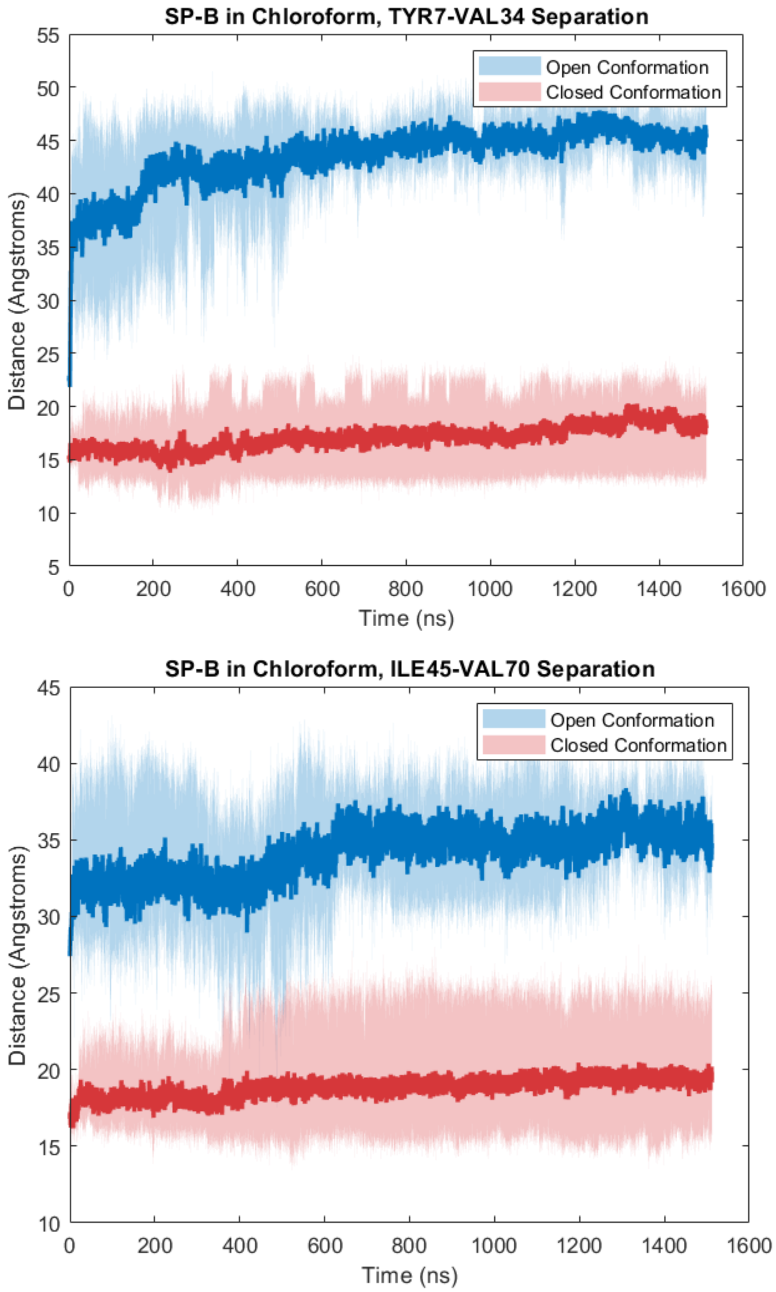

Figure S2: TYR7-VAL34 and ILE45-VAL70 separation in chloroform simulations. Initial open conformation systems shown in blue, initial closed conformation in red. Dark lines represent the mean trajectory over all replicates, while shaded areas show the minimum and maximum values at a given time point.

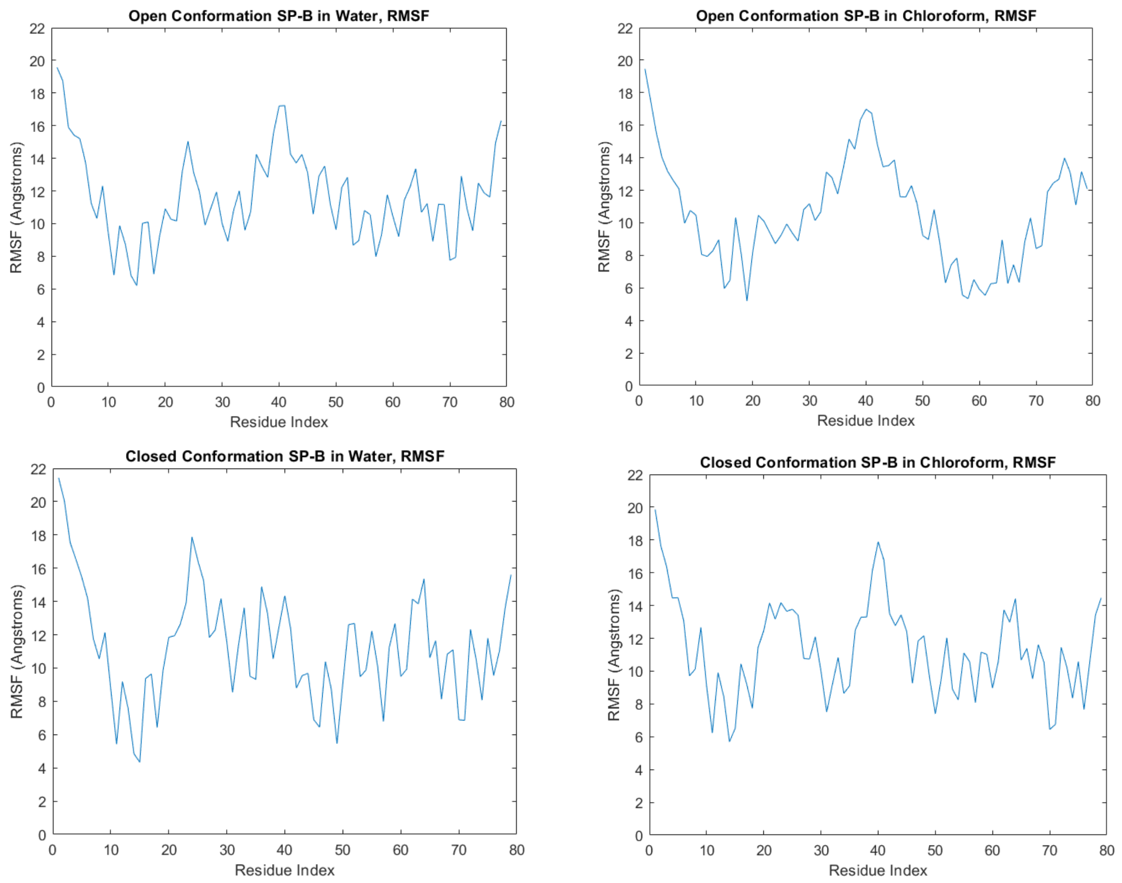

Figure S3: Per residue RMSF in each simulation system.

SP-B Closed PDB

| | | | | | | | | | | |
|---|---|---|---|---|---|---|---|---|---|---|
| ATOM | 1 | N | PHE | 1 | 22.457 | -1.776 | 38.724 | 0.1737 | 1.5500 | N |
| ATOM | 2 | H1 | PHE | 1 | 22.062 | -2.458 | 39.354 | 0.1921 | 1.3000 | H |
| ATOM | 3 | H2 | PHE | 1 | 21.724 | -1.190 | 38.350 | 0.1921 | 1.3000 | H |
| ATOM | 4 | H3 | PHE | 1 | 22.919 | -2.248 | 37.959 | 0.1921 | 1.3000 | H |
| ATOM | 5 | CA | PHE | 1 | 23.429 | -0.951 | 39.436 | 0.0733 | 1.7000 | C |
| ATOM | 6 | HA | PHE | 1 | 22.926 | -0.435 | 40.253 | 0.1041 | 1.2000 | H |
| ATOM | 7 | CB | PHE | 1 | 24.516 | -1.876 | 40.011 | 0.0330 | 1.7000 | C |
| ATOM | 8 | HB2 | PHE | 1 | 25.222 | -1.282 | 40.592 | 0.0104 | 1.2000 | H |
| ATOM | 9 | HB3 | PHE | 1 | 24.047 | -2.578 | 40.702 | 0.0104 | 1.2000 | H |
| ATOM | 10 | CG | PHE | 1 | 25.282 | -2.662 | 38.955 | 0.0031 | 1.7000 | C |
| ATOM | 11 | CD1 | PHE | 1 | 26.409 | -2.132 | 38.390 | -0.1392 | 1.7000 | C |
| ATOM | 12 | HD1 | PHE | 1 | 26.789 | -1.207 | 38.775 | 0.1374 | 1.2000 | H |
| ATOM | 13 | CE1 | PHE | 1 | 27.122 | -2.805 | 37.421 | -0.1602 | 1.7000 | C |
| ATOM | 14 | HE1 | PHE | 1 | 28.020 | -2.369 | 37.007 | 0.1433 | 1.2000 | H |
| ATOM | 15 | CZ | PHE | 1 | 26.708 | -4.041 | 37.002 | -0.1208 | 1.7000 | C |
| ATOM | 16 | HZ | PHE | 1 | 27.269 | -4.579 | 36.252 | 0.1329 | 1.2000 | H |
| ATOM | 17 | CE2 | PHE | 1 | 25.583 | -4.601 | 37.558 | -0.1603 | 1.7000 | C |
| ATOM | 18 | HE2 | PHE | 1 | 25.268 | -5.583 | 37.248 | 0.1433 | 1.2000 | H |
| ATOM | 19 | CD2 | PHE | 1 | 24.866 | -3.916 | 38.529 | -0.1391 | 1.7000 | C |
| ATOM | 20 | HD2 | PHE | 1 | 23.985 | -4.370 | 38.954 | 0.1374 | 1.2000 | H |
| ATOM | 21 | C | PHE | 1 | 24.032 | 0.085 | 38.467 | 0.6123 | 1.7000 | C |
| ATOM | 22 | O | PHE | 1 | 23.907 | -0.092 | 37.258 | -0.5713 | 1.5000 | O |
| ATOM | 23 | N | PRO | 2 | 24.624 | 1.208 | 38.942 | -0.2548 | 1.5500 | N |
| ATOM | 24 | CD | PRO | 2 | 24.570 | 1.561 | 40.383 | 0.0192 | 1.7000 | C |
| ATOM | 25 | HD2 | PRO | 2 | 25.327 | 1.002 | 40.933 | 0.0391 | 1.2000 | H |
| ATOM | 26 | HD3 | PRO | 2 | 23.578 | 1.420 | 40.813 | 0.0391 | 1.2000 | H |
| ATOM | 27 | CG | PRO | 2 | 24.931 | 3.034 | 40.351 | 0.0189 | 1.7000 | C |
| ATOM | 28 | HG2 | PRO | 2 | 25.418 | 3.344 | 41.277 | 0.0213 | 1.2000 | H |
| ATOM | 29 | HG3 | PRO | 2 | 24.028 | 3.625 | 40.187 | 0.0213 | 1.2000 | H |
| ATOM | 30 | CB | PRO | 2 | 25.853 | 3.184 | 39.161 | -0.0070 | 1.7000 | C |
| ATOM | 31 | HB2 | PRO | 2 | 26.863 | 2.901 | 39.462 | 0.0253 | 1.2000 | H |
| ATOM | 32 | HB3 | PRO | 2 | 25.852 | 4.207 | 38.784 | 0.0253 | 1.2000 | H |
| ATOM | 33 | CA | PRO | 2 | 25.302 | 2.205 | 38.117 | -0.0266 | 1.7000 | C |

| | | | | | | | | | | |
|---|---|---|---|---|---|---|---|---|---|---|
| ATOM | 34 | HA | PRO | 2 | 24.578 | 2.704 | 37.471 | 0.0641 | 1.2000 | H |
| ATOM | 35 | C | PRO | 2 | 26.431 | 1.587 | 37.309 | 0.5896 | 1.7000 | C |
| ATOM | 36 | O | PRO | 2 | 27.252 | 0.860 | 37.862 | -0.5748 | 1.5000 | O |
| ATOM | 37 | N | ILE | 3 | 26.540 | 1.952 | 36.040 | -0.4157 | 1.5500 | N |
| ATOM | 38 | H | ILE | 3 | 25.865 | 2.583 | 35.634 | 0.2719 | 1.3000 | H |
| ATOM | 39 | CA | ILE | 3 | 27.603 | 1.424 | 35.189 | -0.0597 | 1.7000 | C |
| ATOM | 40 | HA | ILE | 3 | 28.191 | 0.692 | 35.737 | 0.0869 | 1.2000 | H |
| ATOM | 41 | CB | ILE | 3 | 26.994 | 0.617 | 34.040 | 0.1303 | 1.7000 | C |
| ATOM | 42 | HB | ILE | 3 | 26.328 | 1.264 | 33.463 | 0.0187 | 1.2000 | H |
| ATOM | 43 | CG2 | ILE | 3 | 28.065 | 0.130 | 33.151 | -0.3204 | 1.7000 | C |
| ATOM | 44 | HG21 | ILE | 3 | 27.602 | -0.418 | 32.339 | 0.0882 | 1.2000 | H |
| ATOM | 45 | HG22 | ILE | 3 | 28.625 | 0.939 | 32.686 | 0.0882 | 1.2000 | H |
| ATOM | 46 | HG23 | ILE | 3 | 28.749 | -0.528 | 33.689 | 0.0882 | 1.2000 | H |
| ATOM | 47 | CG1 | ILE | 3 | 26.159 | -0.559 | 34.620 | -0.0430 | 1.7000 | C |
| ATOM | 48 | HG12 | ILE | 3 | 26.828 | -1.247 | 35.137 | 0.0236 | 1.2000 | H |
| ATOM | 49 | HG13 | ILE | 3 | 25.444 | -0.178 | 35.339 | 0.0236 | 1.2000 | H |
| ATOM | 50 | CD1 | ILE | 3 | 25.352 | -1.341 | 33.623 | -0.0660 | 1.7000 | C |
| ATOM | 51 | HD11 | ILE | 3 | 24.789 | -2.114 | 34.145 | 0.0186 | 1.2000 | H |
| ATOM | 52 | HD12 | ILE | 3 | 24.647 | -0.677 | 33.125 | 0.0186 | 1.2000 | H |
| ATOM | 53 | HD13 | ILE | 3 | 26.001 | -1.819 | 32.894 | 0.0186 | 1.2000 | H |
| ATOM | 54 | C | ILE | 3 | 28.525 | 2.525 | 34.635 | 0.5973 | 1.7000 | C |
| ATOM | 55 | O | ILE | 3 | 28.111 | 3.265 | 33.742 | -0.5679 | 1.5000 | O |
| ATOM | 56 | N | PRO | 4 | 29.759 | 2.685 | 35.156 | -0.2548 | 1.5500 | N |
| ATOM | 57 | CD | PRO | 4 | 30.215 | 1.861 | 36.295 | 0.0192 | 1.7000 | C |
| ATOM | 58 | HD2 | PRO | 4 | 30.589 | 0.901 | 35.937 | 0.0391 | 1.2000 | H |
| ATOM | 59 | HD3 | PRO | 4 | 29.454 | 1.738 | 37.063 | 0.0391 | 1.2000 | H |
| ATOM | 60 | CG | PRO | 4 | 31.342 | 2.695 | 36.852 | 0.0189 | 1.7000 | C |
| ATOM | 61 | HG2 | PRO | 4 | 32.092 | 2.073 | 37.343 | 0.0213 | 1.2000 | H |
| ATOM | 62 | HG3 | PRO | 4 | 30.938 | 3.423 | 37.557 | 0.0213 | 1.2000 | H |
| ATOM | 63 | CB | PRO | 4 | 31.917 | 3.418 | 35.662 | -0.0070 | 1.7000 | C |
| ATOM | 64 | HB2 | PRO | 4 | 32.625 | 2.776 | 35.164 | 0.0253 | 1.2000 | H |
| ATOM | 65 | HB3 | PRO | 4 | 32.409 | 4.345 | 35.960 | 0.0253 | 1.2000 | H |
| ATOM | 66 | CA | PRO | 4 | 30.725 | 3.692 | 34.746 | -0.0266 | 1.7000 | C |
| ATOM | 67 | HA | PRO | 4 | 30.318 | 4.684 | 34.949 | 0.0641 | 1.2000 | H |

| | | | | | | | | | | |
|---|---|---|---|---|---|---|---|---|---|---|
| ATOM | 68 | C | PRO | 4 | 31.091 | 3.547 | 33.271 | 0.5896 | 1.7000 | C |
| ATOM | 69 | O | PRO | 4 | 31.234 | 2.431 | 32.752 | -0.5748 | 1.5000 | O |
| ATOM | 70 | N | LEU | 5 | 31.359 | 4.669 | 32.620 | -0.4157 | 1.5500 | N |
| ATOM | 71 | H | LEU | 5 | 31.264 | 5.565 | 33.076 | 0.2719 | 1.3000 | H |
| ATOM | 72 | CA | LEU | 5 | 31.694 | 4.627 | 31.208 | -0.0518 | 1.7000 | C |
| ATOM | 73 | HA | LEU | 5 | 30.849 | 4.159 | 30.708 | 0.0922 | 1.2000 | H |
| ATOM | 74 | CB | LEU | 5 | 31.849 | 6.049 | 30.673 | -0.1102 | 1.7000 | C |
| ATOM | 75 | HB2 | LEU | 5 | 32.527 | 6.605 | 31.318 | 0.0457 | 1.2000 | H |
| ATOM | 76 | HB3 | LEU | 5 | 32.317 | 5.995 | 29.694 | 0.0457 | 1.2000 | H |
| ATOM | 77 | CG | LEU | 5 | 30.561 | 6.821 | 30.553 | 0.3531 | 1.7000 | C |
| ATOM | 78 | HG | LEU | 5 | 30.013 | 6.788 | 31.496 | -0.0361 | 1.2000 | H |
| ATOM | 79 | CD1 | LEU | 5 | 30.859 | 8.271 | 30.224 | -0.4121 | 1.7000 | C |
| ATOM | 80 | HD11 | LEU | 5 | 29.927 | 8.832 | 30.157 | 0.1000 | 1.2000 | H |
| ATOM | 81 | HD12 | LEU | 5 | 31.474 | 8.715 | 31.008 | 0.1000 | 1.2000 | H |
| ATOM | 82 | HD13 | LEU | 5 | 31.383 | 8.346 | 29.273 | 0.1000 | 1.2000 | H |
| ATOM | 83 | CD2 | LEU | 5 | 29.728 | 6.181 | 29.462 | -0.4121 | 1.7000 | C |
| ATOM | 84 | HD21 | LEU | 5 | 28.872 | 6.819 | 29.239 | 0.1000 | 1.2000 | H |
| ATOM | 85 | HD22 | LEU | 5 | 30.309 | 6.056 | 28.547 | 0.1000 | 1.2000 | H |
| ATOM | 86 | HD23 | LEU | 5 | 29.331 | 5.219 | 29.781 | 0.1000 | 1.2000 | H |
| ATOM | 87 | C | LEU | 5 | 32.904 | 3.763 | 30.819 | 0.5973 | 1.7000 | C |
| ATOM | 88 | O | LEU | 5 | 32.787 | 3.054 | 29.825 | -0.5679 | 1.5000 | O |
| ATOM | 89 | N | PRO | 6 | 34.047 | 3.738 | 31.547 | -0.2548 | 1.5500 | N |
| ATOM | 90 | CD | PRO | 6 | 34.276 | 4.637 | 32.696 | 0.0192 | 1.7000 | C |
| ATOM | 91 | HD2 | PRO | 6 | 33.810 | 4.228 | 33.591 | 0.0391 | 1.2000 | H |
| ATOM | 92 | HD3 | PRO | 6 | 33.962 | 5.661 | 32.512 | 0.0391 | 1.2000 | H |
| ATOM | 93 | CG | PRO | 6 | 35.775 | 4.586 | 32.842 | 0.0189 | 1.7000 | C |
| ATOM | 94 | HG2 | PRO | 6 | 36.081 | 4.767 | 33.873 | 0.0213 | 1.2000 | H |
| ATOM | 95 | HG3 | PRO | 6 | 36.228 | 5.325 | 32.178 | 0.0213 | 1.2000 | H |
| ATOM | 96 | CB | PRO | 6 | 36.158 | 3.203 | 32.386 | -0.0070 | 1.7000 | C |
| ATOM | 97 | HB2 | PRO | 6 | 36.007 | 2.502 | 33.209 | 0.0253 | 1.2000 | H |
| ATOM | 98 | HB3 | PRO | 6 | 37.196 | 3.166 | 32.052 | 0.0253 | 1.2000 | H |
| ATOM | 99 | CA | PRO | 6 | 35.194 | 2.893 | 31.239 | -0.0266 | 1.7000 | C |
| ATOM | 100 | HA | PRO | 6 | 35.631 | 3.204 | 30.292 | 0.0641 | 1.2000 | H |
| ATOM | 101 | C | PRO | 6 | 34.870 | 1.405 | 31.199 | 0.5896 | 1.7000 | C |

| | | | | | | | | | | |
|---|---|---|---|---|---|---|---|---|---|---|
| ATOM | 102 | O | PRO | 6 | 35.537 | 0.643 | 30.501 | -0.5748 | 1.5000 | O |
| ATOM | 103 | N | TYR | 7 | 33.811 | 0.985 | 31.894 | -0.4157 | 1.5500 | N |
| ATOM | 104 | H | TYR | 7 | 33.248 | 1.640 | 32.417 | 0.2719 | 1.3000 | H |
| ATOM | 105 | CA | TYR | 7 | 33.452 | -0.421 | 31.927 | -0.0014 | 1.7000 | C |
| ATOM | 106 | HA | TYR | 7 | 34.355 | -1.031 | 31.985 | 0.0876 | 1.2000 | H |
| ATOM | 107 | CB | TYR | 7 | 32.627 | -0.673 | 33.181 | -0.0152 | 1.7000 | C |
| ATOM | 108 | HB2 | TYR | 7 | 31.792 | 0.022 | 33.245 | 0.0295 | 1.2000 | H |
| ATOM | 109 | HB3 | TYR | 7 | 32.213 | -1.681 | 33.128 | 0.0295 | 1.2000 | H |
| ATOM | 110 | CG | TYR | 7 | 33.501 | -0.555 | 34.420 | -0.0011 | 1.7000 | C |
| ATOM | 111 | CD1 | TYR | 7 | 34.061 | 0.659 | 34.710 | -0.1906 | 1.7000 | C |
| ATOM | 112 | HD1 | TYR | 7 | 33.876 | 1.537 | 34.128 | 0.1699 | 1.2000 | H |
| ATOM | 113 | CE1 | TYR | 7 | 34.840 | 0.829 | 35.829 | -0.2341 | 1.7000 | C |
| ATOM | 114 | HE1 | TYR | 7 | 35.261 | 1.797 | 36.059 | 0.1656 | 1.2000 | H |
| ATOM | 115 | CZ | TYR | 7 | 35.052 | -0.241 | 36.681 | 0.3226 | 1.7000 | C |
| ATOM | 116 | OH | TYR | 7 | 35.825 | -0.081 | 37.811 | -0.5579 | 1.5000 | O |
| ATOM | 117 | HH | TYR | 7 | 35.881 | -0.878 | 38.344 | 0.3992 | 1.2000 | H |
| ATOM | 118 | CE2 | TYR | 7 | 34.480 | -1.461 | 36.407 | -0.2341 | 1.7000 | C |
| ATOM | 119 | HE2 | TYR | 7 | 34.626 | -2.290 | 37.083 | 0.1656 | 1.2000 | H |
| ATOM | 120 | CD2 | TYR | 7 | 33.701 | -1.621 | 35.280 | -0.1906 | 1.7000 | C |
| ATOM | 121 | HD2 | TYR | 7 | 33.228 | -2.572 | 35.095 | 0.1699 | 1.2000 | H |
| ATOM | 122 | C | TYR | 7 | 32.737 | -0.744 | 30.636 | 0.5973 | 1.7000 | C |
| ATOM | 123 | O | TYR | 7 | 32.978 | -1.785 | 30.018 | -0.5679 | 1.5000 | O |
| ATOM | 124 | N | CYX | 8 | 31.891 | 0.184 | 30.193 | -0.4157 | 1.5500 | N |
| ATOM | 125 | H | CYX | 8 | 31.734 | 1.027 | 30.727 | 0.2719 | 1.3000 | H |
| ATOM | 126 | CA | CYX | 8 | 31.186 | 0.001 | 28.934 | 0.0429 | 1.7000 | C |
| ATOM | 127 | HA | CYX | 8 | 30.750 | -0.998 | 28.924 | 0.0766 | 1.2000 | H |
| ATOM | 128 | CB | CYX | 8 | 30.068 | 1.023 | 28.758 | -0.0790 | 1.7000 | C |
| ATOM | 129 | HB2 | CYX | 8 | 29.395 | 0.950 | 29.611 | 0.0910 | 1.2000 | H |
| ATOM | 130 | HB3 | CYX | 8 | 30.491 | 2.028 | 28.751 | 0.0910 | 1.2000 | H |
| ATOM | 131 | SG | CYX | 8 | 29.102 | 0.781 | 27.226 | -0.1081 | 1.8000 | S |
| ATOM | 132 | C | CYX | 8 | 32.135 | 0.133 | 27.754 | 0.5973 | 1.7000 | C |
| ATOM | 133 | O | CYX | 8 | 32.066 | -0.641 | 26.791 | -0.5679 | 1.5000 | O |
| ATOM | 134 | N | TRP | 9 | 33.016 | 1.127 | 27.816 | -0.4157 | 1.5500 | N |
| ATOM | 135 | H | TRP | 9 | 33.041 | 1.737 | 28.618 | 0.2719 | 1.3000 | H |

| | | | | | | | | | | |
|---|---|---|---|---|---|---|---|---|---|---|
| ATOM | 136 | CA | TRP | 9 | 33.945 | 1.362 | 26.730 | -0.0275 | 1.7000 | C |
| ATOM | 137 | HA | TRP | 9 | 33.396 | 1.549 | 25.806 | 0.1123 | 1.2000 | H |
| ATOM | 138 | CB | TRP | 9 | 34.833 | 2.558 | 27.058 | -0.0050 | 1.7000 | C |
| ATOM | 139 | HB2 | TRP | 9 | 35.262 | 2.437 | 28.055 | 0.0339 | 1.2000 | H |
| ATOM | 140 | HB3 | TRP | 9 | 35.653 | 2.593 | 26.341 | 0.0339 | 1.2000 | H |
| ATOM | 141 | CG | TRP | 9 | 34.107 | 3.873 | 26.983 | -0.1415 | 1.7000 | C |
| ATOM | 142 | CD1 | TRP | 9 | 32.975 | 4.132 | 26.272 | -0.1638 | 1.7000 | C |
| ATOM | 143 | HD1 | TRP | 9 | 32.442 | 3.425 | 25.655 | 0.2062 | 1.2000 | H |
| ATOM | 144 | NE1 | TRP | 9 | 32.604 | 5.442 | 26.433 | -0.3418 | 1.5500 | N |
| ATOM | 145 | HE1 | TRP | 9 | 31.807 | 5.868 | 25.983 | 0.3412 | 1.3000 | H |
| ATOM | 146 | CE2 | TRP | 9 | 33.497 | 6.057 | 27.267 | 0.1380 | 1.7000 | C |
| ATOM | 147 | CZ2 | TRP | 9 | 33.538 | 7.360 | 27.720 | -0.2601 | 1.7000 | C |
| ATOM | 148 | HZ2 | TRP | 9 | 32.795 | 8.081 | 27.419 | 0.1572 | 1.2000 | H |
| ATOM | 149 | CH2 | TRP | 9 | 34.565 | 7.707 | 28.593 | -0.1134 | 1.7000 | C |
| ATOM | 150 | HH2 | TRP | 9 | 34.620 | 8.720 | 28.966 | 0.1417 | 1.2000 | H |
| ATOM | 151 | CZ3 | TRP | 9 | 35.509 | 6.798 | 28.984 | -0.1972 | 1.7000 | C |
| ATOM | 152 | HZ3 | TRP | 9 | 36.302 | 7.104 | 29.651 | 0.1447 | 1.2000 | H |
| ATOM | 153 | CE3 | TRP | 9 | 35.478 | 5.495 | 28.518 | -0.2387 | 1.7000 | C |
| ATOM | 154 | HE3 | TRP | 9 | 36.236 | 4.788 | 28.816 | 0.1700 | 1.2000 | H |
| ATOM | 155 | CD2 | TRP | 9 | 34.451 | 5.114 | 27.648 | 0.1243 | 1.7000 | C |
| ATOM | 156 | C | TRP | 9 | 34.787 | 0.121 | 26.519 | 0.5973 | 1.7000 | C |
| ATOM | 157 | O | TRP | 9 | 34.943 | -0.341 | 25.381 | -0.5679 | 1.5000 | O |
| ATOM | 158 | N | LEU | 10 | 35.279 | -0.462 | 27.617 | -0.4157 | 1.5500 | N |
| ATOM | 159 | H | LEU | 10 | 35.125 | -0.056 | 28.531 | 0.2719 | 1.3000 | H |
| ATOM | 160 | CA | LEU | 10 | 36.009 | -1.701 | 27.511 | -0.0518 | 1.7000 | C |
| ATOM | 161 | HA | LEU | 10 | 36.741 | -1.583 | 26.716 | 0.0922 | 1.2000 | H |
| ATOM | 162 | CB | LEU | 10 | 36.726 | -2.014 | 28.804 | -0.1102 | 1.7000 | C |
| ATOM | 163 | HB2 | LEU | 10 | 36.008 | -1.902 | 29.618 | 0.0457 | 1.2000 | H |
| ATOM | 164 | HB3 | LEU | 10 | 37.040 | -3.059 | 28.798 | 0.0457 | 1.2000 | H |
| ATOM | 165 | CG | LEU | 10 | 37.906 | -1.143 | 29.060 | 0.3531 | 1.7000 | C |
| ATOM | 166 | HG | LEU | 10 | 37.648 | -0.091 | 28.934 | -0.0361 | 1.2000 | H |
| ATOM | 167 | CD1 | LEU | 10 | 38.368 | -1.356 | 30.437 | -0.4121 | 1.7000 | C |
| ATOM | 168 | HD11 | LEU | 10 | 39.245 | -0.745 | 30.634 | 0.1000 | 1.2000 | H |
| ATOM | 169 | HD12 | LEU | 10 | 37.586 | -1.046 | 31.129 | 0.1000 | 1.2000 | H |

| | | | | | | | | | | |
|---|---|---|---|---|---|---|---|---|---|---|
| ATOM | 170 | HD13 | LEU | 10 | 38.607 | -2.405 | 30.600 | 0.1000 | 1.2000 | H |
| ATOM | 171 | CD2 | LEU | 10 | 38.993 | -1.526 | 28.046 | -0.4121 | 1.7000 | C |
| ATOM | 172 | HD21 | LEU | 10 | 39.926 | -1.031 | 28.294 | 0.1000 | 1.2000 | H |
| ATOM | 173 | HD22 | LEU | 10 | 39.172 | -2.602 | 28.055 | 0.1000 | 1.2000 | H |
| ATOM | 174 | HD23 | LEU | 10 | 38.730 | -1.210 | 27.038 | 0.1000 | 1.2000 | H |
| ATOM | 175 | C | LEU | 10 | 35.137 | -2.849 | 27.085 | 0.5973 | 1.7000 | C |
| ATOM | 176 | O | LEU | 10 | 35.572 | -3.639 | 26.270 | -0.5679 | 1.5000 | O |
| ATOM | 177 | N | CYX | 11 | 33.894 | -2.940 | 27.553 | -0.4157 | 1.5500 | N |
| ATOM | 178 | H | CYX | 11 | 33.536 | -2.273 | 28.222 | 0.2719 | 1.3000 | H |
| ATOM | 179 | CA | CYX | 11 | 33.056 | -4.049 | 27.110 | 0.0429 | 1.7000 | C |
| ATOM | 180 | HA | CYX | 11 | 33.491 | -4.984 | 27.467 | 0.0766 | 1.2000 | H |
| ATOM | 181 | CB | CYX | 11 | 31.643 | -3.927 | 27.663 | -0.0790 | 1.7000 | C |
| ATOM | 182 | HB2 | CYX | 11 | 31.721 | -3.923 | 28.746 | 0.0910 | 1.2000 | H |
| ATOM | 183 | HB3 | CYX | 11 | 31.195 | -2.982 | 27.355 | 0.0910 | 1.2000 | H |
| ATOM | 184 | SG | CYX | 11 | 30.543 | -5.288 | 27.169 | -0.1081 | 1.8000 | S |
| ATOM | 185 | C | CYX | 11 | 32.968 | -4.091 | 25.601 | 0.5973 | 1.7000 | C |
| ATOM | 186 | O | CYX | 11 | 33.165 | -5.133 | 24.976 | -0.5679 | 1.5000 | O |
| ATOM | 187 | N | ARG | 12 | 32.655 | -2.956 | 25.002 | -0.3479 | 1.5500 | N |
| ATOM | 188 | H | ARG | 12 | 32.511 | -2.109 | 25.535 | 0.2747 | 1.3000 | H |
| ATOM | 189 | CA | ARG | 12 | 32.493 | -2.941 | 23.570 | -0.2637 | 1.7000 | C |
| ATOM | 190 | HA | ARG | 12 | 31.848 | -3.774 | 23.281 | 0.1560 | 1.2000 | H |
| ATOM | 191 | CB | ARG | 12 | 31.790 | -1.658 | 23.189 | -0.0007 | 1.7000 | C |
| ATOM | 192 | HB2 | ARG | 12 | 32.323 | -0.807 | 23.619 | 0.0327 | 1.2000 | H |
| ATOM | 193 | HB3 | ARG | 12 | 31.802 | -1.555 | 22.103 | 0.0327 | 1.2000 | H |
| ATOM | 194 | CG | ARG | 12 | 30.335 | -1.648 | 23.661 | 0.0390 | 1.7000 | C |
| ATOM | 195 | HG2 | ARG | 12 | 29.801 | -2.460 | 23.163 | 0.0285 | 1.2000 | H |
| ATOM | 196 | HG3 | ARG | 12 | 30.305 | -1.830 | 24.736 | 0.0285 | 1.2000 | H |
| ATOM | 197 | CD | ARG | 12 | 29.628 | -0.381 | 23.404 | 0.0486 | 1.7000 | C |
| ATOM | 198 | HD2 | ARG | 12 | 30.142 | 0.421 | 23.937 | 0.0687 | 1.2000 | H |
| ATOM | 199 | HD3 | ARG | 12 | 29.644 | -0.167 | 22.335 | 0.0687 | 1.2000 | H |
| ATOM | 200 | NE | ARG | 12 | 28.246 | -0.461 | 23.859 | -0.5295 | 1.5500 | N |
| ATOM | 201 | HE | ARG | 12 | 27.919 | -1.367 | 24.160 | 0.3456 | 1.3000 | H |
| ATOM | 202 | CZ | ARG | 12 | 27.399 | 0.579 | 23.962 | 0.8076 | 1.7000 | C |
| ATOM | 203 | NH1 | ARG | 12 | 27.791 | 1.789 | 23.629 | -0.8627 | 1.5500 | N |

| | | | | | | | | | | |
|---|---|---|---|---|---|---|---|---|---|---|
| ATOM | 204 | HH11 | ARG | 12 | 28.745 | 1.957 | 23.347 | 0.4478 | 1.3000 | H |
| ATOM | 205 | HH12 | ARG | 12 | 27.150 | 2.568 | 23.682 | 0.4478 | 1.3000 | H |
| ATOM | 206 | NH2 | ARG | 12 | 26.172 | 0.380 | 24.409 | -0.8627 | 1.5500 | N |
| ATOM | 207 | HH21 | ARG | 12 | 25.885 | -0.535 | 24.722 | 0.4478 | 1.3000 | H |
| ATOM | 208 | HH22 | ARG | 12 | 25.529 | 1.154 | 24.483 | 0.4478 | 1.3000 | H |
| ATOM | 209 | C | ARG | 12 | 33.824 | -3.123 | 22.826 | 0.7341 | 1.7000 | C |
| ATOM | 210 | O | ARG | 12 | 33.884 | -3.865 | 21.840 | -0.5894 | 1.5000 | O |
| ATOM | 211 | N | ALA | 13 | 34.910 | -2.490 | 23.300 | -0.4157 | 1.5500 | N |
| ATOM | 212 | H | ALA | 13 | 34.847 | -1.901 | 24.118 | 0.2719 | 1.3000 | H |
| ATOM | 213 | CA | ALA | 13 | 36.206 | -2.660 | 22.637 | 0.0337 | 1.7000 | C |
| ATOM | 214 | HA | ALA | 13 | 36.107 | -2.389 | 21.584 | 0.0823 | 1.2000 | H |
| ATOM | 215 | CB | ALA | 13 | 37.251 | -1.755 | 23.273 | -0.1825 | 1.7000 | C |
| ATOM | 216 | HB1 | ALA | 13 | 38.208 | -1.885 | 22.767 | 0.0603 | 1.2000 | H |
| ATOM | 217 | HB2 | ALA | 13 | 36.940 | -0.713 | 23.179 | 0.0603 | 1.2000 | H |
| ATOM | 218 | HB3 | ALA | 13 | 37.368 | -1.995 | 24.331 | 0.0603 | 1.2000 | H |
| ATOM | 219 | C | ALA | 13 | 36.687 | -4.100 | 22.729 | 0.5973 | 1.7000 | C |
| ATOM | 220 | O | ALA | 13 | 37.189 | -4.674 | 21.757 | -0.5679 | 1.5000 | O |
| ATOM | 221 | N | LEU | 14 | 36.496 | -4.682 | 23.910 | -0.4157 | 1.5500 | N |
| ATOM | 222 | H | LEU | 14 | 36.064 | -4.142 | 24.642 | 0.2719 | 1.3000 | H |
| ATOM | 223 | CA | LEU | 14 | 36.908 | -6.028 | 24.243 | -0.0518 | 1.7000 | C |
| ATOM | 224 | HA | LEU | 14 | 37.963 | -6.140 | 24.019 | 0.0922 | 1.2000 | H |
| ATOM | 225 | CB | LEU | 14 | 36.682 | -6.222 | 25.768 | -0.1102 | 1.7000 | C |
| ATOM | 226 | HB2 | LEU | 14 | 37.277 | -5.458 | 26.273 | 0.0457 | 1.2000 | H |
| ATOM | 227 | HB3 | LEU | 14 | 35.629 | -6.019 | 25.956 | 0.0457 | 1.2000 | H |
| ATOM | 228 | CG | LEU | 14 | 36.954 | -7.540 | 26.478 | 0.3531 | 1.7000 | C |
| ATOM | 229 | HG | LEU | 14 | 36.300 | -8.316 | 26.084 | -0.0361 | 1.2000 | H |
| ATOM | 230 | CD1 | LEU | 14 | 38.372 | -7.915 | 26.303 | -0.4121 | 1.7000 | C |
| ATOM | 231 | HD11 | LEU | 14 | 38.609 | -8.789 | 26.905 | 0.1000 | 1.2000 | H |
| ATOM | 232 | HD12 | LEU | 14 | 38.567 | -8.143 | 25.257 | 0.1000 | 1.2000 | H |
| ATOM | 233 | HD13 | LEU | 14 | 39.018 | -7.095 | 26.599 | 0.1000 | 1.2000 | H |
| ATOM | 234 | CD2 | LEU | 14 | 36.655 | -7.345 | 27.984 | -0.4121 | 1.7000 | C |
| ATOM | 235 | HD21 | LEU | 14 | 36.838 | -8.275 | 28.520 | 0.1000 | 1.2000 | H |
| ATOM | 236 | HD22 | LEU | 14 | 37.292 | -6.567 | 28.405 | 0.1000 | 1.2000 | H |
| ATOM | 237 | HD23 | LEU | 14 | 35.612 | -7.062 | 28.123 | 0.1000 | 1.2000 | H |

| | | | | | | | | | | |
|---|---|---|---|---|---|---|---|---|---|---|
| ATOM | 238 | C | LEU | 14 | 36.154 | -7.005 | 23.395 | 0.5973 | 1.7000 | C |
| ATOM | 239 | O | LEU | 14 | 36.763 | -7.778 | 22.666 | -0.5679 | 1.5000 | O |
| ATOM | 240 | N | ILE | 15 | 34.839 | -6.903 | 23.363 | -0.4157 | 1.5500 | N |
| ATOM | 241 | H | ILE | 15 | 34.357 | -6.221 | 23.932 | 0.2719 | 1.3000 | H |
| ATOM | 242 | CA | ILE | 15 | 34.078 | -7.849 | 22.578 | -0.0597 | 1.7000 | C |
| ATOM | 243 | HA | ILE | 15 | 34.320 | -8.855 | 22.929 | 0.0869 | 1.2000 | H |
| ATOM | 244 | CB | ILE | 15 | 32.590 | -7.633 | 22.741 | 0.1303 | 1.7000 | C |
| ATOM | 245 | HB | ILE | 15 | 32.368 | -6.583 | 22.539 | 0.0187 | 1.2000 | H |
| ATOM | 246 | CG2 | ILE | 15 | 31.906 | -8.449 | 21.776 | -0.3204 | 1.7000 | C |
| ATOM | 247 | HG21 | ILE | 15 | 30.832 | -8.448 | 21.920 | 0.0882 | 1.2000 | H |
| ATOM | 248 | HG22 | ILE | 15 | 32.017 | -8.072 | 20.761 | 0.0882 | 1.2000 | H |
| ATOM | 249 | HG23 | ILE | 15 | 32.256 | -9.478 | 21.830 | 0.0882 | 1.2000 | H |
| ATOM | 250 | CG1 | ILE | 15 | 32.143 | -7.984 | 24.108 | -0.0430 | 1.7000 | C |
| ATOM | 251 | HG12 | ILE | 15 | 32.221 | -9.056 | 24.268 | 0.0236 | 1.2000 | H |
| ATOM | 252 | HG13 | ILE | 15 | 32.776 | -7.516 | 24.858 | 0.0236 | 1.2000 | H |
| ATOM | 253 | CD1 | ILE | 15 | 30.751 | -7.546 | 24.334 | -0.0660 | 1.7000 | C |
| ATOM | 254 | HD11 | ILE | 15 | 30.526 | -7.598 | 25.397 | 0.0186 | 1.2000 | H |
| ATOM | 255 | HD12 | ILE | 15 | 30.598 | -6.523 | 23.991 | 0.0186 | 1.2000 | H |
| ATOM | 256 | HD13 | ILE | 15 | 30.062 | -8.208 | 23.826 | 0.0186 | 1.2000 | H |
| ATOM | 257 | C | ILE | 15 | 34.382 | -7.796 | 21.089 | 0.5973 | 1.7000 | C |
| ATOM | 258 | O | ILE | 15 | 34.512 | -8.841 | 20.443 | -0.5679 | 1.5000 | O |
| ATOM | 259 | N | LYS | 16 | 34.476 | -6.598 | 20.510 | -0.3479 | 1.5500 | N |
| ATOM | 260 | H | LYS | 16 | 34.366 | -5.748 | 21.046 | 0.2747 | 1.3000 | H |
| ATOM | 261 | CA | LYS | 16 | 34.745 | -6.525 | 19.081 | -0.2400 | 1.7000 | C |
| ATOM | 262 | HA | LYS | 16 | 33.995 | -7.111 | 18.545 | 0.1426 | 1.2000 | H |
| ATOM | 263 | CB | LYS | 16 | 34.646 | -5.076 | 18.617 | -0.0094 | 1.7000 | C |
| ATOM | 264 | HB2 | LYS | 16 | 35.277 | -4.454 | 19.256 | 0.0362 | 1.2000 | H |
| ATOM | 265 | HB3 | LYS | 16 | 35.036 | -5.013 | 17.599 | 0.0362 | 1.2000 | H |
| ATOM | 266 | CG | LYS | 16 | 33.233 | -4.522 | 18.609 | 0.0187 | 1.7000 | C |
| ATOM | 267 | HG2 | LYS | 16 | 32.633 | -5.102 | 17.906 | 0.0103 | 1.2000 | H |
| ATOM | 268 | HG3 | LYS | 16 | 32.789 | -4.614 | 19.600 | 0.0103 | 1.2000 | H |
| ATOM | 269 | CD | LYS | 16 | 33.216 | -3.067 | 18.186 | -0.0479 | 1.7000 | C |
| ATOM | 270 | HD2 | LYS | 16 | 33.832 | -2.486 | 18.878 | 0.0621 | 1.2000 | H |
| ATOM | 271 | HD3 | LYS | 16 | 33.634 | -2.980 | 17.181 | 0.0621 | 1.2000 | H |

| | | | | | | | | | | |
|---|---|---|---|---|---|---|---|---|---|---|
| ATOM | 272 | CE | LYS | 16 | 31.805 | -2.501 | 18.188 | -0.0143 | 1.7000 | C |
| ATOM | 273 | HE2 | LYS | 16 | 31.195 | -3.076 | 17.489 | 0.1135 | 1.2000 | H |
| ATOM | 274 | HE3 | LYS | 16 | 31.382 | -2.605 | 19.188 | 0.1135 | 1.2000 | H |
| ATOM | 275 | NZ | LYS | 16 | 31.788 | -1.063 | 17.789 | -0.3854 | 1.5500 | N |
| ATOM | 276 | HZ1 | LYS | 16 | 32.192 | -0.961 | 16.868 | 0.3400 | 1.3000 | H |
| ATOM | 277 | HZ2 | LYS | 16 | 30.837 | -0.723 | 17.769 | 0.3400 | 1.3000 | H |
| ATOM | 278 | HZ3 | LYS | 16 | 32.328 | -0.516 | 18.445 | 0.3400 | 1.3000 | H |
| ATOM | 279 | C | LYS | 16 | 36.119 | -7.091 | 18.721 | 0.7341 | 1.7000 | C |
| ATOM | 280 | O | LYS | 16 | 36.260 | -7.830 | 17.735 | -0.5894 | 1.5000 | O |
| ATOM | 281 | N | ARG | 17 | 37.133 | -6.785 | 19.539 | -0.3479 | 1.5500 | N |
| ATOM | 282 | H | ARG | 17 | 36.977 | -6.201 | 20.348 | 0.2747 | 1.3000 | H |
| ATOM | 283 | CA | ARG | 17 | 38.473 | -7.283 | 19.276 | -0.2637 | 1.7000 | C |
| ATOM | 284 | HA | ARG | 17 | 38.744 | -7.083 | 18.238 | 0.1560 | 1.2000 | H |
| ATOM | 285 | CB | ARG | 17 | 39.456 | -6.577 | 20.186 | -0.0007 | 1.7000 | C |
| ATOM | 286 | HB2 | ARG | 17 | 39.095 | -6.654 | 21.211 | 0.0327 | 1.2000 | H |
| ATOM | 287 | HB3 | ARG | 17 | 40.420 | -7.082 | 20.124 | 0.0327 | 1.2000 | H |
| ATOM | 288 | CG | ARG | 17 | 39.669 | -5.109 | 19.851 | 0.0390 | 1.7000 | C |
| ATOM | 289 | HG2 | ARG | 17 | 40.142 | -5.048 | 18.870 | 0.0285 | 1.2000 | H |
| ATOM | 290 | HG3 | ARG | 17 | 38.710 | -4.595 | 19.784 | 0.0285 | 1.2000 | H |
| ATOM | 291 | CD | ARG | 17 | 40.535 | -4.400 | 20.844 | 0.0486 | 1.7000 | C |
| ATOM | 292 | HD2 | ARG | 17 | 40.078 | -4.472 | 21.832 | 0.0687 | 1.2000 | H |
| ATOM | 293 | HD3 | ARG | 17 | 41.516 | -4.877 | 20.870 | 0.0687 | 1.2000 | H |
| ATOM | 294 | NE | ARG | 17 | 40.686 | -2.992 | 20.485 | -0.5295 | 1.5500 | N |
| ATOM | 295 | HE | ARG | 17 | 40.193 | -2.688 | 19.659 | 0.3456 | 1.3000 | H |
| ATOM | 296 | CZ | ARG | 17 | 41.418 | -2.074 | 21.151 | 0.8076 | 1.7000 | C |
| ATOM | 297 | NH1 | ARG | 17 | 42.097 | -2.390 | 22.224 | -0.8627 | 1.5500 | N |
| ATOM | 298 | HH11 | ARG | 17 | 42.675 | -1.699 | 22.677 | 0.4478 | 1.3000 | H |
| ATOM | 299 | HH12 | ARG | 17 | 42.063 | -3.333 | 22.579 | 0.4478 | 1.3000 | H |
| ATOM | 300 | NH2 | ARG | 17 | 41.457 | -0.832 | 20.707 | -0.8627 | 1.5500 | N |
| ATOM | 301 | HH21 | ARG | 17 | 41.001 | -0.590 | 19.842 | 0.4478 | 1.3000 | H |
| ATOM | 302 | HH22 | ARG | 17 | 41.956 | -0.130 | 21.229 | 0.4478 | 1.3000 | H |
| ATOM | 303 | C | ARG | 17 | 38.504 | -8.787 | 19.489 | 0.7341 | 1.7000 | C |
| ATOM | 304 | O | ARG | 17 | 39.096 | -9.535 | 18.707 | -0.5894 | 1.5000 | O |
| ATOM | 305 | N | ILE | 18 | 37.808 | -9.244 | 20.514 | -0.4157 | 1.5500 | N |

| | | | | | | | | | | |
|---|---|---|---|---|---|---|---|---|---|---|
| ATOM | 306 | H | ILE | 18 | 37.311 | -8.609 | 21.117 | 0.2719 | 1.3000 | H |
| ATOM | 307 | CA | ILE | 18 | 37.754 | -10.654 | 20.783 | -0.0597 | 1.7000 | C |
| ATOM | 308 | HA | ILE | 18 | 38.779 | -11.014 | 20.875 | 0.0869 | 1.2000 | H |
| ATOM | 309 | CB | ILE | 18 | 37.038 | -10.929 | 22.092 | 0.1303 | 1.7000 | C |
| ATOM | 310 | HB | ILE | 18 | 36.101 | -10.369 | 22.076 | 0.0187 | 1.2000 | H |
| ATOM | 311 | CG2 | ILE | 18 | 36.709 | -12.281 | 22.208 | -0.3204 | 1.7000 | C |
| ATOM | 312 | HG21 | ILE | 18 | 36.052 | -12.444 | 23.054 | 0.0882 | 1.2000 | H |
| ATOM | 313 | HG22 | ILE | 18 | 36.110 | -12.668 | 21.404 | 0.0882 | 1.2000 | H |
| ATOM | 314 | HG23 | ILE | 18 | 37.601 | -12.890 | 22.303 | 0.0882 | 1.2000 | H |
| ATOM | 315 | CG1 | ILE | 18 | 37.837 | -10.502 | 23.256 | -0.0430 | 1.7000 | C |
| ATOM | 316 | HG12 | ILE | 18 | 38.678 | -11.180 | 23.401 | 0.0236 | 1.2000 | H |
| ATOM | 317 | HG13 | ILE | 18 | 38.279 | -9.522 | 23.111 | 0.0236 | 1.2000 | H |
| ATOM | 318 | CD1 | ILE | 18 | 37.029 | -10.502 | 24.460 | -0.0660 | 1.7000 | C |
| ATOM | 319 | HD11 | ILE | 18 | 37.645 | -10.168 | 25.284 | 0.0186 | 1.2000 | H |
| ATOM | 320 | HD12 | ILE | 18 | 36.168 | -9.844 | 24.351 | 0.0186 | 1.2000 | H |
| ATOM | 321 | HD13 | ILE | 18 | 36.696 | -11.483 | 24.749 | 0.0186 | 1.2000 | H |
| ATOM | 322 | C | ILE | 18 | 37.091 | -11.455 | 19.692 | 0.5973 | 1.7000 | C |
| ATOM | 323 | O | ILE | 18 | 37.658 | -12.448 | 19.258 | -0.5679 | 1.5000 | O |
| ATOM | 324 | N | GLN | 19 | 35.931 | -11.031 | 19.192 | -0.4157 | 1.5500 | N |
| ATOM | 325 | H | GLN | 19 | 35.477 | -10.203 | 19.539 | 0.2719 | 1.3000 | H |
| ATOM | 326 | CA | GLN | 19 | 35.301 | -11.832 | 18.149 | -0.0031 | 1.7000 | C |
| ATOM | 327 | HA | GLN | 19 | 35.137 | -12.838 | 18.531 | 0.0850 | 1.2000 | H |
| ATOM | 328 | CB | GLN | 19 | 33.949 | -11.254 | 17.737 | -0.0036 | 1.7000 | C |
| ATOM | 329 | HB2 | GLN | 19 | 33.352 | -11.126 | 18.636 | 0.0171 | 1.2000 | H |
| ATOM | 330 | HB3 | GLN | 19 | 34.104 | -10.268 | 17.295 | 0.0171 | 1.2000 | H |
| ATOM | 331 | CG | GLN | 19 | 33.140 | -12.137 | 16.737 | -0.0645 | 1.7000 | C |
| ATOM | 332 | HG2 | GLN | 19 | 33.699 | -12.240 | 15.806 | 0.0352 | 1.2000 | H |
| ATOM | 333 | HG3 | GLN | 19 | 33.015 | -13.134 | 17.159 | 0.0352 | 1.2000 | H |
| ATOM | 334 | CD | GLN | 19 | 31.754 | -11.544 | 16.402 | 0.6951 | 1.7000 | C |
| ATOM | 335 | OE1 | GLN | 19 | 31.691 | -10.366 | 16.035 | -0.6086 | 1.5000 | O |
| ATOM | 336 | NE2 | GLN | 19 | 30.655 | -12.321 | 16.519 | -0.9407 | 1.5500 | N |
| ATOM | 337 | HE21 | GLN | 19 | 30.743 | -13.286 | 16.807 | 0.4251 | 1.3000 | H |
| ATOM | 338 | HE22 | GLN | 19 | 29.753 | -11.943 | 16.271 | 0.4251 | 1.3000 | H |
| ATOM | 339 | C | GLN | 19 | 36.239 | -11.943 | 16.961 | 0.5973 | 1.7000 | C |

| | | | | | | | | | | |
|---|---|---|---|---|---|---|---|---|---|---|
| ATOM | 340 | O | GLN | 19 | 36.403 | -13.024 | 16.396 | -0.5679 | 1.5000 | O |
| ATOM | 341 | N | ALA | 20 | 36.918 | -10.844 | 16.615 | -0.4157 | 1.5500 | N |
| ATOM | 342 | H | ALA | 20 | 36.778 | -9.979 | 17.117 | 0.2719 | 1.3000 | H |
| ATOM | 343 | CA | ALA | 20 | 37.860 | -10.860 | 15.503 | 0.0337 | 1.7000 | C |
| ATOM | 344 | HA | ALA | 20 | 37.321 | -11.125 | 14.592 | 0.0823 | 1.2000 | H |
| ATOM | 345 | CB | ALA | 20 | 38.470 | -9.482 | 15.319 | -0.1825 | 1.7000 | C |
| ATOM | 346 | HB1 | ALA | 20 | 39.136 | -9.489 | 14.458 | 0.0603 | 1.2000 | H |
| ATOM | 347 | HB2 | ALA | 20 | 37.684 | -8.750 | 15.145 | 0.0603 | 1.2000 | H |
| ATOM | 348 | HB3 | ALA | 20 | 39.042 | -9.190 | 16.195 | 0.0603 | 1.2000 | H |
| ATOM | 349 | C | ALA | 20 | 38.971 | -11.894 | 15.729 | 0.5973 | 1.7000 | C |
| ATOM | 350 | O | ALA | 20 | 39.444 | -12.531 | 14.785 | -0.5679 | 1.5000 | O |
| ATOM | 351 | N | MET | 21 | 39.371 | -12.072 | 16.989 | -0.4157 | 1.5500 | N |
| ATOM | 352 | H | MET | 21 | 38.937 | -11.533 | 17.723 | 0.2719 | 1.3000 | H |
| ATOM | 353 | CA | MET | 21 | 40.428 | -13.002 | 17.367 | -0.0237 | 1.7000 | C |
| ATOM | 354 | HA | MET | 21 | 41.138 | -13.060 | 16.541 | 0.0880 | 1.2000 | H |
| ATOM | 355 | CB | MET | 21 | 41.190 | -12.446 | 18.569 | 0.0342 | 1.7000 | C |
| ATOM | 356 | HB2 | MET | 21 | 40.486 | -12.239 | 19.375 | 0.0241 | 1.2000 | H |
| ATOM | 357 | HB3 | MET | 21 | 41.901 | -13.193 | 18.925 | 0.0241 | 1.2000 | H |
| ATOM | 358 | CG | MET | 21 | 41.993 | -11.179 | 18.250 | 0.0018 | 1.7000 | C |
| ATOM | 359 | HG2 | MET | 21 | 42.639 | -11.392 | 17.398 | 0.0440 | 1.2000 | H |
| ATOM | 360 | HG3 | MET | 21 | 41.322 | -10.369 | 17.968 | 0.0440 | 1.2000 | H |
| ATOM | 361 | SD | MET | 21 | 43.039 | -10.638 | 19.604 | -0.2737 | 1.8000 | S |
| ATOM | 362 | CE | MET | 21 | 41.854 | -10.020 | 20.756 | -0.0536 | 1.7000 | C |
| ATOM | 363 | HE1 | MET | 21 | 42.378 | -9.601 | 21.614 | 0.0684 | 1.2000 | H |
| ATOM | 364 | HE2 | MET | 21 | 41.260 | -9.241 | 20.282 | 0.0684 | 1.2000 | H |
| ATOM | 365 | HE3 | MET | 21 | 41.203 | -10.827 | 21.089 | 0.0684 | 1.2000 | H |
| ATOM | 366 | C | MET | 21 | 39.973 | -14.448 | 17.639 | 0.5973 | 1.7000 | C |
| ATOM | 367 | O | MET | 21 | 40.808 | -15.279 | 18.011 | -0.5679 | 1.5000 | O |
| ATOM | 368 | N | ILE | 22 | 38.681 | -14.771 | 17.449 | -0.4157 | 1.5500 | N |
| ATOM | 369 | H | ILE | 22 | 38.018 | -14.071 | 17.147 | 0.2719 | 1.3000 | H |
| ATOM | 370 | CA | ILE | 22 | 38.209 | -16.145 | 17.679 | -0.0597 | 1.7000 | C |
| ATOM | 371 | HA | ILE | 22 | 39.046 | -16.793 | 17.943 | 0.0869 | 1.2000 | H |
| ATOM | 372 | CB | ILE | 22 | 37.219 | -16.215 | 18.845 | 0.1303 | 1.7000 | C |
| ATOM | 373 | HB | ILE | 22 | 36.343 | -15.616 | 18.589 | 0.0187 | 1.2000 | H |

| | | | | | | | | | | |
|---|---|---|---|---|---|---|---|---|---|---|
| ATOM | 374 | CG2 | ILE | 22 | 36.780 | -17.673 | 19.086 | -0.3204 | 1.7000 | C |
| ATOM | 375 | HG21 | ILE | 22 | 35.896 | -17.687 | 19.721 | 0.0882 | 1.2000 | H |
| ATOM | 376 | HG22 | ILE | 22 | 36.544 | -18.310 | 18.244 | 0.0882 | 1.2000 | H |
| ATOM | 377 | HG23 | ILE | 22 | 37.585 | -18.193 | 19.600 | 0.0882 | 1.2000 | H |
| ATOM | 378 | CG1 | ILE | 22 | 37.805 | -15.619 | 20.064 | -0.0430 | 1.7000 | C |
| ATOM | 379 | HG12 | ILE | 22 | 38.666 | -16.205 | 20.377 | 0.0236 | 1.2000 | H |
| ATOM | 380 | HG13 | ILE | 22 | 38.183 | -14.624 | 19.892 | 0.0236 | 1.2000 | H |
| ATOM | 381 | CD1 | ILE | 22 | 36.830 | -15.535 | 21.152 | -0.0660 | 1.7000 | C |
| ATOM | 382 | HD11 | ILE | 22 | 37.312 | -15.077 | 22.009 | 0.0186 | 1.2000 | H |
| ATOM | 383 | HD12 | ILE | 22 | 35.985 | -14.927 | 20.845 | 0.0186 | 1.2000 | H |
| ATOM | 384 | HD13 | ILE | 22 | 36.469 | -16.511 | 21.461 | 0.0186 | 1.2000 | H |
| ATOM | 385 | C | ILE | 22 | 37.481 | -16.801 | 16.508 | 0.5973 | 1.7000 | C |
| ATOM | 386 | O | ILE | 22 | 36.256 | -16.899 | 16.543 | -0.5679 | 1.5000 | O |
| ATOM | 387 | N | PRO | 23 | 38.183 | -17.298 | 15.479 | -0.2548 | 1.5500 | N |
| ATOM | 388 | CD | PRO | 23 | 39.637 | -17.089 | 15.383 | 0.0192 | 1.7000 | C |
| ATOM | 389 | HD2 | PRO | 23 | 40.164 | -17.804 | 16.016 | 0.0391 | 1.2000 | H |
| ATOM | 390 | HD3 | PRO | 23 | 39.934 | -16.067 | 15.600 | 0.0391 | 1.2000 | H |
| ATOM | 391 | CG | PRO | 23 | 39.904 | -17.368 | 13.928 | 0.0189 | 1.7000 | C |
| ATOM | 392 | HG2 | PRO | 23 | 40.914 | -17.749 | 13.774 | 0.0213 | 1.2000 | H |
| ATOM | 393 | HG3 | PRO | 23 | 39.758 | -16.452 | 13.352 | 0.0213 | 1.2000 | H |
| ATOM | 394 | CB | PRO | 23 | 38.869 | -18.374 | 13.532 | -0.0070 | 1.7000 | C |
| ATOM | 395 | HB2 | PRO | 23 | 39.223 | -19.367 | 13.816 | 0.0253 | 1.2000 | H |
| ATOM | 396 | HB3 | PRO | 23 | 38.671 | -18.343 | 12.460 | 0.0253 | 1.2000 | H |
| ATOM | 397 | CA | PRO | 23 | 37.617 | -18.002 | 14.343 | -0.0266 | 1.7000 | C |
| ATOM | 398 | HA | PRO | 23 | 36.991 | -17.316 | 13.771 | 0.0641 | 1.2000 | H |
| ATOM | 399 | C | PRO | 23 | 36.838 | -19.282 | 14.671 | 0.5896 | 1.7000 | C |
| ATOM | 400 | O | PRO | 23 | 36.009 | -19.702 | 13.861 | -0.5748 | 1.5000 | O |
| ATOM | 401 | N | LYS | 24 | 37.116 | -19.930 | 15.821 | -0.3479 | 1.5500 | N |
| ATOM | 402 | H | LYS | 24 | 37.776 | -19.556 | 16.485 | 0.2747 | 1.3000 | H |
| ATOM | 403 | CA | LYS | 24 | 36.434 | -21.202 | 16.084 | -0.2400 | 1.7000 | C |
| ATOM | 404 | HA | LYS | 24 | 35.622 | -21.349 | 15.372 | 0.1426 | 1.2000 | H |
| ATOM | 405 | CB | LYS | 24 | 37.423 | -22.351 | 15.849 | -0.0094 | 1.7000 | C |
| ATOM | 406 | HB2 | LYS | 24 | 38.306 | -22.195 | 16.472 | 0.0362 | 1.2000 | H |
| ATOM | 407 | HB3 | LYS | 24 | 36.951 | -23.282 | 16.170 | 0.0362 | 1.2000 | H |

| | | | | | | | | | | |
|---|---|---|---|---|---|---|---|---|---|---|
| ATOM | 408 | CG | LYS | 24 | 37.867 | -22.540 | 14.393 | 0.0187 | 1.7000 | C |
| ATOM | 409 | HG2 | LYS | 24 | 36.986 | -22.715 | 13.772 | 0.0103 | 1.2000 | H |
| ATOM | 410 | HG3 | LYS | 24 | 38.368 | -21.637 | 14.044 | 0.0103 | 1.2000 | H |
| ATOM | 411 | CD | LYS | 24 | 38.824 | -23.713 | 14.244 | -0.0479 | 1.7000 | C |
| ATOM | 412 | HD2 | LYS | 24 | 39.705 | -23.538 | 14.865 | 0.0621 | 1.2000 | H |
| ATOM | 413 | HD3 | LYS | 24 | 38.330 | -24.627 | 14.580 | 0.0621 | 1.2000 | H |
| ATOM | 414 | CE | LYS | 24 | 39.262 | -23.886 | 12.793 | -0.0143 | 1.7000 | C |
| ATOM | 415 | HE2 | LYS | 24 | 38.380 | -24.083 | 12.179 | 0.1135 | 1.2000 | H |
| ATOM | 416 | HE3 | LYS | 24 | 39.725 | -22.957 | 12.451 | 0.1135 | 1.2000 | H |
| ATOM | 417 | NZ | LYS | 24 | 40.233 | -25.004 | 12.634 | -0.3854 | 1.5500 | N |
| ATOM | 418 | HZ1 | LYS | 24 | 41.056 | -24.827 | 13.193 | 0.3400 | 1.3000 | H |
| ATOM | 419 | HZ2 | LYS | 24 | 39.812 | -25.870 | 12.938 | 0.3400 | 1.3000 | H |
| ATOM | 420 | HZ3 | LYS | 24 | 40.501 | -25.089 | 11.664 | 0.3400 | 1.3000 | H |
| ATOM | 421 | C | LYS | 24 | 35.771 | -21.454 | 17.447 | 0.7341 | 1.7000 | C |
| ATOM | 422 | O | LYS | 24 | 34.585 | -21.778 | 17.486 | -0.5894 | 1.5000 | O |
| ATOM | 423 | N | GLY | 25 | 36.513 | -21.386 | 18.564 | -0.4157 | 1.5500 | N |
| ATOM | 424 | H | GLY | 25 | 37.498 | -21.173 | 18.544 | 0.2719 | 1.3000 | H |
| ATOM | 425 | CA | GLY | 25 | 35.858 | -21.826 | 19.803 | -0.0252 | 1.7000 | C |
| ATOM | 426 | HA2 | GLY | 25 | 34.875 | -21.360 | 19.844 | 0.0698 | 1.2000 | H |
| ATOM | 427 | HA3 | GLY | 25 | 35.703 | -22.904 | 19.737 | 0.0698 | 1.2000 | H |
| ATOM | 428 | C | GLY | 25 | 36.507 | -21.530 | 21.166 | 0.5973 | 1.7000 | C |
| ATOM | 429 | O | GLY | 25 | 36.252 | -22.267 | 22.117 | -0.5679 | 1.5000 | O |
| ATOM | 430 | N | ALA | 26 | 37.351 | -20.513 | 21.293 | -0.4157 | 1.5500 | N |
| ATOM | 431 | H | ALA | 26 | 37.546 | -19.888 | 20.525 | 0.2719 | 1.3000 | H |
| ATOM | 432 | CA | ALA | 26 | 37.976 | -20.270 | 22.605 | 0.0337 | 1.7000 | C |
| ATOM | 433 | HA | ALA | 26 | 37.366 | -20.691 | 23.406 | 0.0823 | 1.2000 | H |
| ATOM | 434 | CB | ALA | 26 | 39.341 | -20.926 | 22.664 | -0.1825 | 1.7000 | C |
| ATOM | 435 | HB1 | ALA | 26 | 39.792 | -20.756 | 23.642 | 0.0603 | 1.2000 | H |
| ATOM | 436 | HB2 | ALA | 26 | 39.240 | -22.002 | 22.507 | 0.0603 | 1.2000 | H |
| ATOM | 437 | HB3 | ALA | 26 | 39.990 | -20.508 | 21.892 | 0.0603 | 1.2000 | H |
| ATOM | 438 | C | ALA | 26 | 38.107 | -18.789 | 22.882 | 0.5973 | 1.7000 | C |
| ATOM | 439 | O | ALA | 26 | 38.419 | -18.024 | 21.984 | -0.5679 | 1.5000 | O |
| ATOM | 440 | N | LEU | 27 | 37.928 | -18.407 | 24.149 | -0.4157 | 1.5500 | N |
| ATOM | 441 | H | LEU | 27 | 37.695 | -19.100 | 24.845 | 0.2719 | 1.3000 | H |

| | | | | | | | | | | |
|---|---|---|---|---|---|---|---|---|---|---|
| ATOM | 442 | CA | LEU | 27 | 37.925 | -17.002 | 24.557 | -0.0518 | 1.7000 | C |
| ATOM | 443 | HA | LEU | 27 | 37.961 | -16.411 | 23.645 | 0.0922 | 1.2000 | H |
| ATOM | 444 | CB | LEU | 27 | 36.575 | -16.719 | 25.205 | -0.1102 | 1.7000 | C |
| ATOM | 445 | HB2 | LEU | 27 | 35.809 | -16.923 | 24.470 | 0.0457 | 1.2000 | H |
| ATOM | 446 | HB3 | LEU | 27 | 36.436 | -17.434 | 26.019 | 0.0457 | 1.2000 | H |
| ATOM | 447 | CG | LEU | 27 | 36.379 | -15.381 | 25.754 | 0.3531 | 1.7000 | C |
| ATOM | 448 | HG | LEU | 27 | 37.013 | -15.210 | 26.628 | -0.0361 | 1.2000 | H |
| ATOM | 449 | CD1 | LEU | 27 | 36.623 | -14.433 | 24.752 | -0.4121 | 1.7000 | C |
| ATOM | 450 | HD11 | LEU | 27 | 36.427 | -13.459 | 25.183 | 0.1000 | 1.2000 | H |
| ATOM | 451 | HD12 | LEU | 27 | 37.608 | -14.382 | 24.328 | 0.1000 | 1.2000 | H |
| ATOM | 452 | HD13 | LEU | 27 | 35.935 | -14.576 | 23.915 | 0.1000 | 1.2000 | H |
| ATOM | 453 | CD2 | LEU | 27 | 34.948 | -15.206 | 26.134 | -0.4121 | 1.7000 | C |
| ATOM | 454 | HD21 | LEU | 27 | 34.812 | -14.253 | 26.645 | 0.1000 | 1.2000 | H |
| ATOM | 455 | HD22 | LEU | 27 | 34.288 | -15.226 | 25.266 | 0.1000 | 1.2000 | H |
| ATOM | 456 | HD23 | LEU | 27 | 34.653 | -16.004 | 26.815 | 0.1000 | 1.2000 | H |
| ATOM | 457 | C | LEU | 27 | 39.038 | -16.432 | 25.439 | 0.5973 | 1.7000 | C |
| ATOM | 458 | O | LEU | 27 | 39.426 | -15.276 | 25.258 | -0.5679 | 1.5000 | O |
| ATOM | 459 | N | ALA | 28 | 39.531 | -17.182 | 26.416 | -0.4157 | 1.5500 | N |
| ATOM | 460 | H | ALA | 28 | 39.197 | -18.122 | 26.564 | 0.2719 | 1.3000 | H |
| ATOM | 461 | CA | ALA | 28 | 40.446 | -16.581 | 27.395 | 0.0337 | 1.7000 | C |
| ATOM | 462 | HA | ALA | 28 | 39.897 | -15.820 | 27.953 | 0.0823 | 1.2000 | H |
| ATOM | 463 | CB | ALA | 28 | 40.911 | -17.639 | 28.374 | -0.1825 | 1.7000 | C |
| ATOM | 464 | HB1 | ALA | 28 | 41.554 | -17.191 | 29.130 | 0.0603 | 1.2000 | H |
| ATOM | 465 | HB2 | ALA | 28 | 40.053 | -18.088 | 28.873 | 0.0603 | 1.2000 | H |
| ATOM | 466 | HB3 | ALA | 28 | 41.469 | -18.420 | 27.858 | 0.0603 | 1.2000 | H |
| ATOM | 467 | C | ALA | 28 | 41.676 | -15.918 | 26.768 | 0.5973 | 1.7000 | C |
| ATOM | 468 | O | ALA | 28 | 42.123 | -14.871 | 27.245 | -0.5679 | 1.5000 | O |
| ATOM | 469 | N | VAL | 29 | 42.216 | -16.497 | 25.698 | -0.4157 | 1.5500 | N |
| ATOM | 470 | H | VAL | 29 | 41.830 | -17.349 | 25.320 | 0.2719 | 1.3000 | H |
| ATOM | 471 | CA | VAL | 29 | 43.388 | -15.901 | 25.075 | -0.0875 | 1.7000 | C |
| ATOM | 472 | HA | VAL | 29 | 44.152 | -15.739 | 25.837 | 0.0969 | 1.2000 | H |
| ATOM | 473 | CB | VAL | 29 | 43.962 | -16.813 | 23.991 | 0.2985 | 1.7000 | C |
| ATOM | 474 | HB | VAL | 29 | 43.180 | -17.081 | 23.278 | -0.0297 | 1.2000 | H |
| ATOM | 475 | CG1 | VAL | 29 | 45.066 | -16.072 | 23.246 | -0.3192 | 1.7000 | C |

| | | | | | | | | | | |
|---|---|---|---|---|---|---|---|---|---|---|
| ATOM | 476 | HG11 | VAL | 29 | 45.603 | -16.770 | 22.601 | 0.0791 | 1.2000 | H |
| ATOM | 477 | HG12 | VAL | 29 | 44.651 | -15.300 | 22.599 | 0.0791 | 1.2000 | H |
| ATOM | 478 | HG13 | VAL | 29 | 45.774 | -15.629 | 23.948 | 0.0791 | 1.2000 | H |
| ATOM | 479 | CG2 | VAL | 29 | 44.485 | -18.083 | 24.635 | -0.3192 | 1.7000 | C |
| ATOM | 480 | HG21 | VAL | 29 | 44.929 | -18.726 | 23.876 | 0.0791 | 1.2000 | H |
| ATOM | 481 | HG22 | VAL | 29 | 45.243 | -17.848 | 25.383 | 0.0791 | 1.2000 | H |
| ATOM | 482 | HG23 | VAL | 29 | 43.677 | -18.638 | 25.109 | 0.0791 | 1.2000 | H |
| ATOM | 483 | C | VAL | 29 | 43.045 | -14.563 | 24.456 | 0.5973 | 1.7000 | C |
| ATOM | 484 | O | VAL | 29 | 43.773 | -13.587 | 24.631 | -0.5679 | 1.5000 | O |
| ATOM | 485 | N | ALA | 30 | 41.928 | -14.517 | 23.731 | -0.4157 | 1.5500 | N |
| ATOM | 486 | H | ALA | 30 | 41.357 | -15.342 | 23.626 | 0.2719 | 1.3000 | H |
| ATOM | 487 | CA | ALA | 30 | 41.501 | -13.297 | 23.069 | 0.0337 | 1.7000 | C |
| ATOM | 488 | HA | ALA | 30 | 42.280 | -12.980 | 22.376 | 0.0823 | 1.2000 | H |
| ATOM | 489 | CB | ALA | 30 | 40.246 | -13.570 | 22.280 | -0.1825 | 1.7000 | C |
| ATOM | 490 | HB1 | ALA | 30 | 39.919 | -12.658 | 21.788 | 0.0603 | 1.2000 | H |
| ATOM | 491 | HB2 | ALA | 30 | 40.448 | -14.323 | 21.517 | 0.0603 | 1.2000 | H |
| ATOM | 492 | HB3 | ALA | 30 | 39.464 | -13.936 | 22.942 | 0.0603 | 1.2000 | H |
| ATOM | 493 | C | ALA | 30 | 41.287 | -12.191 | 24.096 | 0.5973 | 1.7000 | C |
| ATOM | 494 | O | ALA | 30 | 41.656 | -11.035 | 23.863 | -0.5679 | 1.5000 | O |
| ATOM | 495 | N | VAL | 31 | 40.752 | -12.556 | 25.265 | -0.4157 | 1.5500 | N |
| ATOM | 496 | H | VAL | 31 | 40.471 | -13.513 | 25.429 | 0.2719 | 1.3000 | H |
| ATOM | 497 | CA | VAL | 31 | 40.568 | -11.566 | 26.313 | -0.0875 | 1.7000 | C |
| ATOM | 498 | HA | VAL | 31 | 39.944 | -10.751 | 25.943 | 0.0969 | 1.2000 | H |
| ATOM | 499 | CB | VAL | 31 | 39.936 | -12.170 | 27.565 | 0.2985 | 1.7000 | C |
| ATOM | 500 | HB | VAL | 31 | 40.493 | -13.051 | 27.878 | -0.0297 | 1.2000 | H |
| ATOM | 501 | CG1 | VAL | 31 | 40.000 | -11.146 | 28.685 | -0.3192 | 1.7000 | C |
| ATOM | 502 | HG11 | VAL | 31 | 39.526 | -11.583 | 29.556 | 0.0791 | 1.2000 | H |
| ATOM | 503 | HG12 | VAL | 31 | 41.017 | -10.936 | 29.021 | 0.0791 | 1.2000 | H |
| ATOM | 504 | HG13 | VAL | 31 | 39.493 | -10.222 | 28.413 | 0.0791 | 1.2000 | H |
| ATOM | 505 | CG2 | VAL | 31 | 38.516 | -12.594 | 27.286 | -0.3192 | 1.7000 | C |
| ATOM | 506 | HG21 | VAL | 31 | 38.481 | -13.253 | 26.427 | 0.0791 | 1.2000 | H |
| ATOM | 507 | HG22 | VAL | 31 | 38.116 | -13.126 | 28.150 | 0.0791 | 1.2000 | H |
| ATOM | 508 | HG23 | VAL | 31 | 37.895 | -11.719 | 27.093 | 0.0791 | 1.2000 | H |
| ATOM | 509 | C | VAL | 31 | 41.889 | -11.012 | 26.758 | 0.5973 | 1.7000 | C |

| | | | | | | | | | | |
|---|---|---|---|---|---|---|---|---|---|---|
| ATOM | 510 | O | VAL | 31 | 42.059 | -9.794 | 26.831 | -0.5679 | 1.5000 | O |
| ATOM | 511 | N | ALA | 32 | 42.841 | -11.910 | 27.029 | -0.4157 | 1.5500 | N |
| ATOM | 512 | H | ALA | 32 | 42.650 | -12.898 | 26.947 | 0.2719 | 1.3000 | H |
| ATOM | 513 | CA | ALA | 32 | 44.142 | -11.485 | 27.497 | 0.0337 | 1.7000 | C |
| ATOM | 514 | HA | ALA | 32 | 44.017 | -10.945 | 28.438 | 0.0823 | 1.2000 | H |
| ATOM | 515 | CB | ALA | 32 | 45.029 | -12.690 | 27.744 | -0.1825 | 1.7000 | C |
| ATOM | 516 | HB1 | ALA | 32 | 45.990 | -12.359 | 28.138 | 0.0603 | 1.2000 | H |
| ATOM | 517 | HB2 | ALA | 32 | 44.557 | -13.348 | 28.475 | 0.0603 | 1.2000 | H |
| ATOM | 518 | HB3 | ALA | 32 | 45.194 | -13.243 | 26.820 | 0.0603 | 1.2000 | H |
| ATOM | 519 | C | ALA | 32 | 44.797 | -10.570 | 26.492 | 0.5973 | 1.7000 | C |
| ATOM | 520 | O | ALA | 32 | 45.382 | -9.554 | 26.863 | -0.5679 | 1.5000 | O |
| ATOM | 521 | N | GLN | 33 | 44.644 | -10.876 | 25.208 | -0.4157 | 1.5500 | N |
| ATOM | 522 | H | GLN | 33 | 44.122 | -11.693 | 24.927 | 0.2719 | 1.3000 | H |
| ATOM | 523 | CA | GLN | 33 | 45.260 | -10.040 | 24.201 | -0.0031 | 1.7000 | C |
| ATOM | 524 | HA | GLN | 33 | 46.331 | -9.972 | 24.408 | 0.0850 | 1.2000 | H |
| ATOM | 525 | CB | GLN | 33 | 45.086 | -10.695 | 22.829 | -0.0036 | 1.7000 | C |
| ATOM | 526 | HB2 | GLN | 33 | 44.038 | -10.955 | 22.687 | 0.0171 | 1.2000 | H |
| ATOM | 527 | HB3 | GLN | 33 | 45.365 | -9.977 | 22.055 | 0.0171 | 1.2000 | H |
| ATOM | 528 | CG | GLN | 33 | 45.952 | -11.955 | 22.650 | -0.0645 | 1.7000 | C |
| ATOM | 529 | HG2 | GLN | 33 | 46.998 | -11.645 | 22.654 | 0.0352 | 1.2000 | H |
| ATOM | 530 | HG3 | GLN | 33 | 45.819 | -12.626 | 23.496 | 0.0352 | 1.2000 | H |
| ATOM | 531 | CD | GLN | 33 | 45.662 | -12.747 | 21.375 | 0.6951 | 1.7000 | C |
| ATOM | 532 | OE1 | GLN | 33 | 44.520 | -13.078 | 21.049 | -0.6086 | 1.5000 | O |
| ATOM | 533 | NE2 | GLN | 33 | 46.724 | -13.056 | 20.631 | -0.9407 | 1.5500 | N |
| ATOM | 534 | HE21 | GLN | 33 | 47.646 | -12.768 | 20.916 | 0.4251 | 1.3000 | H |
| ATOM | 535 | HE22 | GLN | 33 | 46.584 | -13.555 | 19.766 | 0.4251 | 1.3000 | H |
| ATOM | 536 | C | GLN | 33 | 44.687 | -8.627 | 24.254 | 0.5973 | 1.7000 | C |
| ATOM | 537 | O | GLN | 33 | 45.444 | -7.653 | 24.253 | -0.5679 | 1.5000 | O |
| ATOM | 538 | N | VAL | 34 | 43.377 | -8.490 | 24.429 | -0.4157 | 1.5500 | N |
| ATOM | 539 | H | VAL | 34 | 42.763 | -9.292 | 24.479 | 0.2719 | 1.3000 | H |
| ATOM | 540 | CA | VAL | 34 | 42.830 | -7.144 | 24.503 | -0.0875 | 1.7000 | C |
| ATOM | 541 | HA | VAL | 34 | 43.157 | -6.581 | 23.627 | 0.0969 | 1.2000 | H |
| ATOM | 542 | CB | VAL | 34 | 41.313 | -7.120 | 24.563 | 0.2985 | 1.7000 | C |
| ATOM | 543 | HB | VAL | 34 | 40.971 | -7.744 | 25.387 | -0.0297 | 1.2000 | H |

| | | | | | | | | | | |
|---|---|---|---|---|---|---|---|---|---|---|
| ATOM | 544 | CG1 | VAL | 34 | 40.868 | -5.653 | 24.826 | -0.3192 | 1.7000 | C |
| ATOM | 545 | HG11 | VAL | 34 | 39.810 | -5.543 | 24.624 | 0.0791 | 1.2000 | H |
| ATOM | 546 | HG12 | VAL | 34 | 41.005 | -5.368 | 25.869 | 0.0791 | 1.2000 | H |
| ATOM | 547 | HG13 | VAL | 34 | 41.405 | -4.965 | 24.172 | 0.0791 | 1.2000 | H |
| ATOM | 548 | CG2 | VAL | 34 | 40.748 | -7.665 | 23.295 | -0.3192 | 1.7000 | C |
| ATOM | 549 | HG21 | VAL | 34 | 39.670 | -7.521 | 23.280 | 0.0791 | 1.2000 | H |
| ATOM | 550 | HG22 | VAL | 34 | 41.191 | -7.157 | 22.439 | 0.0791 | 1.2000 | H |
| ATOM | 551 | HG23 | VAL | 34 | 40.943 | -8.730 | 23.221 | 0.0791 | 1.2000 | H |
| ATOM | 552 | C | VAL | 34 | 43.280 | -6.410 | 25.736 | 0.5973 | 1.7000 | C |
| ATOM | 553 | O | VAL | 34 | 43.641 | -5.231 | 25.671 | -0.5679 | 1.5000 | O |
| ATOM | 554 | N | CYX | 35 | 43.232 | -7.103 | 26.867 | -0.4157 | 1.5500 | N |
| ATOM | 555 | H | CYX | 35 | 42.921 | -8.065 | 26.863 | 0.2719 | 1.3000 | H |
| ATOM | 556 | CA | CYX | 35 | 43.534 | -6.477 | 28.142 | 0.0429 | 1.7000 | C |
| ATOM | 557 | HA | CYX | 35 | 42.929 | -5.579 | 28.243 | 0.0766 | 1.2000 | H |
| ATOM | 558 | CB | CYX | 35 | 43.155 | -7.424 | 29.275 | -0.0790 | 1.7000 | C |
| ATOM | 559 | HB2 | CYX | 35 | 43.723 | -8.352 | 29.190 | 0.0910 | 1.2000 | H |
| ATOM | 560 | HB3 | CYX | 35 | 43.422 | -6.949 | 30.220 | 0.0910 | 1.2000 | H |
| ATOM | 561 | SG | CYX | 35 | 41.362 | -7.820 | 29.331 | -0.1081 | 1.8000 | S |
| ATOM | 562 | C | CYX | 35 | 45.003 | -6.071 | 28.238 | 0.5973 | 1.7000 | C |
| ATOM | 563 | O | CYX | 35 | 45.348 | -5.142 | 28.967 | -0.5679 | 1.5000 | O |
| ATOM | 564 | N | ARG | 36 | 45.868 | -6.731 | 27.461 | -0.3479 | 1.5500 | N |
| ATOM | 565 | H | ARG | 36 | 45.536 | -7.495 | 26.888 | 0.2747 | 1.3000 | H |
| ATOM | 566 | CA | ARG | 36 | 47.278 | -6.364 | 27.368 | -0.2637 | 1.7000 | C |
| ATOM | 567 | HA | ARG | 36 | 47.650 | -6.121 | 28.365 | 0.1560 | 1.2000 | H |
| ATOM | 568 | CB | ARG | 36 | 48.080 | -7.537 | 26.835 | -0.0007 | 1.7000 | C |
| ATOM | 569 | HB2 | ARG | 36 | 47.611 | -7.901 | 25.920 | 0.0327 | 1.2000 | H |
| ATOM | 570 | HB3 | ARG | 36 | 49.084 | -7.192 | 26.581 | 0.0327 | 1.2000 | H |
| ATOM | 571 | CG | ARG | 36 | 48.214 | -8.690 | 27.834 | 0.0390 | 1.7000 | C |
| ATOM | 572 | HG2 | ARG | 36 | 48.863 | -8.360 | 28.646 | 0.0285 | 1.2000 | H |
| ATOM | 573 | HG3 | ARG | 36 | 47.250 | -8.932 | 28.276 | 0.0285 | 1.2000 | H |
| ATOM | 574 | CD | ARG | 36 | 48.799 | -9.904 | 27.227 | 0.0486 | 1.7000 | C |
| ATOM | 575 | HD2 | ARG | 36 | 48.163 | -10.216 | 26.397 | 0.0687 | 1.2000 | H |
| ATOM | 576 | HD3 | ARG | 36 | 49.793 | -9.670 | 26.843 | 0.0687 | 1.2000 | H |
| ATOM | 577 | NE | ARG | 36 | 48.899 | -10.991 | 28.194 | -0.5295 | 1.5500 | N |

| | | | | | | | | | | |
|---|---|---|---|---|---|---|---|---|---|---|
| ATOM | 578 | HE | ARG | 36 | 48.743 | -10.757 | 29.163 | 0.3456 | 1.3000 | H |
| ATOM | 579 | CZ | ARG | 36 | 49.168 | -12.272 | 27.882 | 0.8076 | 1.7000 | C |
| ATOM | 580 | NH1 | ARG | 36 | 49.356 | -12.618 | 26.629 | -0.8627 | 1.5500 | N |
| ATOM | 581 | HH11 | ARG | 36 | 49.325 | -11.914 | 25.909 | 0.4478 | 1.3000 | H |
| ATOM | 582 | HH12 | ARG | 36 | 49.554 | -13.578 | 26.393 | 0.4478 | 1.3000 | H |
| ATOM | 583 | NH2 | ARG | 36 | 49.240 | -13.183 | 28.836 | -0.8627 | 1.5500 | N |
| ATOM | 584 | HH21 | ARG | 36 | 49.157 | -12.907 | 29.802 | 0.4478 | 1.3000 | H |
| ATOM | 585 | HH22 | ARG | 36 | 49.368 | -14.154 | 28.601 | 0.4478 | 1.3000 | H |
| ATOM | 586 | C | ARG | 36 | 47.473 | -5.115 | 26.492 | 0.7341 | 1.7000 | C |
| ATOM | 587 | O | ARG | 36 | 48.378 | -4.315 | 26.737 | -0.5894 | 1.5000 | O |
| ATOM | 588 | N | VAL | 37 | 46.626 | -4.955 | 25.466 | -0.4157 | 1.5500 | N |
| ATOM | 589 | H | VAL | 37 | 45.910 | -5.650 | 25.309 | 0.2719 | 1.3000 | H |
| ATOM | 590 | CA | VAL | 37 | 46.676 | -3.804 | 24.559 | -0.0875 | 1.7000 | C |
| ATOM | 591 | HA | VAL | 37 | 47.718 | -3.639 | 24.277 | 0.0969 | 1.2000 | H |
| ATOM | 592 | CB | VAL | 37 | 45.877 | -4.097 | 23.275 | 0.2985 | 1.7000 | C |
| ATOM | 593 | HB | VAL | 37 | 44.877 | -4.451 | 23.523 | -0.0297 | 1.2000 | H |
| ATOM | 594 | CG1 | VAL | 37 | 45.758 | -2.844 | 22.425 | -0.3192 | 1.7000 | C |
| ATOM | 595 | HG11 | VAL | 37 | 45.366 | -3.103 | 21.441 | 0.0791 | 1.2000 | H |
| ATOM | 596 | HG12 | VAL | 37 | 45.065 | -2.133 | 22.873 | 0.0791 | 1.2000 | H |
| ATOM | 597 | HG13 | VAL | 37 | 46.734 | -2.374 | 22.294 | 0.0791 | 1.2000 | H |
| ATOM | 598 | CG2 | VAL | 37 | 46.601 | -5.165 | 22.480 | -0.3192 | 1.7000 | C |
| ATOM | 599 | HG21 | VAL | 37 | 45.968 | -5.505 | 21.656 | 0.0791 | 1.2000 | H |
| ATOM | 600 | HG22 | VAL | 37 | 47.528 | -4.762 | 22.071 | 0.0791 | 1.2000 | H |
| ATOM | 601 | HG23 | VAL | 37 | 46.854 | -6.023 | 23.095 | 0.0791 | 1.2000 | H |
| ATOM | 602 | C | VAL | 37 | 46.168 | -2.499 | 25.177 | 0.5973 | 1.7000 | C |
| ATOM | 603 | O | VAL | 37 | 46.753 | -1.433 | 24.967 | -0.5679 | 1.5000 | O |
| ATOM | 604 | N | VAL | 38 | 45.056 | -2.568 | 25.895 | -0.4157 | 1.5500 | N |
| ATOM | 605 | H | VAL | 38 | 44.611 | -3.461 | 26.057 | 0.2719 | 1.3000 | H |
| ATOM | 606 | CA | VAL | 38 | 44.487 | -1.370 | 26.505 | -0.0875 | 1.7000 | C |
| ATOM | 607 | HA | VAL | 38 | 44.381 | -0.618 | 25.726 | 0.0969 | 1.2000 | H |
| ATOM | 608 | CB | VAL | 38 | 43.090 | -1.693 | 27.075 | 0.2985 | 1.7000 | C |
| ATOM | 609 | HB | VAL | 38 | 42.657 | -0.801 | 27.522 | -0.0297 | 1.2000 | H |
| ATOM | 610 | CG1 | VAL | 38 | 42.166 | -2.157 | 25.969 | -0.3192 | 1.7000 | C |
| ATOM | 611 | HG11 | VAL | 38 | 41.191 | -2.411 | 26.381 | 0.0791 | 1.2000 | H |

| | | | | | | | | | | |
|---|---|---|---|---|---|---|---|---|---|---|
| ATOM | 612 | HG12 | VAL | 38 | 42.032 | -1.351 | 25.249 | 0.0791 | 1.2000 | H |
| ATOM | 613 | HG13 | VAL | 38 | 42.551 | -3.027 | 25.446 | 0.0791 | 1.2000 | H |
| ATOM | 614 | CG2 | VAL | 38 | 43.241 | -2.720 | 28.094 | -0.3192 | 1.7000 | C |
| ATOM | 615 | HG21 | VAL | 38 | 42.263 | -2.952 | 28.516 | 0.0791 | 1.2000 | H |
| ATOM | 616 | HG22 | VAL | 38 | 43.653 | -3.633 | 27.683 | 0.0791 | 1.2000 | H |
| ATOM | 617 | HG23 | VAL | 38 | 43.871 | -2.376 | 28.909 | 0.0791 | 1.2000 | H |
| ATOM | 618 | C | VAL | 38 | 45.446 | -0.864 | 27.610 | 0.5973 | 1.7000 | C |
| ATOM | 619 | O | VAL | 38 | 46.231 | -1.650 | 28.134 | -0.5679 | 1.5000 | O |
| ATOM | 620 | N | PRO | 39 | 45.413 | 0.427 | 27.999 | -0.2548 | 1.5500 | N |
| ATOM | 621 | CD | PRO | 39 | 44.509 | 1.405 | 27.336 | 0.0192 | 1.7000 | C |
| ATOM | 622 | HD2 | PRO | 39 | 43.482 | 1.265 | 27.670 | 0.0391 | 1.2000 | H |
| ATOM | 623 | HD3 | PRO | 39 | 44.586 | 1.377 | 26.249 | 0.0391 | 1.2000 | H |
| ATOM | 624 | CG | PRO | 39 | 45.045 | 2.723 | 27.849 | 0.0189 | 1.7000 | C |
| ATOM | 625 | HG2 | PRO | 39 | 44.255 | 3.471 | 27.924 | 0.0213 | 1.2000 | H |
| ATOM | 626 | HG3 | PRO | 39 | 45.833 | 3.075 | 27.181 | 0.0213 | 1.2000 | H |
| ATOM | 627 | CB | PRO | 39 | 45.643 | 2.409 | 29.205 | -0.0070 | 1.7000 | C |
| ATOM | 628 | HB2 | PRO | 39 | 44.844 | 2.383 | 29.949 | 0.0253 | 1.2000 | H |
| ATOM | 629 | HB3 | PRO | 39 | 46.395 | 3.145 | 29.490 | 0.0253 | 1.2000 | H |
| ATOM | 630 | CA | PRO | 39 | 46.261 | 1.019 | 29.032 | -0.0266 | 1.7000 | C |
| ATOM | 631 | HA | PRO | 39 | 47.288 | 1.092 | 28.673 | 0.0641 | 1.2000 | H |
| ATOM | 632 | C | PRO | 39 | 46.174 | 0.212 | 30.311 | 0.5896 | 1.7000 | C |
| ATOM | 633 | O | PRO | 39 | 45.098 | -0.234 | 30.676 | -0.5748 | 1.5000 | O |
| ATOM | 634 | N | LEU | 40 | 47.274 | 0.094 | 31.047 | -0.4157 | 1.5500 | N |
| ATOM | 635 | H | LEU | 40 | 48.140 | 0.497 | 30.721 | 0.2719 | 1.3000 | H |
| ATOM | 636 | CA | LEU | 40 | 47.307 | -0.739 | 32.259 | -0.0518 | 1.7000 | C |
| ATOM | 637 | HA | LEU | 40 | 47.423 | -1.774 | 31.934 | 0.0922 | 1.2000 | H |
| ATOM | 638 | CB | LEU | 40 | 48.515 | -0.344 | 33.113 | -0.1102 | 1.7000 | C |
| ATOM | 639 | HB2 | LEU | 40 | 49.419 | -0.479 | 32.513 | 0.0457 | 1.2000 | H |
| ATOM | 640 | HB3 | LEU | 40 | 48.434 | 0.720 | 33.343 | 0.0457 | 1.2000 | H |
| ATOM | 641 | CG | LEU | 40 | 48.671 | -1.106 | 34.426 | 0.3531 | 1.7000 | C |
| ATOM | 642 | HG | LEU | 40 | 47.780 | -0.996 | 35.044 | -0.0361 | 1.2000 | H |
| ATOM | 643 | CD1 | LEU | 40 | 48.890 | -2.577 | 34.127 | -0.4121 | 1.7000 | C |
| ATOM | 644 | HD11 | LEU | 40 | 49.095 | -3.110 | 35.056 | 0.1000 | 1.2000 | H |
| ATOM | 645 | HD12 | LEU | 40 | 48.000 | -3.018 | 33.679 | 0.1000 | 1.2000 | H |

| | | | | | | | | | | |
|---|---|---|---|---|---|---|---|---|---|---|
| ATOM | 646 | HD13 | LEU | 40 | 49.738 | -2.708 | 33.454 | 0.1000 | 1.2000 | H |
| ATOM | 647 | CD2 | LEU | 40 | 49.833 | -0.518 | 35.205 | -0.4121 | 1.7000 | C |
| ATOM | 648 | HD21 | LEU | 40 | 49.950 | -1.053 | 36.148 | 0.1000 | 1.2000 | H |
| ATOM | 649 | HD22 | LEU | 40 | 50.757 | -0.603 | 34.632 | 0.1000 | 1.2000 | H |
| ATOM | 650 | HD23 | LEU | 40 | 49.640 | 0.533 | 35.424 | 0.1000 | 1.2000 | H |
| ATOM | 651 | C | LEU | 40 | 46.071 | -0.656 | 33.155 | 0.5973 | 1.7000 | C |
| ATOM | 652 | O | LEU | 40 | 45.542 | -1.691 | 33.573 | -0.5679 | 1.5000 | O |
| ATOM | 653 | N | VAL | 41 | 45.597 | 0.548 | 33.451 | -0.4157 | 1.5500 | N |
| ATOM | 654 | H | VAL | 41 | 46.049 | 1.381 | 33.104 | 0.2719 | 1.3000 | H |
| ATOM | 655 | CA | VAL | 41 | 44.454 | 0.676 | 34.345 | -0.0875 | 1.7000 | C |
| ATOM | 656 | HA | VAL | 41 | 44.636 | 0.084 | 35.245 | 0.0969 | 1.2000 | H |
| ATOM | 657 | CB | VAL | 41 | 44.314 | 2.150 | 34.758 | 0.2985 | 1.7000 | C |
| ATOM | 658 | HB | VAL | 41 | 43.457 | 2.257 | 35.424 | -0.0297 | 1.2000 | H |
| ATOM | 659 | CG1 | VAL | 41 | 45.581 | 2.575 | 35.493 | -0.3192 | 1.7000 | C |
| ATOM | 660 | HG11 | VAL | 41 | 45.298 | 3.025 | 36.446 | 0.0791 | 1.2000 | H |
| ATOM | 661 | HG12 | VAL | 41 | 46.237 | 1.730 | 35.712 | 0.0791 | 1.2000 | H |
| ATOM | 662 | HG13 | VAL | 41 | 46.143 | 3.314 | 34.919 | 0.0791 | 1.2000 | H |
| ATOM | 663 | CG2 | VAL | 41 | 44.109 | 3.033 | 33.511 | -0.3192 | 1.7000 | C |
| ATOM | 664 | HG21 | VAL | 41 | 44.067 | 4.078 | 33.821 | 0.0791 | 1.2000 | H |
| ATOM | 665 | HG22 | VAL | 41 | 44.922 | 2.929 | 32.793 | 0.0791 | 1.2000 | H |
| ATOM | 666 | HG23 | VAL | 41 | 43.160 | 2.802 | 33.027 | 0.0791 | 1.2000 | H |
| ATOM | 667 | C | VAL | 41 | 43.172 | 0.153 | 33.691 | 0.5973 | 1.7000 | C |
| ATOM | 668 | O | VAL | 41 | 42.260 | -0.347 | 34.367 | -0.5679 | 1.5000 | O |
| ATOM | 669 | N | ALA | 42 | 43.132 | 0.231 | 32.363 | -0.4157 | 1.5500 | N |
| ATOM | 670 | H | ALA | 42 | 43.915 | 0.629 | 31.868 | 0.2719 | 1.3000 | H |
| ATOM | 671 | CA | ALA | 42 | 42.028 | -0.259 | 31.580 | 0.0337 | 1.7000 | C |
| ATOM | 672 | HA | ALA | 42 | 41.093 | 0.016 | 32.070 | 0.0823 | 1.2000 | H |
| ATOM | 673 | CB | ALA | 42 | 42.040 | 0.348 | 30.194 | -0.1825 | 1.7000 | C |
| ATOM | 674 | HB1 | ALA | 42 | 41.121 | 0.101 | 29.674 | 0.0603 | 1.2000 | H |
| ATOM | 675 | HB2 | ALA | 42 | 42.109 | 1.433 | 30.269 | 0.0603 | 1.2000 | H |
| ATOM | 676 | HB3 | ALA | 42 | 42.882 | -0.023 | 29.619 | 0.0603 | 1.2000 | H |
| ATOM | 677 | C | ALA | 42 | 42.139 | -1.764 | 31.538 | 0.5973 | 1.7000 | C |
| ATOM | 678 | O | ALA | 42 | 41.135 | -2.458 | 31.557 | -0.5679 | 1.5000 | O |
| ATOM | 679 | N | GLY | 43 | 43.382 | -2.259 | 31.511 | -0.4157 | 1.5500 | N |

| | | | | | | | | | | |
|---|---|---|---|---|---|---|---|---|---|---|
| ATOM | 680 | H | GLY | 43 | 44.165 | -1.622 | 31.510 | 0.2719 | 1.3000 | H |
| ATOM | 681 | CA | GLY | 43 | 43.685 | -3.689 | 31.498 | -0.0252 | 1.7000 | C |
| ATOM | 682 | HA2 | GLY | 43 | 43.239 | -4.160 | 30.622 | 0.0698 | 1.2000 | H |
| ATOM | 683 | HA3 | GLY | 43 | 44.767 | -3.821 | 31.469 | 0.0698 | 1.2000 | H |
| ATOM | 684 | C | GLY | 43 | 43.137 | -4.324 | 32.766 | 0.5973 | 1.7000 | C |
| ATOM | 685 | O | GLY | 43 | 42.559 | -5.413 | 32.741 | -0.5679 | 1.5000 | O |
| ATOM | 686 | N | GLY | 44 | 43.268 | -3.599 | 33.879 | -0.4157 | 1.5500 | N |
| ATOM | 687 | H | GLY | 44 | 43.759 | -2.717 | 33.841 | 0.2719 | 1.3000 | H |
| ATOM | 688 | CA | GLY | 44 | 42.730 | -4.028 | 35.164 | -0.0252 | 1.7000 | C |
| ATOM | 689 | HA2 | GLY | 44 | 43.206 | -4.960 | 35.470 | 0.0698 | 1.2000 | H |
| ATOM | 690 | HA3 | GLY | 44 | 42.932 | -3.258 | 35.910 | 0.0698 | 1.2000 | H |
| ATOM | 691 | C | GLY | 44 | 41.222 | -4.233 | 35.046 | 0.5973 | 1.7000 | C |
| ATOM | 692 | O | GLY | 44 | 40.701 | -5.318 | 35.337 | -0.5679 | 1.5000 | O |
| ATOM | 693 | N | ILE | 45 | 40.520 | -3.189 | 34.589 | -0.4157 | 1.5500 | N |
| ATOM | 694 | H | ILE | 45 | 40.986 | -2.323 | 34.358 | 0.2719 | 1.3000 | H |
| ATOM | 695 | CA | ILE | 45 | 39.068 | -3.275 | 34.451 | -0.0597 | 1.7000 | C |
| ATOM | 696 | HA | ILE | 45 | 38.635 | -3.533 | 35.420 | 0.0869 | 1.2000 | H |
| ATOM | 697 | CB | ILE | 45 | 38.460 | -1.944 | 33.973 | 0.1303 | 1.7000 | C |
| ATOM | 698 | HB | ILE | 45 | 39.003 | -1.624 | 33.085 | 0.0187 | 1.2000 | H |
| ATOM | 699 | CG2 | ILE | 45 | 36.992 | -2.149 | 33.612 | -0.3204 | 1.7000 | C |
| ATOM | 700 | HG21 | ILE | 45 | 36.490 | -1.199 | 33.436 | 0.0882 | 1.2000 | H |
| ATOM | 701 | HG22 | ILE | 45 | 36.896 | -2.724 | 32.693 | 0.0882 | 1.2000 | H |
| ATOM | 702 | HG23 | ILE | 45 | 36.473 | -2.663 | 34.421 | 0.0882 | 1.2000 | H |
| ATOM | 703 | CG1 | ILE | 45 | 38.587 | -0.875 | 35.014 | -0.0430 | 1.7000 | C |
| ATOM | 704 | HG12 | ILE | 45 | 37.937 | -1.099 | 35.862 | 0.0236 | 1.2000 | H |
| ATOM | 705 | HG13 | ILE | 45 | 39.613 | -0.840 | 35.382 | 0.0236 | 1.2000 | H |
| ATOM | 706 | CD1 | ILE | 45 | 38.252 | 0.488 | 34.458 | -0.0660 | 1.7000 | C |
| ATOM | 707 | HD11 | ILE | 45 | 38.061 | 0.465 | 33.384 | 0.0186 | 1.2000 | H |
| ATOM | 708 | HD12 | ILE | 45 | 37.379 | 0.889 | 34.967 | 0.0186 | 1.2000 | H |
| ATOM | 709 | HD13 | ILE | 45 | 39.089 | 1.164 | 34.636 | 0.0186 | 1.2000 | H |
| ATOM | 710 | C | ILE | 45 | 38.684 | -4.355 | 33.449 | 0.5973 | 1.7000 | C |
| ATOM | 711 | O | ILE | 45 | 37.807 | -5.168 | 33.721 | -0.5679 | 1.5000 | O |
| ATOM | 712 | N | CYX | 46 | 39.371 | -4.376 | 32.317 | -0.4157 | 1.5500 | N |
| ATOM | 713 | H | CYX | 46 | 40.116 | -3.709 | 32.210 | 0.2719 | 1.3000 | H |

| | | | | | | | | | | |
|---|---|---|---|---|---|---|---|---|---|---|
| ATOM | 714 | CA | CYX | 46 | 39.147 | -5.294 | 31.216 | 0.0429 | 1.7000 | C |
| ATOM | 715 | HA | CYX | 46 | 38.185 | -5.078 | 30.754 | 0.0766 | 1.2000 | H |
| ATOM | 716 | CB | CYX | 46 | 40.258 | -5.066 | 30.187 | -0.0790 | 1.7000 | C |
| ATOM | 717 | HB2 | CYX | 46 | 40.169 | -4.042 | 29.827 | 0.0910 | 1.2000 | H |
| ATOM | 718 | HB3 | CYX | 46 | 41.227 | -5.159 | 30.670 | 0.0910 | 1.2000 | H |
| ATOM | 719 | SG | CYX | 46 | 40.270 | -6.095 | 28.776 | -0.1081 | 1.8000 | S |
| ATOM | 720 | C | CYX | 46 | 39.168 | -6.735 | 31.667 | 0.5973 | 1.7000 | C |
| ATOM | 721 | O | CYX | 46 | 38.242 | -7.505 | 31.378 | -0.5679 | 1.5000 | O |
| ATOM | 722 | N | GLN | 47 | 40.217 | -7.104 | 32.393 | -0.4157 | 1.5500 | N |
| ATOM | 723 | H | GLN | 47 | 40.947 | -6.441 | 32.614 | 0.2719 | 1.3000 | H |
| ATOM | 724 | CA | GLN | 47 | 40.356 | -8.466 | 32.847 | -0.0031 | 1.7000 | C |
| ATOM | 725 | HA | GLN | 47 | 40.263 | -9.138 | 31.991 | 0.0850 | 1.2000 | H |
| ATOM | 726 | CB | GLN | 47 | 41.753 | -8.630 | 33.449 | -0.0036 | 1.7000 | C |
| ATOM | 727 | HB2 | GLN | 47 | 42.483 | -8.271 | 32.721 | 0.0171 | 1.2000 | H |
| ATOM | 728 | HB3 | GLN | 47 | 41.830 | -7.981 | 34.324 | 0.0171 | 1.2000 | H |
| ATOM | 729 | CG | GLN | 47 | 42.179 | -10.035 | 33.873 | -0.0645 | 1.7000 | C |
| ATOM | 730 | HG2 | GLN | 47 | 43.161 | -9.978 | 34.344 | 0.0352 | 1.2000 | H |
| ATOM | 731 | HG3 | GLN | 47 | 41.476 | -10.392 | 34.627 | 0.0352 | 1.2000 | H |
| ATOM | 732 | CD | GLN | 47 | 42.260 | -11.039 | 32.723 | 0.6951 | 1.7000 | C |
| ATOM | 733 | OE1 | GLN | 47 | 42.805 | -10.758 | 31.648 | -0.6086 | 1.5000 | O |
| ATOM | 734 | NE2 | GLN | 47 | 41.768 | -12.246 | 32.980 | -0.9407 | 1.5500 | N |
| ATOM | 735 | HE21 | GLN | 47 | 41.396 | -12.448 | 33.901 | 0.4251 | 1.3000 | H |
| ATOM | 736 | HE22 | GLN | 47 | 41.790 | -12.961 | 32.272 | 0.4251 | 1.3000 | H |
| ATOM | 737 | C | GLN | 47 | 39.259 | -8.784 | 33.846 | 0.5973 | 1.7000 | C |
| ATOM | 738 | O | GLN | 47 | 38.655 | -9.853 | 33.784 | -0.5679 | 1.5000 | O |
| ATOM | 739 | N | CYS | 48 | 38.915 | -7.825 | 34.710 | -0.4157 | 1.5500 | N |
| ATOM | 740 | H | CYS | 48 | 39.413 | -6.945 | 34.727 | 0.2719 | 1.3000 | H |
| ATOM | 741 | CA | CYS | 48 | 37.867 | -8.071 | 35.692 | 0.0213 | 1.7000 | C |
| ATOM | 742 | HA | CYS | 48 | 38.106 | -8.975 | 36.257 | 0.1124 | 1.2000 | H |
| ATOM | 743 | CB | CYS | 48 | 37.778 | -6.895 | 36.659 | -0.1231 | 1.7000 | C |
| ATOM | 744 | HB2 | CYS | 48 | 37.636 | -5.968 | 36.105 | 0.1112 | 1.2000 | H |
| ATOM | 745 | HB3 | CYS | 48 | 36.909 | -7.043 | 37.301 | 0.1112 | 1.2000 | H |
| ATOM | 746 | SG | CYS | 48 | 39.226 | -6.723 | 37.724 | -0.3119 | 1.8000 | S |
| ATOM | 747 | HG | CYS | 48 | 40.095 | -6.473 | 36.752 | 0.1933 | 1.2000 | H |

| | | | | | | | | | | |
|---|---|---|---|---|---|---|---|---|---|---|
| ATOM | 748 | C | CYS | 48 | 36.511 | -8.265 | 35.007 | 0.5973 | 1.7000 | C |
| ATOM | 749 | O | CYS | 48 | 35.714 | -9.133 | 35.393 | -0.5679 | 1.5000 | O |
| ATOM | 750 | N | LEU | 49 | 36.259 | -7.483 | 33.951 | -0.4157 | 1.5500 | N |
| ATOM | 751 | H | LEU | 49 | 36.949 | -6.815 | 33.638 | 0.2719 | 1.3000 | H |
| ATOM | 752 | CA | LEU | 49 | 35.001 | -7.592 | 33.235 | -0.0518 | 1.7000 | C |
| ATOM | 753 | HA | LEU | 49 | 34.165 | -7.435 | 33.918 | 0.0922 | 1.2000 | H |
| ATOM | 754 | CB | LEU | 49 | 34.950 | -6.568 | 32.080 | -0.1102 | 1.7000 | C |
| ATOM | 755 | HB2 | LEU | 49 | 35.838 | -6.711 | 31.466 | 0.0457 | 1.2000 | H |
| ATOM | 756 | HB3 | LEU | 49 | 34.104 | -6.821 | 31.446 | 0.0457 | 1.2000 | H |
| ATOM | 757 | CG | LEU | 49 | 34.856 | -5.061 | 32.417 | 0.3531 | 1.7000 | C |
| ATOM | 758 | HG | LEU | 49 | 35.627 | -4.773 | 33.111 | -0.0361 | 1.2000 | H |
| ATOM | 759 | CD1 | LEU | 49 | 35.024 | -4.261 | 31.123 | -0.4121 | 1.7000 | C |
| ATOM | 760 | HD11 | LEU | 49 | 34.950 | -3.196 | 31.341 | 0.1000 | 1.2000 | H |
| ATOM | 761 | HD12 | LEU | 49 | 36.001 | -4.461 | 30.686 | 0.1000 | 1.2000 | H |
| ATOM | 762 | HD13 | LEU | 49 | 34.251 | -4.528 | 30.402 | 0.1000 | 1.2000 | H |
| ATOM | 763 | CD2 | LEU | 49 | 33.573 | -4.733 | 33.054 | -0.4121 | 1.7000 | C |
| ATOM | 764 | HD21 | LEU | 49 | 33.546 | -3.666 | 33.254 | 0.1000 | 1.2000 | H |
| ATOM | 765 | HD22 | LEU | 49 | 32.755 | -4.966 | 32.384 | 0.1000 | 1.2000 | H |
| ATOM | 766 | HD23 | LEU | 49 | 33.453 | -5.274 | 33.992 | 0.1000 | 1.2000 | H |
| ATOM | 767 | C | LEU | 49 | 34.897 | -8.978 | 32.625 | 0.5973 | 1.7000 | C |
| ATOM | 768 | O | LEU | 49 | 33.852 | -9.627 | 32.699 | -0.5679 | 1.5000 | O |
| ATOM | 769 | N | ALA | 50 | 35.998 | -9.442 | 32.039 | -0.4157 | 1.5500 | N |
| ATOM | 770 | H | ALA | 50 | 36.840 | -8.882 | 32.031 | 0.2719 | 1.3000 | H |
| ATOM | 771 | CA | ALA | 50 | 36.030 | -10.744 | 31.408 | 0.0337 | 1.7000 | C |
| ATOM | 772 | HA | ALA | 50 | 35.209 | -10.805 | 30.692 | 0.0823 | 1.2000 | H |
| ATOM | 773 | CB | ALA | 50 | 37.315 | -10.886 | 30.656 | -0.1825 | 1.7000 | C |
| ATOM | 774 | HB1 | ALA | 50 | 37.323 | -11.849 | 30.147 | 0.0603 | 1.2000 | H |
| ATOM | 775 | HB2 | ALA | 50 | 37.402 | -10.093 | 29.914 | 0.0603 | 1.2000 | H |
| ATOM | 776 | HB3 | ALA | 50 | 38.164 | -10.833 | 31.339 | 0.0603 | 1.2000 | H |
| ATOM | 777 | C | ALA | 50 | 35.891 | -11.897 | 32.397 | 0.5973 | 1.7000 | C |
| ATOM | 778 | O | ALA | 50 | 35.126 | -12.841 | 32.167 | -0.5679 | 1.5000 | O |
| ATOM | 779 | N | GLU | 51 | 36.549 | -11.805 | 33.544 | -0.5163 | 1.5500 | N |
| ATOM | 780 | H | GLU | 51 | 37.161 | -11.026 | 33.740 | 0.2936 | 1.3000 | H |
| ATOM | 781 | CA | GLU | 51 | 36.457 | -12.900 | 34.496 | 0.0397 | 1.7000 | C |

| | | | | | | | | | | |
|---|---|---|---|---|---|---|---|---|---|---|
| ATOM | 782 | HA | GLU | 51 | 36.757 | -13.819 | 33.990 | 0.1105 | 1.2000 | H |
| ATOM | 783 | CB | GLU | 51 | 37.420 | -12.687 | 35.660 | 0.0560 | 1.7000 | C |
| ATOM | 784 | HB2 | GLU | 51 | 37.276 | -11.686 | 36.070 | -0.0173 | 1.2000 | H |
| ATOM | 785 | HB3 | GLU | 51 | 37.203 | -13.418 | 36.440 | -0.0173 | 1.2000 | H |
| ATOM | 786 | CG | GLU | 51 | 38.874 | -12.861 | 35.227 | 0.0136 | 1.7000 | C |
| ATOM | 787 | HG2 | GLU | 51 | 38.998 | -13.888 | 34.879 | -0.0425 | 1.2000 | H |
| ATOM | 788 | HG3 | GLU | 51 | 39.086 | -12.215 | 34.379 | -0.0425 | 1.2000 | H |
| ATOM | 789 | CD | GLU | 51 | 39.897 | -12.609 | 36.278 | 0.8054 | 1.7000 | C |
| ATOM | 790 | OE1 | GLU | 51 | 39.560 | -12.315 | 37.405 | -0.8188 | 1.5000 | O |
| ATOM | 791 | OE2 | GLU | 51 | 41.052 | -12.688 | 35.922 | -0.8188 | 1.5000 | O |
| ATOM | 792 | C | GLU | 51 | 35.021 | -13.099 | 34.958 | 0.5366 | 1.7000 | C |
| ATOM | 793 | O | GLU | 51 | 34.588 | -14.230 | 35.188 | -0.5819 | 1.5000 | O |
| ATOM | 794 | N | ARG | 52 | 34.270 | -12.008 | 35.074 | -0.3479 | 1.5500 | N |
| ATOM | 795 | H | ARG | 52 | 34.666 | -11.096 | 34.891 | 0.2747 | 1.3000 | H |
| ATOM | 796 | CA | ARG | 52 | 32.880 | -12.112 | 35.476 | -0.2637 | 1.7000 | C |
| ATOM | 797 | HA | ARG | 52 | 32.778 | -12.896 | 36.229 | 0.1560 | 1.2000 | H |
| ATOM | 798 | CB | ARG | 52 | 32.430 | -10.807 | 36.103 | -0.0007 | 1.7000 | C |
| ATOM | 799 | HB2 | ARG | 52 | 32.655 | -9.993 | 35.412 | 0.0327 | 1.2000 | H |
| ATOM | 800 | HB3 | ARG | 52 | 31.346 | -10.854 | 36.221 | 0.0327 | 1.2000 | H |
| ATOM | 801 | CG | ARG | 52 | 33.034 | -10.483 | 37.451 | 0.0390 | 1.7000 | C |
| ATOM | 802 | HG2 | ARG | 52 | 32.766 | -11.262 | 38.166 | 0.0285 | 1.2000 | H |
| ATOM | 803 | HG3 | ARG | 52 | 34.121 | -10.460 | 37.362 | 0.0285 | 1.2000 | H |
| ATOM | 804 | CD | ARG | 52 | 32.577 | -9.147 | 37.961 | 0.0486 | 1.7000 | C |
| ATOM | 805 | HD2 | ARG | 52 | 33.112 | -8.915 | 38.883 | 0.0687 | 1.2000 | H |
| ATOM | 806 | HD3 | ARG | 52 | 32.831 | -8.380 | 37.226 | 0.0687 | 1.2000 | H |
| ATOM | 807 | NE | ARG | 52 | 31.130 | -9.113 | 38.204 | -0.5295 | 1.5500 | N |
| ATOM | 808 | HE | ARG | 52 | 30.558 | -8.744 | 37.459 | 0.3456 | 1.3000 | H |
| ATOM | 809 | CZ | ARG | 52 | 30.513 | -9.545 | 39.302 | 0.8076 | 1.7000 | C |
| ATOM | 810 | NH1 | ARG | 52 | 31.180 | -10.057 | 40.317 | -0.8627 | 1.5500 | N |
| ATOM | 811 | HH11 | ARG | 52 | 32.183 | -10.151 | 40.260 | 0.4478 | 1.3000 | H |
| ATOM | 812 | HH12 | ARG | 52 | 30.692 | -10.359 | 41.145 | 0.4478 | 1.3000 | H |
| ATOM | 813 | NH2 | ARG | 52 | 29.210 | -9.440 | 39.320 | -0.8627 | 1.5500 | N |
| ATOM | 814 | HH21 | ARG | 52 | 28.737 | -8.975 | 38.562 | 0.4478 | 1.3000 | H |
| ATOM | 815 | HH22 | ARG | 52 | 28.684 | -9.824 | 40.089 | 0.4478 | 1.3000 | H |

| | | | | | | | | | | |
|---|---|---|---|---|---|---|---|---|---|---|
| ATOM | 816 | C | ARG | 52 | 31.908 | -12.448 | 34.326 | 0.7341 | 1.7000 | C |
| ATOM | 817 | O | ARG | 52 | 30.919 | -13.154 | 34.547 | -0.5894 | 1.5000 | O |
| ATOM | 818 | N | TYR | 53 | 32.165 | -11.934 | 33.109 | -0.4157 | 1.5500 | N |
| ATOM | 819 | H | TYR | 53 | 32.985 | -11.363 | 32.967 | 0.2719 | 1.3000 | H |
| ATOM | 820 | CA | TYR | 53 | 31.188 | -12.065 | 32.023 | -0.0014 | 1.7000 | C |
| ATOM | 821 | HA | TYR | 53 | 30.317 | -12.613 | 32.379 | 0.0876 | 1.2000 | H |
| ATOM | 822 | CB | TYR | 53 | 30.717 | -10.664 | 31.717 | -0.0152 | 1.7000 | C |
| ATOM | 823 | HB2 | TYR | 53 | 31.545 | -10.076 | 31.318 | 0.0295 | 1.2000 | H |
| ATOM | 824 | HB3 | TYR | 53 | 29.938 | -10.702 | 30.962 | 0.0295 | 1.2000 | H |
| ATOM | 825 | CG | TYR | 53 | 30.154 | -10.014 | 32.977 | -0.0011 | 1.7000 | C |
| ATOM | 826 | CD1 | TYR | 53 | 30.776 | -8.913 | 33.510 | -0.1906 | 1.7000 | C |
| ATOM | 827 | HD1 | TYR | 53 | 31.648 | -8.507 | 33.051 | 0.1699 | 1.2000 | H |
| ATOM | 828 | CE1 | TYR | 53 | 30.308 | -8.329 | 34.645 | -0.2341 | 1.7000 | C |
| ATOM | 829 | HE1 | TYR | 53 | 30.807 | -7.463 | 35.054 | 0.1656 | 1.2000 | H |
| ATOM | 830 | CZ | TYR | 53 | 29.196 | -8.850 | 35.276 | 0.3226 | 1.7000 | C |
| ATOM | 831 | OH | TYR | 53 | 28.764 | -8.256 | 36.433 | -0.5579 | 1.5000 | O |
| ATOM | 832 | HH | TYR | 53 | 27.836 | -8.440 | 36.603 | 0.3992 | 1.2000 | H |
| ATOM | 833 | CE2 | TYR | 53 | 28.557 | -9.958 | 34.761 | -0.2341 | 1.7000 | C |
| ATOM | 834 | HE2 | TYR | 53 | 27.696 | -10.375 | 35.262 | 0.1656 | 1.2000 | H |
| ATOM | 835 | CD2 | TYR | 53 | 29.034 | -10.546 | 33.614 | -0.1906 | 1.7000 | C |
| ATOM | 836 | HD2 | TYR | 53 | 28.535 | -11.416 | 33.211 | 0.1699 | 1.2000 | H |
| ATOM | 837 | C | TYR | 53 | 31.602 | -12.810 | 30.732 | 0.5973 | 1.7000 | C |
| ATOM | 838 | O | TYR | 53 | 30.773 | -12.990 | 29.824 | -0.5679 | 1.5000 | O |
| ATOM | 839 | N | SER | 54 | 32.838 | -13.315 | 30.659 | -0.4157 | 1.5500 | N |
| ATOM | 840 | H | SER | 54 | 33.467 | -13.196 | 31.438 | 0.2719 | 1.3000 | H |
| ATOM | 841 | CA | SER | 54 | 33.317 | -14.068 | 29.491 | -0.0249 | 1.7000 | C |
| ATOM | 842 | HA | SER | 54 | 33.208 | -13.439 | 28.608 | 0.0843 | 1.2000 | H |
| ATOM | 843 | CB | SER | 54 | 34.797 | -14.376 | 29.669 | 0.2117 | 1.7000 | C |
| ATOM | 844 | HB2 | SER | 54 | 35.156 | -14.925 | 28.802 | 0.0352 | 1.2000 | H |
| ATOM | 845 | HB3 | SER | 54 | 35.366 | -13.449 | 29.741 | 0.0352 | 1.2000 | H |
| ATOM | 846 | OG | SER | 54 | 35.019 | -15.167 | 30.799 | -0.6546 | 1.5000 | O |
| ATOM | 847 | HG | SER | 54 | 35.021 | -14.593 | 31.578 | 0.4275 | 1.2000 | H |
| ATOM | 848 | C | SER | 54 | 32.499 | -15.351 | 29.272 | 0.5973 | 1.7000 | C |
| ATOM | 849 | O | SER | 54 | 32.415 | -15.886 | 28.158 | -0.5679 | 1.5000 | O |

| | | | | | | | | | | |
|---|---|---|---|---|---|---|---|---|---|---|
| ATOM | 850 | N | VAL | 55 | 31.841 | -15.785 | 30.347 | -0.4157 | 1.5500 | N |
| ATOM | 851 | H | VAL | 55 | 31.974 | -15.285 | 31.214 | 0.2719 | 1.3000 | H |
| ATOM | 852 | CA | VAL | 55 | 30.975 | -16.949 | 30.409 | -0.0875 | 1.7000 | C |
| ATOM | 853 | HA | VAL | 55 | 31.508 | -17.803 | 29.988 | 0.0969 | 1.2000 | H |
| ATOM | 854 | CB | VAL | 55 | 30.659 | -17.243 | 31.884 | 0.2985 | 1.7000 | C |
| ATOM | 855 | HB | VAL | 55 | 30.024 | -18.128 | 31.951 | -0.0297 | 1.2000 | H |
| ATOM | 856 | CG1 | VAL | 55 | 31.955 | -17.510 | 32.620 | -0.3192 | 1.7000 | C |
| ATOM | 857 | HG11 | VAL | 55 | 31.732 | -17.840 | 33.635 | 0.0791 | 1.2000 | H |
| ATOM | 858 | HG12 | VAL | 55 | 32.512 | -18.304 | 32.121 | 0.0791 | 1.2000 | H |
| ATOM | 859 | HG13 | VAL | 55 | 32.579 | -16.618 | 32.686 | 0.0791 | 1.2000 | H |
| ATOM | 860 | CG2 | VAL | 55 | 29.944 | -16.059 | 32.509 | -0.3192 | 1.7000 | C |
| ATOM | 861 | HG21 | VAL | 55 | 29.787 | -16.258 | 33.569 | 0.0791 | 1.2000 | H |
| ATOM | 862 | HG22 | VAL | 55 | 30.516 | -15.136 | 32.422 | 0.0791 | 1.2000 | H |
| ATOM | 863 | HG23 | VAL | 55 | 28.961 | -15.934 | 32.059 | 0.0791 | 1.2000 | H |
| ATOM | 864 | C | VAL | 55 | 29.684 | -16.758 | 29.599 | 0.5973 | 1.7000 | C |
| ATOM | 865 | O | VAL | 55 | 29.014 | -17.731 | 29.258 | -0.5679 | 1.5000 | O |
| ATOM | 866 | N | ILE | 56 | 29.331 | -15.501 | 29.313 | -0.4157 | 1.5500 | N |
| ATOM | 867 | H | ILE | 56 | 29.908 | -14.744 | 29.647 | 0.2719 | 1.3000 | H |
| ATOM | 868 | CA | ILE | 56 | 28.174 | -15.168 | 28.489 | -0.0597 | 1.7000 | C |
| ATOM | 869 | HA | ILE | 56 | 27.488 | -16.014 | 28.434 | 0.0869 | 1.2000 | H |
| ATOM | 870 | CB | ILE | 56 | 27.418 | -13.932 | 29.019 | 0.1303 | 1.7000 | C |
| ATOM | 871 | HB | ILE | 56 | 28.099 | -13.080 | 29.031 | 0.0187 | 1.2000 | H |
| ATOM | 872 | CG2 | ILE | 56 | 26.229 | -13.567 | 28.100 | -0.3204 | 1.7000 | C |
| ATOM | 873 | HG21 | ILE | 56 | 25.616 | -12.800 | 28.573 | 0.0882 | 1.2000 | H |
| ATOM | 874 | HG22 | ILE | 56 | 26.591 | -13.169 | 27.152 | 0.0882 | 1.2000 | H |
| ATOM | 875 | HG23 | ILE | 56 | 25.612 | -14.443 | 27.899 | 0.0882 | 1.2000 | H |
| ATOM | 876 | CG1 | ILE | 56 | 26.975 | -14.186 | 30.460 | -0.0430 | 1.7000 | C |
| ATOM | 877 | HG12 | ILE | 56 | 27.843 | -14.237 | 31.114 | 0.0236 | 1.2000 | H |
| ATOM | 878 | HG13 | ILE | 56 | 26.374 | -13.344 | 30.798 | 0.0236 | 1.2000 | H |
| ATOM | 879 | CD1 | ILE | 56 | 26.140 | -15.421 | 30.628 | -0.0660 | 1.7000 | C |
| ATOM | 880 | HD11 | ILE | 56 | 25.765 | -15.455 | 31.650 | 0.0186 | 1.2000 | H |
| ATOM | 881 | HD12 | ILE | 56 | 25.284 | -15.416 | 29.955 | 0.0186 | 1.2000 | H |
| ATOM | 882 | HD13 | ILE | 56 | 26.731 | -16.322 | 30.468 | 0.0186 | 1.2000 | H |
| ATOM | 883 | C | ILE | 56 | 28.649 | -14.887 | 27.071 | 0.5973 | 1.7000 | C |

| | | | | | | | | | | |
|---|---|---|---|---|---|---|---|---|---|---|
| ATOM | 884 | O | ILE | 56 | 28.029 | -15.311 | 26.076 | -0.5679 | 1.5000 | O |
| ATOM | 885 | N | LEU | 57 | 29.761 | -14.153 | 27.000 | -0.4157 | 1.5500 | N |
| ATOM | 886 | H | LEU | 57 | 30.219 | -13.856 | 27.849 | 0.2719 | 1.3000 | H |
| ATOM | 887 | CA | LEU | 57 | 30.304 | -13.719 | 25.732 | -0.0518 | 1.7000 | C |
| ATOM | 888 | HA | LEU | 57 | 29.536 | -13.094 | 25.274 | 0.0922 | 1.2000 | H |
| ATOM | 889 | CB | LEU | 57 | 31.560 | -12.878 | 25.936 | -0.1102 | 1.7000 | C |
| ATOM | 890 | HB2 | LEU | 57 | 31.297 | -11.976 | 26.493 | 0.0457 | 1.2000 | H |
| ATOM | 891 | HB3 | LEU | 57 | 32.264 | -13.449 | 26.541 | 0.0457 | 1.2000 | H |
| ATOM | 892 | CG | LEU | 57 | 32.255 | -12.501 | 24.635 | 0.3531 | 1.7000 | C |
| ATOM | 893 | HG | LEU | 57 | 32.597 | -13.380 | 24.086 | -0.0361 | 1.2000 | H |
| ATOM | 894 | CD1 | LEU | 57 | 31.301 | -11.749 | 23.814 | -0.4121 | 1.7000 | C |
| ATOM | 895 | HD11 | LEU | 57 | 31.813 | -11.389 | 22.925 | 0.1000 | 1.2000 | H |
| ATOM | 896 | HD12 | LEU | 57 | 30.466 | -12.345 | 23.466 | 0.1000 | 1.2000 | H |
| ATOM | 897 | HD13 | LEU | 57 | 30.917 | -10.890 | 24.359 | 0.1000 | 1.2000 | H |
| ATOM | 898 | CD2 | LEU | 57 | 33.494 | -11.663 | 24.913 | -0.4121 | 1.7000 | C |
| ATOM | 899 | HD21 | LEU | 57 | 33.964 | -11.372 | 23.973 | 0.1000 | 1.2000 | H |
| ATOM | 900 | HD22 | LEU | 57 | 33.231 | -10.768 | 25.475 | 0.1000 | 1.2000 | H |
| ATOM | 901 | HD23 | LEU | 57 | 34.208 | -12.244 | 25.494 | 0.1000 | 1.2000 | H |
| ATOM | 902 | C | LEU | 57 | 30.660 | -14.808 | 24.748 | 0.5973 | 1.7000 | C |
| ATOM | 903 | O | LEU | 57 | 30.210 | -14.731 | 23.611 | -0.5679 | 1.5000 | O |
| ATOM | 904 | N | LEU | 58 | 31.424 | -15.832 | 25.135 | -0.4157 | 1.5500 | N |
| ATOM | 905 | H | LEU | 58 | 31.766 | -15.930 | 26.081 | 0.2719 | 1.3000 | H |
| ATOM | 906 | CA | LEU | 58 | 31.831 | -16.772 | 24.079 | -0.0518 | 1.7000 | C |
| ATOM | 907 | HA | LEU | 58 | 32.345 | -16.178 | 23.326 | 0.0922 | 1.2000 | H |
| ATOM | 908 | CB | LEU | 58 | 32.795 | -17.853 | 24.574 | -0.1102 | 1.7000 | C |
| ATOM | 909 | HB2 | LEU | 58 | 33.650 | -17.387 | 25.053 | 0.0457 | 1.2000 | H |
| ATOM | 910 | HB3 | LEU | 58 | 32.282 | -18.448 | 25.332 | 0.0457 | 1.2000 | H |
| ATOM | 911 | CG | LEU | 58 | 33.289 | -18.796 | 23.453 | 0.3531 | 1.7000 | C |
| ATOM | 912 | HG | LEU | 58 | 32.443 | -19.331 | 23.026 | -0.0361 | 1.2000 | H |
| ATOM | 913 | CD1 | LEU | 58 | 33.996 | -17.963 | 22.363 | -0.4121 | 1.7000 | C |
| ATOM | 914 | HD11 | LEU | 58 | 34.491 | -18.629 | 21.656 | 0.1000 | 1.2000 | H |
| ATOM | 915 | HD12 | LEU | 58 | 33.268 | -17.376 | 21.815 | 0.1000 | 1.2000 | H |
| ATOM | 916 | HD13 | LEU | 58 | 34.730 | -17.288 | 22.778 | 0.1000 | 1.2000 | H |
| ATOM | 917 | CD2 | LEU | 58 | 34.250 | -19.820 | 24.019 | -0.4121 | 1.7000 | C |

| | | | | | | | | | | |
|---|---|---|---|---|---|---|---|---|---|---|
| ATOM | 918 | HD21 | LEU | 58 | 35.269 | -19.436 | 24.010 | 0.1000 | 1.2000 | H |
| ATOM | 919 | HD22 | LEU | 58 | 33.973 | -20.080 | 25.043 | 0.1000 | 1.2000 | H |
| ATOM | 920 | HD23 | LEU | 58 | 34.199 | -20.724 | 23.408 | 0.1000 | 1.2000 | H |
| ATOM | 921 | C | LEU | 58 | 30.631 | -17.456 | 23.417 | 0.5973 | 1.7000 | C |
| ATOM | 922 | O | LEU | 58 | 30.584 | -17.601 | 22.194 | -0.5679 | 1.5000 | O |
| ATOM | 923 | N | ASP | 59 | 29.654 | -17.888 | 24.209 | -0.5163 | 1.5500 | N |
| ATOM | 924 | H | ASP | 59 | 29.710 | -17.793 | 25.212 | 0.2936 | 1.3000 | H |
| ATOM | 925 | CA | ASP | 59 | 28.504 | -18.555 | 23.618 | 0.0381 | 1.7000 | C |
| ATOM | 926 | HA | ASP | 59 | 28.856 | -19.398 | 23.021 | 0.0880 | 1.2000 | H |
| ATOM | 927 | CB | ASP | 59 | 27.566 | -19.091 | 24.699 | -0.0303 | 1.7000 | C |
| ATOM | 928 | HB2 | ASP | 59 | 27.347 | -18.298 | 25.419 | -0.0122 | 1.2000 | H |
| ATOM | 929 | HB3 | ASP | 59 | 26.626 | -19.391 | 24.233 | -0.0122 | 1.2000 | H |
| ATOM | 930 | CG | ASP | 59 | 28.127 | -20.304 | 25.429 | 0.7994 | 1.7000 | C |
| ATOM | 931 | OD1 | ASP | 59 | 29.100 | -20.859 | 24.968 | -0.8014 | 1.5000 | O |
| ATOM | 932 | OD2 | ASP | 59 | 27.558 | -20.690 | 26.420 | -0.8014 | 1.5000 | O |
| ATOM | 933 | C | ASP | 59 | 27.719 | -17.639 | 22.680 | 0.5366 | 1.7000 | C |
| ATOM | 934 | O | ASP | 59 | 27.238 | -18.075 | 21.632 | -0.5819 | 1.5000 | O |
| ATOM | 935 | N | THR | 60 | 27.604 | -16.355 | 23.031 | -0.4157 | 1.5500 | N |
| ATOM | 936 | H | THR | 60 | 28.021 | -16.020 | 23.888 | 0.2719 | 1.3000 | H |
| ATOM | 937 | CA | THR | 60 | 26.853 | -15.438 | 22.184 | -0.0389 | 1.7000 | C |
| ATOM | 938 | HA | THR | 60 | 25.914 | -15.913 | 21.894 | 0.1007 | 1.2000 | H |
| ATOM | 939 | CB | THR | 60 | 26.524 | -14.141 | 22.963 | 0.3654 | 1.7000 | C |
| ATOM | 940 | HB | THR | 60 | 27.435 | -13.595 | 23.212 | 0.0043 | 1.2000 | H |
| ATOM | 941 | CG2 | THR | 60 | 25.573 | -13.256 | 22.132 | -0.2438 | 1.7000 | C |
| ATOM | 942 | HG21 | THR | 60 | 25.060 | -12.549 | 22.785 | 0.0642 | 1.2000 | H |
| ATOM | 943 | HG22 | THR | 60 | 26.135 | -12.686 | 21.406 | 0.0642 | 1.2000 | H |
| ATOM | 944 | HG23 | THR | 60 | 24.826 | -13.865 | 21.622 | 0.0642 | 1.2000 | H |
| ATOM | 945 | OG1 | THR | 60 | 25.822 | -14.483 | 24.170 | -0.6761 | 1.5000 | O |
| ATOM | 946 | HG1 | THR | 60 | 26.407 | -15.005 | 24.726 | 0.4102 | 1.2000 | H |
| ATOM | 947 | C | THR | 60 | 27.678 | -15.150 | 20.904 | 0.5973 | 1.7000 | C |
| ATOM | 948 | O | THR | 60 | 27.149 | -15.143 | 19.776 | -0.5679 | 1.5000 | O |
| ATOM | 949 | N | LEU | 61 | 28.990 | -14.939 | 21.105 | -0.4157 | 1.5500 | N |
| ATOM | 950 | H | LEU | 61 | 29.317 | -15.025 | 22.055 | 0.2719 | 1.3000 | H |
| ATOM | 951 | CA | LEU | 61 | 30.027 | -14.618 | 20.110 | -0.0518 | 1.7000 | C |

| | | | | | | | | | | |
|---|---|---|---|---|---|---|---|---|---|---|
| ATOM | 952 | HA | LEU | 61 | 29.829 | -13.625 | 19.717 | 0.0922 | 1.2000 | H |
| ATOM | 953 | CB | LEU | 61 | 31.383 | -14.590 | 20.814 | -0.1102 | 1.7000 | C |
| ATOM | 954 | HB2 | LEU | 61 | 31.240 | -13.867 | 21.615 | 0.0457 | 1.2000 | H |
| ATOM | 955 | HB3 | LEU | 61 | 31.584 | -15.553 | 21.273 | 0.0457 | 1.2000 | H |
| ATOM | 956 | CG | LEU | 61 | 32.573 | -14.157 | 20.057 | 0.3531 | 1.7000 | C |
| ATOM | 957 | HG | LEU | 61 | 32.297 | -13.370 | 19.358 | -0.0361 | 1.2000 | H |
| ATOM | 958 | CD1 | LEU | 61 | 33.528 | -13.604 | 21.040 | -0.4121 | 1.7000 | C |
| ATOM | 959 | HD11 | LEU | 61 | 34.401 | -13.290 | 20.484 | 0.1000 | 1.2000 | H |
| ATOM | 960 | HD12 | LEU | 61 | 33.121 | -12.724 | 21.534 | 0.1000 | 1.2000 | H |
| ATOM | 961 | HD13 | LEU | 61 | 33.807 | -14.353 | 21.784 | 0.1000 | 1.2000 | H |
| ATOM | 962 | CD2 | LEU | 61 | 33.201 | -15.324 | 19.301 | -0.4121 | 1.7000 | C |
| ATOM | 963 | HD21 | LEU | 61 | 33.435 | -16.148 | 19.971 | 0.1000 | 1.2000 | H |
| ATOM | 964 | HD22 | LEU | 61 | 32.562 | -15.677 | 18.499 | 0.1000 | 1.2000 | H |
| ATOM | 965 | HD23 | LEU | 61 | 34.128 | -14.990 | 18.836 | 0.1000 | 1.2000 | H |
| ATOM | 966 | C | LEU | 61 | 30.066 | -15.583 | 18.939 | 0.5973 | 1.7000 | C |
| ATOM | 967 | O | LEU | 61 | 30.170 | -15.182 | 17.772 | -0.5679 | 1.5000 | O |
| ATOM | 968 | N | LEU | 62 | 29.997 | -16.874 | 19.264 | -0.4157 | 1.5500 | N |
| ATOM | 969 | H | LEU | 62 | 29.935 | -17.129 | 20.241 | 0.2719 | 1.3000 | H |
| ATOM | 970 | CA | LEU | 62 | 30.047 | -17.960 | 18.292 | -0.0518 | 1.7000 | C |
| ATOM | 971 | HA | LEU | 62 | 30.720 | -17.699 | 17.475 | 0.0922 | 1.2000 | H |
| ATOM | 972 | CB | LEU | 62 | 30.546 | -19.223 | 18.993 | -0.1102 | 1.7000 | C |
| ATOM | 973 | HB2 | LEU | 62 | 29.841 | -19.451 | 19.796 | 0.0457 | 1.2000 | H |
| ATOM | 974 | HB3 | LEU | 62 | 30.505 | -20.053 | 18.286 | 0.0457 | 1.2000 | H |
| ATOM | 975 | CG | LEU | 62 | 31.953 | -19.180 | 19.584 | 0.3531 | 1.7000 | C |
| ATOM | 976 | HG | LEU | 62 | 32.037 | -18.338 | 20.262 | -0.0361 | 1.2000 | H |
| ATOM | 977 | CD1 | LEU | 62 | 32.168 | -20.457 | 20.358 | -0.4121 | 1.7000 | C |
| ATOM | 978 | HD11 | LEU | 62 | 33.138 | -20.432 | 20.851 | 0.1000 | 1.2000 | H |
| ATOM | 979 | HD12 | LEU | 62 | 31.398 | -20.566 | 21.124 | 0.1000 | 1.2000 | H |
| ATOM | 980 | HD13 | LEU | 62 | 32.132 | -21.319 | 19.689 | 0.1000 | 1.2000 | H |
| ATOM | 981 | CD2 | LEU | 62 | 33.005 | -19.026 | 18.487 | -0.4121 | 1.7000 | C |
| ATOM | 982 | HD21 | LEU | 62 | 34.002 | -19.111 | 18.918 | 0.1000 | 1.2000 | H |
| ATOM | 983 | HD22 | LEU | 62 | 32.873 | -19.798 | 17.729 | 0.1000 | 1.2000 | H |
| ATOM | 984 | HD23 | LEU | 62 | 32.914 | -18.050 | 18.013 | 0.1000 | 1.2000 | H |
| ATOM | 985 | C | LEU | 62 | 28.663 | -18.268 | 17.707 | 0.5973 | 1.7000 | C |

| | | | | | | | | | | |
|---|---|---|---|---|---|---|---|---|---|---|
| ATOM | 986 | O | LEU | 62 | 28.515 | -19.169 | 16.877 | -0.5679 | 1.5000 | O |
| ATOM | 987 | N | GLY | 63 | 27.650 | -17.550 | 18.187 | -0.4157 | 1.5500 | N |
| ATOM | 988 | H | GLY | 63 | 27.852 | -16.847 | 18.882 | 0.2719 | 1.3000 | H |
| ATOM | 989 | CA | GLY | 63 | 26.258 | -17.737 | 17.834 | -0.0252 | 1.7000 | C |
| ATOM | 990 | HA2 | GLY | 63 | 26.125 | -18.682 | 17.307 | 0.0698 | 1.2000 | H |
| ATOM | 991 | HA3 | GLY | 63 | 25.682 | -17.795 | 18.758 | 0.0698 | 1.2000 | H |
| ATOM | 992 | C | GLY | 63 | 25.696 | -16.611 | 16.970 | 0.5973 | 1.7000 | C |
| ATOM | 993 | O | GLY | 63 | 25.677 | -16.709 | 15.738 | -0.5679 | 1.5000 | O |
| ATOM | 994 | N | ARG | 64 | 25.133 | -15.592 | 17.635 | -0.3479 | 1.5500 | N |
| ATOM | 995 | H | ARG | 64 | 25.183 | -15.573 | 18.644 | 0.2747 | 1.3000 | H |
| ATOM | 996 | CA | ARG | 64 | 24.447 | -14.493 | 16.947 | -0.2637 | 1.7000 | C |
| ATOM | 997 | HA | ARG | 64 | 24.738 | -14.483 | 15.897 | 0.1560 | 1.2000 | H |
| ATOM | 998 | CB | ARG | 64 | 22.935 | -14.667 | 17.045 | -0.0007 | 1.7000 | C |
| ATOM | 999 | HB2 | ARG | 64 | 22.661 | -14.733 | 18.099 | 0.0327 | 1.2000 | H |
| ATOM | 1000 | HB3 | ARG | 64 | 22.422 | -13.799 | 16.625 | 0.0327 | 1.2000 | H |
| ATOM | 1001 | CG | ARG | 64 | 22.357 | -15.889 | 16.318 | 0.0390 | 1.7000 | C |
| ATOM | 1002 | HG2 | ARG | 64 | 22.850 | -16.797 | 16.665 | 0.0285 | 1.2000 | H |
| ATOM | 1003 | HG3 | ARG | 64 | 21.297 | -15.965 | 16.565 | 0.0285 | 1.2000 | H |
| ATOM | 1004 | CD | ARG | 64 | 22.494 | -15.769 | 14.838 | 0.0486 | 1.7000 | C |
| ATOM | 1005 | HD2 | ARG | 64 | 21.991 | -14.858 | 14.511 | 0.0687 | 1.2000 | H |
| ATOM | 1006 | HD3 | ARG | 64 | 23.545 | -15.686 | 14.562 | 0.0687 | 1.2000 | H |
| ATOM | 1007 | NE | ARG | 64 | 21.895 | -16.896 | 14.140 | -0.5295 | 1.5500 | N |
| ATOM | 1008 | HE | ARG | 64 | 20.933 | -16.790 | 13.857 | 0.3456 | 1.3000 | H |
| ATOM | 1009 | CZ | ARG | 64 | 22.520 | -18.057 | 13.832 | 0.8076 | 1.7000 | C |
| ATOM | 1010 | NH1 | ARG | 64 | 23.783 | -18.270 | 14.156 | -0.8627 | 1.5500 | N |
| ATOM | 1011 | HH11 | ARG | 64 | 24.254 | -19.107 | 13.853 | 0.4478 | 1.3000 | H |
| ATOM | 1012 | HH12 | ARG | 64 | 24.295 | -17.576 | 14.683 | 0.4478 | 1.3000 | H |
| ATOM | 1013 | NH2 | ARG | 64 | 21.848 | -19.000 | 13.189 | -0.8627 | 1.5500 | N |
| ATOM | 1014 | HH21 | ARG | 64 | 22.318 | -19.844 | 12.905 | 0.4478 | 1.3000 | H |
| ATOM | 1015 | HH22 | ARG | 64 | 20.868 | -18.874 | 12.986 | 0.4478 | 1.3000 | H |
| ATOM | 1016 | C | ARG | 64 | 24.807 | -13.103 | 17.480 | 0.7341 | 1.7000 | C |
| ATOM | 1017 | O | ARG | 64 | 24.030 | -12.158 | 17.329 | -0.5894 | 1.5000 | O |
| ATOM | 1018 | N | MET | 65 | 25.978 | -12.972 | 18.085 | -0.4157 | 1.5500 | N |
| ATOM | 1019 | H | MET | 65 | 26.582 | -13.776 | 18.182 | 0.2719 | 1.3000 | H |

| | | | | | | | | | | |
|---|---|---|---|---|---|---|---|---|---|---|
| ATOM | 1020 | CA | MET | 65 | 26.417 | -11.703 | 18.660 | -0.0237 | 1.7000 | C |
| ATOM | 1021 | HA | MET | 65 | 25.694 | -11.463 | 19.442 | 0.0880 | 1.2000 | H |
| ATOM | 1022 | CB | MET | 65 | 27.768 | -11.837 | 19.331 | 0.0342 | 1.7000 | C |
| ATOM | 1023 | HB2 | MET | 65 | 27.743 | -12.667 | 20.022 | 0.0241 | 1.2000 | H |
| ATOM | 1024 | HB3 | MET | 65 | 28.509 | -12.066 | 18.564 | 0.0241 | 1.2000 | H |
| ATOM | 1025 | CG | MET | 65 | 28.195 | -10.622 | 20.087 | 0.0018 | 1.7000 | C |
| ATOM | 1026 | HG2 | MET | 65 | 28.394 | -9.800 | 19.400 | 0.0440 | 1.2000 | H |
| ATOM | 1027 | HG3 | MET | 65 | 27.394 | -10.335 | 20.766 | 0.0440 | 1.2000 | H |
| ATOM | 1028 | SD | MET | 65 | 29.615 | -10.874 | 21.007 | -0.2737 | 1.8000 | S |
| ATOM | 1029 | CE | MET | 65 | 30.834 | -10.844 | 19.771 | -0.0536 | 1.7000 | C |
| ATOM | 1030 | HE1 | MET | 65 | 31.811 | -10.987 | 20.232 | 0.0684 | 1.2000 | H |
| ATOM | 1031 | HE2 | MET | 65 | 30.663 | -11.652 | 19.065 | 0.0684 | 1.2000 | H |
| ATOM | 1032 | HE3 | MET | 65 | 30.812 | -9.889 | 19.245 | 0.0684 | 1.2000 | H |
| ATOM | 1033 | C | MET | 65 | 26.493 | -10.522 | 17.716 | 0.5973 | 1.7000 | C |
| ATOM | 1034 | O | MET | 65 | 26.881 | -10.629 | 16.552 | -0.5679 | 1.5000 | O |
| ATOM | 1035 | N | LEU | 66 | 26.133 | -9.387 | 18.290 | -0.4157 | 1.5500 | N |
| ATOM | 1036 | H | LEU | 66 | 25.848 | -9.723 | 19.198 | 0.2719 | 1.3000 | H |
| ATOM | 1037 | CA | LEU | 66 | 25.880 | -7.979 | 18.486 | -0.0518 | 1.7000 | C |
| ATOM | 1038 | HA | LEU | 66 | 26.332 | -7.441 | 17.657 | 0.0922 | 1.2000 | H |
| ATOM | 1039 | CB | LEU | 66 | 24.389 | -7.661 | 18.528 | -0.1102 | 1.7000 | C |
| ATOM | 1040 | HB2 | LEU | 66 | 23.947 | -7.951 | 17.573 | 0.0457 | 1.2000 | H |
| ATOM | 1041 | HB3 | LEU | 66 | 23.911 | -8.266 | 19.299 | 0.0457 | 1.2000 | H |
| ATOM | 1042 | CG | LEU | 66 | 24.094 | -6.190 | 18.788 | 0.3531 | 1.7000 | C |
| ATOM | 1043 | HG | LEU | 66 | 24.555 | -5.897 | 19.724 | -0.0361 | 1.2000 | H |
| ATOM | 1044 | CD1 | LEU | 66 | 24.677 | -5.345 | 17.677 | -0.4121 | 1.7000 | C |
| ATOM | 1045 | HD11 | LEU | 66 | 24.352 | -4.310 | 17.795 | 0.1000 | 1.2000 | H |
| ATOM | 1046 | HD12 | LEU | 66 | 25.764 | -5.351 | 17.711 | 0.1000 | 1.2000 | H |
| ATOM | 1047 | HD13 | LEU | 66 | 24.337 | -5.706 | 16.705 | 0.1000 | 1.2000 | H |
| ATOM | 1048 | CD2 | LEU | 66 | 22.602 | -5.968 | 18.922 | -0.4121 | 1.7000 | C |
| ATOM | 1049 | HD21 | LEU | 66 | 22.399 | -4.917 | 19.133 | 0.1000 | 1.2000 | H |
| ATOM | 1050 | HD22 | LEU | 66 | 22.097 | -6.249 | 17.997 | 0.1000 | 1.2000 | H |
| ATOM | 1051 | HD23 | LEU | 66 | 22.203 | -6.565 | 19.740 | 0.1000 | 1.2000 | H |
| ATOM | 1052 | C | LEU | 66 | 26.501 | -7.614 | 19.833 | 0.5973 | 1.7000 | C |
| ATOM | 1053 | O | LEU | 66 | 25.903 | -7.917 | 20.864 | -0.5679 | 1.5000 | O |

| | | | | | | | | | | |
|---|---|---|---|---|---|---|---|---|---|---|
| ATOM | 1054 | N | PRO | 67 | 27.707 | -7.038 | 19.893 | -0.2548 | 1.5500 | N |
| ATOM | 1055 | CD | PRO | 67 | 28.475 | -6.753 | 18.670 | 0.0192 | 1.7000 | C |
| ATOM | 1056 | HD2 | PRO | 67 | 28.057 | -5.888 | 18.153 | 0.0391 | 1.2000 | H |
| ATOM | 1057 | HD3 | PRO | 67 | 28.544 | -7.621 | 18.013 | 0.0391 | 1.2000 | H |
| ATOM | 1058 | CG | PRO | 67 | 29.840 | -6.417 | 19.235 | 0.0189 | 1.7000 | C |
| ATOM | 1059 | HG2 | PRO | 67 | 30.364 | -5.702 | 18.602 | 0.0213 | 1.2000 | H |
| ATOM | 1060 | HG3 | PRO | 67 | 30.421 | -7.333 | 19.329 | 0.0213 | 1.2000 | H |
| ATOM | 1061 | CB | PRO | 67 | 29.556 | -5.839 | 20.609 | -0.0070 | 1.7000 | C |
| ATOM | 1062 | HB2 | PRO | 67 | 29.253 | -4.795 | 20.504 | 0.0253 | 1.2000 | H |
| ATOM | 1063 | HB3 | PRO | 67 | 30.424 | -5.910 | 21.263 | 0.0253 | 1.2000 | H |
| ATOM | 1064 | CA | PRO | 67 | 28.390 | -6.685 | 21.127 | -0.0266 | 1.7000 | C |
| ATOM | 1065 | HA | PRO | 67 | 28.755 | -7.590 | 21.613 | 0.0641 | 1.2000 | H |
| ATOM | 1066 | C | PRO | 67 | 27.491 | -5.894 | 22.072 | 0.5896 | 1.7000 | C |
| ATOM | 1067 | O | PRO | 67 | 27.576 | -6.051 | 23.289 | -0.5748 | 1.5000 | O |
| ATOM | 1068 | N | GLN | 68 | 26.580 | -5.083 | 21.515 | -0.4157 | 1.5500 | N |
| ATOM | 1069 | H | GLN | 68 | 26.516 | -5.010 | 20.510 | 0.2719 | 1.3000 | H |
| ATOM | 1070 | CA | GLN | 68 | 25.666 | -4.292 | 22.333 | -0.0031 | 1.7000 | C |
| ATOM | 1071 | HA | GLN | 68 | 26.239 | -3.674 | 23.025 | 0.0850 | 1.2000 | H |
| ATOM | 1072 | CB | GLN | 68 | 24.792 | -3.390 | 21.458 | -0.0036 | 1.7000 | C |
| ATOM | 1073 | HB2 | GLN | 68 | 24.269 | -4.008 | 20.732 | 0.0171 | 1.2000 | H |
| ATOM | 1074 | HB3 | GLN | 68 | 24.034 | -2.930 | 22.096 | 0.0171 | 1.2000 | H |
| ATOM | 1075 | CG | GLN | 68 | 25.523 | -2.281 | 20.745 | -0.0645 | 1.7000 | C |
| ATOM | 1076 | HG2 | GLN | 68 | 25.947 | -1.600 | 21.484 | 0.0352 | 1.2000 | H |
| ATOM | 1077 | HG3 | GLN | 68 | 26.338 | -2.696 | 20.150 | 0.0352 | 1.2000 | H |
| ATOM | 1078 | CD | GLN | 68 | 24.593 | -1.510 | 19.815 | 0.6951 | 1.7000 | C |
| ATOM | 1079 | OE1 | GLN | 68 | 23.521 | -2.001 | 19.444 | -0.6086 | 1.5000 | O |
| ATOM | 1080 | NE2 | GLN | 68 | 24.997 | -0.305 | 19.432 | -0.9407 | 1.5500 | N |
| ATOM | 1081 | HE21 | GLN | 68 | 24.443 | 0.197 | 18.754 | 0.4251 | 1.3000 | H |
| ATOM | 1082 | HE22 | GLN | 68 | 25.841 | 0.095 | 19.805 | 0.4251 | 1.3000 | H |
| ATOM | 1083 | C | GLN | 68 | 24.757 | -5.217 | 23.137 | 0.5973 | 1.7000 | C |
| ATOM | 1084 | O | GLN | 68 | 24.399 | -4.916 | 24.277 | -0.5679 | 1.5000 | O |
| ATOM | 1085 | N | LEU | 69 | 24.361 | -6.340 | 22.532 | -0.4157 | 1.5500 | N |
| ATOM | 1086 | H | LEU | 69 | 24.668 | -6.550 | 21.594 | 0.2719 | 1.3000 | H |
| ATOM | 1087 | CA | LEU | 69 | 23.499 | -7.288 | 23.202 | -0.0518 | 1.7000 | C |

| | | | | | | | | | | |
|---|---|---|---|---|---|---|---|---|---|---|
| ATOM | 1088 | HA | LEU | 69 | 22.599 | -6.771 | 23.539 | 0.0922 | 1.2000 | H |
| ATOM | 1089 | CB | LEU | 69 | 23.119 | -8.458 | 22.291 | -0.1102 | 1.7000 | C |
| ATOM | 1090 | HB2 | LEU | 69 | 22.553 | -8.070 | 21.444 | 0.0457 | 1.2000 | H |
| ATOM | 1091 | HB3 | LEU | 69 | 24.021 | -8.923 | 21.903 | 0.0457 | 1.2000 | H |
| ATOM | 1092 | CG | LEU | 69 | 22.322 | -9.553 | 22.957 | 0.3531 | 1.7000 | C |
| ATOM | 1093 | HG | LEU | 69 | 22.877 | -9.951 | 23.803 | -0.0361 | 1.2000 | H |
| ATOM | 1094 | CD1 | LEU | 69 | 21.004 | -9.001 | 23.455 | -0.4121 | 1.7000 | C |
| ATOM | 1095 | HD11 | LEU | 69 | 20.392 | -9.817 | 23.840 | 0.1000 | 1.2000 | H |
| ATOM | 1096 | HD12 | LEU | 69 | 21.161 | -8.295 | 24.269 | 0.1000 | 1.2000 | H |
| ATOM | 1097 | HD13 | LEU | 69 | 20.464 | -8.510 | 22.645 | 0.1000 | 1.2000 | H |
| ATOM | 1098 | CD2 | LEU | 69 | 22.134 | -10.695 | 21.978 | -0.4121 | 1.7000 | C |
| ATOM | 1099 | HD21 | LEU | 69 | 23.102 | -11.090 | 21.669 | 0.1000 | 1.2000 | H |
| ATOM | 1100 | HD22 | LEU | 69 | 21.572 | -11.498 | 22.456 | 0.1000 | 1.2000 | H |
| ATOM | 1101 | HD23 | LEU | 69 | 21.587 | -10.355 | 21.097 | 0.1000 | 1.2000 | H |
| ATOM | 1102 | C | LEU | 69 | 24.229 | -7.845 | 24.373 | 0.5973 | 1.7000 | C |
| ATOM | 1103 | O | LEU | 69 | 23.728 | -7.832 | 25.498 | -0.5679 | 1.5000 | O |
| ATOM | 1104 | N | VAL | 70 | 25.449 | -8.297 | 24.117 | -0.4157 | 1.5500 | N |
| ATOM | 1105 | H | VAL | 70 | 25.830 | -8.282 | 23.182 | 0.2719 | 1.3000 | H |
| ATOM | 1106 | CA | VAL | 70 | 26.177 | -8.931 | 25.186 | -0.0875 | 1.7000 | C |
| ATOM | 1107 | HA | VAL | 70 | 25.576 | -9.757 | 25.568 | 0.0969 | 1.2000 | H |
| ATOM | 1108 | CB | VAL | 70 | 27.521 | -9.485 | 24.746 | 0.2985 | 1.7000 | C |
| ATOM | 1109 | HB | VAL | 70 | 28.073 | -8.705 | 24.231 | -0.0297 | 1.2000 | H |
| ATOM | 1110 | CG1 | VAL | 70 | 28.304 | -9.955 | 25.989 | -0.3192 | 1.7000 | C |
| ATOM | 1111 | HG11 | VAL | 70 | 29.178 | -10.523 | 25.682 | 0.0791 | 1.2000 | H |
| ATOM | 1112 | HG12 | VAL | 70 | 28.659 | -9.110 | 26.577 | 0.0791 | 1.2000 | H |
| ATOM | 1113 | HG13 | VAL | 70 | 27.681 | -10.604 | 26.609 | 0.0791 | 1.2000 | H |
| ATOM | 1114 | CG2 | VAL | 70 | 27.303 | -10.606 | 23.824 | -0.3192 | 1.7000 | C |
| ATOM | 1115 | HG21 | VAL | 70 | 28.259 | -11.042 | 23.539 | 0.0791 | 1.2000 | H |
| ATOM | 1116 | HG22 | VAL | 70 | 26.704 | -11.380 | 24.307 | 0.0791 | 1.2000 | H |
| ATOM | 1117 | HG23 | VAL | 70 | 26.790 | -10.268 | 22.924 | 0.0791 | 1.2000 | H |
| ATOM | 1118 | C | VAL | 70 | 26.412 | -7.980 | 26.317 | 0.5973 | 1.7000 | C |
| ATOM | 1119 | O | VAL | 70 | 26.161 | -8.353 | 27.452 | -0.5679 | 1.5000 | O |
| ATOM | 1120 | N | CYX | 71 | 26.817 | -6.740 | 26.028 | -0.4157 | 1.5500 | N |
| ATOM | 1121 | H | CYX | 71 | 26.981 | -6.465 | 25.071 | 0.2719 | 1.3000 | H |

| | | | | | | | | | | |
|---|---|---|---|---|---|---|---|---|---|---|
| ATOM | 1122 | CA | CYX | 71 | 27.093 | -5.791 | 27.099 | 0.0429 | 1.7000 | C |
| ATOM | 1123 | HA | CYX | 71 | 27.837 | -6.226 | 27.767 | 0.0766 | 1.2000 | H |
| ATOM | 1124 | CB | CYX | 71 | 27.642 | -4.486 | 26.557 | -0.0790 | 1.7000 | C |
| ATOM | 1125 | HB2 | CYX | 71 | 26.925 | -4.064 | 25.852 | 0.0910 | 1.2000 | H |
| ATOM | 1126 | HB3 | CYX | 71 | 27.736 | -3.803 | 27.399 | 0.0910 | 1.2000 | H |
| ATOM | 1127 | SG | CYX | 71 | 29.266 | -4.579 | 25.752 | -0.1081 | 1.8000 | S |
| ATOM | 1128 | C | CYX | 71 | 25.845 | -5.475 | 27.922 | 0.5973 | 1.7000 | C |
| ATOM | 1129 | O | CYX | 71 | 25.942 | -5.109 | 29.105 | -0.5679 | 1.5000 | O |
| ATOM | 1130 | N | ARG | 72 | 24.656 | -5.606 | 27.322 | -0.3479 | 1.5500 | N |
| ATOM | 1131 | H | ARG | 72 | 24.583 | -5.910 | 26.360 | 0.2747 | 1.3000 | H |
| ATOM | 1132 | CA | ARG | 72 | 23.445 | -5.388 | 28.095 | -0.2637 | 1.7000 | C |
| ATOM | 1133 | HA | ARG | 72 | 23.592 | -4.566 | 28.797 | 0.1560 | 1.2000 | H |
| ATOM | 1134 | CB | ARG | 72 | 22.287 | -5.012 | 27.190 | -0.0007 | 1.7000 | C |
| ATOM | 1135 | HB2 | ARG | 72 | 22.207 | -5.748 | 26.388 | 0.0327 | 1.2000 | H |
| ATOM | 1136 | HB3 | ARG | 72 | 21.369 | -5.064 | 27.778 | 0.0327 | 1.2000 | H |
| ATOM | 1137 | CG | ARG | 72 | 22.392 | -3.619 | 26.584 | 0.0390 | 1.7000 | C |
| ATOM | 1138 | HG2 | ARG | 72 | 22.431 | -2.880 | 27.387 | 0.0285 | 1.2000 | H |
| ATOM | 1139 | HG3 | ARG | 72 | 23.317 | -3.530 | 26.020 | 0.0285 | 1.2000 | H |
| ATOM | 1140 | CD | ARG | 72 | 21.251 | -3.314 | 25.683 | 0.0486 | 1.7000 | C |
| ATOM | 1141 | HD2 | ARG | 72 | 21.452 | -2.385 | 25.146 | 0.0687 | 1.2000 | H |
| ATOM | 1142 | HD3 | ARG | 72 | 21.159 | -4.111 | 24.943 | 0.0687 | 1.2000 | H |
| ATOM | 1143 | NE | ARG | 72 | 19.997 | -3.206 | 26.423 | -0.5295 | 1.5500 | N |
| ATOM | 1144 | HE | ARG | 72 | 19.418 | -4.029 | 26.489 | 0.3456 | 1.3000 | H |
| ATOM | 1145 | CZ | ARG | 72 | 19.594 | -2.076 | 27.041 | 0.8076 | 1.7000 | C |
| ATOM | 1146 | NH1 | ARG | 72 | 20.313 | -0.999 | 26.991 | -0.8627 | 1.5500 | N |
| ATOM | 1147 | HH11 | ARG | 72 | 21.157 | -0.971 | 26.448 | 0.4478 | 1.3000 | H |
| ATOM | 1148 | HH12 | ARG | 72 | 20.022 | -0.184 | 27.519 | 0.4478 | 1.3000 | H |
| ATOM | 1149 | NH2 | ARG | 72 | 18.478 | -2.007 | 27.722 | -0.8627 | 1.5500 | N |
| ATOM | 1150 | HH21 | ARG | 72 | 17.847 | -2.790 | 27.756 | 0.4478 | 1.3000 | H |
| ATOM | 1151 | HH22 | ARG | 72 | 18.254 | -1.146 | 28.205 | 0.4478 | 1.3000 | H |
| ATOM | 1152 | C | ARG | 72 | 23.094 | -6.639 | 28.913 | 0.7341 | 1.7000 | C |
| ATOM | 1153 | O | ARG | 72 | 22.742 | -6.535 | 30.089 | -0.5894 | 1.5000 | O |
| ATOM | 1154 | N | LEU | 73 | 23.310 | -7.831 | 28.343 | -0.4157 | 1.5500 | N |
| ATOM | 1155 | H | LEU | 73 | 23.659 | -7.874 | 27.397 | 0.2719 | 1.3000 | H |

| | | | | | | | | | | |
|---|---|---|---|---|---|---|---|---|---|---|
| ATOM | 1156 | CA | LEU | 73 | 22.971 | -9.093 | 29.014 | -0.0518 | 1.7000 | C |
| ATOM | 1157 | HA | LEU | 73 | 21.910 | -9.093 | 29.262 | 0.0922 | 1.2000 | H |
| ATOM | 1158 | CB | LEU | 73 | 23.301 | -10.306 | 28.115 | -0.1102 | 1.7000 | C |
| ATOM | 1159 | HB2 | LEU | 73 | 24.344 | -10.224 | 27.808 | 0.0457 | 1.2000 | H |
| ATOM | 1160 | HB3 | LEU | 73 | 23.227 | -11.210 | 28.724 | 0.0457 | 1.2000 | H |
| ATOM | 1161 | CG | LEU | 73 | 22.445 | -10.520 | 26.854 | 0.3531 | 1.7000 | C |
| ATOM | 1162 | HG | LEU | 73 | 22.425 | -9.627 | 26.247 | -0.0361 | 1.2000 | H |
| ATOM | 1163 | CD1 | LEU | 73 | 23.055 | -11.670 | 26.029 | -0.4121 | 1.7000 | C |
| ATOM | 1164 | HD11 | LEU | 73 | 22.435 | -11.868 | 25.154 | 0.1000 | 1.2000 | H |
| ATOM | 1165 | HD12 | LEU | 73 | 24.058 | -11.405 | 25.696 | 0.1000 | 1.2000 | H |
| ATOM | 1166 | HD13 | LEU | 73 | 23.107 | -12.576 | 26.634 | 0.1000 | 1.2000 | H |
| ATOM | 1167 | CD2 | LEU | 73 | 21.028 | -10.823 | 27.251 | -0.4121 | 1.7000 | C |
| ATOM | 1168 | HD21 | LEU | 73 | 20.442 | -11.050 | 26.360 | 0.1000 | 1.2000 | H |
| ATOM | 1169 | HD22 | LEU | 73 | 20.995 | -11.683 | 27.921 | 0.1000 | 1.2000 | H |
| ATOM | 1170 | HD23 | LEU | 73 | 20.576 | -9.962 | 27.742 | 0.1000 | 1.2000 | H |
| ATOM | 1171 | C | LEU | 73 | 23.753 | -9.273 | 30.304 | 0.5973 | 1.7000 | C |
| ATOM | 1172 | O | LEU | 73 | 23.256 | -9.834 | 31.283 | -0.5679 | 1.5000 | O |
| ATOM | 1173 | N | VAL | 74 | 24.979 | -8.782 | 30.295 | -0.4157 | 1.5500 | N |
| ATOM | 1174 | H | VAL | 74 | 25.320 | -8.348 | 29.450 | 0.2719 | 1.3000 | H |
| ATOM | 1175 | CA | VAL | 74 | 25.900 | -8.915 | 31.409 | -0.0875 | 1.7000 | C |
| ATOM | 1176 | HA | VAL | 74 | 25.610 | -9.751 | 32.050 | 0.0969 | 1.2000 | H |
| ATOM | 1177 | CB | VAL | 74 | 27.266 | -9.185 | 30.832 | 0.2985 | 1.7000 | C |
| ATOM | 1178 | HB | VAL | 74 | 27.974 | -9.351 | 31.632 | -0.0297 | 1.2000 | H |
| ATOM | 1179 | CG1 | VAL | 74 | 27.235 | -10.407 | 29.975 | -0.3192 | 1.7000 | C |
| ATOM | 1180 | HG11 | VAL | 74 | 28.232 | -10.627 | 29.596 | 0.0791 | 1.2000 | H |
| ATOM | 1181 | HG12 | VAL | 74 | 26.900 | -11.246 | 30.579 | 0.0791 | 1.2000 | H |
| ATOM | 1182 | HG13 | VAL | 74 | 26.579 | -10.304 | 29.112 | 0.0791 | 1.2000 | H |
| ATOM | 1183 | CG2 | VAL | 74 | 27.659 | -8.034 | 30.123 | -0.3192 | 1.7000 | C |
| ATOM | 1184 | HG21 | VAL | 74 | 28.534 | -8.261 | 29.518 | 0.0791 | 1.2000 | H |
| ATOM | 1185 | HG22 | VAL | 74 | 26.910 | -7.610 | 29.478 | 0.0791 | 1.2000 | H |
| ATOM | 1186 | HG23 | VAL | 74 | 27.914 | -7.231 | 30.790 | 0.0791 | 1.2000 | H |
| ATOM | 1187 | C | VAL | 74 | 26.009 | -7.654 | 32.267 | 0.5973 | 1.7000 | C |
| ATOM | 1188 | O | VAL | 74 | 26.833 | -7.594 | 33.176 | -0.5679 | 1.5000 | O |
| ATOM | 1189 | N | LEU | 75 | 25.226 | -6.625 | 31.943 | -0.4157 | 1.5500 | N |

| | | | | | | | | | | |
|---|---|---|---|---|---|---|---|---|---|---|
| ATOM | 1190 | H | LEU | 75 | 24.556 | -6.728 | 31.193 | 0.2719 | 1.3000 | H |
| ATOM | 1191 | CA | LEU | 75 | 25.266 | -5.344 | 32.645 | -0.0518 | 1.7000 | C |
| ATOM | 1192 | HA | LEU | 75 | 24.598 | -4.663 | 32.115 | 0.0922 | 1.2000 | H |
| ATOM | 1193 | CB | LEU | 75 | 24.773 | -5.521 | 34.085 | -0.1102 | 1.7000 | C |
| ATOM | 1194 | HB2 | LEU | 75 | 25.463 | -6.149 | 34.647 | 0.0457 | 1.2000 | H |
| ATOM | 1195 | HB3 | LEU | 75 | 24.762 | -4.544 | 34.571 | 0.0457 | 1.2000 | H |
| ATOM | 1196 | CG | LEU | 75 | 23.371 | -6.103 | 34.196 | 0.3531 | 1.7000 | C |
| ATOM | 1197 | HG | LEU | 75 | 23.340 | -7.098 | 33.749 | -0.0361 | 1.2000 | H |
| ATOM | 1198 | CD1 | LEU | 75 | 22.983 | -6.241 | 35.648 | -0.4121 | 1.7000 | C |
| ATOM | 1199 | HD11 | LEU | 75 | 21.979 | -6.661 | 35.718 | 0.1000 | 1.2000 | H |
| ATOM | 1200 | HD12 | LEU | 75 | 23.673 | -6.919 | 36.152 | 0.1000 | 1.2000 | H |
| ATOM | 1201 | HD13 | LEU | 75 | 22.996 | -5.267 | 36.138 | 0.1000 | 1.2000 | H |
| ATOM | 1202 | CD2 | LEU | 75 | 22.434 | -5.210 | 33.452 | -0.4121 | 1.7000 | C |
| ATOM | 1203 | HD21 | LEU | 75 | 21.408 | -5.521 | 33.646 | 0.1000 | 1.2000 | H |
| ATOM | 1204 | HD22 | LEU | 75 | 22.549 | -4.173 | 33.772 | 0.1000 | 1.2000 | H |
| ATOM | 1205 | HD23 | LEU | 75 | 22.595 | -5.289 | 32.381 | 0.1000 | 1.2000 | H |
| ATOM | 1206 | C | LEU | 75 | 26.639 | -4.648 | 32.652 | 0.5973 | 1.7000 | C |
| ATOM | 1207 | O | LEU | 75 | 27.051 | -4.095 | 33.672 | -0.5679 | 1.5000 | O |
| ATOM | 1208 | N | ARG | 76 | 27.365 | -4.692 | 31.526 | -0.3479 | 1.5500 | N |
| ATOM | 1209 | H | ARG | 76 | 27.004 | -5.175 | 30.716 | 0.2747 | 1.3000 | H |
| ATOM | 1210 | CA | ARG | 76 | 28.639 | -3.979 | 31.420 | -0.2637 | 1.7000 | C |
| ATOM | 1211 | HA | ARG | 76 | 29.034 | -3.802 | 32.421 | 0.1560 | 1.2000 | H |
| ATOM | 1212 | CB | ARG | 76 | 29.715 | -4.755 | 30.683 | -0.0007 | 1.7000 | C |
| ATOM | 1213 | HB2 | ARG | 76 | 29.306 | -5.081 | 29.727 | 0.0327 | 1.2000 | H |
| ATOM | 1214 | HB3 | ARG | 76 | 30.545 | -4.070 | 30.496 | 0.0327 | 1.2000 | H |
| ATOM | 1215 | CG | ARG | 76 | 30.296 | -5.981 | 31.431 | 0.0390 | 1.7000 | C |
| ATOM | 1216 | HG2 | ARG | 76 | 30.703 | -5.611 | 32.369 | 0.0285 | 1.2000 | H |
| ATOM | 1217 | HG3 | ARG | 76 | 29.547 | -6.696 | 31.738 | 0.0285 | 1.2000 | H |
| ATOM | 1218 | CD | ARG | 76 | 31.439 | -6.698 | 30.678 | 0.0486 | 1.7000 | C |
| ATOM | 1219 | HD2 | ARG | 76 | 32.172 | -5.951 | 30.373 | 0.0687 | 1.2000 | H |
| ATOM | 1220 | HD3 | ARG | 76 | 31.928 | -7.381 | 31.363 | 0.0687 | 1.2000 | H |
| ATOM | 1221 | NE | ARG | 76 | 31.041 | -7.433 | 29.519 | -0.5295 | 1.5500 | N |
| ATOM | 1222 | HE | ARG | 76 | 30.082 | -7.334 | 29.224 | 0.3456 | 1.3000 | H |
| ATOM | 1223 | CZ | ARG | 76 | 31.857 | -8.232 | 28.791 | 0.8076 | 1.7000 | C |

| | | | | | | | | | | |
|---|---|---|---|---|---|---|---|---|---|---|
| ATOM | 1224 | NH1 | ARG | 76 | 33.107 | -8.399 | 29.128 | -0.8627 | 1.5500 | N |
| ATOM | 1225 | HH11 | ARG | 76 | 33.705 | -8.994 | 28.575 | 0.4478 | 1.3000 | H |
| ATOM | 1226 | HH12 | ARG | 76 | 33.463 | -7.940 | 29.953 | 0.4478 | 1.3000 | H |
| ATOM | 1227 | NH2 | ARG | 76 | 31.406 | -8.858 | 27.738 | -0.8627 | 1.5500 | N |
| ATOM | 1228 | HH21 | ARG | 76 | 30.443 | -8.742 | 27.461 | 0.4478 | 1.3000 | H |
| ATOM | 1229 | HH22 | ARG | 76 | 32.019 | -9.456 | 27.206 | 0.4478 | 1.3000 | H |
| ATOM | 1230 | C | ARG | 76 | 28.428 | -2.595 | 30.811 | 0.7341 | 1.7000 | C |
| ATOM | 1231 | O | ARG | 76 | 29.233 | -1.690 | 31.018 | -0.5894 | 1.5000 | O |
| ATOM | 1232 | N | CYX | 77 | 27.331 | -2.432 | 30.070 | -0.4157 | 1.5500 | N |
| ATOM | 1233 | H | CYX | 77 | 26.692 | -3.206 | 29.953 | 0.2719 | 1.3000 | H |
| ATOM | 1234 | CA | CYX | 77 | 26.939 | -1.126 | 29.529 | 0.0429 | 1.7000 | C |
| ATOM | 1235 | HA | CYX | 77 | 27.685 | -0.388 | 29.818 | 0.0766 | 1.2000 | H |
| ATOM | 1236 | CB | CYX | 77 | 26.873 | -1.083 | 28.010 | -0.0790 | 1.7000 | C |
| ATOM | 1237 | HB2 | CYX | 77 | 26.260 | -1.918 | 27.669 | 0.0910 | 1.2000 | H |
| ATOM | 1238 | HB3 | CYX | 77 | 26.392 | -0.159 | 27.687 | 0.0910 | 1.2000 | H |
| ATOM | 1239 | SG | CYX | 77 | 28.475 | -1.178 | 27.225 | -0.1081 | 1.8000 | S |
| ATOM | 1240 | C | CYX | 77 | 25.634 | -0.636 | 30.121 | 0.5973 | 1.7000 | C |
| ATOM | 1241 | O | CYX | 77 | 24.827 | -1.421 | 30.612 | -0.5679 | 1.5000 | O |
| ATOM | 1242 | N | SER | 78 | 25.432 | 0.677 | 30.075 | -0.4157 | 1.5500 | N |
| ATOM | 1243 | H | SER | 78 | 26.132 | 1.282 | 29.675 | 0.2719 | 1.3000 | H |
| ATOM | 1244 | CA | SER | 78 | 24.222 | 1.282 | 30.613 | -0.0249 | 1.7000 | C |
| ATOM | 1245 | HA | SER | 78 | 24.199 | 1.108 | 31.690 | 0.0843 | 1.2000 | H |
| ATOM | 1246 | CB | SER | 78 | 24.231 | 2.776 | 30.379 | 0.2117 | 1.7000 | C |
| ATOM | 1247 | HB2 | SER | 78 | 25.076 | 3.219 | 30.907 | 0.0352 | 1.2000 | H |
| ATOM | 1248 | HB3 | SER | 78 | 24.332 | 2.980 | 29.312 | 0.0352 | 1.2000 | H |
| ATOM | 1249 | OG | SER | 78 | 23.044 | 3.359 | 30.844 | -0.6546 | 1.5000 | O |
| ATOM | 1250 | HG | SER | 78 | 22.301 | 3.056 | 30.307 | 0.4275 | 1.2000 | H |
| ATOM | 1251 | C | SER | 78 | 22.985 | 0.671 | 29.980 | 0.5973 | 1.7000 | C |
| ATOM | 1252 | O | SER | 78 | 22.967 | 0.399 | 28.775 | -0.5679 | 1.5000 | O |
| ATOM | 1253 | N | MET | 79 | 21.978 | 0.445 | 30.815 | -0.3821 | 1.5500 | N |
| ATOM | 1254 | H | MET | 79 | 22.066 | 0.721 | 31.781 | 0.2681 | 1.3000 | H |
| ATOM | 1255 | CA | MET | 79 | 20.727 | -0.153 | 30.399 | -0.2597 | 1.7000 | C |
| ATOM | 1256 | HA | MET | 79 | 20.908 | -0.791 | 29.537 | 0.1277 | 1.2000 | H |
| ATOM | 1257 | CB | MET | 79 | 20.202 | -1.021 | 31.529 | -0.0236 | 1.7000 | C |

```
ATOM   1258  HB2 MET    79      19.999  -0.393  32.398  0.0480  1.2000       H
ATOM   1259  HB3 MET    79      19.262  -1.480  31.225  0.0480  1.2000       H
ATOM   1260  CG  MET    79      21.195  -2.091  31.927  0.0492  1.7000       C
ATOM   1261  HG2 MET    79      22.115  -1.633  32.291  0.0317  1.2000       H
ATOM   1262  HG3 MET    79      20.771  -2.681  32.740  0.0317  1.2000       H
ATOM   1263  SD  MET    79      21.592  -3.187  30.561 -0.2692  1.8000       S
ATOM   1264  CE  MET    79      20.048  -4.060  30.405 -0.0376  1.7000       C
ATOM   1265  HE1 MET    79      19.268  -3.386  30.057  0.0625  1.2000       H
ATOM   1266  HE2 MET    79      20.162  -4.871  29.689  0.0625  1.2000       H
ATOM   1267  HE3 MET    79      19.767  -4.482  31.370  0.0625  1.2000       H
ATOM   1268  C   MET    79      19.710   0.901  29.972  0.8013  1.7000       C
ATOM   1269  O   MET    79      19.022   0.739  28.957 -0.8105  1.5000       O
ATOM   1270  OXT MET    79      19.753   2.005  30.511 -0.8105  1.5000       O
TER    1271      MET    79
END
```

SP-B Open PDB

| | | | | | | | | | | |
|---|---|---|---|---|---|---|---|---|---|---|
| ATOM | 1 | N | PHE | 1 | 34.501 | -20.742 | -14.102 | 0.1737 | 1.5500 | N |
| ATOM | 2 | H1 | PHE | 1 | 35.098 | -20.743 | -13.287 | 0.1921 | 1.3000 | H |
| ATOM | 3 | H2 | PHE | 1 | 33.628 | -20.287 | -13.872 | 0.1921 | 1.3000 | H |
| ATOM | 4 | H3 | PHE | 1 | 34.323 | -21.693 | -14.393 | 0.1921 | 1.3000 | H |
| ATOM | 5 | CA | PHE | 1 | 35.151 | -20.011 | -15.186 | 0.0733 | 1.7000 | C |
| ATOM | 6 | HA | PHE | 1 | 36.131 | -20.454 | -15.351 | 0.1041 | 1.2000 | H |
| ATOM | 7 | CB | PHE | 1 | 34.345 | -20.133 | -16.485 | 0.0330 | 1.7000 | C |
| ATOM | 8 | HB2 | PHE | 1 | 33.304 | -19.872 | -16.288 | 0.0104 | 1.2000 | H |
| ATOM | 9 | HB3 | PHE | 1 | 34.724 | -19.406 | -17.205 | 0.0104 | 1.2000 | H |
| ATOM | 10 | CG | PHE | 1 | 34.413 | -21.495 | -17.130 | 0.0031 | 1.7000 | C |
| ATOM | 11 | CD1 | PHE | 1 | 33.401 | -22.430 | -16.951 | -0.1392 | 1.7000 | C |
| ATOM | 12 | HD1 | PHE | 1 | 32.539 | -22.186 | -16.347 | 0.1374 | 1.2000 | H |
| ATOM | 13 | CE1 | PHE | 1 | 33.474 | -23.677 | -17.555 | -0.1602 | 1.7000 | C |
| ATOM | 14 | HE1 | PHE | 1 | 32.681 | -24.396 | -17.408 | 0.1433 | 1.2000 | H |
| ATOM | 15 | CZ | PHE | 1 | 34.558 | -24.001 | -18.346 | -0.1208 | 1.7000 | C |
| ATOM | 16 | HZ | PHE | 1 | 34.612 | -24.970 | -18.822 | 0.1329 | 1.2000 | H |
| ATOM | 17 | CE2 | PHE | 1 | 35.570 | -23.080 | -18.534 | -0.1603 | 1.7000 | C |
| ATOM | 18 | HE2 | PHE | 1 | 36.419 | -23.331 | -19.153 | 0.1433 | 1.2000 | H |
| ATOM | 19 | CD2 | PHE | 1 | 35.496 | -21.838 | -17.931 | -0.1391 | 1.7000 | C |
| ATOM | 20 | HD2 | PHE | 1 | 36.291 | -21.125 | -18.095 | 0.1374 | 1.2000 | H |
| ATOM | 21 | C | PHE | 1 | 35.325 | -18.527 | -14.831 | 0.6123 | 1.7000 | C |
| ATOM | 22 | O | PHE | 1 | 34.554 | -18.004 | -14.027 | -0.5713 | 1.5000 | O |
| ATOM | 23 | N | PRO | 2 | 36.352 | -17.833 | -15.366 | -0.2548 | 1.5500 | N |
| ATOM | 24 | CD | PRO | 2 | 37.418 | -18.514 | -16.157 | 0.0192 | 1.7000 | C |
| ATOM | 25 | HD2 | PRO | 2 | 37.067 | -18.705 | -17.172 | 0.0391 | 1.2000 | H |
| ATOM | 26 | HD3 | PRO | 2 | 37.775 | -19.427 | -15.679 | 0.0391 | 1.2000 | H |
| ATOM | 27 | CG | PRO | 2 | 38.516 | -17.470 | -16.169 | 0.0189 | 1.7000 | C |
| ATOM | 28 | HG2 | PRO | 2 | 39.125 | -17.541 | -17.071 | 0.0213 | 1.2000 | H |
| ATOM | 29 | HG3 | PRO | 2 | 39.142 | -17.598 | -15.284 | 0.0213 | 1.2000 | H |
| ATOM | 30 | CB | PRO | 2 | 37.803 | -16.145 | -16.084 | -0.0070 | 1.7000 | C |
| ATOM | 31 | HB2 | PRO | 2 | 37.485 | -15.852 | -17.087 | 0.0253 | 1.2000 | H |
| ATOM | 32 | HB3 | PRO | 2 | 38.445 | -15.373 | -15.658 | 0.0253 | 1.2000 | H |
| ATOM | 33 | CA | PRO | 2 | 36.583 | -16.407 | -15.195 | -0.0266 | 1.7000 | C |

| | | | | | | | | | | |
|---|---|---|---|---|---|---|---|---|---|---|
| ATOM | 34 | HA | PRO | 2 | 36.814 | -16.188 | -14.151 | 0.0641 | 1.2000 | H |
| ATOM | 35 | C | PRO | 2 | 35.390 | -15.605 | -15.681 | 0.5896 | 1.7000 | C |
| ATOM | 36 | O | PRO | 2 | 34.727 | -15.987 | -16.650 | -0.5748 | 1.5000 | O |
| ATOM | 37 | N | ILE | 3 | 35.135 | -14.487 | -15.026 | -0.4157 | 1.5500 | N |
| ATOM | 38 | H | ILE | 3 | 35.727 | -14.207 | -14.257 | 0.2719 | 1.3000 | H |
| ATOM | 39 | CA | ILE | 3 | 34.043 | -13.607 | -15.396 | -0.0597 | 1.7000 | C |
| ATOM | 40 | HA | ILE | 3 | 33.273 | -14.197 | -15.880 | 0.0869 | 1.2000 | H |
| ATOM | 41 | CB | ILE | 3 | 33.434 | -12.985 | -14.142 | 0.1303 | 1.7000 | C |
| ATOM | 42 | HB | ILE | 3 | 34.193 | -12.397 | -13.623 | 0.0187 | 1.2000 | H |
| ATOM | 43 | CG2 | ILE | 3 | 32.318 | -12.076 | -14.524 | -0.3204 | 1.7000 | C |
| ATOM | 44 | HG21 | ILE | 3 | 31.753 | -11.818 | -13.639 | 0.0882 | 1.2000 | H |
| ATOM | 45 | HG22 | ILE | 3 | 32.711 | -11.151 | -14.940 | 0.0882 | 1.2000 | H |
| ATOM | 46 | HG23 | ILE | 3 | 31.647 | -12.546 | -15.241 | 0.0882 | 1.2000 | H |
| ATOM | 47 | CG1 | ILE | 3 | 32.960 | -14.094 | -13.196 | -0.0430 | 1.7000 | C |
| ATOM | 48 | HG12 | ILE | 3 | 33.801 | -14.699 | -12.857 | 0.0236 | 1.2000 | H |
| ATOM | 49 | HG13 | ILE | 3 | 32.534 | -13.637 | -12.305 | 0.0236 | 1.2000 | H |
| ATOM | 50 | CD1 | ILE | 3 | 31.921 | -14.990 | -13.794 | -0.0660 | 1.7000 | C |
| ATOM | 51 | HD11 | ILE | 3 | 31.587 | -15.689 | -13.027 | 0.0186 | 1.2000 | H |
| ATOM | 52 | HD12 | ILE | 3 | 31.052 | -14.425 | -14.128 | 0.0186 | 1.2000 | H |
| ATOM | 53 | HD13 | ILE | 3 | 32.325 | -15.582 | -14.614 | 0.0186 | 1.2000 | H |
| ATOM | 54 | C | ILE | 3 | 34.524 | -12.495 | -16.335 | 0.5973 | 1.7000 | C |
| ATOM | 55 | O | ILE | 3 | 35.524 | -11.849 | -16.033 | -0.5679 | 1.5000 | O |
| ATOM | 56 | N | PRO | 4 | 33.893 | -12.257 | -17.496 | -0.2548 | 1.5500 | N |
| ATOM | 57 | CD | PRO | 4 | 32.784 | -13.106 | -17.982 | 0.0192 | 1.7000 | C |
| ATOM | 58 | HD2 | PRO | 4 | 31.864 | -12.878 | -17.444 | 0.0391 | 1.2000 | H |
| ATOM | 59 | HD3 | PRO | 4 | 33.023 | -14.169 | -17.945 | 0.0391 | 1.2000 | H |
| ATOM | 60 | CG | PRO | 4 | 32.672 | -12.653 | -19.431 | 0.0189 | 1.7000 | C |
| ATOM | 61 | HG2 | PRO | 4 | 31.649 | -12.741 | -19.798 | 0.0213 | 1.2000 | H |
| ATOM | 62 | HG3 | PRO | 4 | 33.346 | -13.249 | -20.049 | 0.0213 | 1.2000 | H |
| ATOM | 63 | CB | PRO | 4 | 33.141 | -11.209 | -19.425 | -0.0070 | 1.7000 | C |
| ATOM | 64 | HB2 | PRO | 4 | 32.317 | -10.569 | -19.104 | 0.0253 | 1.2000 | H |
| ATOM | 65 | HB3 | PRO | 4 | 33.493 | -10.899 | -20.409 | 0.0253 | 1.2000 | H |
| ATOM | 66 | CA | PRO | 4 | 34.275 | -11.191 | -18.396 | -0.0266 | 1.7000 | C |
| ATOM | 67 | HA | PRO | 4 | 35.237 | -11.412 | -18.862 | 0.0641 | 1.2000 | H |

| | | | | | | | | | | |
|---|---|---|---|---|---|---|---|---|---|---|
| ATOM | 68 | C | PRO | 4 | 34.300 | -9.892 | -17.619 | 0.5896 | 1.7000 | C |
| ATOM | 69 | O | PRO | 4 | 33.362 | -9.598 | -16.871 | -0.5748 | 1.5000 | O |
| ATOM | 70 | N | LEU | 5 | 35.325 | -9.079 | -17.829 | -0.4157 | 1.5500 | N |
| ATOM | 71 | H | LEU | 5 | 36.084 | -9.340 | -18.441 | 0.2719 | 1.3000 | H |
| ATOM | 72 | CA | LEU | 5 | 35.398 | -7.834 | -17.087 | -0.0518 | 1.7000 | C |
| ATOM | 73 | HA | LEU | 5 | 35.457 | -8.136 | -16.041 | 0.0922 | 1.2000 | H |
| ATOM | 74 | CB | LEU | 5 | 36.690 | -7.074 | -17.399 | -0.1102 | 1.7000 | C |
| ATOM | 75 | HB2 | LEU | 5 | 37.532 | -7.718 | -17.136 | 0.0457 | 1.2000 | H |
| ATOM | 76 | HB3 | LEU | 5 | 36.762 | -6.886 | -18.468 | 0.0457 | 1.2000 | H |
| ATOM | 77 | CG | LEU | 5 | 36.822 | -5.778 | -16.683 | 0.3531 | 1.7000 | C |
| ATOM | 78 | HG | LEU | 5 | 35.998 | -5.118 | -16.947 | -0.0361 | 1.2000 | H |
| ATOM | 79 | CD1 | LEU | 5 | 36.811 | -6.033 | -15.186 | -0.4121 | 1.7000 | C |
| ATOM | 80 | HD11 | LEU | 5 | 37.069 | -5.114 | -14.661 | 0.1000 | 1.2000 | H |
| ATOM | 81 | HD12 | LEU | 5 | 35.828 | -6.338 | -14.838 | 0.1000 | 1.2000 | H |
| ATOM | 82 | HD13 | LEU | 5 | 37.543 | -6.798 | -14.921 | 0.1000 | 1.2000 | H |
| ATOM | 83 | CD2 | LEU | 5 | 38.111 | -5.114 | -17.103 | -0.4121 | 1.7000 | C |
| ATOM | 84 | HD21 | LEU | 5 | 38.247 | -4.185 | -16.550 | 0.1000 | 1.2000 | H |
| ATOM | 85 | HD22 | LEU | 5 | 38.960 | -5.768 | -16.897 | 0.1000 | 1.2000 | H |
| ATOM | 86 | HD23 | LEU | 5 | 38.083 | -4.885 | -18.169 | 0.1000 | 1.2000 | H |
| ATOM | 87 | C | LEU | 5 | 34.133 | -6.961 | -17.213 | 0.5973 | 1.7000 | C |
| ATOM | 88 | O | LEU | 5 | 33.680 | -6.444 | -16.195 | -0.5679 | 1.5000 | O |
| ATOM | 89 | N | PRO | 6 | 33.534 | -6.745 | -18.410 | -0.2548 | 1.5500 | N |
| ATOM | 90 | CD | PRO | 6 | 34.123 | -7.188 | -19.696 | 0.0192 | 1.7000 | C |
| ATOM | 91 | HD2 | PRO | 6 | 33.868 | -8.230 | -19.888 | 0.0391 | 1.2000 | H |
| ATOM | 92 | HD3 | PRO | 6 | 35.192 | -7.020 | -19.770 | 0.0391 | 1.2000 | H |
| ATOM | 93 | CG | PRO | 6 | 33.410 | -6.305 | -20.681 | 0.0189 | 1.7000 | C |
| ATOM | 94 | HG2 | PRO | 6 | 33.348 | -6.772 | -21.665 | 0.0213 | 1.2000 | H |
| ATOM | 95 | HG3 | PRO | 6 | 33.929 | -5.347 | -20.750 | 0.0213 | 1.2000 | H |
| ATOM | 96 | CB | PRO | 6 | 32.048 | -6.093 | -20.082 | -0.0070 | 1.7000 | C |
| ATOM | 97 | HB2 | PRO | 6 | 31.422 | -6.958 | -20.308 | 0.0253 | 1.2000 | H |
| ATOM | 98 | HB3 | PRO | 6 | 31.580 | -5.184 | -20.463 | 0.0253 | 1.2000 | H |
| ATOM | 99 | CA | PRO | 6 | 32.297 | -6.002 | -18.573 | -0.0266 | 1.7000 | C |
| ATOM | 100 | HA | PRO | 6 | 32.462 | -4.967 | -18.284 | 0.0641 | 1.2000 | H |
| ATOM | 101 | C | PRO | 6 | 31.122 | -6.587 | -17.784 | 0.5896 | 1.7000 | C |

| | | | | | | | | | | |
|---|---|---|---|---|---|---|---|---|---|---|
| ATOM | 102 | O | PRO | 6 | 30.156 | -5.877 | -17.487 | -0.5748 | 1.5000 | O |
| ATOM | 103 | N | TYR | 7 | 31.165 | -7.884 | -17.456 | -0.4157 | 1.5500 | N |
| ATOM | 104 | H | TYR | 7 | 31.963 | -8.456 | -17.694 | 0.2719 | 1.3000 | H |
| ATOM | 105 | CA | TYR | 7 | 30.072 | -8.467 | -16.700 | -0.0014 | 1.7000 | C |
| ATOM | 106 | HA | TYR | 7 | 29.128 | -8.082 | -17.089 | 0.0876 | 1.2000 | H |
| ATOM | 107 | CB | TYR | 7 | 30.026 | -9.982 | -16.798 | -0.0152 | 1.7000 | C |
| ATOM | 108 | HB2 | TYR | 7 | 30.049 | -10.272 | -17.850 | 0.0295 | 1.2000 | H |
| ATOM | 109 | HB3 | TYR | 7 | 30.900 | -10.412 | -16.315 | 0.0295 | 1.2000 | H |
| ATOM | 110 | CG | TYR | 7 | 28.769 | -10.533 | -16.174 | -0.0011 | 1.7000 | C |
| ATOM | 111 | CD1 | TYR | 7 | 27.570 | -10.416 | -16.871 | -0.1906 | 1.7000 | C |
| ATOM | 112 | HD1 | TYR | 7 | 27.552 | -9.947 | -17.845 | 0.1699 | 1.2000 | H |
| ATOM | 113 | CE1 | TYR | 7 | 26.400 | -10.897 | -16.323 | -0.2341 | 1.7000 | C |
| ATOM | 114 | HE1 | TYR | 7 | 25.470 | -10.809 | -16.866 | 0.1656 | 1.2000 | H |
| ATOM | 115 | CZ | TYR | 7 | 26.430 | -11.497 | -15.076 | 0.3226 | 1.7000 | C |
| ATOM | 116 | OH | TYR | 7 | 25.265 | -11.978 | -14.520 | -0.5579 | 1.5000 | O |
| ATOM | 117 | HH | TYR | 7 | 24.537 | -11.992 | -15.144 | 0.3992 | 1.2000 | H |
| ATOM | 118 | CE2 | TYR | 7 | 27.616 | -11.609 | -14.393 | -0.2341 | 1.7000 | C |
| ATOM | 119 | HE2 | TYR | 7 | 27.633 | -12.090 | -13.431 | 0.1656 | 1.2000 | H |
| ATOM | 120 | CD2 | TYR | 7 | 28.782 | -11.130 | -14.932 | -0.1906 | 1.7000 | C |
| ATOM | 121 | HD2 | TYR | 7 | 29.702 | -11.224 | -14.383 | 0.1699 | 1.2000 | H |
| ATOM | 122 | C | TYR | 7 | 30.239 | -7.985 | -15.273 | 0.5973 | 1.7000 | C |
| ATOM | 123 | O | TYR | 7 | 29.272 | -7.583 | -14.620 | -0.5679 | 1.5000 | O |
| ATOM | 124 | N | CYX | 8 | 31.491 | -8.023 | -14.798 | -0.4157 | 1.5500 | N |
| ATOM | 125 | H | CYX | 8 | 32.245 | -8.368 | -15.376 | 0.2719 | 1.3000 | H |
| ATOM | 126 | CA | CYX | 8 | 31.797 | -7.511 | -13.470 | 0.0429 | 1.7000 | C |
| ATOM | 127 | HA | CYX | 8 | 31.226 | -8.076 | -12.735 | 0.0766 | 1.2000 | H |
| ATOM | 128 | CB | CYX | 8 | 33.283 | -7.643 | -13.137 | -0.0790 | 1.7000 | C |
| ATOM | 129 | HB2 | CYX | 8 | 33.568 | -8.694 | -13.201 | 0.0910 | 1.2000 | H |
| ATOM | 130 | HB3 | CYX | 8 | 33.872 | -7.092 | -13.864 | 0.0910 | 1.2000 | H |
| ATOM | 131 | SG | CYX | 8 | 33.703 | -7.016 | -11.485 | -0.1081 | 1.8000 | S |
| ATOM | 132 | C | CYX | 8 | 31.399 | -6.049 | -13.381 | 0.5973 | 1.7000 | C |
| ATOM | 133 | O | CYX | 8 | 30.749 | -5.615 | -12.428 | -0.5679 | 1.5000 | O |
| ATOM | 134 | N | TRP | 9 | 31.770 | -5.264 | -14.394 | -0.4157 | 1.5500 | N |
| ATOM | 135 | H | TRP | 9 | 32.298 | -5.634 | -15.172 | 0.2719 | 1.3000 | H |

| | | | | | | | | | | |
|---|---|---|---|---|---|---|---|---|---|---|
| ATOM | 136 | CA | TRP | 9 | 31.407 | -3.859 | -14.376 | -0.0275 | 1.7000 | C |
| ATOM | 137 | HA | TRP | 9 | 31.870 | -3.373 | -13.518 | 0.1123 | 1.2000 | H |
| ATOM | 138 | CB | TRP | 9 | 31.850 | -3.167 | -15.659 | -0.0050 | 1.7000 | C |
| ATOM | 139 | HB2 | TRP | 9 | 31.518 | -3.746 | -16.520 | 0.0339 | 1.2000 | H |
| ATOM | 140 | HB3 | TRP | 9 | 31.365 | -2.191 | -15.712 | 0.0339 | 1.2000 | H |
| ATOM | 141 | CG | TRP | 9 | 33.317 | -2.930 | -15.764 | -0.1415 | 1.7000 | C |
| ATOM | 142 | CD1 | TRP | 9 | 34.218 | -2.903 | -14.746 | -0.1638 | 1.7000 | C |
| ATOM | 143 | HD1 | TRP | 9 | 33.991 | -3.056 | -13.703 | 0.2062 | 1.2000 | H |
| ATOM | 144 | NE1 | TRP | 9 | 35.471 | -2.630 | -15.236 | -0.3418 | 1.5500 | N |
| ATOM | 145 | HE1 | TRP | 9 | 36.305 | -2.562 | -14.672 | 0.3412 | 1.3000 | H |
| ATOM | 146 | CE2 | TRP | 9 | 35.391 | -2.481 | -16.597 | 0.1380 | 1.7000 | C |
| ATOM | 147 | CZ2 | TRP | 9 | 36.378 | -2.202 | -17.524 | -0.2601 | 1.7000 | C |
| ATOM | 148 | HZ2 | TRP | 9 | 37.402 | -2.044 | -17.222 | 0.1572 | 1.2000 | H |
| ATOM | 149 | CH2 | TRP | 9 | 36.006 | -2.124 | -18.860 | -0.1134 | 1.7000 | C |
| ATOM | 150 | HH2 | TRP | 9 | 36.759 | -1.919 | -19.606 | 0.1417 | 1.2000 | H |
| ATOM | 151 | CZ3 | TRP | 9 | 34.711 | -2.305 | -19.251 | -0.1972 | 1.7000 | C |
| ATOM | 152 | HZ3 | TRP | 9 | 34.454 | -2.224 | -20.297 | 0.1447 | 1.2000 | H |
| ATOM | 153 | CE3 | TRP | 9 | 33.721 | -2.570 | -18.325 | -0.2387 | 1.7000 | C |
| ATOM | 154 | HE3 | TRP | 9 | 32.697 | -2.703 | -18.641 | 0.1700 | 1.2000 | H |
| ATOM | 155 | CD2 | TRP | 9 | 34.067 | -2.669 | -16.973 | 0.1243 | 1.7000 | C |
| ATOM | 156 | C | TRP | 9 | 29.896 | -3.702 | -14.236 | 0.5973 | 1.7000 | C |
| ATOM | 157 | O | TRP | 9 | 29.430 | -2.899 | -13.420 | -0.5679 | 1.5000 | O |
| ATOM | 158 | N | LEU | 10 | 29.124 | -4.497 | -14.985 | -0.4157 | 1.5500 | N |
| ATOM | 159 | H | LEU | 10 | 29.535 | -5.145 | -15.643 | 0.2719 | 1.3000 | H |
| ATOM | 160 | CA | LEU | 10 | 27.678 | -4.434 | -14.869 | -0.0518 | 1.7000 | C |
| ATOM | 161 | HA | LEU | 10 | 27.380 | -3.412 | -15.106 | 0.0922 | 1.2000 | H |
| ATOM | 162 | CB | LEU | 10 | 26.978 | -5.376 | -15.844 | -0.1102 | 1.7000 | C |
| ATOM | 163 | HB2 | LEU | 10 | 27.203 | -5.053 | -16.862 | 0.0457 | 1.2000 | H |
| ATOM | 164 | HB3 | LEU | 10 | 27.380 | -6.380 | -15.724 | 0.0457 | 1.2000 | H |
| ATOM | 165 | CG | LEU | 10 | 25.446 | -5.447 | -15.654 | 0.3531 | 1.7000 | C |
| ATOM | 166 | HG | LEU | 10 | 25.194 | -5.783 | -14.649 | -0.0361 | 1.2000 | H |
| ATOM | 167 | CD1 | LEU | 10 | 24.834 | -4.066 | -15.881 | -0.4121 | 1.7000 | C |
| ATOM | 168 | HD11 | LEU | 10 | 23.748 | -4.138 | -15.820 | 0.1000 | 1.2000 | H |
| ATOM | 169 | HD12 | LEU | 10 | 25.163 | -3.354 | -15.127 | 0.1000 | 1.2000 | H |

| | | | | | | | | | | |
|---|---|---|---|---|---|---|---|---|---|---|
| ATOM | 170 | HD13 | LEU | 10 | 25.106 | -3.694 | -16.870 | 0.1000 | 1.2000 | H |
| ATOM | 171 | CD2 | LEU | 10 | 24.870 | -6.483 | -16.604 | -0.4121 | 1.7000 | C |
| ATOM | 172 | HD21 | LEU | 10 | 25.299 | -7.463 | -16.388 | 0.1000 | 1.2000 | H |
| ATOM | 173 | HD22 | LEU | 10 | 23.788 | -6.539 | -16.477 | 0.1000 | 1.2000 | H |
| ATOM | 174 | HD23 | LEU | 10 | 25.095 | -6.213 | -17.637 | 0.1000 | 1.2000 | H |
| ATOM | 175 | C | LEU | 10 | 27.172 | -4.765 | -13.485 | 0.5973 | 1.7000 | C |
| ATOM | 176 | O | LEU | 10 | 26.362 | -4.015 | -12.940 | -0.5679 | 1.5000 | O |
| ATOM | 177 | N | CYX | 11 | 27.611 | -5.881 | -12.894 | -0.4157 | 1.5500 | N |
| ATOM | 178 | H | CYX | 11 | 28.271 | -6.503 | -13.341 | 0.2719 | 1.3000 | H |
| ATOM | 179 | CA | CYX | 11 | 27.032 | -6.205 | -11.596 | 0.0429 | 1.7000 | C |
| ATOM | 180 | HA | CYX | 11 | 25.947 | -6.236 | -11.690 | 0.0766 | 1.2000 | H |
| ATOM | 181 | CB | CYX | 11 | 27.504 | -7.575 | -11.114 | -0.0790 | 1.7000 | C |
| ATOM | 182 | HB2 | CYX | 11 | 26.907 | -7.870 | -10.254 | 0.0910 | 1.2000 | H |
| ATOM | 183 | HB3 | CYX | 11 | 27.334 | -8.300 | -11.911 | 0.0910 | 1.2000 | H |
| ATOM | 184 | SG | CYX | 11 | 29.237 | -7.633 | -10.646 | -0.1081 | 1.8000 | S |
| ATOM | 185 | C | CYX | 11 | 27.410 | -5.160 | -10.557 | 0.5973 | 1.7000 | C |
| ATOM | 186 | O | CYX | 11 | 26.617 | -4.841 | -9.668 | -0.5679 | 1.5000 | O |
| ATOM | 187 | N | ARG | 12 | 28.567 | -4.526 | -10.732 | -0.3479 | 1.5500 | N |
| ATOM | 188 | H | ARG | 12 | 29.185 | -4.781 | -11.491 | 0.2747 | 1.3000 | H |
| ATOM | 189 | CA | ARG | 12 | 28.971 | -3.506 | -9.796 | -0.2637 | 1.7000 | C |
| ATOM | 190 | HA | ARG | 12 | 28.802 | -3.869 | -8.781 | 0.1560 | 1.2000 | H |
| ATOM | 191 | CB | ARG | 12 | 30.446 | -3.209 | -9.947 | -0.0007 | 1.7000 | C |
| ATOM | 192 | HB2 | ARG | 12 | 30.667 | -2.994 | -10.994 | 0.0327 | 1.2000 | H |
| ATOM | 193 | HB3 | ARG | 12 | 30.678 | -2.318 | -9.362 | 0.0327 | 1.2000 | H |
| ATOM | 194 | CG | ARG | 12 | 31.329 | -4.354 | -9.464 | 0.0390 | 1.7000 | C |
| ATOM | 195 | HG2 | ARG | 12 | 31.172 | -4.505 | -8.397 | 0.0285 | 1.2000 | H |
| ATOM | 196 | HG3 | ARG | 12 | 31.044 | -5.277 | -9.963 | 0.0285 | 1.2000 | H |
| ATOM | 197 | CD | ARG | 12 | 32.752 | -4.130 | -9.716 | 0.0486 | 1.7000 | C |
| ATOM | 198 | HD2 | ARG | 12 | 33.293 | -5.032 | -9.429 | 0.0687 | 1.2000 | H |
| ATOM | 199 | HD3 | ARG | 12 | 32.894 | -3.966 | -10.786 | 0.0687 | 1.2000 | H |
| ATOM | 200 | NE | ARG | 12 | 33.299 | -3.014 | -8.967 | -0.5295 | 1.5500 | N |
| ATOM | 201 | HE | ARG | 12 | 32.721 | -2.606 | -8.247 | 0.3456 | 1.3000 | H |
| ATOM | 202 | CZ | ARG | 12 | 34.533 | -2.522 | -9.156 | 0.8076 | 1.7000 | C |
| ATOM | 203 | NH1 | ARG | 12 | 35.313 | -3.062 | -10.068 | -0.8627 | 1.5500 | N |

| | | | | | | | | | | |
|---|---|---|---|---|---|---|---|---|---|---|
| ATOM | 204 | HH11 | ARG | 12 | 36.242 | -2.699 | -10.222 | 0.4478 | 1.3000 | H |
| ATOM | 205 | HH12 | ARG | 12 | 34.983 | -3.845 | -10.612 | 0.4478 | 1.3000 | H |
| ATOM | 206 | NH2 | ARG | 12 | 34.965 | -1.505 | -8.431 | -0.8627 | 1.5500 | N |
| ATOM | 207 | HH21 | ARG | 12 | 35.876 | -1.111 | -8.605 | 0.4478 | 1.3000 | H |
| ATOM | 208 | HH22 | ARG | 12 | 34.390 | -1.145 | -7.685 | 0.4478 | 1.3000 | H |
| ATOM | 209 | C | ARG | 12 | 28.104 | -2.275 | -9.995 | 0.7341 | 1.7000 | C |
| ATOM | 210 | O | ARG | 12 | 27.714 | -1.625 | -9.024 | -0.5894 | 1.5000 | O |
| ATOM | 211 | N | ALA | 13 | 27.775 | -1.960 | -11.253 | -0.4157 | 1.5500 | N |
| ATOM | 212 | H | ALA | 13 | 28.107 | -2.526 | -12.021 | 0.2719 | 1.3000 | H |
| ATOM | 213 | CA | ALA | 13 | 26.904 | -0.833 | -11.546 | 0.0337 | 1.7000 | C |
| ATOM | 214 | HA | ALA | 13 | 27.345 | 0.073 | -11.127 | 0.0823 | 1.2000 | H |
| ATOM | 215 | CB | ALA | 13 | 26.750 | -0.648 | -13.044 | -0.1825 | 1.7000 | C |
| ATOM | 216 | HB1 | ALA | 13 | 26.137 | 0.232 | -13.240 | 0.0603 | 1.2000 | H |
| ATOM | 217 | HB2 | ALA | 13 | 27.730 | -0.502 | -13.500 | 0.0603 | 1.2000 | H |
| ATOM | 218 | HB3 | ALA | 13 | 26.273 | -1.516 | -13.496 | 0.0603 | 1.2000 | H |
| ATOM | 219 | C | ALA | 13 | 25.535 | -1.049 | -10.920 | 0.5973 | 1.7000 | C |
| ATOM | 220 | O | ALA | 13 | 24.947 | -0.125 | -10.352 | -0.5679 | 1.5000 | O |
| ATOM | 221 | N | LEU | 14 | 25.043 | -2.292 | -10.984 | -0.4157 | 1.5500 | N |
| ATOM | 222 | H | LEU | 14 | 25.559 | -3.026 | -11.449 | 0.2719 | 1.3000 | H |
| ATOM | 223 | CA | LEU | 14 | 23.734 | -2.595 | -10.431 | -0.0518 | 1.7000 | C |
| ATOM | 224 | HA | LEU | 14 | 23.000 | -1.899 | -10.835 | 0.0922 | 1.2000 | H |
| ATOM | 225 | CB | LEU | 14 | 23.342 | -4.030 | -10.791 | -0.1102 | 1.7000 | C |
| ATOM | 226 | HB2 | LEU | 14 | 24.125 | -4.706 | -10.448 | 0.0457 | 1.2000 | H |
| ATOM | 227 | HB3 | LEU | 14 | 22.430 | -4.286 | -10.249 | 0.0457 | 1.2000 | H |
| ATOM | 228 | CG | LEU | 14 | 23.103 | -4.265 | -12.307 | 0.3531 | 1.7000 | C |
| ATOM | 229 | HG | LEU | 14 | 23.970 | -3.952 | -12.875 | -0.0361 | 1.2000 | H |
| ATOM | 230 | CD1 | LEU | 14 | 22.880 | -5.746 | -12.572 | -0.4121 | 1.7000 | C |
| ATOM | 231 | HD11 | LEU | 14 | 22.740 | -5.911 | -13.641 | 0.1000 | 1.2000 | H |
| ATOM | 232 | HD12 | LEU | 14 | 23.745 | -6.320 | -12.244 | 0.1000 | 1.2000 | H |
| ATOM | 233 | HD13 | LEU | 14 | 21.994 | -6.093 | -12.039 | 0.1000 | 1.2000 | H |
| ATOM | 234 | CD2 | LEU | 14 | 21.943 | -3.433 | -12.772 | -0.4121 | 1.7000 | C |
| ATOM | 235 | HD21 | LEU | 14 | 21.702 | -3.692 | -13.804 | 0.1000 | 1.2000 | H |
| ATOM | 236 | HD22 | LEU | 14 | 21.067 | -3.619 | -12.150 | 0.1000 | 1.2000 | H |
| ATOM | 237 | HD23 | LEU | 14 | 22.195 | -2.375 | -12.749 | 0.1000 | 1.2000 | H |

| | | | | | | | | | | |
|---|---|---|---|---|---|---|---|---|---|---|
| ATOM | 238 | C | LEU | 14 | 23.779 | -2.380 | -8.923 | 0.5973 | 1.7000 | C |
| ATOM | 239 | O | LEU | 14 | 22.864 | -1.788 | -8.342 | -0.5679 | 1.5000 | O |
| ATOM | 240 | N | ILE | 15 | 24.888 | -2.783 | -8.300 | -0.4157 | 1.5500 | N |
| ATOM | 241 | H | ILE | 15 | 25.622 | -3.247 | -8.817 | 0.2719 | 1.3000 | H |
| ATOM | 242 | CA | ILE | 15 | 25.061 | -2.563 | -6.877 | -0.0597 | 1.7000 | C |
| ATOM | 243 | HA | ILE | 15 | 24.206 | -2.998 | -6.357 | 0.0869 | 1.2000 | H |
| ATOM | 244 | CB | ILE | 15 | 26.324 | -3.232 | -6.343 | 0.1303 | 1.7000 | C |
| ATOM | 245 | HB | ILE | 15 | 27.152 | -2.993 | -7.005 | 0.0187 | 1.2000 | H |
| ATOM | 246 | CG2 | ILE | 15 | 26.641 | -2.642 | -4.993 | -0.3204 | 1.7000 | C |
| ATOM | 247 | HG21 | ILE | 15 | 27.470 | -3.186 | -4.554 | 0.0882 | 1.2000 | H |
| ATOM | 248 | HG22 | ILE | 15 | 26.987 | -1.611 | -5.063 | 0.0882 | 1.2000 | H |
| ATOM | 249 | HG23 | ILE | 15 | 25.780 | -2.713 | -4.328 | 0.0882 | 1.2000 | H |
| ATOM | 250 | CG1 | ILE | 15 | 26.137 | -4.789 | -6.362 | -0.0430 | 1.7000 | C |
| ATOM | 251 | HG12 | ILE | 15 | 25.414 | -5.062 | -5.592 | 0.0236 | 1.2000 | H |
| ATOM | 252 | HG13 | ILE | 15 | 25.717 | -5.099 | -7.316 | 0.0236 | 1.2000 | H |
| ATOM | 253 | CD1 | ILE | 15 | 27.407 | -5.601 | -6.137 | -0.0660 | 1.7000 | C |
| ATOM | 254 | HD11 | ILE | 15 | 27.183 | -6.663 | -6.239 | 0.0186 | 1.2000 | H |
| ATOM | 255 | HD12 | ILE | 15 | 28.151 | -5.318 | -6.880 | 0.0186 | 1.2000 | H |
| ATOM | 256 | HD13 | ILE | 15 | 27.805 | -5.421 | -5.142 | 0.0186 | 1.2000 | H |
| ATOM | 257 | C | ILE | 15 | 25.091 | -1.090 | -6.542 | 0.5973 | 1.7000 | C |
| ATOM | 258 | O | ILE | 15 | 24.454 | -0.667 | -5.579 | -0.5679 | 1.5000 | O |
| ATOM | 259 | N | LYS | 16 | 25.822 | -0.295 | -7.319 | -0.3479 | 1.5500 | N |
| ATOM | 260 | H | LYS | 16 | 26.329 | -0.670 | -8.107 | 0.2747 | 1.3000 | H |
| ATOM | 261 | CA | LYS | 16 | 25.889 | 1.128 | -7.030 | -0.2400 | 1.7000 | C |
| ATOM | 262 | HA | LYS | 16 | 26.258 | 1.266 | -6.012 | 0.1426 | 1.2000 | H |
| ATOM | 263 | CB | LYS | 16 | 26.846 | 1.832 | -7.989 | -0.0094 | 1.7000 | C |
| ATOM | 264 | HB2 | LYS | 16 | 26.577 | 1.575 | -9.015 | 0.0362 | 1.2000 | H |
| ATOM | 265 | HB3 | LYS | 16 | 26.714 | 2.909 | -7.871 | 0.0362 | 1.2000 | H |
| ATOM | 266 | CG | LYS | 16 | 28.320 | 1.517 | -7.761 | 0.0187 | 1.7000 | C |
| ATOM | 267 | HG2 | LYS | 16 | 28.600 | 1.836 | -6.756 | 0.0103 | 1.2000 | H |
| ATOM | 268 | HG3 | LYS | 16 | 28.493 | 0.446 | -7.839 | 0.0103 | 1.2000 | H |
| ATOM | 269 | CD | LYS | 16 | 29.190 | 2.243 | -8.776 | -0.0479 | 1.7000 | C |
| ATOM | 270 | HD2 | LYS | 16 | 28.903 | 1.928 | -9.781 | 0.0621 | 1.2000 | H |
| ATOM | 271 | HD3 | LYS | 16 | 29.027 | 3.318 | -8.683 | 0.0621 | 1.2000 | H |

| | | | | | | | | | | |
|---|---|---|---|---|---|---|---|---|---|---|
| ATOM | 272 | CE | LYS | 16 | 30.663 | 1.945 | -8.567 | -0.0143 | 1.7000 | C |
| ATOM | 273 | HE2 | LYS | 16 | 30.954 | 2.261 | -7.563 | 0.1135 | 1.2000 | H |
| ATOM | 274 | HE3 | LYS | 16 | 30.815 | 0.868 | -8.653 | 0.1135 | 1.2000 | H |
| ATOM | 275 | NZ | LYS | 16 | 31.514 | 2.644 | -9.576 | -0.3854 | 1.5500 | N |
| ATOM | 276 | HZ1 | LYS | 16 | 31.467 | 3.643 | -9.425 | 0.3400 | 1.3000 | H |
| ATOM | 277 | HZ2 | LYS | 16 | 32.474 | 2.344 | -9.488 | 0.3400 | 1.3000 | H |
| ATOM | 278 | HZ3 | LYS | 16 | 31.183 | 2.436 | -10.508 | 0.3400 | 1.3000 | H |
| ATOM | 279 | C | LYS | 16 | 24.511 | 1.772 | -7.115 | 0.7341 | 1.7000 | C |
| ATOM | 280 | O | LYS | 16 | 24.170 | 2.627 | -6.293 | -0.5894 | 1.5000 | O |
| ATOM | 281 | N | ARG | 17 | 23.687 | 1.339 | -8.075 | -0.3479 | 1.5500 | N |
| ATOM | 282 | H | ARG | 17 | 23.988 | 0.623 | -8.722 | 0.2747 | 1.3000 | H |
| ATOM | 283 | CA | ARG | 17 | 22.345 | 1.897 | -8.198 | -0.2637 | 1.7000 | C |
| ATOM | 284 | HA | ARG | 17 | 22.403 | 2.984 | -8.281 | 0.1560 | 1.2000 | H |
| ATOM | 285 | CB | ARG | 17 | 21.662 | 1.336 | -9.434 | -0.0007 | 1.7000 | C |
| ATOM | 286 | HB2 | ARG | 17 | 21.700 | 0.247 | -9.402 | 0.0327 | 1.2000 | H |
| ATOM | 287 | HB3 | ARG | 17 | 20.611 | 1.630 | -9.412 | 0.0327 | 1.2000 | H |
| ATOM | 288 | CG | ARG | 17 | 22.257 | 1.820 | -10.754 | 0.0390 | 1.7000 | C |
| ATOM | 289 | HG2 | ARG | 17 | 22.070 | 2.890 | -10.854 | 0.0285 | 1.2000 | H |
| ATOM | 290 | HG3 | ARG | 17 | 23.333 | 1.671 | -10.756 | 0.0285 | 1.2000 | H |
| ATOM | 291 | CD | ARG | 17 | 21.664 | 1.114 | -11.917 | 0.0486 | 1.7000 | C |
| ATOM | 292 | HD2 | ARG | 17 | 21.759 | 0.039 | -11.754 | 0.0687 | 1.2000 | H |
| ATOM | 293 | HD3 | ARG | 17 | 20.605 | 1.368 | -11.988 | 0.0687 | 1.2000 | H |
| ATOM | 294 | NE | ARG | 17 | 22.329 | 1.460 | -13.162 | -0.5295 | 1.5500 | N |
| ATOM | 295 | HE | ARG | 17 | 22.997 | 2.215 | -13.133 | 0.3456 | 1.3000 | H |
| ATOM | 296 | CZ | ARG | 17 | 22.122 | 0.832 | -14.333 | 0.8076 | 1.7000 | C |
| ATOM | 297 | NH1 | ARG | 17 | 21.248 | -0.147 | -14.403 | -0.8627 | 1.5500 | N |
| ATOM | 298 | HH11 | ARG | 17 | 20.744 | -0.431 | -13.577 | 0.4478 | 1.3000 | H |
| ATOM | 299 | HH12 | ARG | 17 | 21.067 | -0.600 | -15.286 | 0.4478 | 1.3000 | H |
| ATOM | 300 | NH2 | ARG | 17 | 22.793 | 1.197 | -15.411 | -0.8627 | 1.5500 | N |
| ATOM | 301 | HH21 | ARG | 17 | 22.629 | 0.731 | -16.290 | 0.4478 | 1.3000 | H |
| ATOM | 302 | HH22 | ARG | 17 | 23.454 | 1.957 | -15.361 | 0.4478 | 1.3000 | H |
| ATOM | 303 | C | ARG | 17 | 21.537 | 1.550 | -6.948 | 0.7341 | 1.7000 | C |
| ATOM | 304 | O | ARG | 17 | 20.844 | 2.400 | -6.380 | -0.5894 | 1.5000 | O |
| ATOM | 305 | N | ILE | 18 | 21.693 | 0.318 | -6.464 | -0.4157 | 1.5500 | N |

| | | | | | | | | | | |
|---|---|---|---|---|---|---|---|---|---|---|
| ATOM | 306 | H | ILE | 18 | 22.292 | -0.346 | -6.935 | 0.2719 | 1.3000 | H |
| ATOM | 307 | CA | ILE | 18 | 20.987 | -0.092 | -5.265 | -0.0597 | 1.7000 | C |
| ATOM | 308 | HA | ILE | 18 | 19.920 | 0.086 | -5.404 | 0.0869 | 1.2000 | H |
| ATOM | 309 | CB | ILE | 18 | 21.197 | -1.591 | -4.990 | 0.1303 | 1.7000 | C |
| ATOM | 310 | HB | ILE | 18 | 22.262 | -1.817 | -5.034 | 0.0187 | 1.2000 | H |
| ATOM | 311 | CG2 | ILE | 18 | 20.706 | -1.951 | -3.591 | -0.3204 | 1.7000 | C |
| ATOM | 312 | HG21 | ILE | 18 | 20.745 | -3.025 | -3.426 | 0.0882 | 1.2000 | H |
| ATOM | 313 | HG22 | ILE | 18 | 21.347 | -1.499 | -2.836 | 0.0882 | 1.2000 | H |
| ATOM | 314 | HG23 | ILE | 18 | 19.679 | -1.610 | -3.450 | 0.0882 | 1.2000 | H |
| ATOM | 315 | CG1 | ILE | 18 | 20.487 | -2.410 | -6.081 | -0.0430 | 1.7000 | C |
| ATOM | 316 | HG12 | ILE | 18 | 19.409 | -2.335 | -5.928 | 0.0236 | 1.2000 | H |
| ATOM | 317 | HG13 | ILE | 18 | 20.698 | -2.003 | -7.068 | 0.0236 | 1.2000 | H |
| ATOM | 318 | CD1 | ILE | 18 | 20.869 | -3.872 | -6.095 | -0.0660 | 1.7000 | C |
| ATOM | 319 | HD11 | ILE | 18 | 20.282 | -4.385 | -6.858 | 0.0186 | 1.2000 | H |
| ATOM | 320 | HD12 | ILE | 18 | 21.926 | -3.974 | -6.341 | 0.0186 | 1.2000 | H |
| ATOM | 321 | HD13 | ILE | 18 | 20.672 | -4.348 | -5.137 | 0.0186 | 1.2000 | H |
| ATOM | 322 | C | ILE | 18 | 21.446 | 0.726 | -4.075 | 0.5973 | 1.7000 | C |
| ATOM | 323 | O | ILE | 18 | 20.620 | 1.268 | -3.339 | -0.5679 | 1.5000 | O |
| ATOM | 324 | N | GLN | 19 | 22.756 | 0.869 | -3.911 | -0.4157 | 1.5500 | N |
| ATOM | 325 | H | GLN | 19 | 23.397 | 0.437 | -4.559 | 0.2719 | 1.3000 | H |
| ATOM | 326 | CA | GLN | 19 | 23.291 | 1.608 | -2.785 | -0.0031 | 1.7000 | C |
| ATOM | 327 | HA | GLN | 19 | 22.917 | 1.164 | -1.863 | 0.0850 | 1.2000 | H |
| ATOM | 328 | CB | GLN | 19 | 24.806 | 1.533 | -2.776 | -0.0036 | 1.7000 | C |
| ATOM | 329 | HB2 | GLN | 19 | 25.171 | 1.850 | -3.754 | 0.0171 | 1.2000 | H |
| ATOM | 330 | HB3 | GLN | 19 | 25.184 | 2.239 | -2.034 | 0.0171 | 1.2000 | H |
| ATOM | 331 | CG | GLN | 19 | 25.360 | 0.174 | -2.443 | -0.0645 | 1.7000 | C |
| ATOM | 332 | HG2 | GLN | 19 | 25.036 | -0.111 | -1.442 | 0.0352 | 1.2000 | H |
| ATOM | 333 | HG3 | GLN | 19 | 24.975 | -0.576 | -3.129 | 0.0352 | 1.2000 | H |
| ATOM | 334 | CD | GLN | 19 | 26.842 | 0.178 | -2.532 | 0.6951 | 1.7000 | C |
| ATOM | 335 | OE1 | GLN | 19 | 27.433 | 0.674 | -3.499 | -0.6086 | 1.5000 | O |
| ATOM | 336 | NE2 | GLN | 19 | 27.465 | -0.339 | -1.506 | -0.9407 | 1.5500 | N |
| ATOM | 337 | HE21 | GLN | 19 | 26.947 | -0.706 | -0.721 | 0.4251 | 1.3000 | H |
| ATOM | 338 | HE22 | GLN | 19 | 28.474 | -0.326 | -1.484 | 0.4251 | 1.3000 | H |
| ATOM | 339 | C | GLN | 19 | 22.851 | 3.056 | -2.816 | 0.5973 | 1.7000 | C |

| | | | | | | | | | | |
|---|---|---|---|---|---|---|---|---|---|---|
| ATOM | 340 | O | GLN | 19 | 22.557 | 3.645 | -1.782 | -0.5679 | 1.5000 | O |
| ATOM | 341 | N | ALA | 20 | 22.779 | 3.640 | -4.008 | -0.4157 | 1.5500 | N |
| ATOM | 342 | H | ALA | 20 | 23.025 | 3.124 | -4.840 | 0.2719 | 1.3000 | H |
| ATOM | 343 | CA | ALA | 20 | 22.327 | 5.015 | -4.148 | 0.0337 | 1.7000 | C |
| ATOM | 344 | HA | ALA | 20 | 22.983 | 5.662 | -3.564 | 0.0823 | 1.2000 | H |
| ATOM | 345 | CB | ALA | 20 | 22.397 | 5.441 | -5.603 | -0.1825 | 1.7000 | C |
| ATOM | 346 | HB1 | ALA | 20 | 22.095 | 6.485 | -5.692 | 0.0603 | 1.2000 | H |
| ATOM | 347 | HB2 | ALA | 20 | 23.420 | 5.340 | -5.967 | 0.0603 | 1.2000 | H |
| ATOM | 348 | HB3 | ALA | 20 | 21.736 | 4.829 | -6.214 | 0.0603 | 1.2000 | H |
| ATOM | 349 | C | ALA | 20 | 20.899 | 5.171 | -3.630 | 0.5973 | 1.7000 | C |
| ATOM | 350 | O | ALA | 20 | 20.552 | 6.199 | -3.044 | -0.5679 | 1.5000 | O |
| ATOM | 351 | N | MET | 21 | 20.075 | 4.143 | -3.846 | -0.4157 | 1.5500 | N |
| ATOM | 352 | H | MET | 21 | 20.424 | 3.325 | -4.326 | 0.2719 | 1.3000 | H |
| ATOM | 353 | CA | MET | 21 | 18.680 | 4.140 | -3.428 | -0.0237 | 1.7000 | C |
| ATOM | 354 | HA | MET | 21 | 18.283 | 5.152 | -3.525 | 0.0880 | 1.2000 | H |
| ATOM | 355 | CB | MET | 21 | 17.893 | 3.244 | -4.377 | 0.0342 | 1.7000 | C |
| ATOM | 356 | HB2 | MET | 21 | 18.353 | 2.257 | -4.409 | 0.0241 | 1.2000 | H |
| ATOM | 357 | HB3 | MET | 21 | 16.884 | 3.128 | -3.979 | 0.0241 | 1.2000 | H |
| ATOM | 358 | CG | MET | 21 | 17.765 | 3.797 | -5.797 | 0.0018 | 1.7000 | C |
| ATOM | 359 | HG2 | MET | 21 | 17.349 | 4.803 | -5.736 | 0.0440 | 1.2000 | H |
| ATOM | 360 | HG3 | MET | 21 | 18.747 | 3.864 | -6.263 | 0.0440 | 1.2000 | H |
| ATOM | 361 | SD | MET | 21 | 16.676 | 2.810 | -6.837 | -0.2737 | 1.8000 | S |
| ATOM | 362 | CE | MET | 21 | 17.725 | 1.388 | -7.143 | -0.0536 | 1.7000 | C |
| ATOM | 363 | HE1 | MET | 21 | 17.213 | 0.703 | -7.819 | 0.0684 | 1.2000 | H |
| ATOM | 364 | HE2 | MET | 21 | 18.656 | 1.714 | -7.603 | 0.0684 | 1.2000 | H |
| ATOM | 365 | HE3 | MET | 21 | 17.935 | 0.874 | -6.206 | 0.0684 | 1.2000 | H |
| ATOM | 366 | C | MET | 21 | 18.445 | 3.698 | -1.975 | 0.5973 | 1.7000 | C |
| ATOM | 367 | O | MET | 21 | 17.452 | 4.111 | -1.361 | -0.5679 | 1.5000 | O |
| ATOM | 368 | N | ILE | 22 | 19.359 | 2.909 | -1.394 | -0.4157 | 1.5500 | N |
| ATOM | 369 | H | ILE | 22 | 20.174 | 2.601 | -1.906 | 0.2719 | 1.3000 | H |
| ATOM | 370 | CA | ILE | 22 | 19.122 | 2.409 | -0.038 | -0.0597 | 1.7000 | C |
| ATOM | 371 | HA | ILE | 22 | 18.223 | 1.794 | -0.103 | 0.0869 | 1.2000 | H |
| ATOM | 372 | CB | ILE | 22 | 20.238 | 1.478 | 0.505 | 0.1303 | 1.7000 | C |
| ATOM | 373 | HB | ILE | 22 | 21.205 | 1.938 | 0.331 | 0.0187 | 1.2000 | H |

| | | | | | | | | | | |
|---|---|---|---|---|---|---|---|---|---|---|
| ATOM | 374 | CG2 | ILE | 22 | 20.064 | 1.326 | 1.973 | -0.3204 | 1.7000 | C |
| ATOM | 375 | HG21 | ILE | 22 | 20.770 | 0.599 | 2.364 | 0.0882 | 1.2000 | H |
| ATOM | 376 | HG22 | ILE | 22 | 20.284 | 2.241 | 2.519 | 0.0882 | 1.2000 | H |
| ATOM | 377 | HG23 | ILE | 22 | 19.054 | 0.996 | 2.204 | 0.0882 | 1.2000 | H |
| ATOM | 378 | CG1 | ILE | 22 | 20.253 | 0.142 | -0.213 | -0.0430 | 1.7000 | C |
| ATOM | 379 | HG12 | ILE | 22 | 19.347 | -0.417 | 0.001 | 0.0236 | 1.2000 | H |
| ATOM | 380 | HG13 | ILE | 22 | 20.259 | 0.303 | -1.277 | 0.0236 | 1.2000 | H |
| ATOM | 381 | CD1 | ILE | 22 | 21.478 | -0.696 | 0.125 | -0.0660 | 1.7000 | C |
| ATOM | 382 | HD11 | ILE | 22 | 21.502 | -1.572 | -0.520 | 0.0186 | 1.2000 | H |
| ATOM | 383 | HD12 | ILE | 22 | 22.390 | -0.122 | -0.032 | 0.0186 | 1.2000 | H |
| ATOM | 384 | HD13 | ILE | 22 | 21.433 | -1.043 | 1.155 | 0.0186 | 1.2000 | H |
| ATOM | 385 | C | ILE | 22 | 18.812 | 3.473 | 1.027 | 0.5973 | 1.7000 | C |
| ATOM | 386 | O | ILE | 22 | 17.865 | 3.275 | 1.780 | -0.5679 | 1.5000 | O |
| ATOM | 387 | N | PRO | 23 | 19.549 | 4.599 | 1.145 | -0.2548 | 1.5500 | N |
| ATOM | 388 | CD | PRO | 23 | 20.764 | 4.826 | 0.340 | 0.0192 | 1.7000 | C |
| ATOM | 389 | HD2 | PRO | 23 | 20.502 | 5.213 | -0.644 | 0.0391 | 1.2000 | H |
| ATOM | 390 | HD3 | PRO | 23 | 21.393 | 3.948 | 0.284 | 0.0391 | 1.2000 | H |
| ATOM | 391 | CG | PRO | 23 | 21.484 | 5.870 | 1.130 | 0.0189 | 1.7000 | C |
| ATOM | 392 | HG2 | PRO | 23 | 22.127 | 6.479 | 0.493 | 0.0213 | 1.2000 | H |
| ATOM | 393 | HG3 | PRO | 23 | 22.076 | 5.390 | 1.912 | 0.0213 | 1.2000 | H |
| ATOM | 394 | CB | PRO | 23 | 20.394 | 6.686 | 1.759 | -0.0070 | 1.7000 | C |
| ATOM | 395 | HB2 | PRO | 23 | 20.031 | 7.412 | 1.029 | 0.0253 | 1.2000 | H |
| ATOM | 396 | HB3 | PRO | 23 | 20.746 | 7.200 | 2.654 | 0.0253 | 1.2000 | H |
| ATOM | 397 | CA | PRO | 23 | 19.290 | 5.675 | 2.092 | -0.0266 | 1.7000 | C |
| ATOM | 398 | HA | PRO | 23 | 19.423 | 5.297 | 3.107 | 0.0641 | 1.2000 | H |
| ATOM | 399 | C | PRO | 23 | 17.910 | 6.336 | 1.952 | 0.5896 | 1.7000 | C |
| ATOM | 400 | O | PRO | 23 | 17.486 | 7.058 | 2.857 | -0.5748 | 1.5000 | O |
| ATOM | 401 | N | LYS | 24 | 17.250 | 6.172 | 0.797 | -0.3479 | 1.5500 | N |
| ATOM | 402 | H | LYS | 24 | 17.637 | 5.582 | 0.074 | 0.2747 | 1.3000 | H |
| ATOM | 403 | CA | LYS | 24 | 15.963 | 6.815 | 0.561 | -0.2400 | 1.7000 | C |
| ATOM | 404 | HA | LYS | 24 | 15.732 | 7.512 | 1.364 | 0.1426 | 1.2000 | H |
| ATOM | 405 | CB | LYS | 24 | 16.030 | 7.605 | -0.745 | -0.0094 | 1.7000 | C |
| ATOM | 406 | HB2 | LYS | 24 | 16.235 | 6.912 | -1.563 | 0.0362 | 1.2000 | H |
| ATOM | 407 | HB3 | LYS | 24 | 15.056 | 8.060 | -0.935 | 0.0362 | 1.2000 | H |

| | | | | | | | | | | |
|---|---|---|---|---|---|---|---|---|---|---|
| ATOM | 408 | CG | LYS | 24 | 17.097 | 8.711 | -0.774 | 0.0187 | 1.7000 | C |
| ATOM | 409 | HG2 | LYS | 24 | 18.086 | 8.286 | -0.599 | 0.0103 | 1.2000 | H |
| ATOM | 410 | HG3 | LYS | 24 | 17.095 | 9.159 | -1.769 | 0.0103 | 1.2000 | H |
| ATOM | 411 | CD | LYS | 24 | 16.817 | 9.806 | 0.255 | -0.0479 | 1.7000 | C |
| ATOM | 412 | HD2 | LYS | 24 | 15.816 | 10.208 | 0.091 | 0.0621 | 1.2000 | H |
| ATOM | 413 | HD3 | LYS | 24 | 16.870 | 9.387 | 1.260 | 0.0621 | 1.2000 | H |
| ATOM | 414 | CE | LYS | 24 | 17.836 | 10.942 | 0.155 | -0.0143 | 1.7000 | C |
| ATOM | 415 | HE2 | LYS | 24 | 18.837 | 10.531 | 0.301 | 0.1135 | 1.2000 | H |
| ATOM | 416 | HE3 | LYS | 24 | 17.781 | 11.377 | -0.844 | 0.1135 | 1.2000 | H |
| ATOM | 417 | NZ | LYS | 24 | 17.588 | 12.007 | 1.172 | -0.3854 | 1.5500 | N |
| ATOM | 418 | HZ1 | LYS | 24 | 18.265 | 12.749 | 1.063 | 0.3400 | 1.3000 | H |
| ATOM | 419 | HZ2 | LYS | 24 | 16.657 | 12.382 | 1.052 | 0.3400 | 1.3000 | H |
| ATOM | 420 | HZ3 | LYS | 24 | 17.672 | 11.615 | 2.100 | 0.3400 | 1.3000 | H |
| ATOM | 421 | C | LYS | 24 | 14.773 | 5.851 | 0.498 | 0.7341 | 1.7000 | C |
| ATOM | 422 | O | LYS | 24 | 13.636 | 6.252 | 0.763 | -0.5894 | 1.5000 | O |
| ATOM | 423 | N | GLY | 25 | 15.015 | 4.601 | 0.104 | -0.4157 | 1.5500 | N |
| ATOM | 424 | H | GLY | 25 | 15.958 | 4.303 | -0.099 | 0.2719 | 1.3000 | H |
| ATOM | 425 | CA | GLY | 25 | 13.917 | 3.663 | -0.103 | -0.0252 | 1.7000 | C |
| ATOM | 426 | HA2 | GLY | 25 | 13.091 | 4.166 | -0.608 | 0.0698 | 1.2000 | H |
| ATOM | 427 | HA3 | GLY | 25 | 14.268 | 2.870 | -0.763 | 0.0698 | 1.2000 | H |
| ATOM | 428 | C | GLY | 25 | 13.397 | 3.021 | 1.182 | 0.5973 | 1.7000 | C |
| ATOM | 429 | O | GLY | 25 | 13.863 | 3.315 | 2.282 | -0.5679 | 1.5000 | O |
| ATOM | 430 | N | ALA | 26 | 12.402 | 2.142 | 1.023 | -0.4157 | 1.5500 | N |
| ATOM | 431 | H | ALA | 26 | 12.057 | 1.948 | 0.094 | 0.2719 | 1.3000 | H |
| ATOM | 432 | CA | ALA | 26 | 11.748 | 1.461 | 2.143 | 0.0337 | 1.7000 | C |
| ATOM | 433 | HA | ALA | 26 | 12.411 | 1.469 | 3.010 | 0.0823 | 1.2000 | H |
| ATOM | 434 | CB | ALA | 26 | 10.485 | 2.184 | 2.532 | -0.1825 | 1.7000 | C |
| ATOM | 435 | HB1 | ALA | 26 | 10.040 | 1.681 | 3.386 | 0.0603 | 1.2000 | H |
| ATOM | 436 | HB2 | ALA | 26 | 10.727 | 3.207 | 2.823 | 0.0603 | 1.2000 | H |
| ATOM | 437 | HB3 | ALA | 26 | 9.780 | 2.214 | 1.701 | 0.0603 | 1.2000 | H |
| ATOM | 438 | C | ALA | 26 | 11.445 | 0.001 | 1.812 | 0.5973 | 1.7000 | C |
| ATOM | 439 | O | ALA | 26 | 10.504 | -0.183 | 1.043 | -0.5679 | 1.5000 | O |
| ATOM | 440 | N | LEU | 27 | 12.564 | -0.694 | 1.659 | -0.4157 | 1.5500 | N |
| ATOM | 441 | H | LEU | 27 | 13.440 | -0.243 | 1.881 | 0.2719 | 1.3000 | H |

| | | | | | | | | | | |
|---|---|---|---|---|---|---|---|---|---|---|
| ATOM | 442 | CA | LEU | 27 | 12.494 | -2.119 | 1.965 | -0.0518 | 1.7000 | C |
| ATOM | 443 | HA | LEU | 27 | 12.062 | -2.596 | 1.086 | 0.0922 | 1.2000 | H |
| ATOM | 444 | CB | LEU | 27 | 13.889 | -2.681 | 2.182 | -0.1102 | 1.7000 | C |
| ATOM | 445 | HB2 | LEU | 27 | 14.486 | -2.488 | 1.288 | 0.0457 | 1.2000 | H |
| ATOM | 446 | HB3 | LEU | 27 | 14.339 | -2.132 | 3.007 | 0.0457 | 1.2000 | H |
| ATOM | 447 | CG | LEU | 27 | 13.941 | -4.151 | 2.513 | 0.3531 | 1.7000 | C |
| ATOM | 448 | HG | LEU | 27 | 13.378 | -4.364 | 3.422 | -0.0361 | 1.2000 | H |
| ATOM | 449 | CD1 | LEU | 27 | 13.365 | -4.948 | 1.367 | -0.4121 | 1.7000 | C |
| ATOM | 450 | HD11 | LEU | 27 | 13.468 | -6.013 | 1.574 | 0.1000 | 1.2000 | H |
| ATOM | 451 | HD12 | LEU | 27 | 12.306 | -4.742 | 1.231 | 0.1000 | 1.2000 | H |
| ATOM | 452 | HD13 | LEU | 27 | 13.893 | -4.720 | 0.440 | 0.1000 | 1.2000 | H |
| ATOM | 453 | CD2 | LEU | 27 | 15.356 | -4.534 | 2.759 | -0.4121 | 1.7000 | C |
| ATOM | 454 | HD21 | LEU | 27 | 15.397 | -5.562 | 3.110 | 0.1000 | 1.2000 | H |
| ATOM | 455 | HD22 | LEU | 27 | 15.945 | -4.436 | 1.846 | 0.1000 | 1.2000 | H |
| ATOM | 456 | HD23 | LEU | 27 | 15.787 | -3.896 | 3.526 | 0.1000 | 1.2000 | H |
| ATOM | 457 | C | LEU | 27 | 11.627 | -2.478 | 3.166 | 0.5973 | 1.7000 | C |
| ATOM | 458 | O | LEU | 27 | 10.832 | -3.402 | 3.053 | -0.5679 | 1.5000 | O |
| ATOM | 459 | N | ALA | 28 | 11.706 | -1.750 | 4.287 | -0.4157 | 1.5500 | N |
| ATOM | 460 | H | ALA | 28 | 12.366 | -0.992 | 4.385 | 0.2719 | 1.3000 | H |
| ATOM | 461 | CA | ALA | 28 | 10.851 | -2.125 | 5.416 | 0.0337 | 1.7000 | C |
| ATOM | 462 | HA | ALA | 28 | 11.125 | -3.124 | 5.760 | 0.0823 | 1.2000 | H |
| ATOM | 463 | CB | ALA | 28 | 11.003 | -1.150 | 6.562 | -0.1825 | 1.7000 | C |
| ATOM | 464 | HB1 | ALA | 28 | 10.394 | -1.477 | 7.405 | 0.0603 | 1.2000 | H |
| ATOM | 465 | HB2 | ALA | 28 | 12.035 | -1.076 | 6.878 | 0.0603 | 1.2000 | H |
| ATOM | 466 | HB3 | ALA | 28 | 10.673 | -0.156 | 6.257 | 0.0603 | 1.2000 | H |
| ATOM | 467 | C | ALA | 28 | 9.391 | -2.120 | 4.993 | 0.5973 | 1.7000 | C |
| ATOM | 468 | O | ALA | 28 | 8.623 | -3.003 | 5.370 | -0.5679 | 1.5000 | O |
| ATOM | 469 | N | VAL | 29 | 9.017 | -1.135 | 4.174 | -0.4157 | 1.5500 | N |
| ATOM | 470 | H | VAL | 29 | 9.685 | -0.435 | 3.888 | 0.2719 | 1.3000 | H |
| ATOM | 471 | CA | VAL | 29 | 7.656 | -1.036 | 3.688 | -0.0875 | 1.7000 | C |
| ATOM | 472 | HA | VAL | 29 | 6.981 | -1.198 | 4.529 | 0.0969 | 1.2000 | H |
| ATOM | 473 | CB | VAL | 29 | 7.354 | 0.338 | 3.111 | 0.2985 | 1.7000 | C |
| ATOM | 474 | HB | VAL | 29 | 8.102 | 0.605 | 2.366 | -0.0297 | 1.2000 | H |
| ATOM | 475 | CG1 | VAL | 29 | 5.995 | 0.319 | 2.452 | -0.3192 | 1.7000 | C |

| | | | | | | | | | | |
|---|---|---|---|---|---|---|---|---|---|---|
| ATOM | 476 | HG11 | VAL | 29 | 5.702 | 1.339 | 2.199 | 0.0791 | 1.2000 | H |
| ATOM | 477 | HG12 | VAL | 29 | 6.021 | -0.249 | 1.522 | 0.0791 | 1.2000 | H |
| ATOM | 478 | HG13 | VAL | 29 | 5.244 | -0.100 | 3.124 | 0.0791 | 1.2000 | H |
| ATOM | 479 | CG2 | VAL | 29 | 7.364 | 1.356 | 4.247 | -0.3192 | 1.7000 | C |
| ATOM | 480 | HG21 | VAL | 29 | 7.208 | 2.357 | 3.842 | 0.0791 | 1.2000 | H |
| ATOM | 481 | HG22 | VAL | 29 | 6.565 | 1.136 | 4.957 | 0.0791 | 1.2000 | H |
| ATOM | 482 | HG23 | VAL | 29 | 8.313 | 1.345 | 4.782 | 0.0791 | 1.2000 | H |
| ATOM | 483 | C | VAL | 29 | 7.366 | -2.095 | 2.659 | 0.5973 | 1.7000 | C |
| ATOM | 484 | O | VAL | 29 | 6.346 | -2.770 | 2.747 | -0.5679 | 1.5000 | O |
| ATOM | 485 | N | ALA | 30 | 8.270 | -2.310 | 1.706 | -0.4157 | 1.5500 | N |
| ATOM | 486 | H | ALA | 30 | 9.117 | -1.764 | 1.639 | 0.2719 | 1.3000 | H |
| ATOM | 487 | CA | ALA | 30 | 7.986 | -3.344 | 0.730 | 0.0337 | 1.7000 | C |
| ATOM | 488 | HA | ALA | 30 | 7.101 | -3.062 | 0.157 | 0.0823 | 1.2000 | H |
| ATOM | 489 | CB | ALA | 30 | 9.156 | -3.490 | -0.221 | -0.1825 | 1.7000 | C |
| ATOM | 490 | HB1 | ALA | 30 | 8.928 | -4.253 | -0.966 | 0.0603 | 1.2000 | H |
| ATOM | 491 | HB2 | ALA | 30 | 9.337 | -2.542 | -0.730 | 0.0603 | 1.2000 | H |
| ATOM | 492 | HB3 | ALA | 30 | 10.054 | -3.781 | 0.324 | 0.0603 | 1.2000 | H |
| ATOM | 493 | C | ALA | 30 | 7.724 | -4.678 | 1.428 | 0.5973 | 1.7000 | C |
| ATOM | 494 | O | ALA | 30 | 6.780 | -5.391 | 1.080 | -0.5679 | 1.5000 | O |
| ATOM | 495 | N | VAL | 31 | 8.501 | -4.988 | 2.464 | -0.4157 | 1.5500 | N |
| ATOM | 496 | H | VAL | 31 | 9.248 | -4.374 | 2.753 | 0.2719 | 1.3000 | H |
| ATOM | 497 | CA | VAL | 31 | 8.288 | -6.232 | 3.178 | -0.0875 | 1.7000 | C |
| ATOM | 498 | HA | VAL | 31 | 8.178 | -7.035 | 2.447 | 0.0969 | 1.2000 | H |
| ATOM | 499 | CB | VAL | 31 | 9.467 | -6.588 | 4.080 | 0.2985 | 1.7000 | C |
| ATOM | 500 | HB | VAL | 31 | 9.682 | -5.755 | 4.743 | -0.0297 | 1.2000 | H |
| ATOM | 501 | CG1 | VAL | 31 | 9.102 | -7.831 | 4.895 | -0.3192 | 1.7000 | C |
| ATOM | 502 | HG11 | VAL | 31 | 9.969 | -8.145 | 5.477 | 0.0791 | 1.2000 | H |
| ATOM | 503 | HG12 | VAL | 31 | 8.290 | -7.641 | 5.592 | 0.0791 | 1.2000 | H |
| ATOM | 504 | HG13 | VAL | 31 | 8.818 | -8.648 | 4.230 | 0.0791 | 1.2000 | H |
| ATOM | 505 | CG2 | VAL | 31 | 10.697 | -6.833 | 3.249 | -0.3192 | 1.7000 | C |
| ATOM | 506 | HG21 | VAL | 31 | 11.539 | -7.060 | 3.903 | 0.0791 | 1.2000 | H |
| ATOM | 507 | HG22 | VAL | 31 | 10.534 | -7.677 | 2.577 | 0.0791 | 1.2000 | H |
| ATOM | 508 | HG23 | VAL | 31 | 10.947 | -5.956 | 2.659 | 0.0791 | 1.2000 | H |
| ATOM | 509 | C | VAL | 31 | 6.989 | -6.158 | 3.981 | 0.5973 | 1.7000 | C |

| | | | | | | | | | | |
|---|---|---|---|---|---|---|---|---|---|---|
| ATOM | 510 | O | VAL | 31 | 6.216 | -7.112 | 3.986 | -0.5679 | 1.5000 | O |
| ATOM | 511 | N | ALA | 32 | 6.722 | -5.024 | 4.635 | -0.4157 | 1.5500 | N |
| ATOM | 512 | H | ALA | 32 | 7.375 | -4.254 | 4.602 | 0.2719 | 1.3000 | H |
| ATOM | 513 | CA | ALA | 32 | 5.492 | -4.864 | 5.403 | 0.0337 | 1.7000 | C |
| ATOM | 514 | HA | ALA | 32 | 5.482 | -5.616 | 6.192 | 0.0823 | 1.2000 | H |
| ATOM | 515 | CB | ALA | 32 | 5.441 | -3.490 | 6.041 | -0.1825 | 1.7000 | C |
| ATOM | 516 | HB1 | ALA | 32 | 4.522 | -3.386 | 6.619 | 0.0603 | 1.2000 | H |
| ATOM | 517 | HB2 | ALA | 32 | 6.291 | -3.361 | 6.712 | 0.0603 | 1.2000 | H |
| ATOM | 518 | HB3 | ALA | 32 | 5.459 | -2.716 | 5.276 | 0.0603 | 1.2000 | H |
| ATOM | 519 | C | ALA | 32 | 4.266 | -5.075 | 4.506 | 0.5973 | 1.7000 | C |
| ATOM | 520 | O | ALA | 32 | 3.216 | -5.536 | 4.967 | -0.5679 | 1.5000 | O |
| ATOM | 521 | N | GLN | 33 | 4.373 | -4.680 | 3.232 | -0.4157 | 1.5500 | N |
| ATOM | 522 | H | GLN | 33 | 5.244 | -4.289 | 2.902 | 0.2719 | 1.3000 | H |
| ATOM | 523 | CA | GLN | 33 | 3.306 | -4.901 | 2.265 | -0.0031 | 1.7000 | C |
| ATOM | 524 | HA | GLN | 33 | 2.360 | -4.573 | 2.698 | 0.0850 | 1.2000 | H |
| ATOM | 525 | CB | GLN | 33 | 3.562 | -4.084 | 0.997 | -0.0036 | 1.7000 | C |
| ATOM | 526 | HB2 | GLN | 33 | 4.559 | -4.312 | 0.625 | 0.0171 | 1.2000 | H |
| ATOM | 527 | HB3 | GLN | 33 | 2.842 | -4.385 | 0.234 | 0.0171 | 1.2000 | H |
| ATOM | 528 | CG | GLN | 33 | 3.427 | -2.579 | 1.214 | -0.0645 | 1.7000 | C |
| ATOM | 529 | HG2 | GLN | 33 | 2.381 | -2.360 | 1.436 | 0.0352 | 1.2000 | H |
| ATOM | 530 | HG3 | GLN | 33 | 4.010 | -2.258 | 2.074 | 0.0352 | 1.2000 | H |
| ATOM | 531 | CD | GLN | 33 | 3.854 | -1.759 | 0.011 | 0.6951 | 1.7000 | C |
| ATOM | 532 | OE1 | GLN | 33 | 4.552 | -2.238 | -0.889 | -0.6086 | 1.5000 | O |
| ATOM | 533 | NE2 | GLN | 33 | 3.437 | -0.496 | -0.010 | -0.9407 | 1.5500 | N |
| ATOM | 534 | HE21 | GLN | 33 | 2.884 | -0.140 | 0.754 | 0.4251 | 1.3000 | H |
| ATOM | 535 | HE22 | GLN | 33 | 3.691 | 0.098 | -0.782 | 0.4251 | 1.3000 | H |
| ATOM | 536 | C | GLN | 33 | 3.196 | -6.402 | 1.964 | 0.5973 | 1.7000 | C |
| ATOM | 537 | O | GLN | 33 | 2.089 | -6.950 | 1.890 | -0.5679 | 1.5000 | O |
| ATOM | 538 | N | VAL | 34 | 4.346 | -7.091 | 1.870 | -0.4157 | 1.5500 | N |
| ATOM | 539 | H | VAL | 34 | 5.234 | -6.614 | 1.956 | 0.2719 | 1.3000 | H |
| ATOM | 540 | CA | VAL | 34 | 4.356 | -8.536 | 1.640 | -0.0875 | 1.7000 | C |
| ATOM | 541 | HA | VAL | 34 | 3.810 | -8.745 | 0.719 | 0.0969 | 1.2000 | H |
| ATOM | 542 | CB | VAL | 34 | 5.770 | -9.134 | 1.518 | 0.2985 | 1.7000 | C |
| ATOM | 543 | HB | VAL | 34 | 6.392 | -8.853 | 2.361 | -0.0297 | 1.2000 | H |

| | | | | | | | | | | |
|---|---|---|---|---|---|---|---|---|---|---|
| ATOM | 544 | CG1 | VAL | 34 | 5.660 | -10.652 | 1.512 | -0.3192 | 1.7000 | C |
| ATOM | 545 | HG11 | VAL | 34 | 6.622 | -11.081 | 1.230 | 0.0791 | 1.2000 | H |
| ATOM | 546 | HG12 | VAL | 34 | 5.421 | -11.039 | 2.503 | 0.0791 | 1.2000 | H |
| ATOM | 547 | HG13 | VAL | 34 | 4.911 | -10.983 | 0.790 | 0.0791 | 1.2000 | H |
| ATOM | 548 | CG2 | VAL | 34 | 6.434 | -8.652 | 0.250 | -0.3192 | 1.7000 | C |
| ATOM | 549 | HG21 | VAL | 34 | 7.477 | -8.971 | 0.237 | 0.0791 | 1.2000 | H |
| ATOM | 550 | HG22 | VAL | 34 | 5.925 | -9.071 | -0.619 | 0.0791 | 1.2000 | H |
| ATOM | 551 | HG23 | VAL | 34 | 6.399 | -7.573 | 0.173 | 0.0791 | 1.2000 | H |
| ATOM | 552 | C | VAL | 34 | 3.652 | -9.238 | 2.775 | 0.5973 | 1.7000 | C |
| ATOM | 553 | O | VAL | 34 | 2.856 | -10.141 | 2.546 | -0.5679 | 1.5000 | O |
| ATOM | 554 | N | CYX | 35 | 3.868 | -8.771 | 4.005 | -0.4157 | 1.5500 | N |
| ATOM | 555 | H | CYX | 35 | 4.517 | -8.007 | 4.131 | 0.2719 | 1.3000 | H |
| ATOM | 556 | CA | CYX | 35 | 3.292 | -9.371 | 5.207 | 0.0429 | 1.7000 | C |
| ATOM | 557 | HA | CYX | 35 | 3.731 | -10.361 | 5.335 | 0.0766 | 1.2000 | H |
| ATOM | 558 | CB | CYX | 35 | 3.618 | -8.533 | 6.440 | -0.0790 | 1.7000 | C |
| ATOM | 559 | HB2 | CYX | 35 | 3.274 | -7.515 | 6.286 | 0.0910 | 1.2000 | H |
| ATOM | 560 | HB3 | CYX | 35 | 3.069 | -8.934 | 7.292 | 0.0910 | 1.2000 | H |
| ATOM | 561 | SG | CYX | 35 | 5.369 | -8.505 | 6.882 | -0.1081 | 1.8000 | S |
| ATOM | 562 | C | CYX | 35 | 1.773 | -9.544 | 5.130 | 0.5973 | 1.7000 | C |
| ATOM | 563 | O | CYX | 35 | 1.188 | -10.269 | 5.925 | -0.5679 | 1.5000 | O |
| ATOM | 564 | N | ARG | 36 | 1.095 | -8.939 | 4.165 | -0.3479 | 1.5500 | N |
| ATOM | 565 | H | ARG | 36 | 1.576 | -8.366 | 3.486 | 0.2747 | 1.3000 | H |
| ATOM | 566 | CA | ARG | 36 | -0.344 | -9.124 | 4.047 | -0.2637 | 1.7000 | C |
| ATOM | 567 | HA | ARG | 36 | -0.816 | -8.869 | 4.997 | 0.1560 | 1.2000 | H |
| ATOM | 568 | CB | ARG | 36 | -0.899 | -8.213 | 2.972 | -0.0007 | 1.7000 | C |
| ATOM | 569 | HB2 | ARG | 36 | -0.350 | -8.400 | 2.047 | 0.0327 | 1.2000 | H |
| ATOM | 570 | HB3 | ARG | 36 | -1.944 | -8.473 | 2.796 | 0.0327 | 1.2000 | H |
| ATOM | 571 | CG | ARG | 36 | -0.842 | -6.730 | 3.325 | 0.0390 | 1.7000 | C |
| ATOM | 572 | HG2 | ARG | 36 | -1.497 | -6.552 | 4.179 | 0.0285 | 1.2000 | H |
| ATOM | 573 | HG3 | ARG | 36 | 0.170 | -6.445 | 3.615 | 0.0285 | 1.2000 | H |
| ATOM | 574 | CD | ARG | 36 | -1.287 | -5.864 | 2.199 | 0.0486 | 1.7000 | C |
| ATOM | 575 | HD2 | ARG | 36 | -2.283 | -6.177 | 1.883 | 0.0687 | 1.2000 | H |
| ATOM | 576 | HD3 | ARG | 36 | -1.342 | -4.834 | 2.555 | 0.0687 | 1.2000 | H |
| ATOM | 577 | NE | ARG | 36 | -0.369 | -5.921 | 1.065 | -0.5295 | 1.5500 | N |

| | | | | | | | | | | |
|---|---|---|---|---|---|---|---|---|---|---|
| ATOM | 578 | HE | ARG | 36 | 0.469 | -6.472 | 1.176 | 0.3456 | 1.3000 | H |
| ATOM | 579 | CZ | ARG | 36 | -0.589 | -5.310 | -0.122 | 0.8076 | 1.7000 | C |
| ATOM | 580 | NH1 | ARG | 36 | -1.688 | -4.604 | -0.297 | -0.8627 | 1.5500 | N |
| ATOM | 581 | HH11 | ARG | 36 | -1.868 | -4.157 | -1.183 | 0.4478 | 1.3000 | H |
| ATOM | 582 | HH12 | ARG | 36 | -2.357 | -4.527 | 0.454 | 0.4478 | 1.3000 | H |
| ATOM | 583 | NH2 | ARG | 36 | 0.285 | -5.411 | -1.112 | -0.8627 | 1.5500 | N |
| ATOM | 584 | HH21 | ARG | 36 | 0.114 | -4.947 | -1.991 | 0.4478 | 1.3000 | H |
| ATOM | 585 | HH22 | ARG | 36 | 1.126 | -5.955 | -0.984 | 0.4478 | 1.3000 | H |
| ATOM | 586 | C | ARG | 36 | -0.698 | -10.594 | 3.729 | 0.7341 | 1.7000 | C |
| ATOM | 587 | O | ARG | 36 | -1.862 | -10.987 | 3.818 | -0.5894 | 1.5000 | O |
| ATOM | 588 | N | VAL | 37 | 0.312 | -11.402 | 3.357 | -0.4157 | 1.5500 | N |
| ATOM | 589 | H | VAL | 37 | 1.244 | -11.019 | 3.285 | 0.2719 | 1.3000 | H |
| ATOM | 590 | CA | VAL | 37 | 0.139 | -12.812 | 3.019 | -0.0875 | 1.7000 | C |
| ATOM | 591 | HA | VAL | 37 | -0.858 | -12.964 | 2.603 | 0.0969 | 1.2000 | H |
| ATOM | 592 | CB | VAL | 37 | 1.180 | -13.270 | 1.980 | 0.2985 | 1.7000 | C |
| ATOM | 593 | HB | VAL | 37 | 0.927 | -14.278 | 1.647 | -0.0297 | 1.2000 | H |
| ATOM | 594 | CG1 | VAL | 37 | 1.101 | -12.378 | 0.749 | -0.3192 | 1.7000 | C |
| ATOM | 595 | HG11 | VAL | 37 | 1.779 | -12.760 | -0.015 | 0.0791 | 1.2000 | H |
| ATOM | 596 | HG12 | VAL | 37 | 0.087 | -12.396 | 0.349 | 0.0791 | 1.2000 | H |
| ATOM | 597 | HG13 | VAL | 37 | 1.376 | -11.350 | 0.969 | 0.0791 | 1.2000 | H |
| ATOM | 598 | CG2 | VAL | 37 | 2.572 | -13.329 | 2.629 | -0.3192 | 1.7000 | C |
| ATOM | 599 | HG21 | VAL | 37 | 3.334 | -13.340 | 1.850 | 0.0791 | 1.2000 | H |
| ATOM | 600 | HG22 | VAL | 37 | 2.756 | -12.484 | 3.288 | 0.0791 | 1.2000 | H |
| ATOM | 601 | HG23 | VAL | 37 | 2.684 | -14.248 | 3.205 | 0.0791 | 1.2000 | H |
| ATOM | 602 | C | VAL | 37 | 0.310 | -13.751 | 4.223 | 0.5973 | 1.7000 | C |
| ATOM | 603 | O | VAL | 37 | 0.247 | -14.970 | 4.051 | -0.5679 | 1.5000 | O |
| ATOM | 604 | N | VAL | 38 | 0.525 | -13.221 | 5.435 | -0.4157 | 1.5500 | N |
| ATOM | 605 | H | VAL | 38 | 0.556 | -12.221 | 5.576 | 0.2719 | 1.3000 | H |
| ATOM | 606 | CA | VAL | 38 | 0.709 | -14.125 | 6.572 | -0.0875 | 1.7000 | C |
| ATOM | 607 | HA | VAL | 38 | 0.860 | -15.158 | 6.265 | 0.0969 | 1.2000 | H |
| ATOM | 608 | CB | VAL | 38 | 1.908 | -13.700 | 7.469 | 0.2985 | 1.7000 | C |
| ATOM | 609 | HB | VAL | 38 | 2.100 | -14.495 | 8.187 | -0.0297 | 1.2000 | H |
| ATOM | 610 | CG1 | VAL | 38 | 3.167 | -13.597 | 6.623 | -0.3192 | 1.7000 | C |
| ATOM | 611 | HG11 | VAL | 38 | 4.032 | -13.451 | 7.271 | 0.0791 | 1.2000 | H |

| | | | | | | | | | | |
|---|---|---|---|---|---|---|---|---|---|---|
| ATOM | 612 | HG12 | VAL | 38 | 3.309 | -14.516 | 6.053 | 0.0791 | 1.2000 | H |
| ATOM | 613 | HG13 | VAL | 38 | 3.101 | -12.752 | 5.937 | 0.0791 | 1.2000 | H |
| ATOM | 614 | CG2 | VAL | 38 | 1.591 | -12.474 | 8.251 | -0.3192 | 1.7000 | C |
| ATOM | 615 | HG21 | VAL | 38 | 2.512 | -11.978 | 8.555 | 0.0791 | 1.2000 | H |
| ATOM | 616 | HG22 | VAL | 38 | 0.972 | -11.779 | 7.700 | 0.0791 | 1.2000 | H |
| ATOM | 617 | HG23 | VAL | 38 | 1.048 | -12.761 | 9.152 | 0.0791 | 1.2000 | H |
| ATOM | 618 | C | VAL | 38 | -0.603 | -13.984 | 7.346 | 0.5973 | 1.7000 | C |
| ATOM | 619 | O | VAL | 38 | -1.372 | -13.084 | 7.032 | -0.5679 | 1.5000 | O |
| ATOM | 620 | N | PRO | 39 | -0.966 | -14.875 | 8.282 | -0.2548 | 1.5500 | N |
| ATOM | 621 | CD | PRO | 39 | -0.142 | -16.053 | 8.620 | 0.0192 | 1.7000 | C |
| ATOM | 622 | HD2 | PRO | 39 | 0.686 | -15.762 | 9.266 | 0.0391 | 1.2000 | H |
| ATOM | 623 | HD3 | PRO | 39 | 0.202 | -16.594 | 7.738 | 0.0391 | 1.2000 | H |
| ATOM | 624 | CG | PRO | 39 | -1.132 | -16.893 | 9.406 | 0.0189 | 1.7000 | C |
| ATOM | 625 | HG2 | PRO | 39 | -0.628 | -17.521 | 10.141 | 0.0213 | 1.2000 | H |
| ATOM | 626 | HG3 | PRO | 39 | -1.713 | -17.510 | 8.718 | 0.0213 | 1.2000 | H |
| ATOM | 627 | CB | PRO | 39 | -2.049 | -15.875 | 10.072 | -0.0070 | 1.7000 | C |
| ATOM | 628 | HB2 | PRO | 39 | -1.554 | -15.477 | 10.960 | 0.0253 | 1.2000 | H |
| ATOM | 629 | HB3 | PRO | 39 | -3.008 | -16.317 | 10.343 | 0.0253 | 1.2000 | H |
| ATOM | 630 | CA | PRO | 39 | -2.217 | -14.782 | 9.014 | -0.0266 | 1.7000 | C |
| ATOM | 631 | HA | PRO | 39 | -3.051 | -15.017 | 8.350 | 0.0641 | 1.2000 | H |
| ATOM | 632 | C | PRO | 39 | -2.369 | -13.406 | 9.632 | 0.5896 | 1.7000 | C |
| ATOM | 633 | O | PRO | 39 | -1.444 | -12.893 | 10.272 | -0.5748 | 1.5000 | O |
| ATOM | 634 | N | LEU | 40 | -3.547 | -12.816 | 9.488 | -0.4157 | 1.5500 | N |
| ATOM | 635 | H | LEU | 40 | -4.296 | -13.256 | 8.974 | 0.2719 | 1.3000 | H |
| ATOM | 636 | CA | LEU | 40 | -3.718 | -11.472 | 9.996 | -0.0518 | 1.7000 | C |
| ATOM | 637 | HA | LEU | 40 | -2.735 | -11.097 | 10.220 | 0.0922 | 1.2000 | H |
| ATOM | 638 | CB | LEU | 40 | -4.275 | -10.577 | 8.884 | -0.1102 | 1.7000 | C |
| ATOM | 639 | HB2 | LEU | 40 | -5.238 | -10.979 | 8.565 | 0.0457 | 1.2000 | H |
| ATOM | 640 | HB3 | LEU | 40 | -4.453 | -9.578 | 9.277 | 0.0457 | 1.2000 | H |
| ATOM | 641 | CG | LEU | 40 | -3.345 | -10.471 | 7.658 | 0.3531 | 1.7000 | C |
| ATOM | 642 | HG | LEU | 40 | -3.221 | -11.455 | 7.214 | -0.0361 | 1.2000 | H |
| ATOM | 643 | CD1 | LEU | 40 | -3.979 | -9.593 | 6.607 | -0.4121 | 1.7000 | C |
| ATOM | 644 | HD11 | LEU | 40 | -3.330 | -9.543 | 5.732 | 0.1000 | 1.2000 | H |
| ATOM | 645 | HD12 | LEU | 40 | -4.939 | -10.012 | 6.304 | 0.1000 | 1.2000 | H |

| | | | | | | | | | | |
|---|---|---|---|---|---|---|---|---|---|---|
| ATOM | 646 | HD13 | LEU | 40 | -4.131 | -8.587 | 6.998 | 0.1000 | 1.2000 | H |
| ATOM | 647 | CD2 | LEU | 40 | -1.963 | -9.926 | 8.106 | -0.4121 | 1.7000 | C |
| ATOM | 648 | HD21 | LEU | 40 | -1.369 | -9.692 | 7.223 | 0.1000 | 1.2000 | H |
| ATOM | 649 | HD22 | LEU | 40 | -2.082 | -9.018 | 8.697 | 0.1000 | 1.2000 | H |
| ATOM | 650 | HD23 | LEU | 40 | -1.409 | -10.675 | 8.668 | 0.1000 | 1.2000 | H |
| ATOM | 651 | C | LEU | 40 | -4.547 | -11.377 | 11.286 | 0.5973 | 1.7000 | C |
| ATOM | 652 | O | LEU | 40 | -5.443 | -12.191 | 11.498 | -0.5679 | 1.5000 | O |
| ATOM | 653 | N | VAL | 41 | -4.237 | -10.398 | 12.163 | -0.4157 | 1.5500 | N |
| ATOM | 654 | H | VAL | 41 | -4.770 | -10.337 | 13.018 | 0.2719 | 1.3000 | H |
| ATOM | 655 | CA | VAL | 41 | -3.169 | -9.408 | 11.957 | -0.0875 | 1.7000 | C |
| ATOM | 656 | HA | VAL | 41 | -2.927 | -9.292 | 10.905 | 0.0969 | 1.2000 | H |
| ATOM | 657 | CB | VAL | 41 | -3.645 | -8.019 | 12.387 | 0.2985 | 1.7000 | C |
| ATOM | 658 | HB | VAL | 41 | -3.879 | -8.008 | 13.453 | -0.0297 | 1.2000 | H |
| ATOM | 659 | CG1 | VAL | 41 | -2.558 | -6.998 | 12.096 | -0.3192 | 1.7000 | C |
| ATOM | 660 | HG11 | VAL | 41 | -3.018 | -6.027 | 11.908 | 0.0791 | 1.2000 | H |
| ATOM | 661 | HG12 | VAL | 41 | -1.896 | -6.820 | 12.941 | 0.0791 | 1.2000 | H |
| ATOM | 662 | HG13 | VAL | 41 | -1.990 | -7.257 | 11.203 | 0.0791 | 1.2000 | H |
| ATOM | 663 | CG2 | VAL | 41 | -4.911 | -7.667 | 11.626 | -0.3192 | 1.7000 | C |
| ATOM | 664 | HG21 | VAL | 41 | -5.241 | -6.668 | 11.914 | 0.0791 | 1.2000 | H |
| ATOM | 665 | HG22 | VAL | 41 | -4.724 | -7.671 | 10.552 | 0.0791 | 1.2000 | H |
| ATOM | 666 | HG23 | VAL | 41 | -5.724 | -8.354 | 11.860 | 0.0791 | 1.2000 | H |
| ATOM | 667 | C | VAL | 41 | -1.872 | -9.735 | 12.697 | 0.5973 | 1.7000 | C |
| ATOM | 668 | O | VAL | 41 | -1.461 | -8.993 | 13.589 | -0.5679 | 1.5000 | O |
| ATOM | 669 | N | ALA | 42 | -1.117 | -10.723 | 12.194 | -0.4157 | 1.5500 | N |
| ATOM | 670 | H | ALA | 42 | -1.460 | -11.286 | 11.428 | 0.2719 | 1.3000 | H |
| ATOM | 671 | CA | ALA | 42 | 0.212 | -11.006 | 12.707 | 0.0337 | 1.7000 | C |
| ATOM | 672 | HA | ALA | 42 | 0.501 | -10.262 | 13.449 | 0.0823 | 1.2000 | H |
| ATOM | 673 | CB | ALA | 42 | 0.209 | -12.351 | 13.401 | -0.1825 | 1.7000 | C |
| ATOM | 674 | HB1 | ALA | 42 | 1.178 | -12.532 | 13.866 | 0.0603 | 1.2000 | H |
| ATOM | 675 | HB2 | ALA | 42 | -0.557 | -12.365 | 14.177 | 0.0603 | 1.2000 | H |
| ATOM | 676 | HB3 | ALA | 42 | 0.004 | -13.145 | 12.682 | 0.0603 | 1.2000 | H |
| ATOM | 677 | C | ALA | 42 | 1.312 | -11.022 | 11.623 | 0.5973 | 1.7000 | C |
| ATOM | 678 | O | ALA | 42 | 2.443 | -10.936 | 12.100 | -0.5679 | 1.5000 | O |
| ATOM | 679 | N | GLY | 43 | 1.084 | -10.296 | 10.509 | -0.4157 | 1.5500 | N |

| | | | | | | | | | | |
|---|---|---|---|---|---|---|---|---|---|---|
| ATOM | 680 | H | GLY | 43 | 1.683 | -10.488 | 9.720 | 0.2719 | 1.3000 | H |
| ATOM | 681 | CA | GLY | 43 | 0.624 | -8.908 | 10.570 | -0.0252 | 1.7000 | C |
| ATOM | 682 | HA2 | GLY | 43 | 0.601 | -8.515 | 9.553 | 0.0698 | 1.2000 | H |
| ATOM | 683 | HA3 | GLY | 43 | -0.372 | -8.790 | 10.969 | 0.0698 | 1.2000 | H |
| ATOM | 684 | C | GLY | 43 | 1.624 | -8.106 | 11.353 | 0.5973 | 1.7000 | C |
| ATOM | 685 | O | GLY | 43 | 2.741 | -7.912 | 10.891 | -0.5679 | 1.5000 | O |
| ATOM | 686 | N | GLY | 44 | 1.229 | -7.646 | 12.538 | -0.4157 | 1.5500 | N |
| ATOM | 687 | H | GLY | 44 | 0.297 | -7.840 | 12.878 | 0.2719 | 1.3000 | H |
| ATOM | 688 | CA | GLY | 44 | 2.109 | -6.832 | 13.365 | -0.0252 | 1.7000 | C |
| ATOM | 689 | HA2 | GLY | 44 | 2.290 | -5.883 | 12.858 | 0.0698 | 1.2000 | H |
| ATOM | 690 | HA3 | GLY | 44 | 1.596 | -6.627 | 14.304 | 0.0698 | 1.2000 | H |
| ATOM | 691 | C | GLY | 44 | 3.467 | -7.491 | 13.692 | 0.5973 | 1.7000 | C |
| ATOM | 692 | O | GLY | 44 | 4.466 | -6.790 | 13.876 | -0.5679 | 1.5000 | O |
| ATOM | 693 | N | ILE | 45 | 3.534 | -8.826 | 13.784 | -0.4157 | 1.5500 | N |
| ATOM | 694 | H | ILE | 45 | 2.727 | -9.409 | 13.613 | 0.2719 | 1.3000 | H |
| ATOM | 695 | CA | ILE | 45 | 4.809 | -9.447 | 14.094 | -0.0597 | 1.7000 | C |
| ATOM | 696 | HA | ILE | 45 | 5.309 | -8.885 | 14.884 | 0.0869 | 1.2000 | H |
| ATOM | 697 | CB | ILE | 45 | 4.655 | -10.895 | 14.564 | 0.1303 | 1.7000 | C |
| ATOM | 698 | HB | ILE | 45 | 4.108 | -11.471 | 13.821 | 0.0187 | 1.2000 | H |
| ATOM | 699 | CG2 | ILE | 45 | 6.039 | -11.517 | 14.710 | -0.3204 | 1.7000 | C |
| ATOM | 700 | HG21 | ILE | 45 | 5.966 | -12.505 | 15.162 | 0.0882 | 1.2000 | H |
| ATOM | 701 | HG22 | ILE | 45 | 6.511 | -11.653 | 13.737 | 0.0882 | 1.2000 | H |
| ATOM | 702 | HG23 | ILE | 45 | 6.674 | -10.891 | 15.339 | 0.0882 | 1.2000 | H |
| ATOM | 703 | CG1 | ILE | 45 | 3.889 | -10.933 | 15.881 | -0.0430 | 1.7000 | C |
| ATOM | 704 | HG12 | ILE | 45 | 4.489 | -10.495 | 16.680 | 0.0236 | 1.2000 | H |
| ATOM | 705 | HG13 | ILE | 45 | 2.979 | -10.341 | 15.785 | 0.0236 | 1.2000 | H |
| ATOM | 706 | CD1 | ILE | 45 | 3.453 | -12.318 | 16.279 | -0.0660 | 1.7000 | C |
| ATOM | 707 | HD11 | ILE | 45 | 2.672 | -12.246 | 17.037 | 0.0186 | 1.2000 | H |
| ATOM | 708 | HD12 | ILE | 45 | 3.057 | -12.858 | 15.419 | 0.0186 | 1.2000 | H |
| ATOM | 709 | HD13 | ILE | 45 | 4.292 | -12.871 | 16.700 | 0.0186 | 1.2000 | H |
| ATOM | 710 | C | ILE | 45 | 5.664 | -9.408 | 12.850 | 0.5973 | 1.7000 | C |
| ATOM | 711 | O | ILE | 45 | 6.796 | -8.935 | 12.876 | -0.5679 | 1.5000 | O |
| ATOM | 712 | N | CYX | 46 | 5.095 | -9.841 | 11.734 | -0.4157 | 1.5500 | N |
| ATOM | 713 | H | CYX | 46 | 4.148 | -10.191 | 11.749 | 0.2719 | 1.3000 | H |

| | | | | | | | | | | |
|---|---|---|---|---|---|---|---|---|---|---|
| ATOM | 714 | CA | CYX | 46 | 5.814 | -9.819 | 10.476 | 0.0429 | 1.7000 | C |
| ATOM | 715 | HA | CYX | 46 | 6.668 | -10.494 | 10.545 | 0.0766 | 1.2000 | H |
| ATOM | 716 | CB | CYX | 46 | 4.909 | -10.294 | 9.343 | -0.0790 | 1.7000 | C |
| ATOM | 717 | HB2 | CYX | 46 | 4.581 | -11.305 | 9.586 | 0.0910 | 1.2000 | H |
| ATOM | 718 | HB3 | CYX | 46 | 4.024 | -9.661 | 9.292 | 0.0910 | 1.2000 | H |
| ATOM | 719 | SG | CYX | 46 | 5.689 | -10.352 | 7.690 | -0.1081 | 1.8000 | S |
| ATOM | 720 | C | CYX | 46 | 6.321 | -8.413 | 10.194 | 0.5973 | 1.7000 | C |
| ATOM | 721 | O | CYX | 46 | 7.471 | -8.224 | 9.805 | -0.5679 | 1.5000 | O |
| ATOM | 722 | N | GLN | 47 | 5.462 | -7.422 | 10.372 | -0.4157 | 1.5500 | N |
| ATOM | 723 | H | GLN | 47 | 4.529 | -7.620 | 10.699 | 0.2719 | 1.3000 | H |
| ATOM | 724 | CA | GLN | 47 | 5.813 | -6.047 | 10.102 | -0.0031 | 1.7000 | C |
| ATOM | 725 | HA | GLN | 47 | 6.222 | -5.998 | 9.092 | 0.0850 | 1.2000 | H |
| ATOM | 726 | CB | GLN | 47 | 4.542 | -5.217 | 10.115 | -0.0036 | 1.7000 | C |
| ATOM | 727 | HB2 | GLN | 47 | 4.005 | -5.387 | 11.050 | 0.0171 | 1.2000 | H |
| ATOM | 728 | HB3 | GLN | 47 | 4.799 | -4.159 | 10.046 | 0.0171 | 1.2000 | H |
| ATOM | 729 | CG | GLN | 47 | 3.666 | -5.587 | 8.919 | -0.0645 | 1.7000 | C |
| ATOM | 730 | HG2 | GLN | 47 | 4.209 | -5.307 | 8.019 | 0.0352 | 1.2000 | H |
| ATOM | 731 | HG3 | GLN | 47 | 3.514 | -6.662 | 8.862 | 0.0352 | 1.2000 | H |
| ATOM | 732 | CD | GLN | 47 | 2.327 | -4.950 | 8.879 | 0.6951 | 1.7000 | C |
| ATOM | 733 | OE1 | GLN | 47 | 1.750 | -4.564 | 9.906 | -0.6086 | 1.5000 | O |
| ATOM | 734 | NE2 | GLN | 47 | 1.786 | -4.831 | 7.662 | -0.9407 | 1.5500 | N |
| ATOM | 735 | HE21 | GLN | 47 | 2.294 | -5.137 | 6.844 | 0.4251 | 1.3000 | H |
| ATOM | 736 | HE22 | GLN | 47 | 0.872 | -4.416 | 7.574 | 0.4251 | 1.3000 | H |
| ATOM | 737 | C | GLN | 47 | 6.902 | -5.514 | 11.045 | 0.5973 | 1.7000 | C |
| ATOM | 738 | O | GLN | 47 | 7.792 | -4.778 | 10.604 | -0.5679 | 1.5000 | O |
| ATOM | 739 | N | CYS | 48 | 6.908 | -5.889 | 12.337 | -0.4157 | 1.5500 | N |
| ATOM | 740 | H | CYS | 48 | 6.202 | -6.493 | 12.735 | 0.2719 | 1.3000 | H |
| ATOM | 741 | CA | CYS | 48 | 8.020 | -5.385 | 13.145 | 0.0213 | 1.7000 | C |
| ATOM | 742 | HA | CYS | 48 | 8.114 | -4.306 | 13.010 | 0.1124 | 1.2000 | H |
| ATOM | 743 | CB | CYS | 48 | 7.836 | -5.664 | 14.641 | -0.1231 | 1.7000 | C |
| ATOM | 744 | HB2 | CYS | 48 | 8.562 | -5.059 | 15.185 | 0.1112 | 1.2000 | H |
| ATOM | 745 | HB3 | CYS | 48 | 6.839 | -5.342 | 14.943 | 0.1112 | 1.2000 | H |
| ATOM | 746 | SG | CYS | 48 | 8.079 | -7.396 | 15.138 | -0.3119 | 1.8000 | S |
| ATOM | 747 | HG | CYS | 48 | 7.001 | -7.872 | 14.506 | 0.1933 | 1.2000 | H |

| | | | | | | | | | | |
|---|---|---|---|---|---|---|---|---|---|---|
| ATOM | 748 | C | CYS | 48 | 9.309 | -6.071 | 12.682 | 0.5973 | 1.7000 | C |
| ATOM | 749 | O | CYS | 48 | 10.398 | -5.486 | 12.744 | -0.5679 | 1.5000 | O |
| ATOM | 750 | N | LEU | 49 | 9.173 | -7.285 | 12.135 | -0.4157 | 1.5500 | N |
| ATOM | 751 | H | LEU | 49 | 8.262 | -7.721 | 12.083 | 0.2719 | 1.3000 | H |
| ATOM | 752 | CA | LEU | 49 | 10.320 | -8.009 | 11.636 | -0.0518 | 1.7000 | C |
| ATOM | 753 | HA | LEU | 49 | 11.154 | -7.878 | 12.323 | 0.0922 | 1.2000 | H |
| ATOM | 754 | CB | LEU | 49 | 10.012 | -9.503 | 11.505 | -0.1102 | 1.7000 | C |
| ATOM | 755 | HB2 | LEU | 49 | 9.148 | -9.639 | 10.861 | 0.0457 | 1.2000 | H |
| ATOM | 756 | HB3 | LEU | 49 | 10.856 | -9.981 | 11.004 | 0.0457 | 1.2000 | H |
| ATOM | 757 | CG | LEU | 49 | 9.755 | -10.235 | 12.835 | 0.3531 | 1.7000 | C |
| ATOM | 758 | HG | LEU | 49 | 8.970 | -9.746 | 13.398 | -0.0361 | 1.2000 | H |
| ATOM | 759 | CD1 | LEU | 49 | 9.321 | -11.651 | 12.546 | -0.4121 | 1.7000 | C |
| ATOM | 760 | HD11 | LEU | 49 | 9.123 | -12.173 | 13.482 | 0.1000 | 1.2000 | H |
| ATOM | 761 | HD12 | LEU | 49 | 8.406 | -11.645 | 11.953 | 0.1000 | 1.2000 | H |
| ATOM | 762 | HD13 | LEU | 49 | 10.101 | -12.182 | 11.999 | 0.1000 | 1.2000 | H |
| ATOM | 763 | CD2 | LEU | 49 | 10.998 | -10.191 | 13.680 | -0.4121 | 1.7000 | C |
| ATOM | 764 | HD21 | LEU | 49 | 10.863 | -10.824 | 14.558 | 0.1000 | 1.2000 | H |
| ATOM | 765 | HD22 | LEU | 49 | 11.858 | -10.554 | 13.116 | 0.1000 | 1.2000 | H |
| ATOM | 766 | HD23 | LEU | 49 | 11.185 | -9.178 | 14.028 | 0.1000 | 1.2000 | H |
| ATOM | 767 | C | LEU | 49 | 10.741 | -7.407 | 10.307 | 0.5973 | 1.7000 | C |
| ATOM | 768 | O | LEU | 49 | 11.924 | -7.385 | 9.993 | -0.5679 | 1.5000 | O |
| ATOM | 769 | N | ALA | 50 | 9.781 | -6.881 | 9.535 | -0.4157 | 1.5500 | N |
| ATOM | 770 | H | ALA | 50 | 8.821 | -6.908 | 9.846 | 0.2719 | 1.3000 | H |
| ATOM | 771 | CA | ALA | 50 | 10.066 | -6.239 | 8.255 | 0.0337 | 1.7000 | C |
| ATOM | 772 | HA | ALA | 50 | 10.528 | -6.964 | 7.585 | 0.0823 | 1.2000 | H |
| ATOM | 773 | CB | ALA | 50 | 8.775 | -5.729 | 7.629 | -0.1825 | 1.7000 | C |
| ATOM | 774 | HB1 | ALA | 50 | 8.982 | -5.300 | 6.657 | 0.0603 | 1.2000 | H |
| ATOM | 775 | HB2 | ALA | 50 | 8.056 | -6.541 | 7.532 | 0.0603 | 1.2000 | H |
| ATOM | 776 | HB3 | ALA | 50 | 8.336 | -4.940 | 8.233 | 0.0603 | 1.2000 | H |
| ATOM | 777 | C | ALA | 50 | 11.009 | -5.068 | 8.457 | 0.5973 | 1.7000 | C |
| ATOM | 778 | O | ALA | 50 | 11.971 | -4.874 | 7.703 | -0.5679 | 1.5000 | O |
| ATOM | 779 | N | GLU | 51 | 10.767 | -4.311 | 9.526 | -0.5163 | 1.5500 | N |
| ATOM | 780 | H | GLU | 51 | 9.970 | -4.491 | 10.121 | 0.2936 | 1.3000 | H |
| ATOM | 781 | CA | GLU | 51 | 11.635 | -3.186 | 9.814 | 0.0397 | 1.7000 | C |

| | | | | | | | | | | |
|---|---|---|---|---|---|---|---|---|---|---|
| ATOM | 782 | HA | GLU | 51 | 11.775 | -2.608 | 8.901 | 0.1105 | 1.2000 | H |
| ATOM | 783 | CB | GLU | 51 | 11.020 | -2.255 | 10.859 | 0.0560 | 1.7000 | C |
| ATOM | 784 | HB2 | GLU | 51 | 10.050 | -1.924 | 10.486 | -0.0173 | 1.2000 | H |
| ATOM | 785 | HB3 | GLU | 51 | 10.860 | -2.807 | 11.786 | -0.0173 | 1.2000 | H |
| ATOM | 786 | CG | GLU | 51 | 11.877 | -1.012 | 11.158 | 0.0136 | 1.7000 | C |
| ATOM | 787 | HG2 | GLU | 51 | 12.832 | -1.329 | 11.581 | -0.0425 | 1.2000 | H |
| ATOM | 788 | HG3 | GLU | 51 | 12.077 | -0.493 | 10.219 | -0.0425 | 1.2000 | H |
| ATOM | 789 | CD | GLU | 51 | 11.230 | -0.033 | 12.126 | 0.8054 | 1.7000 | C |
| ATOM | 790 | OE1 | GLU | 51 | 10.136 | -0.288 | 12.570 | -0.8188 | 1.5000 | O |
| ATOM | 791 | OE2 | GLU | 51 | 11.838 | 0.973 | 12.413 | -0.8188 | 1.5000 | O |
| ATOM | 792 | C | GLU | 51 | 12.997 | -3.675 | 10.273 | 0.5366 | 1.7000 | C |
| ATOM | 793 | O | GLU | 51 | 14.018 | -3.255 | 9.721 | -0.5819 | 1.5000 | O |
| ATOM | 794 | N | ARG | 52 | 13.024 | -4.609 | 11.232 | -0.3479 | 1.5500 | N |
| ATOM | 795 | H | ARG | 52 | 12.159 | -4.943 | 11.635 | 0.2747 | 1.3000 | H |
| ATOM | 796 | CA | ARG | 52 | 14.295 | -5.090 | 11.773 | -0.2637 | 1.7000 | C |
| ATOM | 797 | HA | ARG | 52 | 14.847 | -4.252 | 12.203 | 0.1560 | 1.2000 | H |
| ATOM | 798 | CB | ARG | 52 | 14.009 | -6.115 | 12.856 | -0.0007 | 1.7000 | C |
| ATOM | 799 | HB2 | ARG | 52 | 13.336 | -6.875 | 12.455 | 0.0327 | 1.2000 | H |
| ATOM | 800 | HB3 | ARG | 52 | 14.944 | -6.608 | 13.127 | 0.0327 | 1.2000 | H |
| ATOM | 801 | CG | ARG | 52 | 13.402 | -5.520 | 14.129 | 0.0390 | 1.7000 | C |
| ATOM | 802 | HG2 | ARG | 52 | 14.155 | -4.901 | 14.619 | 0.0285 | 1.2000 | H |
| ATOM | 803 | HG3 | ARG | 52 | 12.560 | -4.879 | 13.875 | 0.0285 | 1.2000 | H |
| ATOM | 804 | CD | ARG | 52 | 12.942 | -6.570 | 15.077 | 0.0486 | 1.7000 | C |
| ATOM | 805 | HD2 | ARG | 52 | 12.227 | -7.206 | 14.555 | 0.0687 | 1.2000 | H |
| ATOM | 806 | HD3 | ARG | 52 | 13.795 | -7.175 | 15.386 | 0.0687 | 1.2000 | H |
| ATOM | 807 | NE | ARG | 52 | 12.303 | -6.003 | 16.253 | -0.5295 | 1.5500 | N |
| ATOM | 808 | HE | ARG | 52 | 12.430 | -5.014 | 16.411 | 0.3456 | 1.3000 | H |
| ATOM | 809 | CZ | ARG | 52 | 11.539 | -6.700 | 17.119 | 0.8076 | 1.7000 | C |
| ATOM | 810 | NH1 | ARG | 52 | 11.334 | -7.985 | 16.934 | -0.8627 | 1.5500 | N |
| ATOM | 811 | HH11 | ARG | 52 | 10.786 | -8.509 | 17.599 | 0.4478 | 1.3000 | H |
| ATOM | 812 | HH12 | ARG | 52 | 11.734 | -8.447 | 16.132 | 0.4478 | 1.3000 | H |
| ATOM | 813 | NH2 | ARG | 52 | 10.993 | -6.092 | 18.157 | -0.8627 | 1.5500 | N |
| ATOM | 814 | HH21 | ARG | 52 | 11.137 | -5.103 | 18.295 | 0.4478 | 1.3000 | H |
| ATOM | 815 | HH22 | ARG | 52 | 10.413 | -6.612 | 18.797 | 0.4478 | 1.3000 | H |

| | | | | | | | | | | |
|---|---|---|---|---|---|---|---|---|---|---|
| ATOM | 816 | C | ARG | 52 | 15.146 | -5.707 | 10.665 | 0.7341 | 1.7000 | C |
| ATOM | 817 | O | ARG | 52 | 16.358 | -5.480 | 10.590 | -0.5894 | 1.5000 | O |
| ATOM | 818 | N | TYR | 53 | 14.488 | -6.440 | 9.769 | -0.4157 | 1.5500 | N |
| ATOM | 819 | H | TYR | 53 | 13.492 | -6.541 | 9.868 | 0.2719 | 1.3000 | H |
| ATOM | 820 | CA | TYR | 53 | 15.114 | -7.050 | 8.617 | -0.0014 | 1.7000 | C |
| ATOM | 821 | HA | TYR | 53 | 15.874 | -7.755 | 8.957 | 0.0876 | 1.2000 | H |
| ATOM | 822 | CB | TYR | 53 | 14.073 | -7.810 | 7.790 | -0.0152 | 1.7000 | C |
| ATOM | 823 | HB2 | TYR | 53 | 13.723 | -8.660 | 8.378 | 0.0295 | 1.2000 | H |
| ATOM | 824 | HB3 | TYR | 53 | 13.212 | -7.166 | 7.618 | 0.0295 | 1.2000 | H |
| ATOM | 825 | CG | TYR | 53 | 14.555 | -8.331 | 6.444 | -0.0011 | 1.7000 | C |
| ATOM | 826 | CD1 | TYR | 53 | 15.324 | -9.479 | 6.360 | -0.1906 | 1.7000 | C |
| ATOM | 827 | HD1 | TYR | 53 | 15.603 | -10.015 | 7.256 | 0.1699 | 1.2000 | H |
| ATOM | 828 | CE1 | TYR | 53 | 15.731 | -9.947 | 5.121 | -0.2341 | 1.7000 | C |
| ATOM | 829 | HE1 | TYR | 53 | 16.331 | -10.842 | 5.051 | 0.1656 | 1.2000 | H |
| ATOM | 830 | CZ | TYR | 53 | 15.361 | -9.265 | 3.965 | 0.3226 | 1.7000 | C |
| ATOM | 831 | OH | TYR | 53 | 15.759 | -9.745 | 2.736 | -0.5579 | 1.5000 | O |
| ATOM | 832 | HH | TYR | 53 | 15.385 | -9.247 | 2.003 | 0.3992 | 1.2000 | H |
| ATOM | 833 | CE2 | TYR | 53 | 14.597 | -8.124 | 4.048 | -0.2341 | 1.7000 | C |
| ATOM | 834 | HE2 | TYR | 53 | 14.299 | -7.608 | 3.148 | 0.1656 | 1.2000 | H |
| ATOM | 835 | CD2 | TYR | 53 | 14.191 | -7.652 | 5.279 | -0.1906 | 1.7000 | C |
| ATOM | 836 | HD2 | TYR | 53 | 13.589 | -6.756 | 5.338 | 0.1699 | 1.2000 | H |
| ATOM | 837 | C | TYR | 53 | 15.775 | -6.008 | 7.760 | 0.5973 | 1.7000 | C |
| ATOM | 838 | O | TYR | 53 | 16.954 | -6.143 | 7.418 | -0.5679 | 1.5000 | O |
| ATOM | 839 | N | SER | 54 | 15.025 | -4.959 | 7.397 | -0.4157 | 1.5500 | N |
| ATOM | 840 | H | SER | 54 | 14.063 | -4.873 | 7.696 | 0.2719 | 1.3000 | H |
| ATOM | 841 | CA | SER | 54 | 15.608 | -3.952 | 6.546 | -0.0249 | 1.7000 | C |
| ATOM | 842 | HA | SER | 54 | 16.006 | -4.438 | 5.656 | 0.0843 | 1.2000 | H |
| ATOM | 843 | CB | SER | 54 | 14.562 | -2.949 | 6.124 | 0.2117 | 1.7000 | C |
| ATOM | 844 | HB2 | SER | 54 | 14.969 | -2.319 | 5.334 | 0.0352 | 1.2000 | H |
| ATOM | 845 | HB3 | SER | 54 | 13.686 | -3.473 | 5.740 | 0.0352 | 1.2000 | H |
| ATOM | 846 | OG | SER | 54 | 14.187 | -2.132 | 7.196 | -0.6546 | 1.5000 | O |
| ATOM | 847 | HG | SER | 54 | 13.934 | -2.683 | 7.944 | 0.4275 | 1.2000 | H |
| ATOM | 848 | C | SER | 54 | 16.731 | -3.203 | 7.235 | 0.5973 | 1.7000 | C |
| ATOM | 849 | O | SER | 54 | 17.718 | -2.856 | 6.605 | -0.5679 | 1.5000 | O |

| | | | | | | | | | | |
|---|---|---|---|---|---|---|---|---|---|---|
| ATOM | 850 | N | VAL | 55 | 16.675 | -3.049 | 8.553 | -0.4157 | 1.5500 | N |
| ATOM | 851 | H | VAL | 55 | 15.882 | -3.362 | 9.095 | 0.2719 | 1.3000 | H |
| ATOM | 852 | CA | VAL | 55 | 17.774 | -2.351 | 9.196 | -0.0875 | 1.7000 | C |
| ATOM | 853 | HA | VAL | 55 | 17.933 | -1.393 | 8.697 | 0.0969 | 1.2000 | H |
| ATOM | 854 | CB | VAL | 55 | 17.479 | -2.078 | 10.681 | 0.2985 | 1.7000 | C |
| ATOM | 855 | HB | VAL | 55 | 17.179 | -3.000 | 11.180 | -0.0297 | 1.2000 | H |
| ATOM | 856 | CG1 | VAL | 55 | 18.733 | -1.535 | 11.361 | -0.3192 | 1.7000 | C |
| ATOM | 857 | HG11 | VAL | 55 | 18.484 | -1.196 | 12.367 | 0.0791 | 1.2000 | H |
| ATOM | 858 | HG12 | VAL | 55 | 19.496 | -2.308 | 11.455 | 0.0791 | 1.2000 | H |
| ATOM | 859 | HG13 | VAL | 55 | 19.138 | -0.692 | 10.799 | 0.0791 | 1.2000 | H |
| ATOM | 860 | CG2 | VAL | 55 | 16.345 | -1.060 | 10.800 | -0.3192 | 1.7000 | C |
| ATOM | 861 | HG21 | VAL | 55 | 16.080 | -0.924 | 11.849 | 0.0791 | 1.2000 | H |
| ATOM | 862 | HG22 | VAL | 55 | 16.664 | -0.101 | 10.390 | 0.0791 | 1.2000 | H |
| ATOM | 863 | HG23 | VAL | 55 | 15.462 | -1.382 | 10.260 | 0.0791 | 1.2000 | H |
| ATOM | 864 | C | VAL | 55 | 19.042 | -3.166 | 9.069 | 0.5973 | 1.7000 | C |
| ATOM | 865 | O | VAL | 55 | 20.081 | -2.667 | 8.636 | -0.5679 | 1.5000 | O |
| ATOM | 866 | N | ILE | 56 | 18.951 | -4.452 | 9.371 | -0.4157 | 1.5500 | N |
| ATOM | 867 | H | ILE | 56 | 18.078 | -4.845 | 9.694 | 0.2719 | 1.3000 | H |
| ATOM | 868 | CA | ILE | 56 | 20.126 | -5.288 | 9.296 | -0.0597 | 1.7000 | C |
| ATOM | 869 | HA | ILE | 56 | 20.915 | -4.847 | 9.909 | 0.0869 | 1.2000 | H |
| ATOM | 870 | CB | ILE | 56 | 19.806 | -6.687 | 9.837 | 0.1303 | 1.7000 | C |
| ATOM | 871 | HB | ILE | 56 | 18.920 | -7.068 | 9.324 | 0.0187 | 1.2000 | H |
| ATOM | 872 | CG2 | ILE | 56 | 20.968 | -7.637 | 9.561 | -0.3204 | 1.7000 | C |
| ATOM | 873 | HG21 | ILE | 56 | 20.795 | -8.595 | 10.050 | 0.0882 | 1.2000 | H |
| ATOM | 874 | HG22 | ILE | 56 | 21.067 | -7.839 | 8.494 | 0.0882 | 1.2000 | H |
| ATOM | 875 | HG23 | ILE | 56 | 21.900 | -7.213 | 9.940 | 0.0882 | 1.2000 | H |
| ATOM | 876 | CG1 | ILE | 56 | 19.512 | -6.600 | 11.341 | -0.0430 | 1.7000 | C |
| ATOM | 877 | HG12 | ILE | 56 | 20.427 | -6.369 | 11.889 | 0.0236 | 1.2000 | H |
| ATOM | 878 | HG13 | ILE | 56 | 18.808 | -5.792 | 11.532 | 0.0236 | 1.2000 | H |
| ATOM | 879 | CD1 | ILE | 56 | 18.893 | -7.852 | 11.904 | -0.0660 | 1.7000 | C |
| ATOM | 880 | HD11 | ILE | 56 | 18.366 | -7.611 | 12.828 | 0.0186 | 1.2000 | H |
| ATOM | 881 | HD12 | ILE | 56 | 18.181 | -8.280 | 11.197 | 0.0186 | 1.2000 | H |
| ATOM | 882 | HD13 | ILE | 56 | 19.669 | -8.585 | 12.127 | 0.0186 | 1.2000 | H |
| ATOM | 883 | C | ILE | 56 | 20.636 | -5.390 | 7.865 | 0.5973 | 1.7000 | C |

| | | | | | | | | | | |
|---|---|---|---|---|---|---|---|---|---|---|
| ATOM | 884 | O | ILE | 56 | 21.825 | -5.172 | 7.597 | -0.5679 | 1.5000 | O |
| ATOM | 885 | N | LEU | 57 | 19.737 | -5.694 | 6.928 | -0.4157 | 1.5500 | N |
| ATOM | 886 | H | LEU | 57 | 18.769 | -5.833 | 7.176 | 0.2719 | 1.3000 | H |
| ATOM | 887 | CA | LEU | 57 | 20.151 | -5.863 | 5.550 | -0.0518 | 1.7000 | C |
| ATOM | 888 | HA | LEU | 57 | 20.968 | -6.584 | 5.557 | 0.0922 | 1.2000 | H |
| ATOM | 889 | CB | LEU | 57 | 19.022 | -6.416 | 4.686 | -0.1102 | 1.7000 | C |
| ATOM | 890 | HB2 | LEU | 57 | 18.719 | -7.386 | 5.084 | 0.0457 | 1.2000 | H |
| ATOM | 891 | HB3 | LEU | 57 | 18.164 | -5.745 | 4.758 | 0.0457 | 1.2000 | H |
| ATOM | 892 | CG | LEU | 57 | 19.408 | -6.563 | 3.212 | 0.3531 | 1.7000 | C |
| ATOM | 893 | HG | LEU | 57 | 19.678 | -5.599 | 2.784 | -0.0361 | 1.2000 | H |
| ATOM | 894 | CD1 | LEU | 57 | 20.592 | -7.518 | 3.099 | -0.4121 | 1.7000 | C |
| ATOM | 895 | HD11 | LEU | 57 | 20.794 | -7.719 | 2.047 | 0.1000 | 1.2000 | H |
| ATOM | 896 | HD12 | LEU | 57 | 21.496 | -7.090 | 3.527 | 0.1000 | 1.2000 | H |
| ATOM | 897 | HD13 | LEU | 57 | 20.367 | -8.461 | 3.598 | 0.1000 | 1.2000 | H |
| ATOM | 898 | CD2 | LEU | 57 | 18.239 | -7.073 | 2.428 | -0.4121 | 1.7000 | C |
| ATOM | 899 | HD21 | LEU | 57 | 18.511 | -7.161 | 1.376 | 0.1000 | 1.2000 | H |
| ATOM | 900 | HD22 | LEU | 57 | 17.932 | -8.050 | 2.801 | 0.1000 | 1.2000 | H |
| ATOM | 901 | HD23 | LEU | 57 | 17.410 | -6.378 | 2.512 | 0.1000 | 1.2000 | H |
| ATOM | 902 | C | LEU | 57 | 20.649 | -4.593 | 4.902 | 0.5973 | 1.7000 | C |
| ATOM | 903 | O | LEU | 57 | 21.701 | -4.597 | 4.269 | -0.5679 | 1.5000 | O |
| ATOM | 904 | N | LEU | 58 | 19.908 | -3.502 | 5.024 | -0.4157 | 1.5500 | N |
| ATOM | 905 | H | LEU | 58 | 19.062 | -3.510 | 5.572 | 0.2719 | 1.3000 | H |
| ATOM | 906 | CA | LEU | 58 | 20.304 | -2.303 | 4.322 | -0.0518 | 1.7000 | C |
| ATOM | 907 | HA | LEU | 58 | 20.456 | -2.533 | 3.268 | 0.0922 | 1.2000 | H |
| ATOM | 908 | CB | LEU | 58 | 19.217 | -1.248 | 4.453 | -0.1102 | 1.7000 | C |
| ATOM | 909 | HB2 | LEU | 58 | 19.052 | -1.041 | 5.511 | 0.0457 | 1.2000 | H |
| ATOM | 910 | HB3 | LEU | 58 | 19.590 | -0.323 | 4.027 | 0.0457 | 1.2000 | H |
| ATOM | 911 | CG | LEU | 58 | 17.859 | -1.604 | 3.776 | 0.3531 | 1.7000 | C |
| ATOM | 912 | HG | LEU | 58 | 17.439 | -2.492 | 4.238 | -0.0361 | 1.2000 | H |
| ATOM | 913 | CD1 | LEU | 58 | 16.884 | -0.453 | 4.028 | -0.4121 | 1.7000 | C |
| ATOM | 914 | HD11 | LEU | 58 | 15.925 | -0.663 | 3.562 | 0.1000 | 1.2000 | H |
| ATOM | 915 | HD12 | LEU | 58 | 16.733 | -0.328 | 5.101 | 0.1000 | 1.2000 | H |
| ATOM | 916 | HD13 | LEU | 58 | 17.277 | 0.476 | 3.615 | 0.1000 | 1.2000 | H |
| ATOM | 917 | CD2 | LEU | 58 | 18.043 | -1.939 | 2.325 | -0.4121 | 1.7000 | C |

| | | | | | | | | | | |
|---|---|---|---|---|---|---|---|---|---|---|
| ATOM | 918 | HD21 | LEU | 58 | 17.074 | -1.980 | 1.827 | 0.1000 | 1.2000 | H |
| ATOM | 919 | HD22 | LEU | 58 | 18.656 | -1.184 | 1.852 | 0.1000 | 1.2000 | H |
| ATOM | 920 | HD23 | LEU | 58 | 18.522 | -2.912 | 2.221 | 0.1000 | 1.2000 | H |
| ATOM | 921 | C | LEU | 58 | 21.625 | -1.774 | 4.869 | 0.5973 | 1.7000 | C |
| ATOM | 922 | O | LEU | 58 | 22.497 | -1.350 | 4.100 | -0.5679 | 1.5000 | O |
| ATOM | 923 | N | ASP | 59 | 21.827 | -1.859 | 6.197 | -0.5163 | 1.5500 | N |
| ATOM | 924 | H | ASP | 59 | 21.105 | -2.196 | 6.820 | 0.2936 | 1.3000 | H |
| ATOM | 925 | CA | ASP | 59 | 23.096 | -1.412 | 6.751 | 0.0381 | 1.7000 | C |
| ATOM | 926 | HA | ASP | 59 | 23.306 | -0.403 | 6.392 | 0.0880 | 1.2000 | H |
| ATOM | 927 | CB | ASP | 59 | 23.074 | -1.381 | 8.281 | -0.0303 | 1.7000 | C |
| ATOM | 928 | HB2 | ASP | 59 | 22.717 | -2.339 | 8.662 | -0.0122 | 1.2000 | H |
| ATOM | 929 | HB3 | ASP | 59 | 24.096 | -1.242 | 8.638 | -0.0122 | 1.2000 | H |
| ATOM | 930 | CG | ASP | 59 | 22.224 | -0.236 | 8.861 | 0.7994 | 1.7000 | C |
| ATOM | 931 | OD1 | ASP | 59 | 21.845 | 0.642 | 8.113 | -0.8014 | 1.5000 | O |
| ATOM | 932 | OD2 | ASP | 59 | 22.001 | -0.227 | 10.054 | -0.8014 | 1.5000 | O |
| ATOM | 933 | C | ASP | 59 | 24.218 | -2.319 | 6.265 | 0.5366 | 1.7000 | C |
| ATOM | 934 | O | ASP | 59 | 25.313 | -1.839 | 5.951 | -0.5819 | 1.5000 | O |
| ATOM | 935 | N | THR | 60 | 23.938 | -3.627 | 6.163 | -0.4157 | 1.5500 | N |
| ATOM | 936 | H | THR | 60 | 23.024 | -3.968 | 6.428 | 0.2719 | 1.3000 | H |
| ATOM | 937 | CA | THR | 60 | 24.925 | -4.575 | 5.684 | -0.0389 | 1.7000 | C |
| ATOM | 938 | HA | THR | 60 | 25.817 | -4.495 | 6.308 | 0.1007 | 1.2000 | H |
| ATOM | 939 | CB | THR | 60 | 24.411 | -6.027 | 5.749 | 0.3654 | 1.7000 | C |
| ATOM | 940 | HB | THR | 60 | 23.482 | -6.115 | 5.189 | 0.0043 | 1.2000 | H |
| ATOM | 941 | CG2 | THR | 60 | 25.435 | -6.978 | 5.124 | -0.2438 | 1.7000 | C |
| ATOM | 942 | HG21 | THR | 60 | 25.151 | -8.006 | 5.348 | 0.0642 | 1.2000 | H |
| ATOM | 943 | HG22 | THR | 60 | 25.458 | -6.866 | 4.040 | 0.0642 | 1.2000 | H |
| ATOM | 944 | HG23 | THR | 60 | 26.427 | -6.789 | 5.536 | 0.0642 | 1.2000 | H |
| ATOM | 945 | OG1 | THR | 60 | 24.146 | -6.390 | 7.114 | -0.6761 | 1.5000 | O |
| ATOM | 946 | HG1 | THR | 60 | 23.408 | -5.855 | 7.431 | 0.4102 | 1.2000 | H |
| ATOM | 947 | C | THR | 60 | 25.310 | -4.275 | 4.255 | 0.5973 | 1.7000 | C |
| ATOM | 948 | O | THR | 60 | 26.493 | -4.248 | 3.925 | -0.5679 | 1.5000 | O |
| ATOM | 949 | N | LEU | 61 | 24.329 | -4.015 | 3.401 | -0.4157 | 1.5500 | N |
| ATOM | 950 | H | LEU | 61 | 23.369 | -3.990 | 3.713 | 0.2719 | 1.3000 | H |
| ATOM | 951 | CA | LEU | 61 | 24.633 | -3.771 | 2.007 | -0.0518 | 1.7000 | C |

| | | | | | | | | | | |
|---|---|---|---|---|---|---|---|---|---|---|
| ATOM | 952 | HA | LEU | 61 | 25.190 | -4.618 | 1.610 | 0.0922 | 1.2000 | H |
| ATOM | 953 | CB | LEU | 61 | 23.321 | -3.589 | 1.232 | -0.1102 | 1.7000 | C |
| ATOM | 954 | HB2 | LEU | 61 | 22.739 | -2.826 | 1.748 | 0.0457 | 1.2000 | H |
| ATOM | 955 | HB3 | LEU | 61 | 23.548 | -3.205 | 0.236 | 0.0457 | 1.2000 | H |
| ATOM | 956 | CG | LEU | 61 | 22.448 | -4.846 | 1.089 | 0.3531 | 1.7000 | C |
| ATOM | 957 | HG | LEU | 61 | 22.268 | -5.291 | 2.059 | -0.0361 | 1.2000 | H |
| ATOM | 958 | CD1 | LEU | 61 | 21.109 | -4.462 | 0.474 | -0.4121 | 1.7000 | C |
| ATOM | 959 | HD11 | LEU | 61 | 20.476 | -5.345 | 0.387 | 0.1000 | 1.2000 | H |
| ATOM | 960 | HD12 | LEU | 61 | 20.606 | -3.734 | 1.110 | 0.1000 | 1.2000 | H |
| ATOM | 961 | HD13 | LEU | 61 | 21.257 | -4.032 | -0.517 | 0.1000 | 1.2000 | H |
| ATOM | 962 | CD2 | LEU | 61 | 23.154 | -5.868 | 0.257 | -0.4121 | 1.7000 | C |
| ATOM | 963 | HD21 | LEU | 61 | 24.071 | -6.194 | 0.742 | 0.1000 | 1.2000 | H |
| ATOM | 964 | HD22 | LEU | 61 | 22.511 | -6.740 | 0.137 | 0.1000 | 1.2000 | H |
| ATOM | 965 | HD23 | LEU | 61 | 23.379 | -5.464 | -0.730 | 0.1000 | 1.2000 | H |
| ATOM | 966 | C | LEU | 61 | 25.503 | -2.529 | 1.837 | 0.5973 | 1.7000 | C |
| ATOM | 967 | O | LEU | 61 | 26.486 | -2.550 | 1.083 | -0.5679 | 1.5000 | O |
| ATOM | 968 | N | LEU | 62 | 25.196 | -1.470 | 2.591 | -0.4157 | 1.5500 | N |
| ATOM | 969 | H | LEU | 62 | 24.403 | -1.494 | 3.219 | 0.2719 | 1.3000 | H |
| ATOM | 970 | CA | LEU | 62 | 25.979 | -0.247 | 2.484 | -0.0518 | 1.7000 | C |
| ATOM | 971 | HA | LEU | 62 | 26.145 | -0.030 | 1.430 | 0.0922 | 1.2000 | H |
| ATOM | 972 | CB | LEU | 62 | 25.216 | 0.926 | 3.114 | -0.1102 | 1.7000 | C |
| ATOM | 973 | HB2 | LEU | 62 | 24.939 | 0.643 | 4.131 | 0.0457 | 1.2000 | H |
| ATOM | 974 | HB3 | LEU | 62 | 25.896 | 1.776 | 3.192 | 0.0457 | 1.2000 | H |
| ATOM | 975 | CG | LEU | 62 | 23.945 | 1.390 | 2.368 | 0.3531 | 1.7000 | C |
| ATOM | 976 | HG | LEU | 62 | 23.269 | 0.547 | 2.228 | -0.0361 | 1.2000 | H |
| ATOM | 977 | CD1 | LEU | 62 | 23.233 | 2.453 | 3.191 | -0.4121 | 1.7000 | C |
| ATOM | 978 | HD11 | LEU | 62 | 22.413 | 2.882 | 2.620 | 0.1000 | 1.2000 | H |
| ATOM | 979 | HD12 | LEU | 62 | 22.839 | 2.007 | 4.106 | 0.1000 | 1.2000 | H |
| ATOM | 980 | HD13 | LEU | 62 | 23.926 | 3.253 | 3.452 | 0.1000 | 1.2000 | H |
| ATOM | 981 | CD2 | LEU | 62 | 24.339 | 1.940 | 1.005 | -0.4121 | 1.7000 | C |
| ATOM | 982 | HD21 | LEU | 62 | 23.470 | 2.371 | 0.511 | 0.1000 | 1.2000 | H |
| ATOM | 983 | HD22 | LEU | 62 | 25.098 | 2.716 | 1.114 | 0.1000 | 1.2000 | H |
| ATOM | 984 | HD23 | LEU | 62 | 24.723 | 1.145 | 0.374 | 0.1000 | 1.2000 | H |
| ATOM | 985 | C | LEU | 62 | 27.364 | -0.363 | 3.125 | 0.5973 | 1.7000 | C |

| | | | | | | | | | | |
|---|---|---|---|---|---|---|---|---|---|---|
| ATOM | 986 | O | LEU | 62 | 28.340 | 0.175 | 2.597 | -0.5679 | 1.5000 | O |
| ATOM | 987 | N | GLY | 63 | 27.464 | -1.064 | 4.257 | -0.4157 | 1.5500 | N |
| ATOM | 988 | H | GLY | 63 | 26.642 | -1.481 | 4.670 | 0.2719 | 1.3000 | H |
| ATOM | 989 | CA | GLY | 63 | 28.740 | -1.194 | 4.952 | -0.0252 | 1.7000 | C |
| ATOM | 990 | HA2 | GLY | 63 | 29.252 | -0.231 | 4.970 | 0.0698 | 1.2000 | H |
| ATOM | 991 | HA3 | GLY | 63 | 28.532 | -1.488 | 5.981 | 0.0698 | 1.2000 | H |
| ATOM | 992 | C | GLY | 63 | 29.662 | -2.245 | 4.333 | 0.5973 | 1.7000 | C |
| ATOM | 993 | O | GLY | 63 | 30.886 | -2.119 | 4.393 | -0.5679 | 1.5000 | O |
| ATOM | 994 | N | ARG | 64 | 29.079 | -3.271 | 3.720 | -0.3479 | 1.5500 | N |
| ATOM | 995 | H | ARG | 64 | 28.072 | -3.311 | 3.677 | 0.2747 | 1.3000 | H |
| ATOM | 996 | CA | ARG | 64 | 29.836 | -4.372 | 3.146 | -0.2637 | 1.7000 | C |
| ATOM | 997 | HA | ARG | 64 | 30.674 | -4.601 | 3.807 | 0.1560 | 1.2000 | H |
| ATOM | 998 | CB | ARG | 64 | 28.962 | -5.620 | 3.045 | -0.0007 | 1.7000 | C |
| ATOM | 999 | HB2 | ARG | 64 | 28.597 | -5.857 | 4.046 | 0.0327 | 1.2000 | H |
| ATOM | 1000 | HB3 | ARG | 64 | 28.107 | -5.390 | 2.409 | 0.0327 | 1.2000 | H |
| ATOM | 1001 | CG | ARG | 64 | 29.635 | -6.850 | 2.486 | 0.0390 | 1.7000 | C |
| ATOM | 1002 | HG2 | ARG | 64 | 29.901 | -6.682 | 1.442 | 0.0285 | 1.2000 | H |
| ATOM | 1003 | HG3 | ARG | 64 | 30.547 | -7.043 | 3.052 | 0.0285 | 1.2000 | H |
| ATOM | 1004 | CD | ARG | 64 | 28.762 | -8.063 | 2.565 | 0.0486 | 1.7000 | C |
| ATOM | 1005 | HD2 | ARG | 64 | 29.334 | -8.924 | 2.216 | 0.0687 | 1.2000 | H |
| ATOM | 1006 | HD3 | ARG | 64 | 28.492 | -8.229 | 3.609 | 0.0687 | 1.2000 | H |
| ATOM | 1007 | NE | ARG | 64 | 27.558 | -7.946 | 1.761 | -0.5295 | 1.5500 | N |
| ATOM | 1008 | HE | ARG | 64 | 27.501 | -7.168 | 1.121 | 0.3456 | 1.3000 | H |
| ATOM | 1009 | CZ | ARG | 64 | 26.518 | -8.802 | 1.825 | 0.8076 | 1.7000 | C |
| ATOM | 1010 | NH1 | ARG | 64 | 26.558 | -9.823 | 2.652 | -0.8627 | 1.5500 | N |
| ATOM | 1011 | HH11 | ARG | 64 | 27.370 | -9.967 | 3.233 | 0.4478 | 1.3000 | H |
| ATOM | 1012 | HH12 | ARG | 64 | 25.775 | -10.455 | 2.715 | 0.4478 | 1.3000 | H |
| ATOM | 1013 | NH2 | ARG | 64 | 25.460 | -8.625 | 1.060 | -0.8627 | 1.5500 | N |
| ATOM | 1014 | HH21 | ARG | 64 | 25.422 | -7.843 | 0.424 | 0.4478 | 1.3000 | H |
| ATOM | 1015 | HH22 | ARG | 64 | 24.689 | -9.273 | 1.110 | 0.4478 | 1.3000 | H |
| ATOM | 1016 | C | ARG | 64 | 30.398 | -4.052 | 1.775 | 0.7341 | 1.7000 | C |
| ATOM | 1017 | O | ARG | 64 | 31.569 | -4.312 | 1.503 | -0.5894 | 1.5000 | O |
| ATOM | 1018 | N | MET | 65 | 29.578 | -3.502 | 0.889 | -0.4157 | 1.5500 | N |
| ATOM | 1019 | H | MET | 65 | 28.627 | -3.263 | 1.133 | 0.2719 | 1.3000 | H |

| | | | | | | | | | | |
|---|---|---|---|---|---|---|---|---|---|---|
| ATOM | 1020 | CA | MET | 65 | 30.054 | -3.280 | -0.463 | -0.0237 | 1.7000 | C |
| ATOM | 1021 | HA | MET | 65 | 30.724 | -4.097 | -0.714 | 0.0880 | 1.2000 | H |
| ATOM | 1022 | CB | MET | 65 | 28.852 | -3.384 | -1.391 | 0.0342 | 1.7000 | C |
| ATOM | 1023 | HB2 | MET | 65 | 28.083 | -2.712 | -1.030 | 0.0241 | 1.2000 | H |
| ATOM | 1024 | HB3 | MET | 65 | 29.125 | -3.092 | -2.403 | 0.0241 | 1.2000 | H |
| ATOM | 1025 | CG | MET | 65 | 28.290 | -4.725 | -1.429 | 0.0018 | 1.7000 | C |
| ATOM | 1026 | HG2 | MET | 65 | 29.068 | -5.395 | -1.792 | 0.0440 | 1.2000 | H |
| ATOM | 1027 | HG3 | MET | 65 | 27.999 | -5.040 | -0.427 | 0.0440 | 1.2000 | H |
| ATOM | 1028 | SD | MET | 65 | 26.881 | -4.898 | -2.496 | -0.2737 | 1.8000 | S |
| ATOM | 1029 | CE | MET | 65 | 25.665 | -3.952 | -1.663 | -0.0536 | 1.7000 | C |
| ATOM | 1030 | HE1 | MET | 65 | 24.714 | -4.042 | -2.188 | 0.0684 | 1.2000 | H |
| ATOM | 1031 | HE2 | MET | 65 | 25.556 | -4.326 | -0.647 | 0.0684 | 1.2000 | H |
| ATOM | 1032 | HE3 | MET | 65 | 25.955 | -2.902 | -1.642 | 0.0684 | 1.2000 | H |
| ATOM | 1033 | C | MET | 65 | 30.796 | -1.938 | -0.631 | 0.5973 | 1.7000 | C |
| ATOM | 1034 | O | MET | 65 | 30.124 | -1.037 | -1.139 | -0.5679 | 1.5000 | O |
| ATOM | 1035 | N | LEU | 66 | 32.097 | -2.018 | -1.000 | -0.4157 | 1.5500 | N |
| ATOM | 1036 | H | LEU | 66 | 32.615 | -1.164 | -0.852 | 0.2719 | 1.3000 | H |
| ATOM | 1037 | CA | LEU | 66 | 32.732 | -2.893 | -2.021 | -0.0518 | 1.7000 | C |
| ATOM | 1038 | HA | LEU | 66 | 33.366 | -2.258 | -2.635 | 0.0922 | 1.2000 | H |
| ATOM | 1039 | CB | LEU | 66 | 33.618 | -3.923 | -1.319 | -0.1102 | 1.7000 | C |
| ATOM | 1040 | HB2 | LEU | 66 | 33.003 | -4.499 | -0.628 | 0.0457 | 1.2000 | H |
| ATOM | 1041 | HB3 | LEU | 66 | 34.014 | -4.613 | -2.067 | 0.0457 | 1.2000 | H |
| ATOM | 1042 | CG | LEU | 66 | 34.795 | -3.382 | -0.565 | 0.3531 | 1.7000 | C |
| ATOM | 1043 | HG | LEU | 66 | 34.470 | -2.625 | 0.150 | -0.0361 | 1.2000 | H |
| ATOM | 1044 | CD1 | LEU | 66 | 35.453 | -4.517 | 0.200 | -0.4121 | 1.7000 | C |
| ATOM | 1045 | HD11 | LEU | 66 | 36.310 | -4.135 | 0.756 | 0.1000 | 1.2000 | H |
| ATOM | 1046 | HD12 | LEU | 66 | 34.746 | -4.940 | 0.914 | 0.1000 | 1.2000 | H |
| ATOM | 1047 | HD13 | LEU | 66 | 35.791 | -5.297 | -0.483 | 0.1000 | 1.2000 | H |
| ATOM | 1048 | CD2 | LEU | 66 | 35.754 | -2.765 | -1.544 | -0.4121 | 1.7000 | C |
| ATOM | 1049 | HD21 | LEU | 66 | 36.650 | -2.446 | -1.010 | 0.1000 | 1.2000 | H |
| ATOM | 1050 | HD22 | LEU | 66 | 36.047 | -3.495 | -2.298 | 0.1000 | 1.2000 | H |
| ATOM | 1051 | HD23 | LEU | 66 | 35.329 | -1.881 | -2.016 | 0.1000 | 1.2000 | H |
| ATOM | 1052 | C | LEU | 66 | 31.844 | -3.670 | -2.991 | 0.5973 | 1.7000 | C |
| ATOM | 1053 | O | LEU | 66 | 31.674 | -4.876 | -2.795 | -0.5679 | 1.5000 | O |

| | | | | | | | | | | |
|---|---|---|---|---|---|---|---|---|---|---|
| ATOM | 1054 | N | PRO | 67 | 31.271 | -3.063 | -4.033 | -0.2548 | 1.5500 | N |
| ATOM | 1055 | CD | PRO | 67 | 31.402 | -1.614 | -4.251 | 0.0192 | 1.7000 | C |
| ATOM | 1056 | HD2 | PRO | 67 | 32.395 | -1.376 | -4.630 | 0.0391 | 1.2000 | H |
| ATOM | 1057 | HD3 | PRO | 67 | 31.170 | -1.026 | -3.367 | 0.0391 | 1.2000 | H |
| ATOM | 1058 | CG | PRO | 67 | 30.355 | -1.381 | -5.329 | 0.0189 | 1.7000 | C |
| ATOM | 1059 | HG2 | PRO | 67 | 30.617 | -0.536 | -5.966 | 0.0213 | 1.2000 | H |
| ATOM | 1060 | HG3 | PRO | 67 | 29.384 | -1.214 | -4.861 | 0.0213 | 1.2000 | H |
| ATOM | 1061 | CB | PRO | 67 | 30.331 | -2.679 | -6.105 | -0.0070 | 1.7000 | C |
| ATOM | 1062 | HB2 | PRO | 67 | 31.185 | -2.697 | -6.783 | 0.0253 | 1.2000 | H |
| ATOM | 1063 | HB3 | PRO | 67 | 29.406 | -2.795 | -6.666 | 0.0253 | 1.2000 | H |
| ATOM | 1064 | CA | PRO | 67 | 30.479 | -3.757 | -5.027 | -0.0266 | 1.7000 | C |
| ATOM | 1065 | HA | PRO | 67 | 29.512 | -4.021 | -4.603 | 0.0641 | 1.2000 | H |
| ATOM | 1066 | C | PRO | 67 | 31.198 | -4.998 | -5.552 | 0.5896 | 1.7000 | C |
| ATOM | 1067 | O | PRO | 67 | 30.576 | -6.040 | -5.775 | -0.5748 | 1.5000 | O |
| ATOM | 1068 | N | GLN | 68 | 32.526 | -4.903 | -5.705 | -0.4157 | 1.5500 | N |
| ATOM | 1069 | H | GLN | 68 | 32.978 | -4.057 | -5.418 | 0.2719 | 1.3000 | H |
| ATOM | 1070 | CA | GLN | 68 | 33.324 | -6.030 | -6.172 | -0.0031 | 1.7000 | C |
| ATOM | 1071 | HA | GLN | 68 | 32.864 | -6.406 | -7.087 | 0.0850 | 1.2000 | H |
| ATOM | 1072 | CB | GLN | 68 | 34.741 | -5.570 | -6.534 | -0.0036 | 1.7000 | C |
| ATOM | 1073 | HB2 | GLN | 68 | 35.288 | -6.449 | -6.878 | 0.0171 | 1.2000 | H |
| ATOM | 1074 | HB3 | GLN | 68 | 34.700 | -4.883 | -7.378 | 0.0171 | 1.2000 | H |
| ATOM | 1075 | CG | GLN | 68 | 35.547 | -4.938 | -5.395 | -0.0645 | 1.7000 | C |
| ATOM | 1076 | HG2 | GLN | 68 | 35.329 | -5.405 | -4.436 | 0.0352 | 1.2000 | H |
| ATOM | 1077 | HG3 | GLN | 68 | 36.605 | -5.106 | -5.606 | 0.0352 | 1.2000 | H |
| ATOM | 1078 | CD | GLN | 68 | 35.309 | -3.437 | -5.307 | 0.6951 | 1.7000 | C |
| ATOM | 1079 | OE1 | GLN | 68 | 34.225 | -2.934 | -5.616 | -0.6086 | 1.5000 | O |
| ATOM | 1080 | NE2 | GLN | 68 | 36.337 | -2.707 | -4.891 | -0.9407 | 1.5500 | N |
| ATOM | 1081 | HE21 | GLN | 68 | 37.205 | -3.152 | -4.632 | 0.4251 | 1.3000 | H |
| ATOM | 1082 | HE22 | GLN | 68 | 36.231 | -1.708 | -4.792 | 0.4251 | 1.3000 | H |
| ATOM | 1083 | C | GLN | 68 | 33.378 | -7.192 | -5.186 | 0.5973 | 1.7000 | C |
| ATOM | 1084 | O | GLN | 68 | 33.589 | -8.336 | -5.584 | -0.5679 | 1.5000 | O |
| ATOM | 1085 | N | LEU | 69 | 33.172 | -6.923 | -3.898 | -0.4157 | 1.5500 | N |
| ATOM | 1086 | H | LEU | 69 | 32.986 | -5.981 | -3.587 | 0.2719 | 1.3000 | H |
| ATOM | 1087 | CA | LEU | 69 | 33.204 | -7.995 | -2.927 | -0.0518 | 1.7000 | C |

| | | | | | | | | | | |
|---|---|---|---|---|---|---|---|---|---|---|
| ATOM | 1088 | HA | LEU | 69 | 34.112 | -8.582 | -3.075 | 0.0922 | 1.2000 | H |
| ATOM | 1089 | CB | LEU | 69 | 33.172 | -7.460 | -1.497 | -0.1102 | 1.7000 | C |
| ATOM | 1090 | HB2 | LEU | 69 | 34.070 | -6.862 | -1.342 | 0.0457 | 1.2000 | H |
| ATOM | 1091 | HB3 | LEU | 69 | 32.311 | -6.804 | -1.383 | 0.0457 | 1.2000 | H |
| ATOM | 1092 | CG | LEU | 69 | 33.094 | -8.504 | -0.413 | 0.3531 | 1.7000 | C |
| ATOM | 1093 | HG | LEU | 69 | 32.192 | -9.100 | -0.540 | -0.0361 | 1.2000 | H |
| ATOM | 1094 | CD1 | LEU | 69 | 34.301 | -9.411 | -0.481 | -0.4121 | 1.7000 | C |
| ATOM | 1095 | HD11 | LEU | 69 | 34.290 | -10.093 | 0.370 | 0.1000 | 1.2000 | H |
| ATOM | 1096 | HD12 | LEU | 69 | 34.278 | -10.016 | -1.387 | 0.1000 | 1.2000 | H |
| ATOM | 1097 | HD13 | LEU | 69 | 35.222 | -8.827 | -0.451 | 0.1000 | 1.2000 | H |
| ATOM | 1098 | CD2 | LEU | 69 | 32.993 | -7.804 | 0.924 | -0.4121 | 1.7000 | C |
| ATOM | 1099 | HD21 | LEU | 69 | 32.840 | -8.540 | 1.714 | 0.1000 | 1.2000 | H |
| ATOM | 1100 | HD22 | LEU | 69 | 33.910 | -7.251 | 1.131 | 0.1000 | 1.2000 | H |
| ATOM | 1101 | HD23 | LEU | 69 | 32.154 | -7.110 | 0.926 | 0.1000 | 1.2000 | H |
| ATOM | 1102 | C | LEU | 69 | 32.012 | -8.857 | -3.147 | 0.5973 | 1.7000 | C |
| ATOM | 1103 | O | LEU | 69 | 32.126 | -10.065 | -3.352 | -0.5679 | 1.5000 | O |
| ATOM | 1104 | N | VAL | 70 | 30.855 | -8.222 | -3.159 | -0.4157 | 1.5500 | N |
| ATOM | 1105 | H | VAL | 70 | 30.800 | -7.224 | -3.008 | 0.2719 | 1.3000 | H |
| ATOM | 1106 | CA | VAL | 70 | 29.652 | -8.999 | -3.334 | -0.0875 | 1.7000 | C |
| ATOM | 1107 | HA | VAL | 70 | 29.694 | -9.818 | -2.615 | 0.0969 | 1.2000 | H |
| ATOM | 1108 | CB | VAL | 70 | 28.405 | -8.211 | -3.019 | 0.2985 | 1.7000 | C |
| ATOM | 1109 | HB | VAL | 70 | 28.432 | -7.238 | -3.510 | -0.0297 | 1.2000 | H |
| ATOM | 1110 | CG1 | VAL | 70 | 27.174 | -8.977 | -3.454 | -0.3192 | 1.7000 | C |
| ATOM | 1111 | HG11 | VAL | 70 | 26.281 | -8.469 | -3.087 | 0.0791 | 1.2000 | H |
| ATOM | 1112 | HG12 | VAL | 70 | 27.098 | -9.008 | -4.541 | 0.0791 | 1.2000 | H |
| ATOM | 1113 | HG13 | VAL | 70 | 27.188 | -9.993 | -3.055 | 0.0791 | 1.2000 | H |
| ATOM | 1114 | CG2 | VAL | 70 | 28.361 | -8.054 | -1.502 | -0.3192 | 1.7000 | C |
| ATOM | 1115 | HG21 | VAL | 70 | 27.473 | -7.491 | -1.217 | 0.0791 | 1.2000 | H |
| ATOM | 1116 | HG22 | VAL | 70 | 28.316 | -9.031 | -1.018 | 0.0791 | 1.2000 | H |
| ATOM | 1117 | HG23 | VAL | 70 | 29.243 | -7.526 | -1.139 | 0.0791 | 1.2000 | H |
| ATOM | 1118 | C | VAL | 70 | 29.591 | -9.656 | -4.678 | 0.5973 | 1.7000 | C |
| ATOM | 1119 | O | VAL | 70 | 29.242 | -10.830 | -4.744 | -0.5679 | 1.5000 | O |
| ATOM | 1120 | N | CYX | 71 | 29.963 | -8.958 | -5.750 | -0.4157 | 1.5500 | N |
| ATOM | 1121 | H | CYX | 71 | 30.252 | -7.992 | -5.689 | 0.2719 | 1.3000 | H |

| | | | | | | | | | | |
|---|---|---|---|---|---|---|---|---|---|---|
| ATOM | 1122 | CA | CYX | 71 | 29.918 | -9.635 | -7.027 | 0.0429 | 1.7000 | C |
| ATOM | 1123 | HA | CYX | 71 | 28.896 | -9.961 | -7.222 | 0.0766 | 1.2000 | H |
| ATOM | 1124 | CB | CYX | 71 | 30.349 | -8.753 | -8.164 | -0.0790 | 1.7000 | C |
| ATOM | 1125 | HB2 | CYX | 71 | 31.326 | -8.322 | -7.944 | 0.0910 | 1.2000 | H |
| ATOM | 1126 | HB3 | CYX | 71 | 30.451 | -9.398 | -9.035 | 0.0910 | 1.2000 | H |
| ATOM | 1127 | SG | CYX | 71 | 29.253 | -7.468 | -8.619 | -0.1081 | 1.8000 | S |
| ATOM | 1128 | C | CYX | 71 | 30.825 | -10.861 | -7.022 | 0.5973 | 1.7000 | C |
| ATOM | 1129 | O | CYX | 71 | 30.462 | -11.904 | -7.565 | -0.5679 | 1.5000 | O |
| ATOM | 1130 | N | ARG | 72 | 32.007 | -10.781 | -6.408 | -0.3479 | 1.5500 | N |
| ATOM | 1131 | H | ARG | 72 | 32.306 | -9.936 | -5.940 | 0.2747 | 1.3000 | H |
| ATOM | 1132 | CA | ARG | 72 | 32.849 | -11.967 | -6.380 | -0.2637 | 1.7000 | C |
| ATOM | 1133 | HA | ARG | 72 | 32.946 | -12.358 | -7.391 | 0.1560 | 1.2000 | H |
| ATOM | 1134 | CB | ARG | 72 | 34.240 | -11.612 | -5.880 | -0.0007 | 1.7000 | C |
| ATOM | 1135 | HB2 | ARG | 72 | 34.156 | -11.063 | -4.941 | 0.0327 | 1.2000 | H |
| ATOM | 1136 | HB3 | ARG | 72 | 34.786 | -12.536 | -5.679 | 0.0327 | 1.2000 | H |
| ATOM | 1137 | CG | ARG | 72 | 35.055 | -10.790 | -6.885 | 0.0390 | 1.7000 | C |
| ATOM | 1138 | HG2 | ARG | 72 | 35.260 | -11.419 | -7.747 | 0.0285 | 1.2000 | H |
| ATOM | 1139 | HG3 | ARG | 72 | 34.482 | -9.932 | -7.231 | 0.0285 | 1.2000 | H |
| ATOM | 1140 | CD | ARG | 72 | 36.349 | -10.339 | -6.361 | 0.0486 | 1.7000 | C |
| ATOM | 1141 | HD2 | ARG | 72 | 36.176 | -9.720 | -5.479 | 0.0687 | 1.2000 | H |
| ATOM | 1142 | HD3 | ARG | 72 | 36.942 | -11.209 | -6.075 | 0.0687 | 1.2000 | H |
| ATOM | 1143 | NE | ARG | 72 | 37.078 | -9.563 | -7.360 | -0.5295 | 1.5500 | N |
| ATOM | 1144 | HE | ARG | 72 | 36.676 | -9.514 | -8.285 | 0.3456 | 1.3000 | H |
| ATOM | 1145 | CZ | ARG | 72 | 38.248 | -8.939 | -7.143 | 0.8076 | 1.7000 | C |
| ATOM | 1146 | NH1 | ARG | 72 | 38.819 | -9.006 | -5.959 | -0.8627 | 1.5500 | N |
| ATOM | 1147 | HH11 | ARG | 72 | 39.705 | -8.552 | -5.799 | 0.4478 | 1.3000 | H |
| ATOM | 1148 | HH12 | ARG | 72 | 38.389 | -9.551 | -5.228 | 0.4478 | 1.3000 | H |
| ATOM | 1149 | NH2 | ARG | 72 | 38.827 | -8.257 | -8.115 | -0.8627 | 1.5500 | N |
| ATOM | 1150 | HH21 | ARG | 72 | 39.727 | -7.826 | -7.965 | 0.4478 | 1.3000 | H |
| ATOM | 1151 | HH22 | ARG | 72 | 38.411 | -8.236 | -9.035 | 0.4478 | 1.3000 | H |
| ATOM | 1152 | C | ARG | 72 | 32.215 | -13.082 | -5.535 | 0.7341 | 1.7000 | C |
| ATOM | 1153 | O | ARG | 72 | 32.220 | -14.248 | -5.934 | -0.5894 | 1.5000 | O |
| ATOM | 1154 | N | LEU | 73 | 31.576 | -12.718 | -4.419 | -0.4157 | 1.5500 | N |
| ATOM | 1155 | H | LEU | 73 | 31.555 | -11.745 | -4.151 | 0.2719 | 1.3000 | H |

| | | | | | | | | | | |
|---|---|---|---|---|---|---|---|---|---|---|
| ATOM | 1156 | CA | LEU | 73 | 30.948 | -13.694 | -3.524 | -0.0518 | 1.7000 | C |
| ATOM | 1157 | HA | LEU | 73 | 31.697 | -14.432 | -3.238 | 0.0922 | 1.2000 | H |
| ATOM | 1158 | CB | LEU | 73 | 30.413 | -12.995 | -2.260 | -0.1102 | 1.7000 | C |
| ATOM | 1159 | HB2 | LEU | 73 | 29.761 | -12.179 | -2.568 | 0.0457 | 1.2000 | H |
| ATOM | 1160 | HB3 | LEU | 73 | 29.790 | -13.706 | -1.713 | 0.0457 | 1.2000 | H |
| ATOM | 1161 | CG | LEU | 73 | 31.485 | -12.435 | -1.274 | 0.3531 | 1.7000 | C |
| ATOM | 1162 | HG | LEU | 73 | 32.196 | -11.805 | -1.790 | -0.0361 | 1.2000 | H |
| ATOM | 1163 | CD1 | LEU | 73 | 30.796 | -11.610 | -0.174 | -0.4121 | 1.7000 | C |
| ATOM | 1164 | HD11 | LEU | 73 | 31.543 | -11.239 | 0.529 | 0.1000 | 1.2000 | H |
| ATOM | 1165 | HD12 | LEU | 73 | 30.278 | -10.761 | -0.617 | 0.1000 | 1.2000 | H |
| ATOM | 1166 | HD13 | LEU | 73 | 30.077 | -12.229 | 0.364 | 0.1000 | 1.2000 | H |
| ATOM | 1167 | CD2 | LEU | 73 | 32.264 | -13.581 | -0.685 | -0.4121 | 1.7000 | C |
| ATOM | 1168 | HD21 | LEU | 73 | 32.955 | -13.202 | 0.069 | 0.1000 | 1.2000 | H |
| ATOM | 1169 | HD22 | LEU | 73 | 31.591 | -14.299 | -0.215 | 0.1000 | 1.2000 | H |
| ATOM | 1170 | HD23 | LEU | 73 | 32.852 | -14.082 | -1.453 | 0.1000 | 1.2000 | H |
| ATOM | 1171 | C | LEU | 73 | 29.809 | -14.460 | -4.202 | 0.5973 | 1.7000 | C |
| ATOM | 1172 | O | LEU | 73 | 29.588 | -15.638 | -3.911 | -0.5679 | 1.5000 | O |
| ATOM | 1173 | N | VAL | 74 | 29.106 | -13.801 | -5.123 | -0.4157 | 1.5500 | N |
| ATOM | 1174 | H | VAL | 74 | 29.333 | -12.839 | -5.329 | 0.2719 | 1.3000 | H |
| ATOM | 1175 | CA | VAL | 74 | 27.981 | -14.420 | -5.824 | -0.0875 | 1.7000 | C |
| ATOM | 1176 | HA | VAL | 74 | 27.633 | -15.296 | -5.275 | 0.0969 | 1.2000 | H |
| ATOM | 1177 | CB | VAL | 74 | 26.805 | -13.445 | -5.957 | 0.2985 | 1.7000 | C |
| ATOM | 1178 | HB | VAL | 74 | 25.970 | -13.962 | -6.433 | -0.0297 | 1.2000 | H |
| ATOM | 1179 | CG1 | VAL | 74 | 26.334 | -12.984 | -4.588 | -0.3192 | 1.7000 | C |
| ATOM | 1180 | HG11 | VAL | 74 | 25.460 | -12.342 | -4.700 | 0.0791 | 1.2000 | H |
| ATOM | 1181 | HG12 | VAL | 74 | 26.057 | -13.850 | -3.985 | 0.0791 | 1.2000 | H |
| ATOM | 1182 | HG13 | VAL | 74 | 27.113 | -12.427 | -4.067 | 0.0791 | 1.2000 | H |
| ATOM | 1183 | CG2 | VAL | 74 | 27.218 | -12.322 | -6.803 | -0.3192 | 1.7000 | C |
| ATOM | 1184 | HG21 | VAL | 74 | 26.381 | -12.090 | -7.456 | 0.0791 | 1.2000 | H |
| ATOM | 1185 | HG22 | VAL | 74 | 27.396 | -11.431 | -6.208 | 0.0791 | 1.2000 | H |
| ATOM | 1186 | HG23 | VAL | 74 | 28.077 | -12.494 | -7.439 | 0.0791 | 1.2000 | H |
| ATOM | 1187 | C | VAL | 74 | 28.362 | -14.874 | -7.239 | 0.5973 | 1.7000 | C |
| ATOM | 1188 | O | VAL | 74 | 27.484 | -15.183 | -8.047 | -0.5679 | 1.5000 | O |
| ATOM | 1189 | N | LEU | 75 | 29.665 | -14.886 | -7.545 | -0.4157 | 1.5500 | N |

| | | | | | | | | | | |
|---|---|---|---|---|---|---|---|---|---|---|
| ATOM | 1190 | H | LEU | 75 | 30.329 | -14.621 | -6.831 | 0.2719 | 1.3000 | H |
| ATOM | 1191 | CA | LEU | 75 | 30.214 | -15.312 | -8.834 | -0.0518 | 1.7000 | C |
| ATOM | 1192 | HA | LEU | 75 | 31.297 | -15.218 | -8.768 | 0.0922 | 1.2000 | H |
| ATOM | 1193 | CB | LEU | 75 | 29.854 | -16.783 | -9.079 | -0.1102 | 1.7000 | C |
| ATOM | 1194 | HB2 | LEU | 75 | 28.785 | -16.888 | -9.260 | 0.0457 | 1.2000 | H |
| ATOM | 1195 | HB3 | LEU | 75 | 30.361 | -17.113 | -9.988 | 0.0457 | 1.2000 | H |
| ATOM | 1196 | CG | LEU | 75 | 30.270 | -17.744 | -7.950 | 0.3531 | 1.7000 | C |
| ATOM | 1197 | HG | LEU | 75 | 29.751 | -17.481 | -7.028 | -0.0361 | 1.2000 | H |
| ATOM | 1198 | CD1 | LEU | 75 | 29.859 | -19.155 | -8.317 | -0.4121 | 1.7000 | C |
| ATOM | 1199 | HD11 | LEU | 75 | 30.112 | -19.834 | -7.502 | 0.1000 | 1.2000 | H |
| ATOM | 1200 | HD12 | LEU | 75 | 28.781 | -19.195 | -8.480 | 0.1000 | 1.2000 | H |
| ATOM | 1201 | HD13 | LEU | 75 | 30.373 | -19.477 | -9.223 | 0.1000 | 1.2000 | H |
| ATOM | 1202 | CD2 | LEU | 75 | 31.762 | -17.627 | -7.717 | -0.4121 | 1.7000 | C |
| ATOM | 1203 | HD21 | LEU | 75 | 32.080 | -18.400 | -7.017 | 0.1000 | 1.2000 | H |
| ATOM | 1204 | HD22 | LEU | 75 | 32.307 | -17.754 | -8.653 | 0.1000 | 1.2000 | H |
| ATOM | 1205 | HD23 | LEU | 75 | 32.010 | -16.663 | -7.274 | 0.1000 | 1.2000 | H |
| ATOM | 1206 | C | LEU | 75 | 29.793 | -14.479 | -10.059 | 0.5973 | 1.7000 | C |
| ATOM | 1207 | O | LEU | 75 | 29.614 | -15.021 | -11.152 | -0.5679 | 1.5000 | O |
| ATOM | 1208 | N | ARG | 76 | 29.673 | -13.158 | -9.889 | -0.3479 | 1.5500 | N |
| ATOM | 1209 | H | ARG | 76 | 29.797 | -12.775 | -8.966 | 0.2747 | 1.3000 | H |
| ATOM | 1210 | CA | ARG | 76 | 29.368 | -12.238 | -10.987 | -0.2637 | 1.7000 | C |
| ATOM | 1211 | HA | ARG | 76 | 29.151 | -12.802 | -11.894 | 0.1560 | 1.2000 | H |
| ATOM | 1212 | CB | ARG | 76 | 28.166 | -11.355 | -10.679 | -0.0007 | 1.7000 | C |
| ATOM | 1213 | HB2 | ARG | 76 | 28.356 | -10.825 | -9.744 | 0.0327 | 1.2000 | H |
| ATOM | 1214 | HB3 | ARG | 76 | 28.071 | -10.609 | -11.468 | 0.0327 | 1.2000 | H |
| ATOM | 1215 | CG | ARG | 76 | 26.815 | -12.064 | -10.592 | 0.0390 | 1.7000 | C |
| ATOM | 1216 | HG2 | ARG | 76 | 26.670 | -12.676 | -11.482 | 0.0285 | 1.2000 | H |
| ATOM | 1217 | HG3 | ARG | 76 | 26.802 | -12.726 | -9.726 | 0.0285 | 1.2000 | H |
| ATOM | 1218 | CD | ARG | 76 | 25.681 | -11.087 | -10.501 | 0.0486 | 1.7000 | C |
| ATOM | 1219 | HD2 | ARG | 76 | 25.704 | -10.440 | -11.380 | 0.0687 | 1.2000 | H |
| ATOM | 1220 | HD3 | ARG | 76 | 24.745 | -11.647 | -10.511 | 0.0687 | 1.2000 | H |
| ATOM | 1221 | NE | ARG | 76 | 25.727 | -10.268 | -9.288 | -0.5295 | 1.5500 | N |
| ATOM | 1222 | HE | ARG | 76 | 26.449 | -10.488 | -8.618 | 0.3456 | 1.3000 | H |
| ATOM | 1223 | CZ | ARG | 76 | 24.891 | -9.238 | -9.029 | 0.8076 | 1.7000 | C |

| | | | | | | | | | | |
|---|---|---|---|---|---|---|---|---|---|---|
| ATOM | 1224 | NH1 | ARG | 76 | 23.943 | -8.938 | -9.890 | -0.8627 | 1.5500 | N |
| ATOM | 1225 | HH11 | ARG | 76 | 23.853 | -9.480 | -10.737 | 0.4478 | 1.3000 | H |
| ATOM | 1226 | HH12 | ARG | 76 | 23.292 | -8.194 | -9.692 | 0.4478 | 1.3000 | H |
| ATOM | 1227 | NH2 | ARG | 76 | 25.016 | -8.526 | -7.920 | -0.8627 | 1.5500 | N |
| ATOM | 1228 | HH21 | ARG | 76 | 25.743 | -8.748 | -7.257 | 0.4478 | 1.3000 | H |
| ATOM | 1229 | HH22 | ARG | 76 | 24.378 | -7.766 | -7.738 | 0.4478 | 1.3000 | H |
| ATOM | 1230 | C | ARG | 76 | 30.559 | -11.334 | -11.305 | 0.7341 | 1.7000 | C |
| ATOM | 1231 | O | ARG | 76 | 30.443 | -10.383 | -12.073 | -0.5894 | 1.5000 | O |
| ATOM | 1232 | N | CYX | 77 | 31.706 | -11.650 | -10.718 | -0.4157 | 1.5500 | N |
| ATOM | 1233 | H | CYX | 77 | 31.737 | -12.453 | -10.106 | 0.2719 | 1.3000 | H |
| ATOM | 1234 | CA | CYX | 77 | 32.949 | -10.901 | -10.899 | 0.0429 | 1.7000 | C |
| ATOM | 1235 | HA | CYX | 77 | 33.056 | -10.664 | -11.958 | 0.0766 | 1.2000 | H |
| ATOM | 1236 | CB | CYX | 77 | 32.962 | -9.599 | -10.119 | -0.0790 | 1.7000 | C |
| ATOM | 1237 | HB2 | CYX | 77 | 32.086 | -9.015 | -10.397 | 0.0910 | 1.2000 | H |
| ATOM | 1238 | HB3 | CYX | 77 | 32.905 | -9.829 | -9.055 | 0.0910 | 1.2000 | H |
| ATOM | 1239 | SG | CYX | 77 | 34.405 | -8.591 | -10.406 | -0.1081 | 1.8000 | S |
| ATOM | 1240 | C | CYX | 77 | 34.142 | -11.728 | -10.472 | 0.5973 | 1.7000 | C |
| ATOM | 1241 | O | CYX | 77 | 34.039 | -12.547 | -9.561 | -0.5679 | 1.5000 | O |
| ATOM | 1242 | N | SER | 78 | 35.269 | -11.542 | -11.141 | -0.4157 | 1.5500 | N |
| ATOM | 1243 | H | SER | 78 | 35.317 | -10.882 | -11.905 | 0.2719 | 1.3000 | H |
| ATOM | 1244 | CA | SER | 78 | 36.491 | -12.210 | -10.732 | -0.0249 | 1.7000 | C |
| ATOM | 1245 | HA | SER | 78 | 36.487 | -12.376 | -9.654 | 0.0843 | 1.2000 | H |
| ATOM | 1246 | CB | SER | 78 | 36.625 | -13.553 | -11.431 | 0.2117 | 1.7000 | C |
| ATOM | 1247 | HB2 | SER | 78 | 37.474 | -14.094 | -11.013 | 0.0352 | 1.2000 | H |
| ATOM | 1248 | HB3 | SER | 78 | 35.722 | -14.143 | -11.271 | 0.0352 | 1.2000 | H |
| ATOM | 1249 | OG | SER | 78 | 36.833 | -13.377 | -12.811 | -0.6546 | 1.5000 | O |
| ATOM | 1250 | HG | SER | 78 | 37.668 | -12.912 | -12.944 | 0.4275 | 1.2000 | H |
| ATOM | 1251 | C | SER | 78 | 37.693 | -11.354 | -11.084 | 0.5973 | 1.7000 | C |
| ATOM | 1252 | O | SER | 78 | 37.609 | -10.494 | -11.959 | -0.5679 | 1.5000 | O |
| ATOM | 1253 | N | MET | 79 | 38.777 | -11.598 | -10.361 | -0.3821 | 1.5500 | N |
| ATOM | 1254 | H | MET | 79 | 38.720 | -12.367 | -9.717 | 0.2681 | 1.3000 | H |
| ATOM | 1255 | CA | MET | 79 | 40.101 | -11.032 | -10.571 | -0.2597 | 1.7000 | C |
| ATOM | 1256 | HA | MET | 79 | 40.153 | -10.087 | -10.032 | 0.1277 | 1.2000 | H |
| ATOM | 1257 | CB | MET | 79 | 41.114 | -11.979 | -9.959 | -0.0236 | 1.7000 | C |

```
ATOM   1258  HB2 MET    79      40.995 -12.983 -10.369  0.0480  1.2000       H
ATOM   1259  HB3 MET    79      42.115 -11.624 -10.209  0.0480  1.2000       H
ATOM   1260  CG  MET    79      40.992 -12.016  -8.441  0.0492  1.7000       C
ATOM   1261  HG2 MET    79      41.834 -12.579  -8.038  0.0317  1.2000       H
ATOM   1262  HG3 MET    79      41.031 -11.001  -8.045  0.0317  1.2000       H
ATOM   1263  SD  MET    79      39.450 -12.820  -7.903 -0.2692  1.8000       S
ATOM   1264  CE  MET    79      39.633 -12.787  -6.127 -0.0376  1.7000       C
ATOM   1265  HE1 MET    79      38.750 -13.226  -5.663  0.0625  1.2000       H
ATOM   1266  HE2 MET    79      40.514 -13.361  -5.841  0.0625  1.2000       H
ATOM   1267  HE3 MET    79      39.746 -11.757  -5.790  0.0625  1.2000       H
ATOM   1268  C   MET    79      40.395 -10.713 -12.032  0.8013  1.7000       C
ATOM   1269  O   MET    79      41.008  -9.686 -12.324 -0.8105  1.5000       O
ATOM   1270  OXT MET    79      40.395 -11.660 -12.820 -0.8105  1.5000       O
TER    1271      MET    79
END
```
